# Gene Evolutionary Trajectories in *M. tuberculosis* Reveal Temporal Signs of Selection

**DOI:** 10.1101/2021.07.15.452434

**Authors:** Álvaro Chiner-Oms, Mariana G. López, Iñaki Comas

## Abstract

Genetic differences between different *Mycobacterium tuberculosis* complex (MTBC) strains determine their ability to transmit within different host populations, their latency times, and their drug-resistance profiles. Said differences usually emerge through *de novo* mutations and are maintained or discarded by the balance of evolutionary forces. Using a dataset of ~5,000 strains representing global MTBC diversity, we determined the past and present selective forces that have shaped the current variability observed in the pathogen population. We identified regions that have evolved under changing types of selection since the time of the MTBC common ancestor. Our approach highlighted striking differences in the genome regions relevant for host-pathogen interaction and, in particular, suggested an adaptive role for the sensor protein of two-component systems. In addition, we applied our approach to successfully identify potential determinants of resistance to drugs administered as second-line tuberculosis treatments.

## Introduction

The *Mycobacterium tuberculosis* complex (MTBC) is a genetically monomorphic group of bacteria ^1,2^ whose members cause tuberculosis in humans and animals. The MTBC comprises both human-associated (L1, L2, L3, L4, L5, L6, L7, L8, and L9) and animal-associated (A1, A2, A3, and A4) clades ^3–7^. Due to the absence of mobile genetic elements and measurable recombination among strains and other species ^8–10^, chromosomal mutations represent the source of MTBC genetic diversity. The maximum genetic distance between any two MTBC strains is around 2,500 single nucleotide polymorphisms (SNPs). Strikingly, studies have highlighted large phenotypic differences between strains involving traits like gene expression, drug resistance, transmissibility, and immune response despite this limited variation. In some cases, the mutations driving phenotypic differences have been identified - for example, non-synonymous variants in genes such as *rpoB, katG*, or *embB* cause drug-resistant phenotypes ^11–13^. Furthermore, single mutations in regulatory elements can induce alterations to downstream gene expression, which can foster differential virulence characteristics ^14,15^. Finally, specific gene mutations may affect transmission ^9^, host tropism within the complex ^16^, and the host immune response ^17^. However, many of the genomic determinants of these phenotypes remain elusive despite robust evidence that they are driven by genetic differences between strains ^18,19^.

Several types of evolutionary forces play crucial roles in the fixation of mutations in bacterial populations. Previous research has provided evidence for the ongoing positive selection of specific genes and regions ^9,20–23^, while other studies have reported ongoing purifying selection of specific genomic regions, especially in epitopes and essential genes ^24^. Additionally, there exists some evidence that genetic drift may have significant functional and evolutionary consequences ^25^.

Detecting selection in MTBC at the genome-wide level remains a challenging task due to limited genetic diversity. The significant accumulation of non-synonymous substitutions has been previously used to characterize patterns of mutation accumulation in large categories of genes ^24,26^; however, these studies employed a limited number of strains. Of note, the number of MTBC sequences has undergone a recent and rapid expansion, with studies involving hundreds to thousands of strains. The large number of available sequences has allowed, for example, the estimation of the ratio of non-synonymous to synonymous substitutions (dN/dS) signatures in more than 10,000 strains ^27^, thereby allowing the identification of novel targets of selection with some probably related to host-pathogen interactions. Host-pathogen interaction signals are specially challenging as they are likely obscured by the force exerted by antimicrobial therapies. Weaker signals are also expected in genes related to second-line drugs related to the relative under-use of related treatments and the low abundance of associated resistant strains in genome databases ^28^.

We reasoned that to detect signs of selection, we should focus on when and/or where they occurred in the phylogenetic tree instead of averaging signs across the phylogeny. In this new study, we developed a methodology to study temporal signs of selection in MTBC genes and identified positive selection in a larger number of genes than previously described. This allowed the identification of past and currently unknown players in MTBC evolution, particularly two-component systems, related to host adaptation and second-line drug resistance. This new methodology can be applied to other tuberculosis settings to explore signs of selection associated with changing selective pressures and could be extremely useful to unravel hidden details in the evolution of other human pathogens.

## Results

### Dataset Preparation

We downloaded all samples described in Brites *et al. ^4^*, Coll *et al. ^29^*, Stucki *et al. ^30^*, Guerra-Assunçao *et al. ^31^*, Zignol *et al. ^32^*, Bos *et al. ^33^*, Ates *et al. ^34^*, Comas *et al. ^10^*, Comas *et al. ^35^*, Borrell *et al.^36^*, and Cancino-Muñoz *et al.^37^* and obtained whole-genome sequencing data from 9,240 samples comprising the primary human- and animal-adapted MTBC lineages. We mapped Fastq files for each sample against the inferred ancestor of the MTBC and extracted genomic variants (**Methods**), from which we derived a multiple sequence alignment and a phylogeny. The huge size of the phylogeny and the multifasta file obtained made unaffordable certain parts of the planned subsequent computational analyses, hence we used Treemer to prune the tree down to 4,958 leaves (**Table S1**) while maintaining 95% of the original genetic diversity. With this final set of selected samples, we reconstructed a multiple sequence alignment and a phylogeny (**Figure S1a**).

We mapped each genomic variant to the inferred phylogeny using PAUP (Phylogenetic Analysis Using Parsimony). This step provides information regarding the branch in which every mutation appeared, which allows the identification of homoplastic variants - those that appeared multiple times in different branches of the phylogeny - and the relative ‘age’ of every mutation, calculated as the node-to-root genetic distance.

### Scars of Past Selection and Drift in Almost Half of the MTBC Genome

As a first step, we calculated the pN/pS values for genes that possessed up to ten identified variants (n=3,690). A previous study stated a mean pN/pS value for the complete MTBC genome considerably under 1 ^38^. In agreement with this result, we found that 90% of the genes evaluated possess a pN/pS value less than 1 (**Figure 1a**, pN/pS IQR 0.477-0.804), suggesting ongoing evolution under purifying selection. A high pN/pS may reflect the recent origin of the MTBC, given the time-dependent nature of the accumulation of non-synonymous variants ^39^.

**Figure 1.**
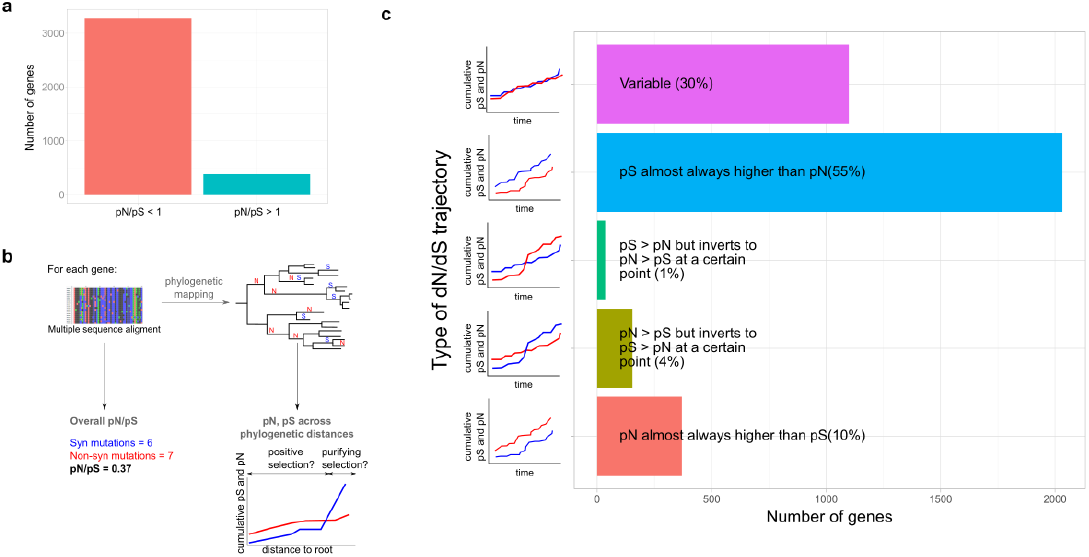
Gene-by-Gene Calculation of pN/pS Over Phylogenetic Time. **a**. Bar plot showing the number of genes currently displaying a pN/pS > 1 and a pN/pS < 1. **b**. From the alignment, we inferred the current pN/pS; however, when mapping different mutations onto the phylogeny, we inferred how the pN and pS rates changed over time. **c**. Five categories grouping studied genes according to their trajectories.

Of note, the pN/pS value for a gene results from the pN and pS values calculated with all gene mutations found across the phylogeny (what we term the ‘overall pN/pS’ in **Figure 1b**). This value does not reflect changes in selective pressures over time and lineages as the pathogen has potentially faced different environmental “challenges.” As we estimated the genetic distance to the root for each mutation as a relative measure of time, we calculated temporal trajectories for pN/pS for each gene during MTBC evolution (**Methods**, **Figure 1b**). By doing so, we classified all genes according to their pN and pS trajectories over time into five different categories (**Figure 1c**, **Figure S2a**, **Table S2):** (i) pS almost always higher than pN (n=2,032); (ii) pN almost always higher than pS (n=154); (iii) pS > pN but inverts to pN > pS at a certain point (n=35); (iv) pN > pS but inverts to pS > pN at a certain point (n=370); and (v) complex pN and pS trajectories with multiple cross-points, which don’t support proper categorization (n=1,099). If our classification reflects differences in the selection pattern over time, we expect that those genes with stable trajectories (‘always higher’/‘always lower’) will have accumulated low variances in pN/pS when pooling timepoints. Conversely, we expect changing trajectories to display high variance between timepoints (**Methods**, **Figure S2b**). As predicted, we failed to observe significant differences in variance (Welch t-test, p-value > 0.05) in genes belonging to the ‘pN almost always higher’ or ‘pS almost always higher’ categories. In both cases, the pN/pS cumulative variation has a value around zero. However, categories with changing trajectories displayed significant differences (Welch t-test, p-value << 0.01), using ‘pS almost always higher category’ as the reference category.

In summary, and in contrast with the observation that 90% of genes possess an overall pN/pS < 1, only 55% of genes (n=2,032) maintained a pN/pS value below a value of 1 since divergence from the MTBC common ancestor. This set of 2,032 genes is overrepresented for experimentally confirmed essential genes in both *in vivo* and *in vitro* conditions (chi-square test, p-values 0.003 and <2.2E-16, respectively). In contrast, 45% of the genes (n = 1,658), mainly those initially classified as being under purifying selection, may have faced other types of selective pressures or genetic drift.

These results suggest that many genes have been subjected to periods of non-synonymous substitution accumulation. Distinguishing between genetic drift and positive selection at a particular time point remains challenging. We expect founder effects to play a crucial role during the early evolution of MTBC, and they may drive a number of the unstable trajectories observed. However, given that MTBC is clonal, positive selection and genetic drift are both expected to have a functional impact. Our analysis identifies a set of genes that shows a pN/pS > 1 near the root but changed to pN/pS < 1 near the leaves (n=370), suggesting that selection and/or founder effects favored the fixation of non-synonymous mutations at early times but that the gene functionality remained conserved at later times. We found that this gene category was enriched for conserved hypotheticals (fisher test, p-value = 0.02) and protein and peptide secretion (fisher-test, p-value = 0.05). Intriguingly, we also discovered that certain genes that fell into this category encode known MTBC epitopes (which we will explore below). Of particular note, the presence of 154 genes almost always exhibiting a pN higher than pS. This gene category is enriched for non-essential *in vitro* genes (chi-square test, p-value=0.005) from three main categories; antibiotic production and resistance (fisher-test, p-value=0.02), conserved hypotheticals (fisher-test, p-value=0.02), and unknown functions (fisher-test, p-value=0.03). The mix of genes with a clearly identified function and hypothetical genes suggests that, in some cases, positive selection has been acting through the evolutionary story of some genes while others are likely under genetic drift.

### Evolutionary Trajectories Identify Sensor Proteins of Two-Component Systems Under Positive Selection

An increasing number of non-synonymous mutations that start to grow near the leaves may indicate the action of more recent selective forces and suggest unpurged transitory polymorphisms. To distinguish between the two possibilities, we examined the group of genes with a pS > pN in the internal branches but a pN > pS near the leaves (n=58, **Table S2**). Antibiotic resistance genes represent a clear instance of recent positive selection, and we hypothesized that their initial trajectory should reflect conservation of gene function, as they usually perform relevant biological functions and only recently started to diversify due to antibiotic selective pressure. Encouragingly, data for the antimicrobial resistance genes such as *rpoB, katG, embB, gidB*, and *rpsL* supported this hypothesis. The genetic distance from the root at which we detected a change in the selective pressures correlates with the time at which each antibiotic became a treatment for tuberculosis. Genes related to resistance to the most recently employed drugs began to accumulate non-synonymous variants at a higher genetic distance to the root than those used in early periods. This point is placed at 1.566483e-04 for *gidB* (streptomycin, first antibiotic used in tuberculosis treatment in 1946), 1.637692e-04 for *katG* (isoniazid, use began in 1952), and 1.774088E-04 for both *embB* (ethambutol, 1966) and *rpoB* (rifampicin, 1972). These results suggest that our approach possesses sufficient sensitivity to detect recent instances of positive selection.

Among those genes unrelated to antimicrobial resistance, we found several components of toxin-antitoxin systems, including *vapC29, vapB3, vapC35, vapB40, vapC22*, and *vapC47*, which are critical for the adaptation of bacteria to different stressful conditions. For example, VapC22 has a significant role in virulence and innate immune responses in particular ^40^. Other significant virulence regulators in MTBC are the two-component systems (2CS), which are critical players in extended transcriptional networks. 2CSs comprise a sensor protein coupled to a transcription factor - the sensor protein activates the transcription factor in response to a specific stimuli to trigger a regulatory cascade. We have previously described *phoR*, which encodes the sensor component of the PhoPR 2CS, as an important player in MTBC evolution ^9^ as illustrated by the high levels of accumulation of non-synonymous variants over time. Our data shows that *kdpD*, a gene that encodes the sensor component of the KdpDE 2CS, displays a similar pattern, with a dN/dS value that reached ~2 at some points during MTBC evolution. In both 2CSs, the genes encoding the regulatory protein (*phoP* and *kdpE*) display high conservation at the functional level, with the pS values consistently higher than the pN values. For the NarLS 2CS, both the regulatory protein (*narL*) and the sensor protein (*narS*) exhibit changing patterns towards recent positive selection; however, as for the other described 2CSs, the sensor domain of *narS* accumulates more non-synonymous variants (fisher test, p-value = 0.036). Our analysis suggests that sensor proteins of 2CSs allow MTBC strains to adapt to varying environments during host-pathogen interaction.

### Epitope Mutations are Older and Show Divergent Evolutionary Trajectories Compared to the Rest of the Antigen

Contrary to many other pathogens, the *M. tuberculosis* genome regions recognized by the host tend to be conserved, albeit with some exceptions ^24,41^. Given our new results revealing past “scars” of selection in MTBC genes, we analyzed the pN/pS trajectory of a total of 179 antigens harboring 1,556 epitopes ^42^. Specifically, we aimed to evaluate a hypothesis that epitope and non-epitope regions of the antigen experience different selective pressures and that the former most likely reflect interactions with the immune system while the latter reflects the evolution of gene function.

Our results revealed that ~60% of the antigens analyzed exhibited a pN/pS value of < 1 across phylogenetic history, providing evidence for their conservation since their diversification of the MTBC from a common ancestor (**Table S3**). Of note, a relevant proportion of antigens (11%) accumulated a high number of non-synonymous variants in internal branches, which now appear to be conserved (**Table S3**). For example, the *mpt64* gene encodes for a known antigen employed in diagnostic tests. When mapping the genetic variants in the MTBC phylogeny, most non-synonymous mutations map to the L5 ancestral branch in a large clade of the L1.2.2 sublineage and a group of L4.10 strains (**Figure 2a, b**). Other antigens, such as *eccD2*, Rv1866, *fadD21*, or Rv2575, exhibited a similar pattern. Apart from human-adapted clades, specific antigens accumulated non-synonymous mutations in deep branches of the animal-adapted lineages, such as Rv2575 or *IlvB1*. This suggests that these antigens were under positive selection or genetic drift driven by founder effects when MTBC diversified.

**Figure 2.**
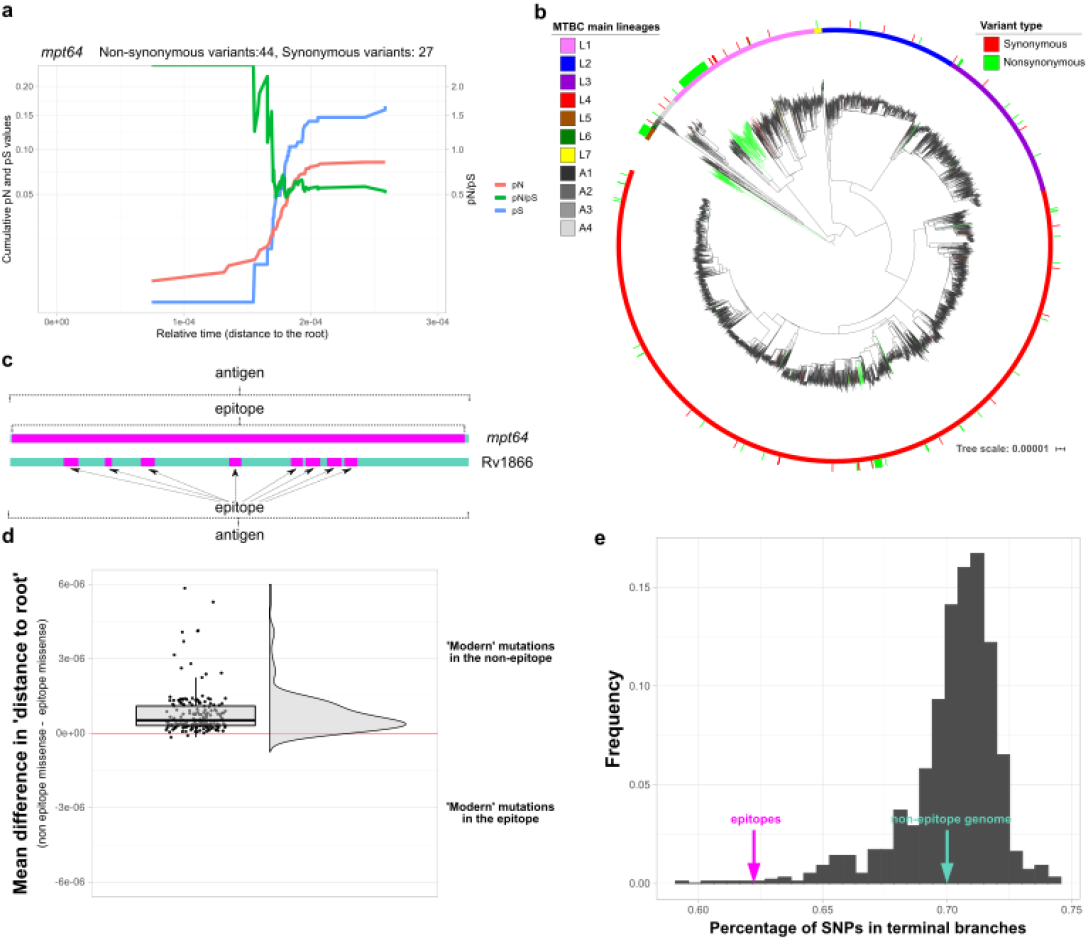
Specific Antigenic Genes Show Signs of Early Positive Selection. **a.** Cumulative pN, pS, and pN/pS trajectories over time for the *mpt64* antigen (Rv1980c). The x-axis represents the genetic distance of each node to the root. The left y-axis represents the cumulative pN (red line) and pS (blue line) values. The right y-axis represents the pN/pS. **b.** Maximum-likelihood MTBC phylogeny with mapped *mpt64* variants. The sticks in the outer circle mark the strains with variants identified (red synonymous, green non-synonymous). Deep non-synonymous mutations can be found in deep nodes of L1 and L5. **c.** Some epitopes comprise the entire antigen (such as in *mpt64*), while in genes such as Rv1866, the epitope represents a small subset of regions embedded in the antigen. **d.** Raincloud plot of the mean differences in the distance (to root) value between the non-epitope and the epitope mutations for each antigen. **e**. Distribution of SNPs found in terminal branches for 1,000 randomly selected sets of non-epitope fragments (grey bars). The percentage of SNPs observed in the epitopes differs from this distribution (~62%, z-score = −4,28, pink arrow), while the percentage of SNPs found in the rest of the genome remains similar to the distribution (~70%, green arrow).

For another group of antigens (27%), the pN/pS value failed to show a definitive trajectory (**Table S3**). Specific antigens showed a pattern of pN/pS value of ~ 1 since the diversification of the MTBC from a common ancestor. This pattern could reflect two different causes: genetic drift or differential selective pressures in different MTBC clades/lineages, which could be masked when calculating a common pN/pS for all lineages. The second option is defined by an accumulation of non-synonymous mutations in specific MTBC clades and synonymous mutations in other clades. As a result, the overall pN/pS value would be ~ 1. We observed this scenario, for example, in the *lpqL, mce2A*, and *esxH* genes; in these cases, we found an elevated accumulation of non-synonymous mutations in deep branches of the L1, L2, and *M. africanum* lineages, although they are highly conserved in modern lineages. Other genes exhibited a similar pattern (**Table S3**), while others could have evolved under the effect of genetic drift.

In general, the evolution of antigens does not essentially differ from other genes in their respective functional categories. When we compared the trajectories of the antigens against such genes, we failed to encounter statistical differences between the distributions (Fisher test, BH adjusted p-value > 0.05).

Of note, antigens have a myriad of distinct functions, but the immune system only recognizes specific regions of the antigens - the epitopes. In some cases, epitope regions cover the entire antigen (as for *mpt64*), so selection acts on the antigen and epitope equally. In other cases, epitopes represent only a small fraction of the antigen and may be subject to different selective pressures than the rest of the gene (**Figure 2c**). When exploring whether selection at the epitope level drives different antigen trajectories, we encountered the Rv1866 locus as a clear example. This antigen has a pN/pS value of >1 near the root, but its value changes to <1 near the leaves, suggesting the action of distinct types of selection across the phylogeny; however, the epitopes contained are highly conserved with a pN/pS value of <1 during the complete trajectory.

In most cases (**Table S3**), the evolutionary trajectories of epitopes seem to be unlinked to the rest of the antigen, with most epitopes being conserved. We hypothesized that epitopes might reflect past selection events to adapt to different populations during the initial expansion of the MTBC. In general, the mean relative phylogenetic age (measured as the genetic distance to the root) of the non-synonymous variants present in the epitopes is older than the non-synonymous variants of the non-epitope regions of the antigen. This phenomenon can be observed when pooling all epitope vs. non-epitope variants (Welch t-test, p-value = 8e-07) and when splitting by different genes (**Figure 2d**) (although with considerable overlap, as expected). Consequently, we expect fewer mutations to accumulate at phylogeny tips if epitope conservation becomes more important at a later stage. The proportion of mutations in epitopes falling in terminal branches (62%) is significantly lower than in sets of regions of the same size randomly selected from the non-epitope genome (70%, z-score = −4.28, P(x<Z) = 0.00001, **Figure 2e**). This suggests the more robust nature of negative selection on epitopes than the rest of the genome in circulating strains.

Thus our results provide further evidence for the generally unlinked nature of gene and epitope evolution, which had been previously established in smaller sets of samples ^24,38^. In addition, we demonstrate that interaction with the immune system likely drives epitope conservation (as it is the only function in common among epitopes), while non-epitope regions reflect the selection of the gene’s biological function. Finally, mutations in epitopes mainly reflect older fixation events while the rest of the genome accumulates mutations more rapidly.

### Novel Candidate Drug Resistance Regions Revealed by a Dataset Enriched for MDR/XDR-TB-associated Strains

Identifying genes involved in resistance to second- and third-line drugs and new and repurpose drugs remains challenging. We reasoned that if our approach was powerful enough to identify changing selective pressures due to the introduction of first-line antibiotics, we should detect changes in genes associated with the treatment of multidrug-resistant (MDR)- and extensively drug-resistant (XDR)-tuberculosis patients. We assembled and compared a dataset enriched for MDR (n=312) and XDR (n=132) strains and additional sensitive controls to our global dataset (**Figure S1b**, **Table S1**). Our analysis revealed instances of genes with an increased pN value towards the leaves of the tree for the MDR/XDR dataset compared to the global dataset. Our approach correctly identified genes associated with MDR, such as *gyrA* (quinolones), *ethA* (ethionamide), and *rpoC*, which compensates for the fitness cost of MDR strains (**Figure 3a**). Importantly, we also identified less-well-studied genes with a similar profile, including Rv0552, Rv1730c, *alr* (Rv3423c), *eccC4* (Rv3447), *eccCa1* (Rv3870) (**Figure 3a**), and Rv3883c (*mycP1*). To formally evaluate their association to different drugs, we generated computational models (**Methods**, **Figure S1b**) to link the observed drug-resistant phenotypes with mutations in genes with a changing pN/pS pattern. Well-known resistance-conferring genes such as *rpoB, katG*, or *rpsL* exhibit a strong statistical association with drug-resistant phenotypes, as expected (**Table S4**, **Figure 3b**). Corroborating our observations, the identified less-well-studied genes displayed a significant association with resistant phenotypes for second-line drugs. For example, Rv1730 weakly associated with fluoroquinolone-resistant phenotypes (Wald test, p-value = 0.02), *alr* with D-cycloserine and fluoroquinolones (Wald test, p-value = 0.007 and p-value = 0.001), *eccC4* with fluoroquinolones (Wald test, p-value = 0.01), and *eccCa1* with D-cycloserine and aminoglycoside injectable agents (Wald test, p-value = 0.04 and p-value = 0.002).

**Figure 3.**
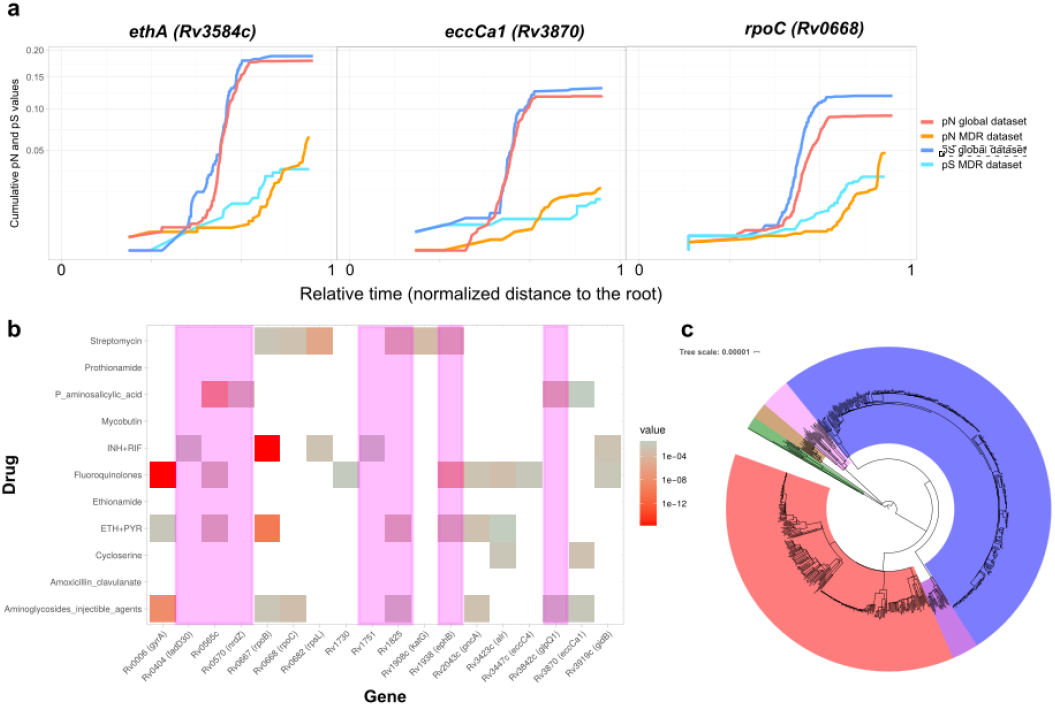
Identification of Genes Related to Second-line Antibiotic Resistance. **a.** Three genes showing signs of ongoing positive selection in the MDR-enriched dataset but ongoing purifying selection in the global dataset. The x-axis represents the node-to-root genetic distance normalized in the 0-1 range to merge data from both trees as a measure of relative time. The y-axis represents the cumulative dN and dS values. **b.** A computational model has been constructed for each antituberculosis drug to identify specific gene mutations associated with resistance. In the matrix, rows represent antibiotics and columns represent genes suspected to be under positive selection in the MDR-enriched dataset. Colored cells (from gray to red) indicate a statistically significant association between non-synonymous mutations found in the genes and resistant phenotypes. Genes marked in pink show a strong association with drug-resistant phenotypes due to phylogenetic variants, suggesting that the association may be spurious. **c.** Maximum-likelihood phylogeny constructed with the MDR-enriched dataset showing an overrepresentation of L2 (blue) and L4 (red) strains.

Of note, our analysis did have certain limitations; for example, given the combined therapy administered in tuberculosis treatment, the same gene may correlate with several antibiotics. Likewise, given the enrichment of this dataset with L4 and L2 strains (**Figure 3c**), non-synonymous phylogenetic variants in genes such as *fadD30* (Rv0404), Rv0565c, *nrdZ* (Rv0570), Rv1751, Rv1825, *ephB* (Rv1938), and *glpQ1* (Rv3842c) appear to be associated with drug-resistant phenotypes but are likely neutral markers, a previously reported phenomenon ^43^. The identification of previously uncharacterized genes represents the overall value of the analysis, with results requiring corroboration by fine-grain *in vitro* experiments.

### Selection Also Acts in Non-coding Regions

Beyond mutations affecting coding regions, we (and others) have established the importance of mutations in intergenic regions in shaping the pathogen’s phenotype, as they can alter gene regulation. Hence, natural selection can also target these positions. Using a Poisson distribution, we identified 290 intergenic regions possessing more mutations than expected by chance (BH adjusted p-value < 0.05). 270 of the intergenic regions harbor homoplastic mutations, representing a good correlate of positive selection in MTBC. Certain mutations had been previously categorized as resistance-conferring variants, including 1673425C>T (upstream *fabG1*), 4243221C>T (between *embC-embA*), or 2715342C>G (upstream *eis*) (**Table S5**). We found other mutations in intergenic regions suspected of being related to drug resistance; however, the exact mutations were not present in the PhyReSse and ReseqTB catalogs.

We also calculated the ratio of intergenic variants per intergenic site compared to the ratio of synonymous variants per synonymous site of the flanking genes (pI/pS) for each intergenic region as a measure of selective pressure, as previously proposed by Thorpe et al. ^44^. We found a mean pI/pS value of 1.03 (95% CI: 0.98 - 1.07), near the expected value of 1 when under no selection; however, 123 intergenic regions appeared as outliers of this distribution (**Table S6**) as they exhibit pI/pS values greater than 2.058 (calculated as Q3 + 1.5*IQR ^45^). A gene set enrichment analysis (GSEA) of gene ontology (GO) functions of flanking genes of these intergenic regions demonstrated that the most overrepresented functions (Hypergeometric test, BH corrected p-value < 0.05) are responses to acid chemicals, REDOX processes, and regulation of DNA templated transcription. The identification of REDOX is in agreement with oxidative metabolism playing a role in macrophage survival and drug resistance ^46–48^. A previous study reported that changes in regulatory regions (mostly intergenic) could significantly affect the transcription rates of downstream genes ^49^. Therefore, the positive selection of these regions may not be surprising.

## Discussion

Pathogen diversity reflects a balance between evolutionary forces. In the case of the virtually clonal MTBC (^9^, highly diverse and highly conserved genes can be identified despite low genetic diversity ^1^, thereby suggesting the activity of distinct evolutionary forces. While metrics such as pN/pS present with certain limitations ^39^, they allow the identification of the footprints of evolutionary forces. pN/pS has the power to identify selection at the genome-wide level ^26,27^, including traces of positive selection in specific genes, gene categories, and/or lineages ^23,50,51^. Analyses revealed an average pN/pS value across the MTBC genome of around 0.7, well below the value of 1 expected for any organism but high compared to others. This likely reflects the recent emergence of MTBC with the presence of many transitory polymorphisms ^39^ and the impact of genetic drift in the form of bottlenecks and founder effects (Herbergh 2008). However, the balance of evolutionary forces shaping genetic diversity is dynamic, and what was under positive selection or drift in the past may be under negative selection in the present and *vice versa*. This idea is illustrated in our work by the striking discovery of scars of elevated non-synonymous rates in almost half of the MTBC genome, contrasting with previous reports (Coscolla et al. 2015; Pepperell et al. 2013).

Our analyses identified different temporal evolutionary dynamics in *M. tuberculosis* genes. In one important category, genes are subjected to positive selection or genetic drift early in MTBC evolution but to purifying selection near the leaves. A prominent example of this phenomenon is the accumulation of early non-synonymous variants in epitopes such as *mpt64*. Deep mutations may reflect past events such as founder effects or drift, but our analysis suggests that mutations in epitopes are older when compared to other regions of the genome and that epitope evolution is not linked to the evolution of the rest of the antigen and functional category. These observations are compatible with scenarios suggesting early co-evolution of host and pathogen populations ^5^.

We also identified genes subjected to purifying selection in the past but to current positive selection. The abrupt shift in the pN/pS values in resistance-conferring genes illustrates the impact of antibiotic treatments on MTBC evolution. While our novel approach detected an increase in the pN/pS in a set of genes in MDR and XDR strains, we did not observe this increase in strains not exposed to second-line drugs. This finding allowed the proposal of a set of candidate genes that confer resistance to second-line antitubercular drugs. Previous reports have suggested that genes such as *alr* or *eccCa1* can confer resistance to MDR treatments ^15,52–56^; however, novel genes identified in this study highlight our incomplete understanding of the genetic basis of resistance, in particular for second-line and new drugs. Our approach also detected genes unrelated to antibiotic resistance that have been subjected to recent positive selection, a finding missed when applying averaged pN/pS ratios. We commonly encountered the sensor component of 2CSs in this gene-set, and our previous data established robust signs of recent positive selection in *phoR*, the sensor component of the PhoPR 2CS ^9,57^. This finding suggested that non-synonymous mutations in *phoR* participate in host adaptation by regulating *PhoP*, a major regulator of MTBC physiology and virulence. We now show a similar occurrence in two other sensor proteins - KdpD and NarS. Thus, the accumulation of non-synonymous mutations in sensor proteins may represent a common strategy used by mycobacteria to adapt to the changing environment during infection.

In addition to coding regions, we also found traces of selection in non-coding sequences, which agrees with previous findings ^44^. While identifying selection pressures on intergenic regions remains challenging, given the problematic interpretation of the functional effect of variants that fall in these areas, homoplastic mutations and the comparison of variants against surrounding genes provide a good framework. Variants accumulation in these regions can impact the regulation of nearby gene expression ^16,49^. Again, drug resistance appears to represent the strongest selective force; however, variants found in these regions also impact transcription factor activity and oxidative metabolism

We are aware of the limitations of our current study. The study of past traces of selection in MTBC members remains challenging due to the low genetic diversity present; however, we attempted to maximize genetic diversity to gain resolution by including a broad representation of the main MTBC lineages. Unfortunately, subtle traces of selection affecting small subclades or groups of strains can be masked using this strategy – indeed, this is illustrated by our study when lineage-positive signs of selection fail to appear in our analysis. For example, Menardo *et al*. have described a high number of non-synonymous mutations in the epitopes of *esxH* ^41^. This finding is not reflected when considering all lineages but only when we search lineage by lineage (**Table S3**). Further analysis focusing on specific subclades may illuminate differential evolutionary pressures within the MTBC. Furthermore, we only analyze mutations fixed in the phylogeny, so we only infer an approximate picture of the evolutionary forces that have shaped complex evolution in the past. In addition, the low variability present in the MTBC, strain subsampling, and lack of metadata/dates for most deposited genomes make absolute dating for some studied mutations extremely challenging. We are also aware that, in some cases, genetic drift may be mistaken with other selection forces; however, this does not preclude those changes from having a functional effect ^25^.

Finally, we note that our approach can be used as a blueprint to study the evolution of several bacterial species. For example, the *Salmonella* genus includes strains exhibiting high host-specificity and those with the general ability to infect many hosts ^58^. The gene-by-gene evaluation of past and current selective pressures could shed light on the genomic determinants that drive differing specificity. The same approach could be valid with *Helicobacter pylori*, a pathogenic bacteria that causes gastric infections and is highly specialized at infecting human hosts ^59,60^. MTBC displays virtually no recombination or ongoing horizontal gene transfer (which is not the case of *H. Pylori* or *Salmonella*), making the interpretation of the results more straightforward; however, we anticipate that, taking into account population structure, our approach can be adapted to answer a range of evolutionary questions in pathogen evolution.

## Methods

### Variant Analysis Pipeline and Phylogenetic Reconstruction

All samples were analyzed using our variant analysis pipeline, which has been extensively described in a previous publication ^61^. Briefly, FASTQ files were trimmed to remove low-quality reads using fastp ^62^ (version 0.12.5, arguments --cut_by_quality3, --cut_window_size=10, -- cut_mean_quality=20, --length_required=50, --correction) and aligned to the most likely inferred ancestor of MTBC ^24^ using the BWA-MEM algorithm ^63^. Potential optical and PCR duplicates were removed with Picard tools ^64^, while reads with a MAPQ value < 60 were also discarded. Variant calling was performed using SAMtools ^65^, VarScan ^66^, and GATK ^67^. A pileup file was created with SAMtools from the BAM files, and VarScan was then used to extract variant positions from this pileup file (version 2.3.7, arguments --p-value 0.01 --min-coverage 20 --min-reads2 20 --min-avg-qual 25 --min-strands2 2 --min-var-freq 0.90), while GATK was used to extract INDELS (version 3.8-1-0-gf15c1c3ef, HaplotypeCaller and SelectVariants functions). To remove mapping errors, detected variants were discarded if found in INDEL areas or areas of high variant accumulation (more than three variants in a 10-bp defined window). Variants were then annotated using SnpEff (version 4.2) ^68^. Variants associated with proline-glutamate/proline-proline-glutamate (PE/PPE) genes, phages, or repeated sequences were also filtered out (**Table S7**) as they tend to accumulate false-positive SNPs owing to mapping errors. Finally, with the selected high-quality variant calls, a non-redundant variant list was created and used to retrieve the most likely allele at each genomic sequence to generate a variant alignment.

The first phylogeny was constructed with all samples that passed a minimum depth coverage threshold (median 25x) and had no mixed infections (n=9,240). This initial phylogeny was constructed using MEGA-CC ^69^ and the Neighbor-Joining algorithm. Later, we pruned the phylogeny with Treemer ^70^ to obtain a smaller tree for subsequent computational analyses. A reduction of just 5% of the initial genetic diversity led to the selection of 4,958 samples. With these selected samples, a maximum likelihood phylogeny was constructed using IQTREE ^71^ (version 1.6.10) with the GTR model of evolution, taking into account the invariant sites and with an ultrafast bootstrap ^72^ of 1,000 replicates.

### Phylogenetic Variant Mapping and pN/pS Trajectories

After phylogenetic reconstruction, the mutations called in the 4,958 samples (n=368,719) were mapped onto the phylogeny. For his purpose, the ancestral state of each polymorphism in each node was reconstructed using PAUP ^73^ with a weight matrix that punished reversions with a 10X multiplier. From this information, the phylogenetic node at which each variant appeared was obtained. Later, a relative age derived from the branch length information for each variant was assigned for each variant. This relative age is the genetic distance from the ancestral node to the bisection point of the target branch on which the variant appears. Finally, the cumulative pN and pS trajectories were calculated for each gene using the potential synonymous and non-synonymous sites inferred using the SNAP tool ^74^ and plotting the pN and pS values at each timepoint, taking into account the variants that appeared before this timepoint.

Initially, we classified the genes according to their pN/pS trajectories with the following criteria:

I. genes with a cumulative pN/pS < 1 at more than 95% of the sampled points were classified as ‘pS almost always higher than pN’
II. genes with a cumulative pN/pS > 1 at more than 95% of the sampled times were classified as ‘pN almost always higher than pS’
III. genes in which the cumulative pN/pS changed from >1 to <1 or vice versa more than three times were classified as ‘variable’
IV. genes in which the cumulative pN/pS changed from >1 to <1 or vice versa less than four times and that the cumulative pN/pS started being <1 but ended >1 were classified as ‘pS > pN but inverts to pN > pS at a certain point’
V. genes in which the cumulative pN/pS changed from >1 to <1 or vice versa less than four times and that the cumulative pN/pS started being >1 but ended <1 were classified as ‘pN > pS but inverts to pS > pN at a certain point’.

This classification was reviewed manually at a later stage. Genes with less than ten mutations were not considered for subsequent analyses.

The cumulative pN/pS variation for each gene was calculated as:

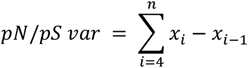

with *x* the cumulative pN/pS value at each of the sampled *i* points. The first three values of each gene’s cumulative pN/pS value were not considered, as the initial values can show significant differences due to a low number of mutations.

### Epitope and Antigen Analysis

All linear epitopes (n=1,556) found in the IEDB database ^42^ that belong to *M. tuberculosis* in August 2019 were downloaded. All linear epitopes with overlapping coordinates with regards to the H37Rv reference strain were merged into unique non-overlapping ‘contigs’ (n=718). The potential synonymous and non-synonymous sites were inferred using the SNAP tool ^74^. All genes containing such epitopes were considered antigens, except those genes not considered in the variant calling step, as explained above (PE/PPE, phages).

The percentage of SNPs that occur in these 718 regions that appear in terminal branches of the phylogeny were determined using the information derived from PAUP. The percentage of SNPs in the rest of the genome (not considering these 718 regions) that fall in terminal branches were also determined. To evaluate if the difference between these values was statistically significant, 718 segments of the non-epitope genome with the same length as the epitope regions set, 1,000 times, were selected. For each iteration, the percentage of SNPs found in terminal branches was calculated and plotted in a distribution. Finally, a z-score (see below) between the distribution and the value observed for the epitopes was calculated.

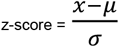

### Gene Set Enrichment Analysis

Several approaches for functional category enrichment were performed to compare genes present in our sets of interest against other genes. For the essentiality enrichment, the *in vivo* ^75^ and *in vitro* ^76^ classification of genes was used, and the enrichment in these categories tested with Fisher tests. For GO enrichment, the Bingo tool ^77^ was used with a hypergeometric test (sampling without replacement) and the Benjamini-Hochberg correction for multiple testing comparisons. Finally, the enrichment of the functional categories was also evaluated ^78^ employing Fisher tests corrected with the Benjamini-Hochberg procedure.

### Drug-resistant Dataset Preparation and Analysis

Drug-resistant strains were downloaded from the TBportals database ^79^ on October 22, 2019 (n=656). Samples were classified according to their drug-resistant phenotype and then passed through the variant analysis pipeline described above. A maximum-likelihood phylogeny was constructed using IQTREE with the previously described options, including samples from the Comas *et al*., 2013 study to achieve nodes from lineages underrepresented in the TBportals database.

The pN/pS trajectories were calculated and classified as explained for the other dataset

A matrix was next created that included phenotypic information for each tested drug (resistant/susceptible) and the presence/absence of non-synonymous mutations in the gene set classified as having a trajectory in which the pS > pN but inverts to pN > pS at a certain point for each sample. A set of binomial logistic regression models was constructed with this data, explaining the observed phenotypes based on the presence of non-synonymous mutations on selected genes. These models were trimmed *a posteriori* following a backward stepwise methodology, selecting the set of regressors that show the best Akaike Information Criterion.

## Acknowledgments

This project received funding from the European Research Council (ERC) under the European Union’s Horizon 2020 research and innovation program (Grant agreement No. 101001038 (TB-RECONNECT)). In addition, this work was funded by projects PID2019-104477RB-I00 from Ministerio de Ciencia (Spanish Government) and AICO/2018/113 from Generalitat Valenciana.

## Author Contributions

I.C. conceived this work. A.C.O. and M.G.L. analyzed the data. A.C.O. wrote the first version of the draft. A.C.O., I.C., M.G.L. critically reviewed and contributed to the final version of the paper.

## Supplementary Material

**Figure S1.**
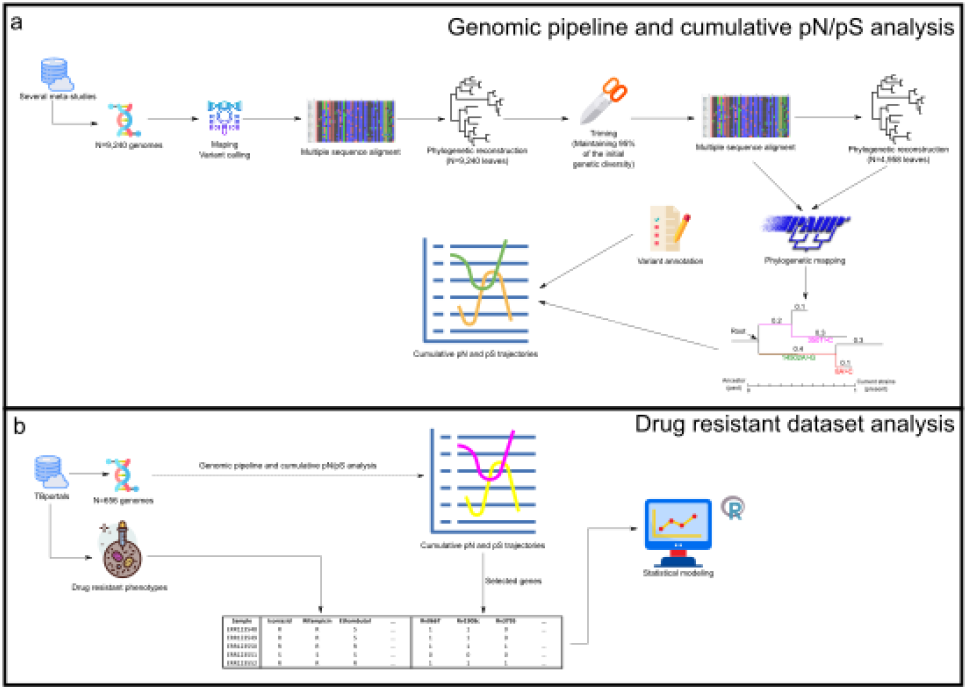
Workflow Followed in Different Analyses. **a.** From public repositories, we downloaded more than 9,000 MTBC genomes. After reconstructing a phylogenetic tree, the dataset underwent a trimming process to reduce the number of samples while maintaining as much genetic diversity as possible. From these reduced datasets, we reconstructed a tree and an alignment. PAUP mapped each detected polymorphism into the phylogeny. Finally, knowing the annotation of the polymorphisms and the branch in which they appeared allowed us to generate pN/pS trajectories. **b.** TBportals was used to obtain a dataset enriched for resistant strains. The same approach as described above was applied (except for the trimming step), thereby obtaining pN/pS trajectories for each gene based on the information of this new dataset. We also downloaded drug-susceptibility test (DST) information for each resistant strain. Combining both the genomic and the phenotypic information allowed the generation of computational models linking the observed phenotypes to mutations in specific genes.

**Figure S2.**
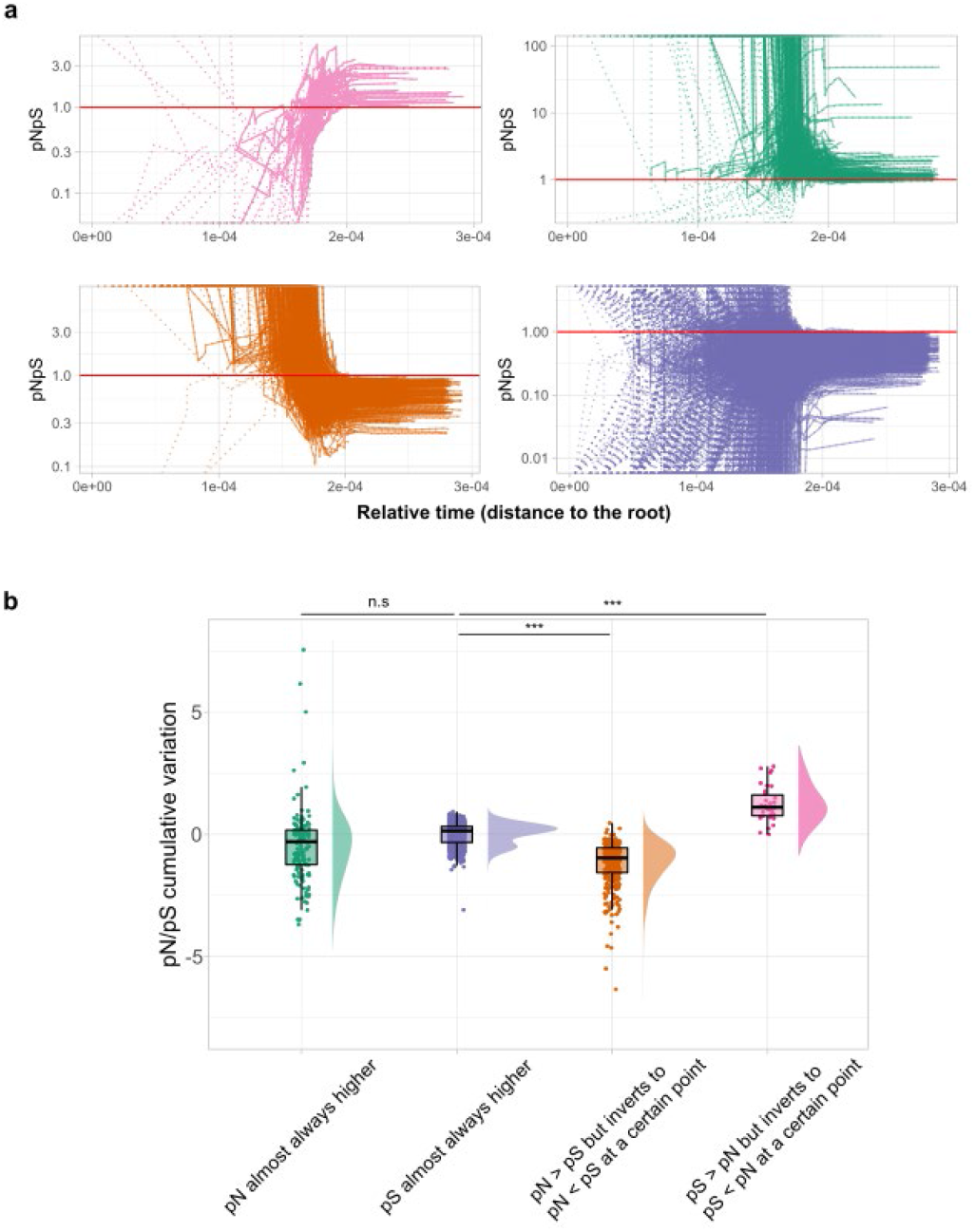
Classification of Genes According to pN/pS Trajectory. **a.** pN/pS variation across the phylogeny, from root to tips. Each line corresponds to a different gene. Genes were classified as: (i) pS almost always higher than pN (blue); (ii) pN almost always higher than pS (green); (iii) pS > pN but inverts to pN > pS at a certain point (pink); (iv) pN > pS but inverts to pS > pN at a certain point (orange); (v) pN and pS had complex trajectories (not plotted). The red horizontal line marks pN/pS = 1. The first three values of the trajectory (dashed in the plots) were not considered for classification, and the rest of the analysis as they present with high variability due to a small number of mutations. **b.** Cumulative pN/pS variation distribution for each gene category. Categories reflecting ‘stable’ trajectories (‘pN almost always higher’ and ‘pS almost always higher’) accumulated low variance in pN/pS and displayed no significant differences (Welch t-test, p-value > 0.05). In both cases, the pN/pS cumulative variation is around zero. In contrast, categories with changing trajectories display significant differences (Welch t-test, p-value << 0.01), using ‘pS almost always higher category’ as the reference category.

**Table S1.** Samples used in the analyses, including accession numbers and the main phylogenetic lineage

**Table S2.** Classification of genes in the five main categories defined in the main results

**Table S3.** Classification of the antigens/epitopes studied, including the categories proposed for each of the features and the lineages in which they show differential trajectories.

**Table S4.** P-values of the computational models generated - Genes marked in yellow display significant values, probably due to phylogenetic markers.

**Table S5.** Homoplastic variants called in the intergenic regions analyzed.

**Table S6.** 123 intergenic regions that exhibited pI/pS values that are outliers of the genomic pI/pS distribution. Observed and expected mutations in the intergenic regions, probability of observing SNPs by chance (Poisson distribution), and the pI/pS calculated are shown.

**Table S7**. Genomic regions not considered for analysis.

**Data S1.** Plots of all trajectories calculated.

