## Supplementary figures and images for "Gene Evolutionary Trajectories in *M. tuberculosis* Reveal Temporal Signs of Selection"

### Rv0104.png

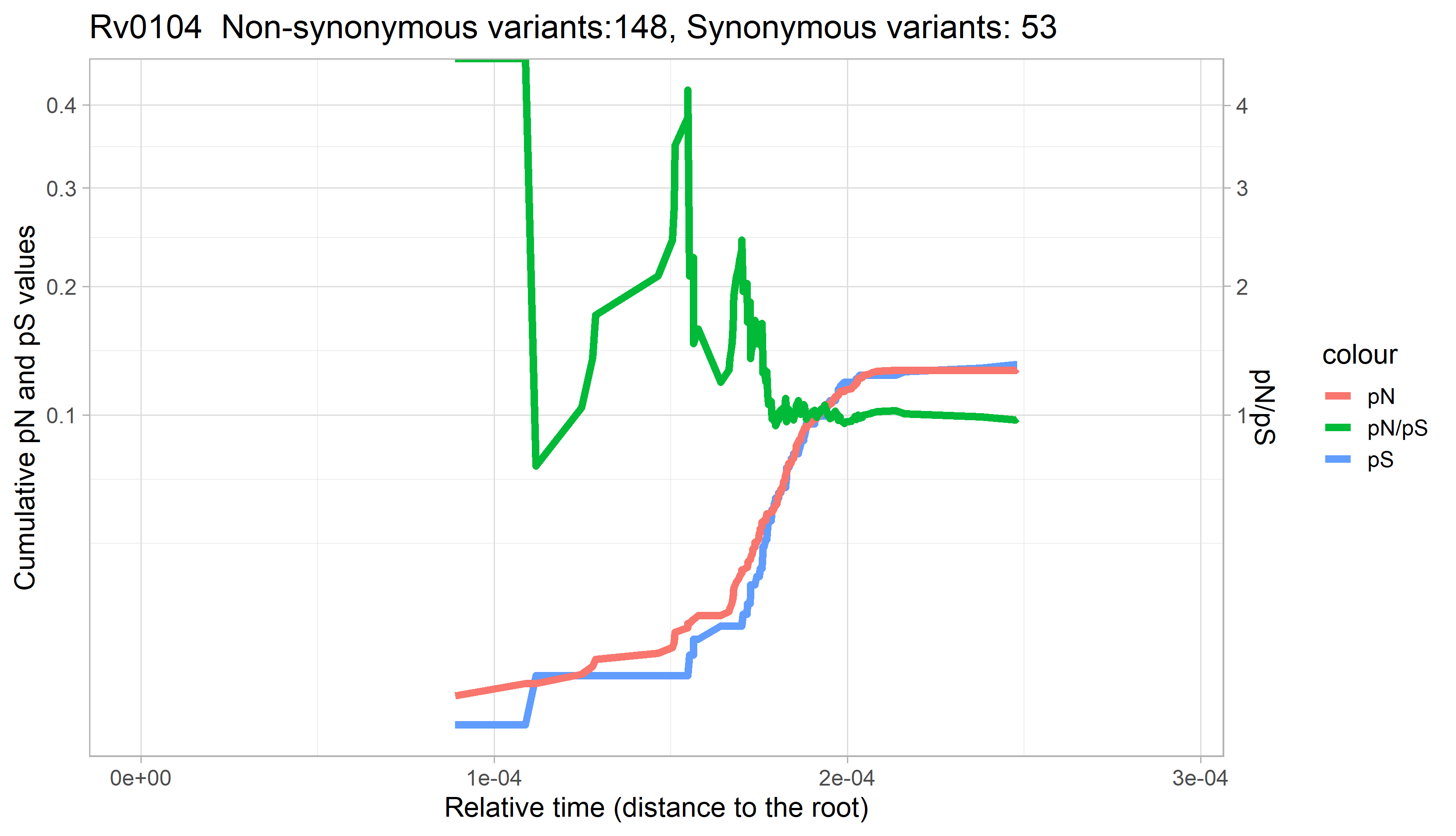

### Rv0105c.png

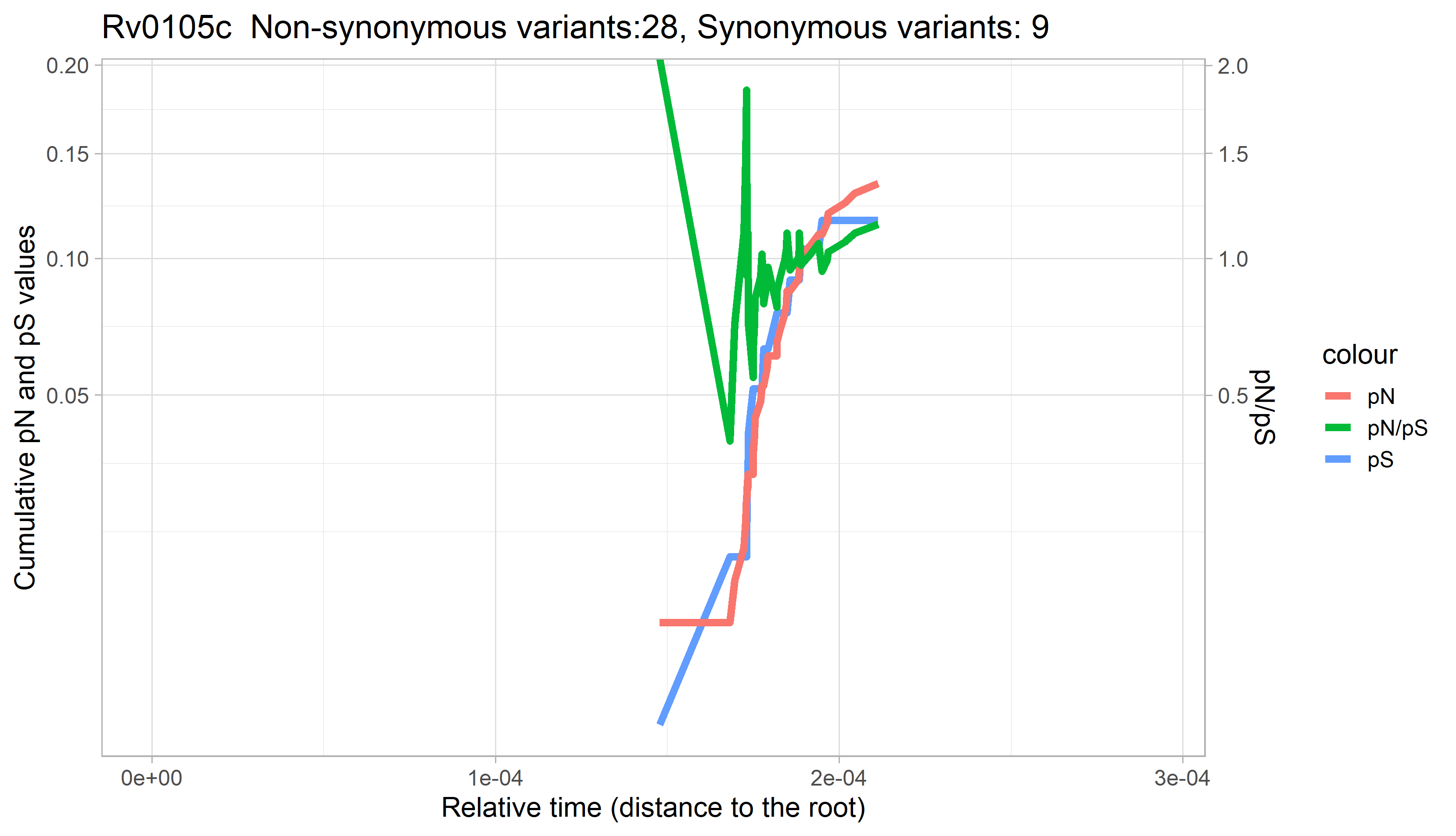

### Rv0106.png

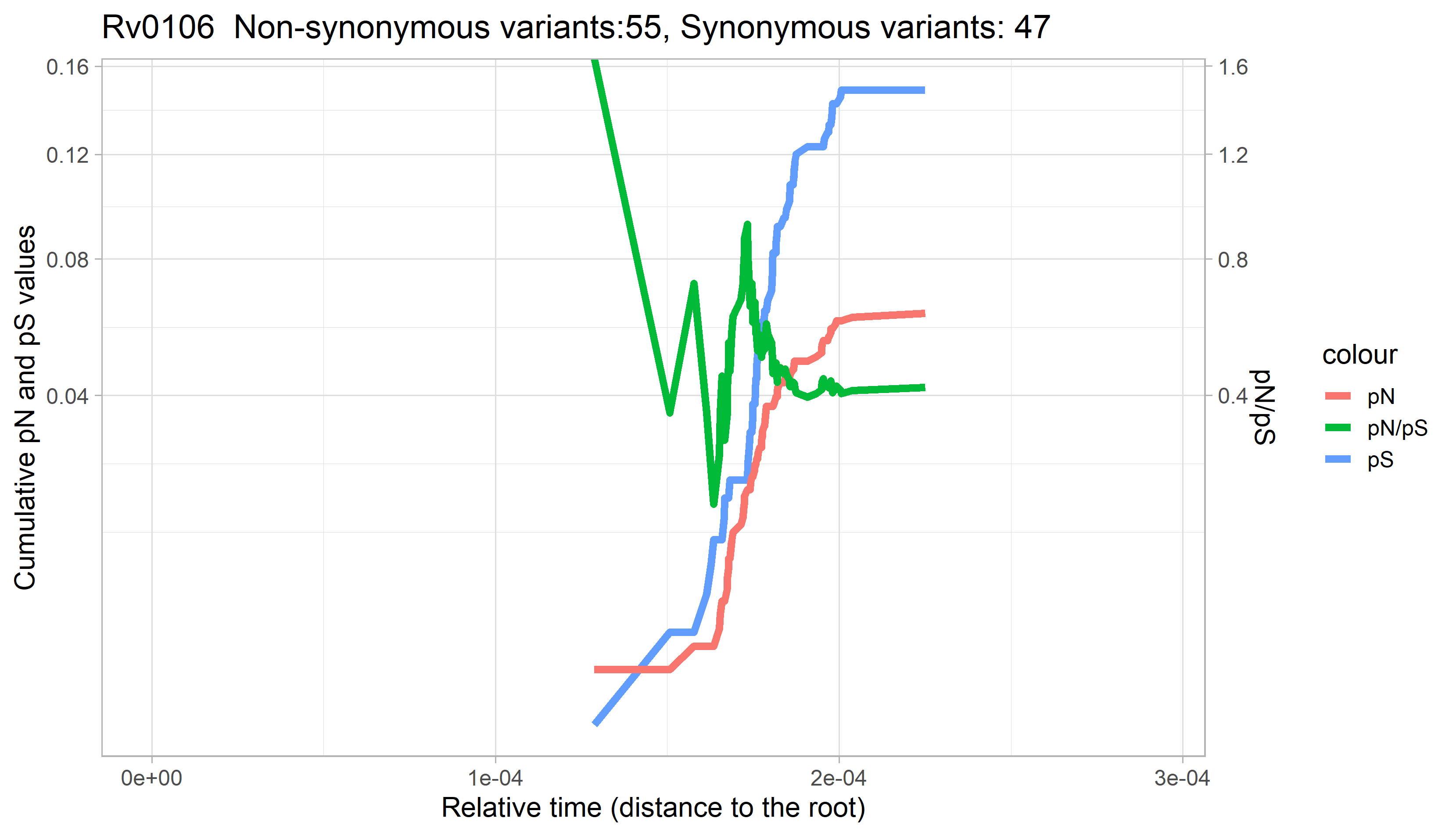

### Rv0107c.png

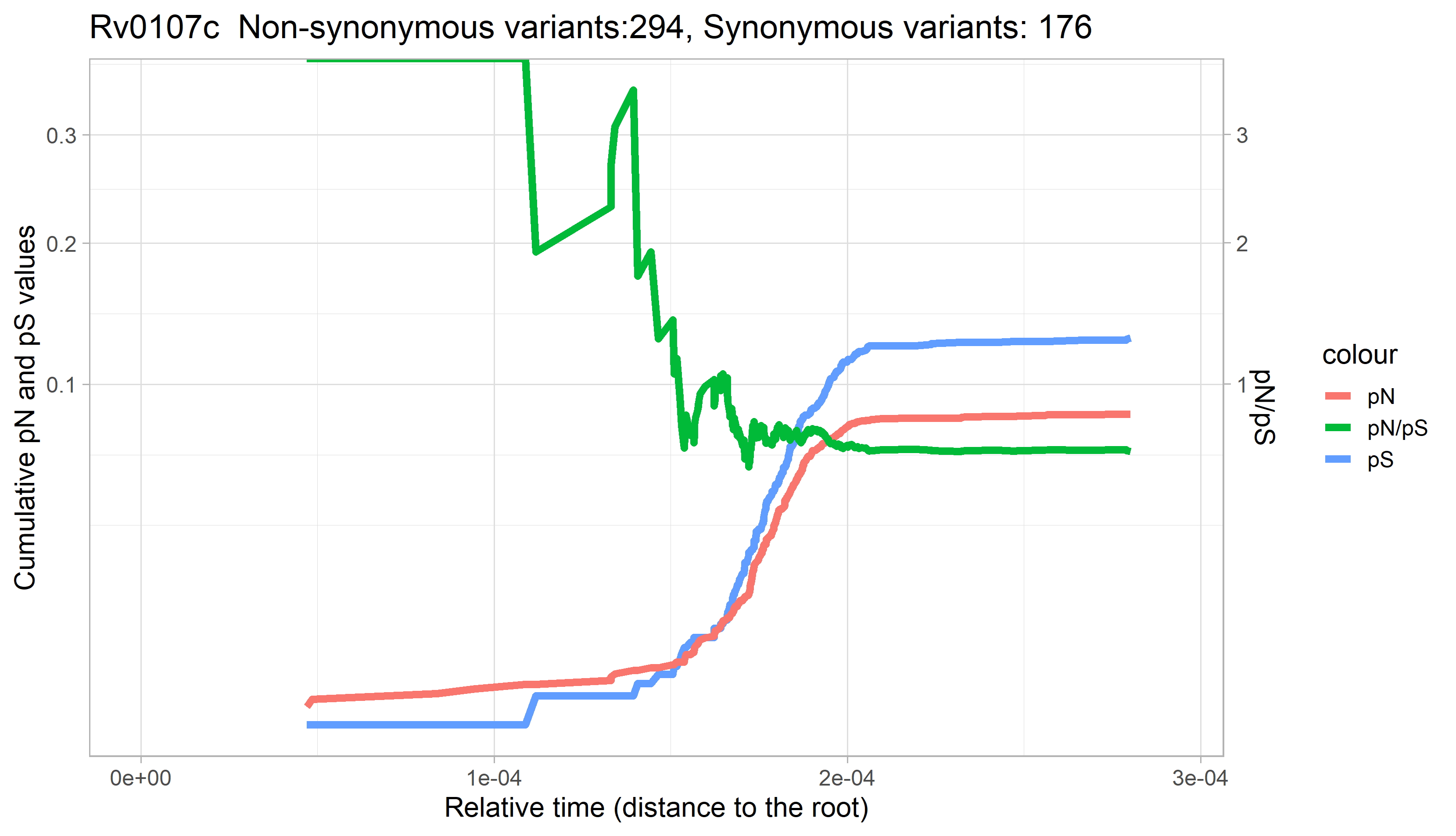

### Rv0108c.png

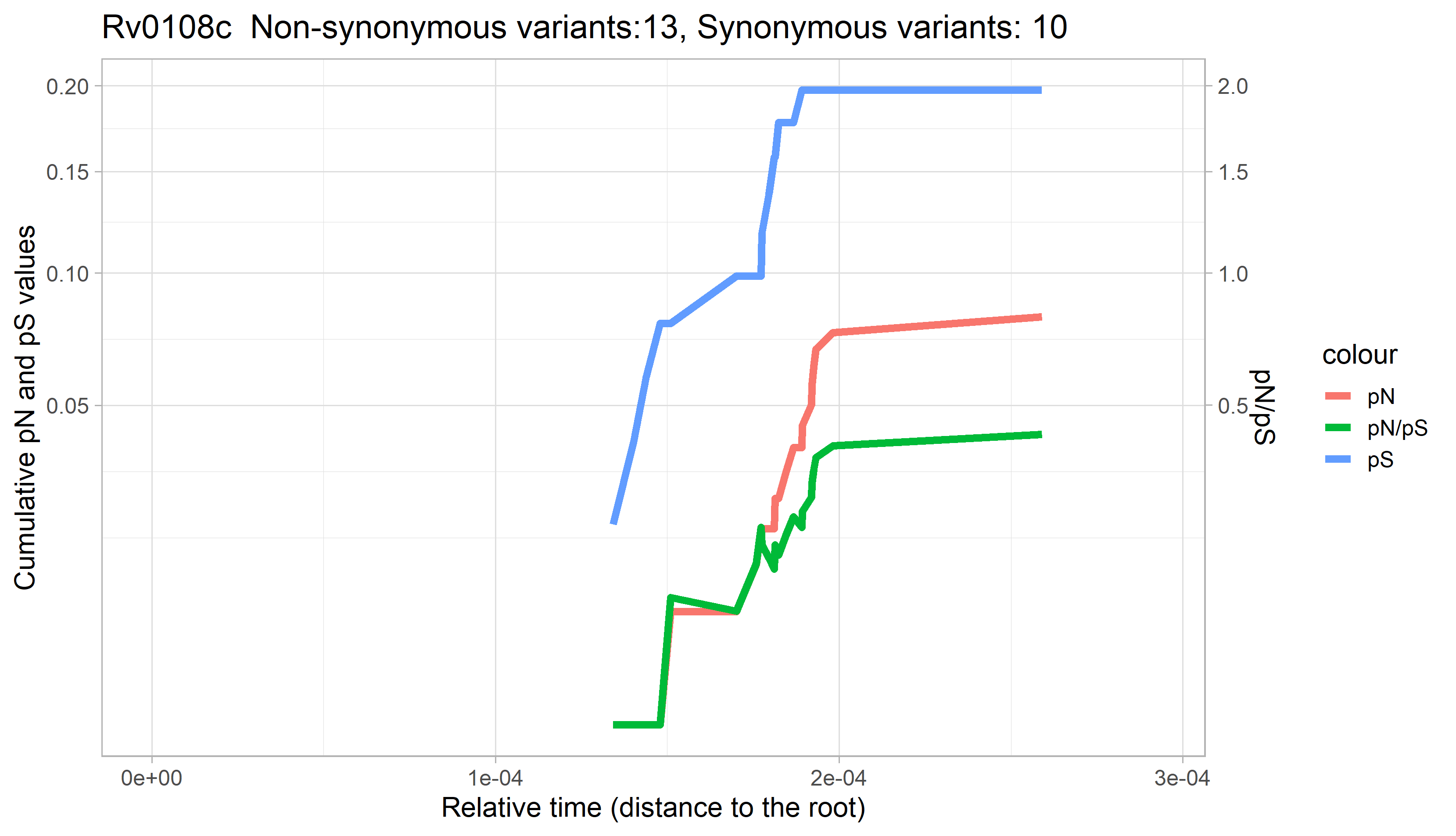

### Rv0110.png

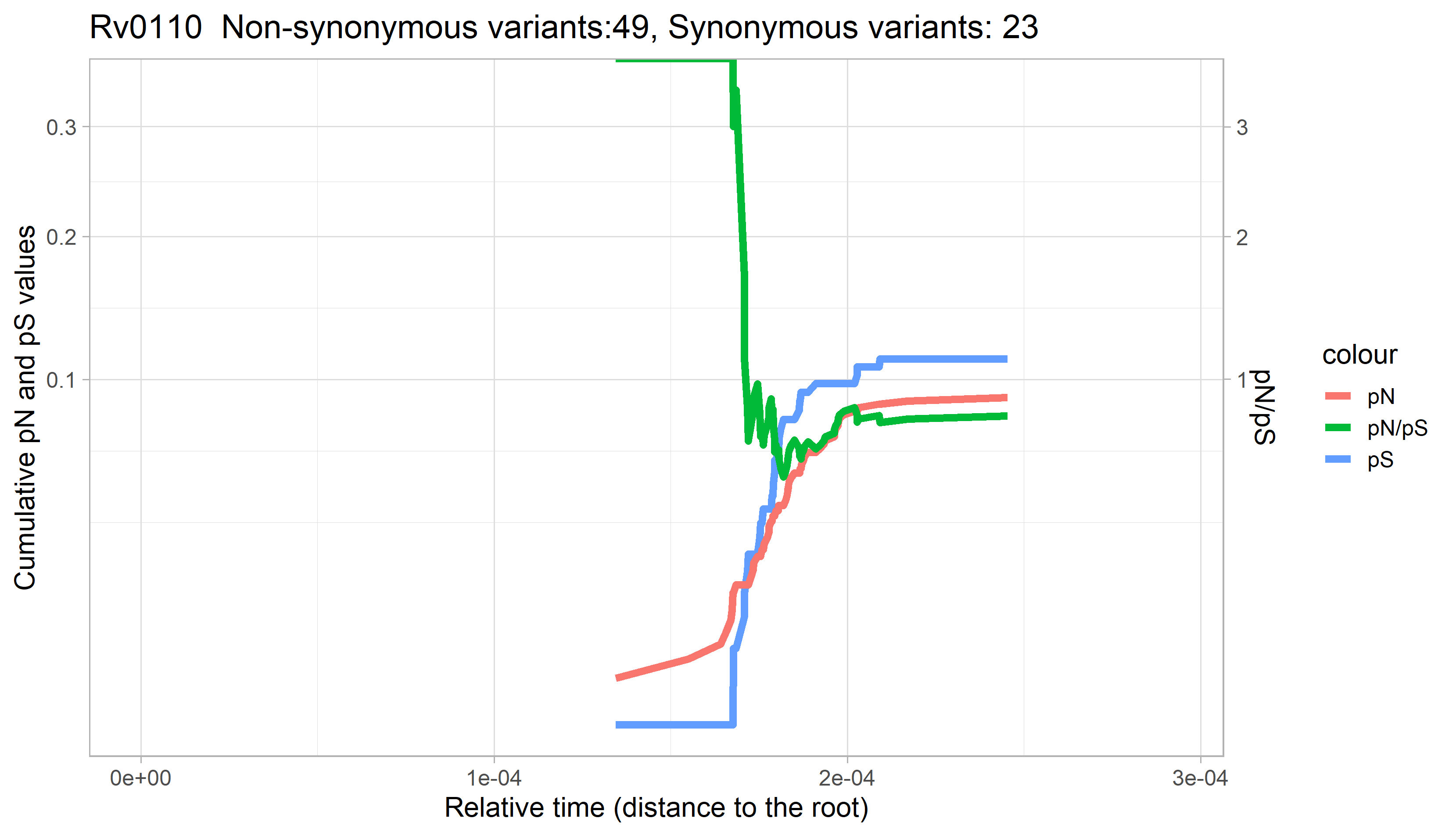

### Rv0111.png

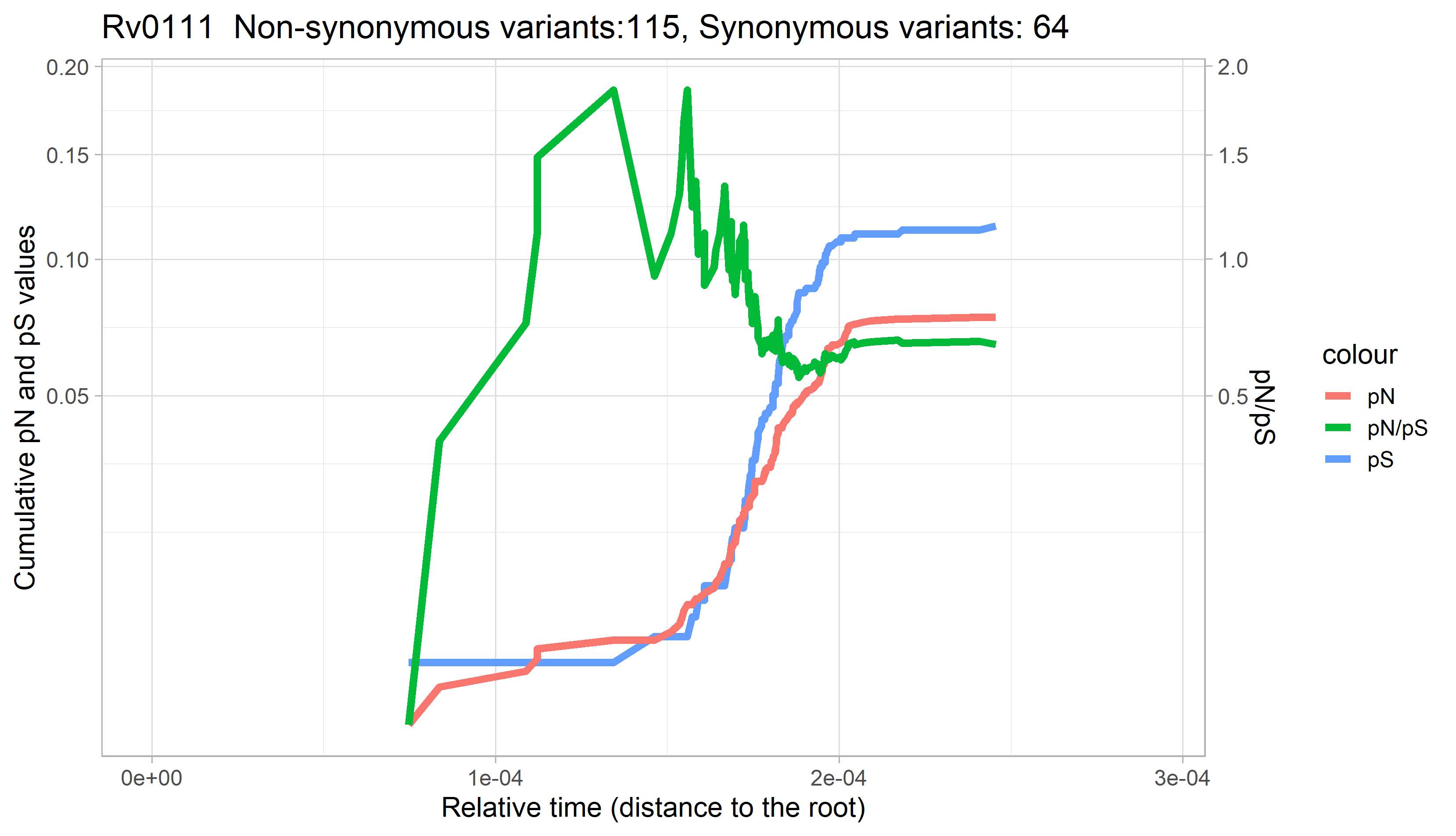

### Rv0112.png

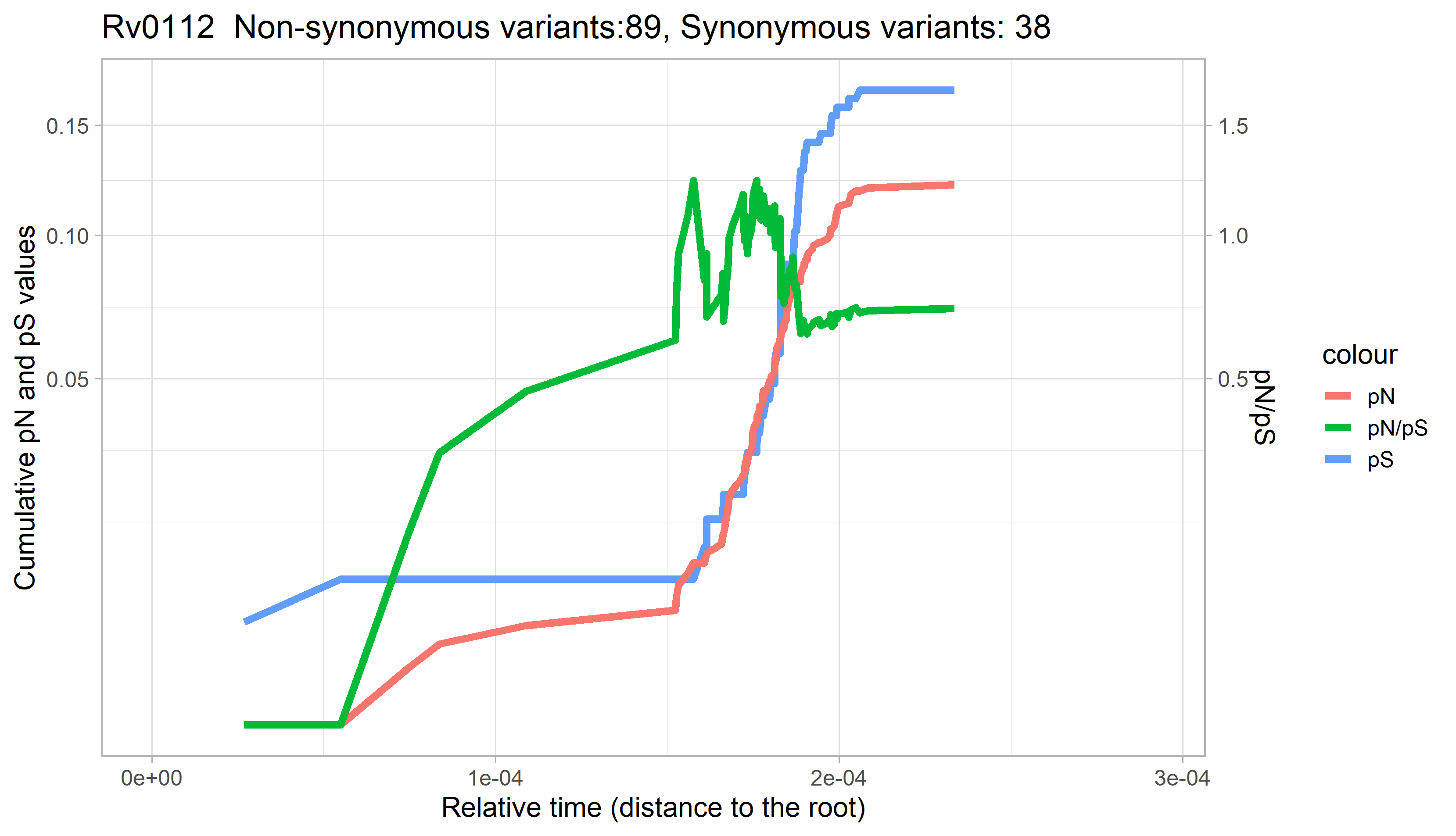

### Rv0113.png

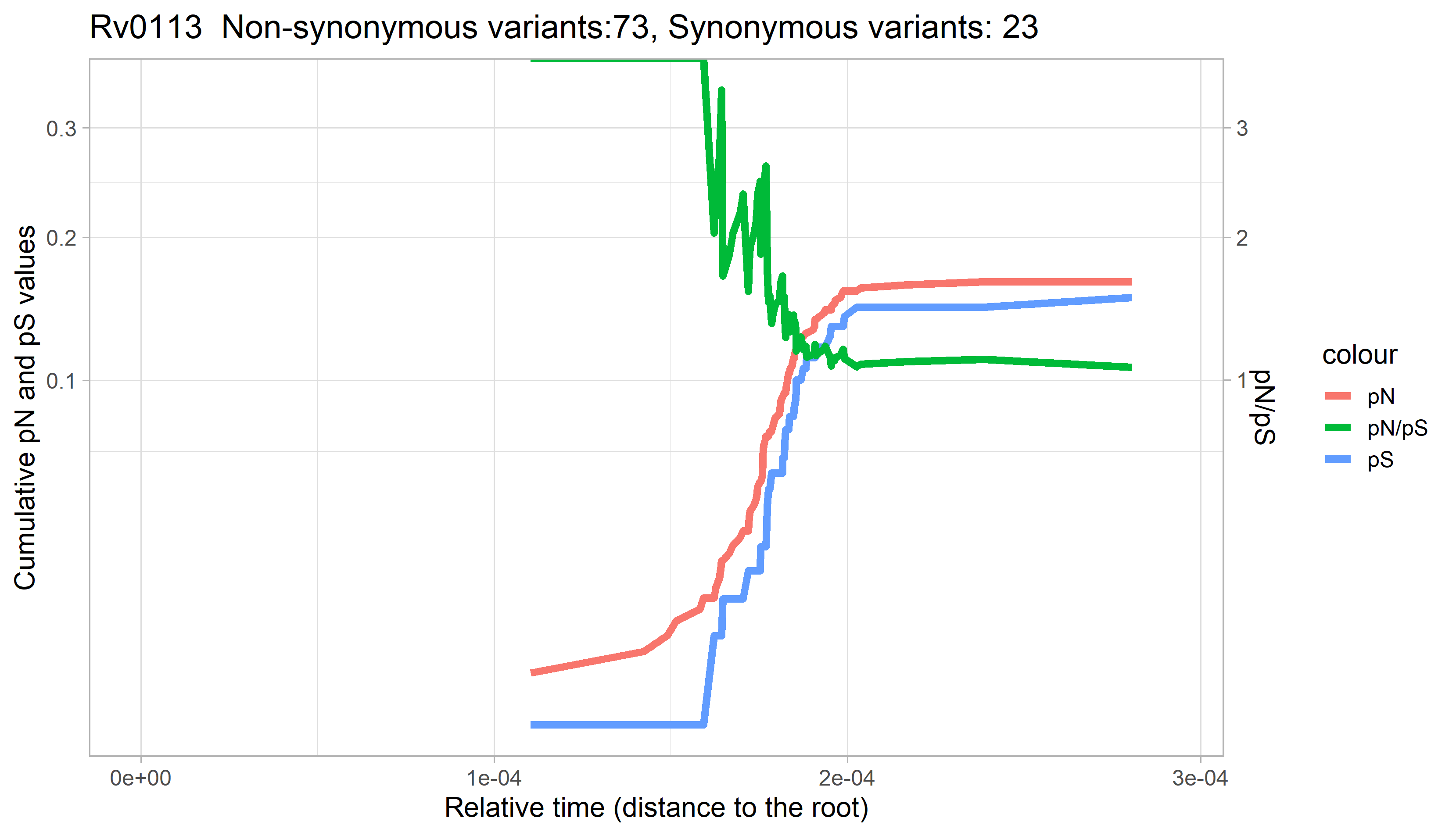

### Rv0114.png

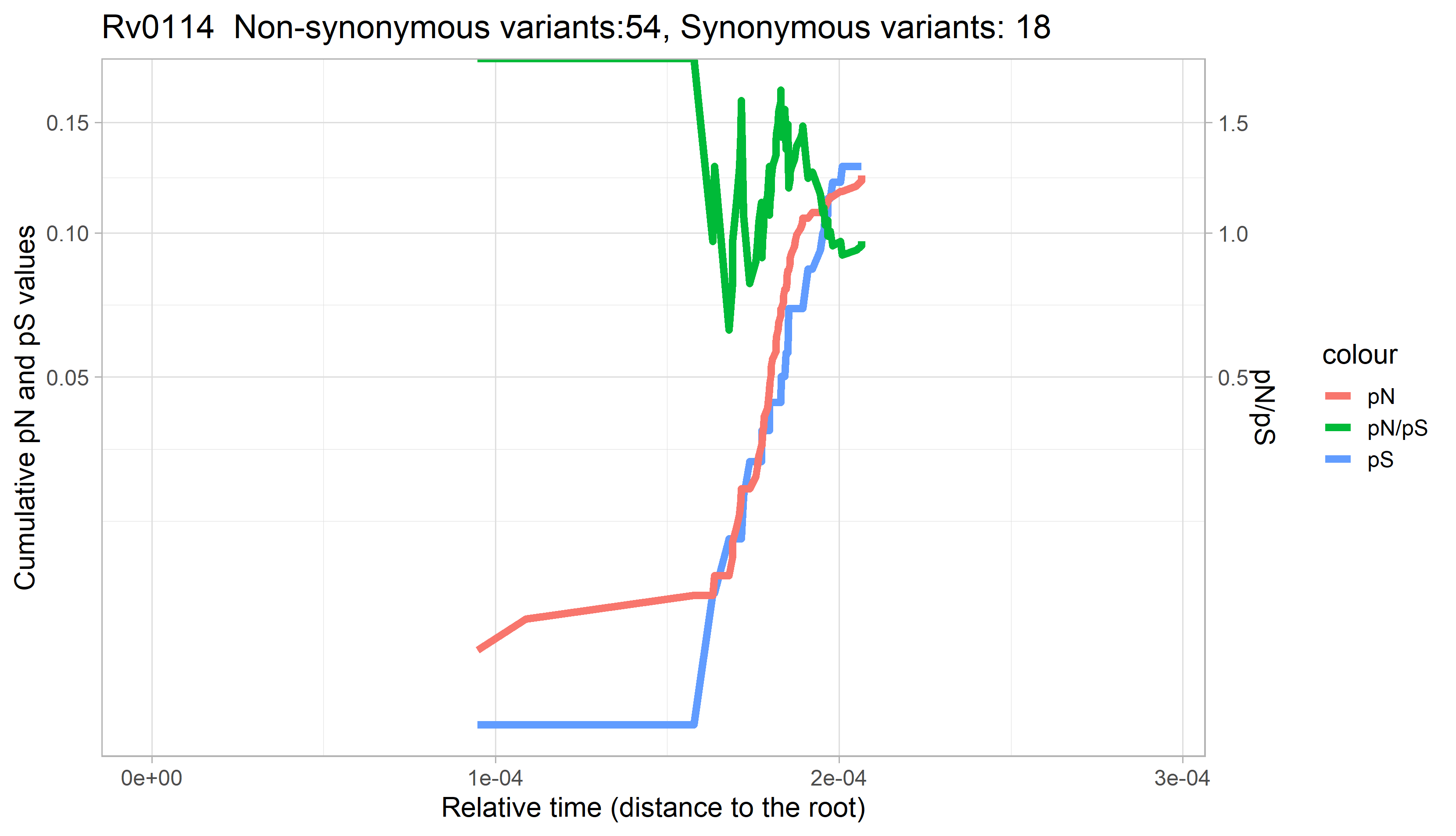

### Rv0115.png

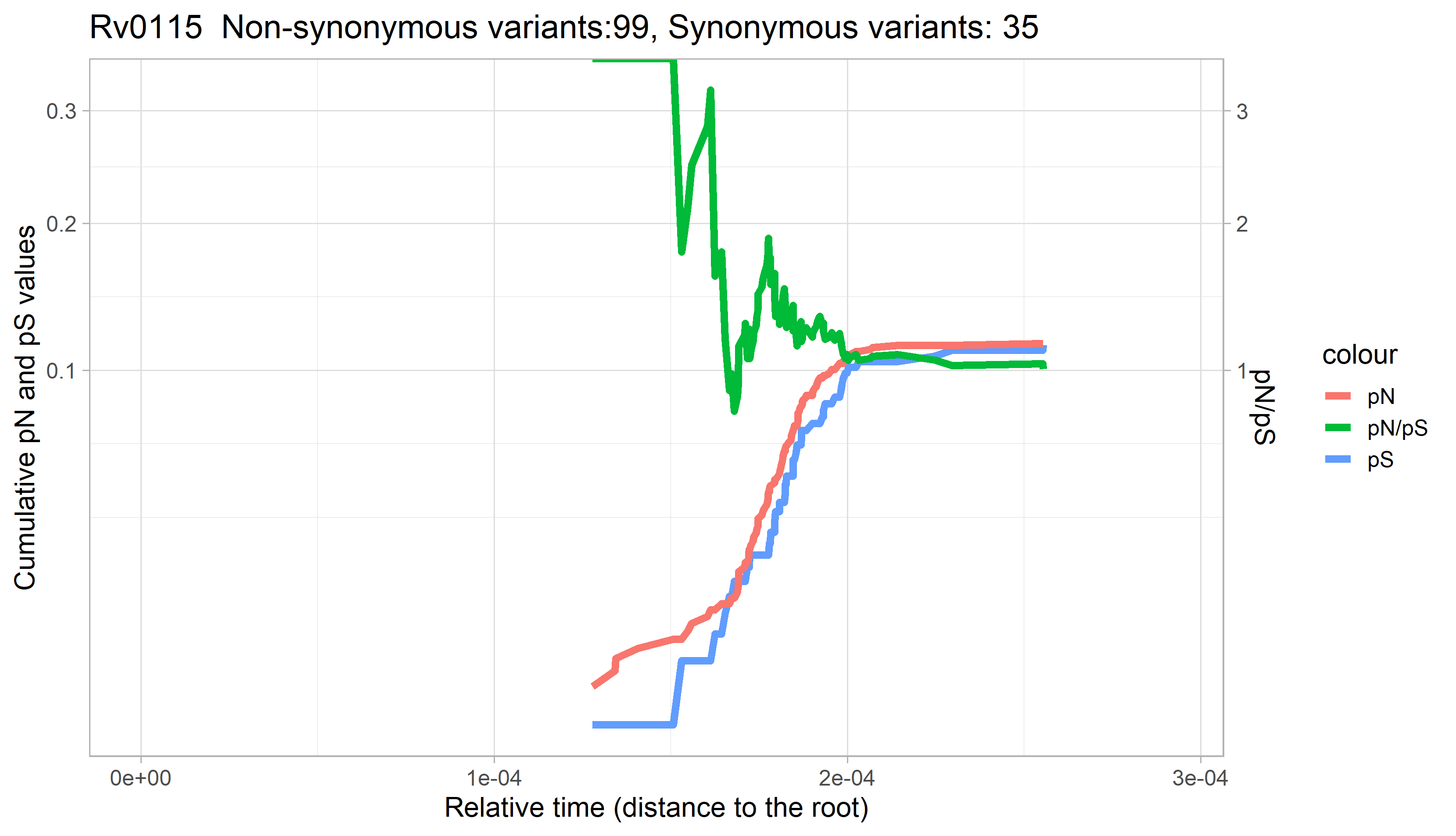

### Rv0116c.png

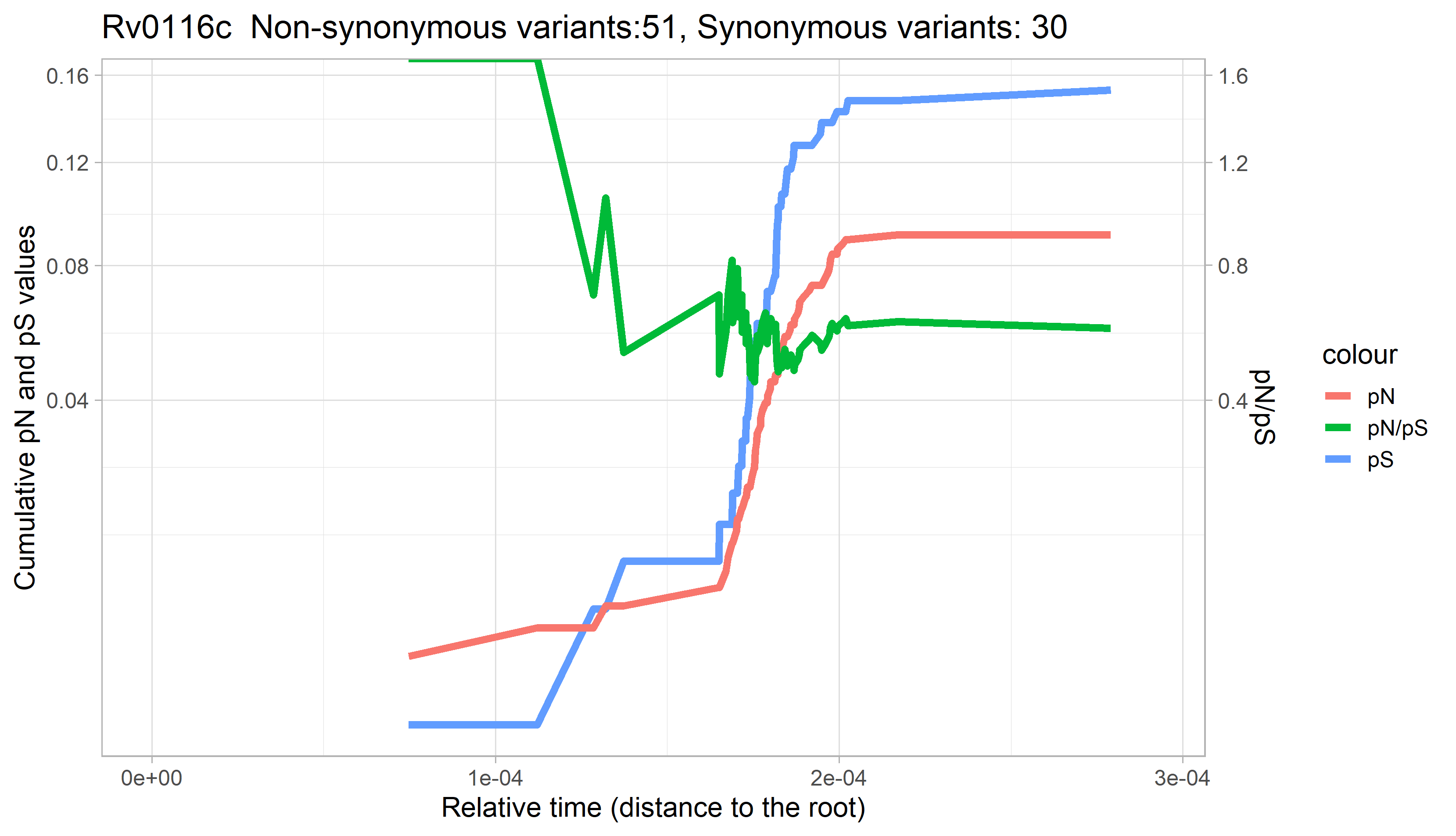

### Rv0117.png

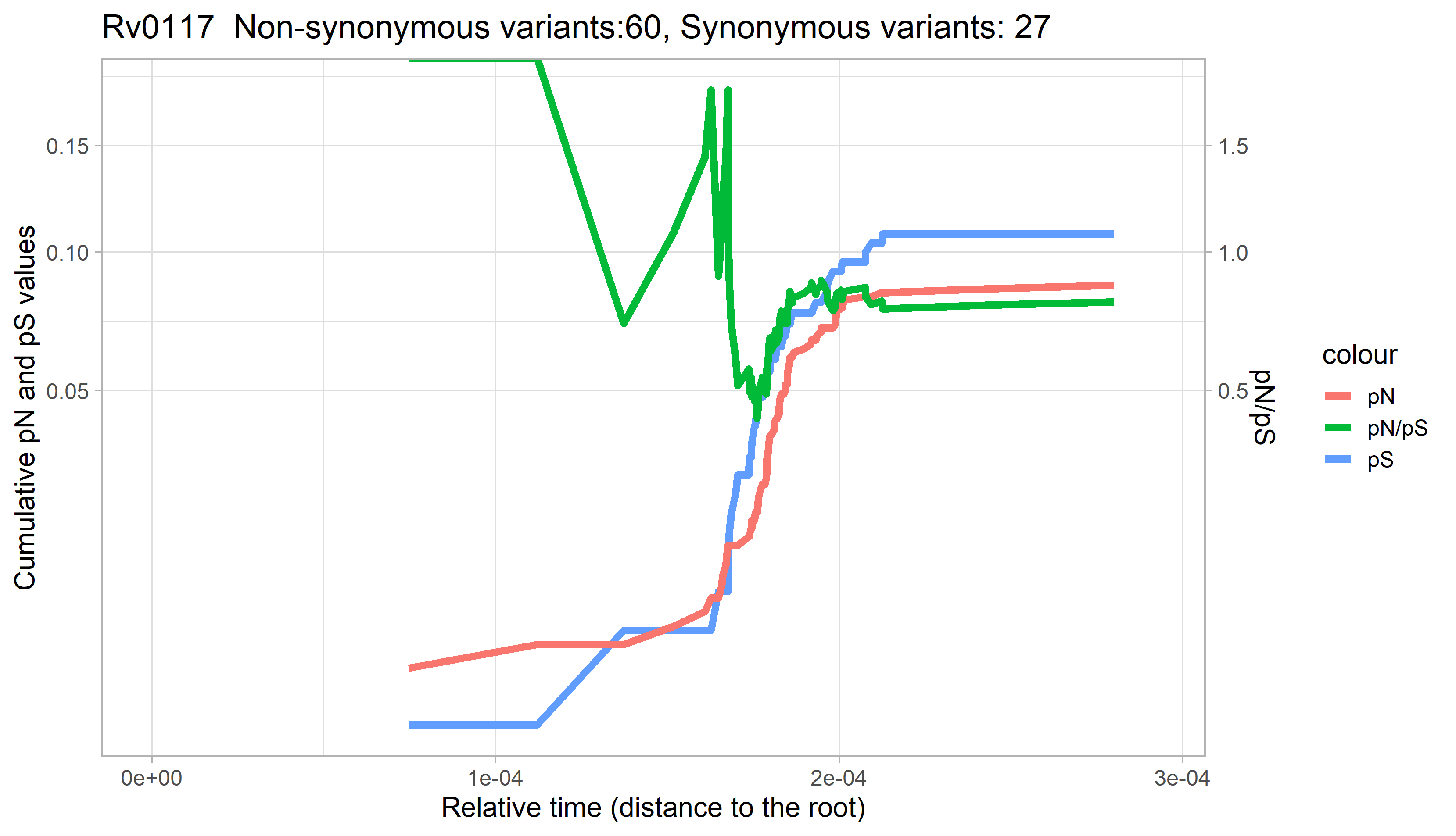

### Rv0118c.png

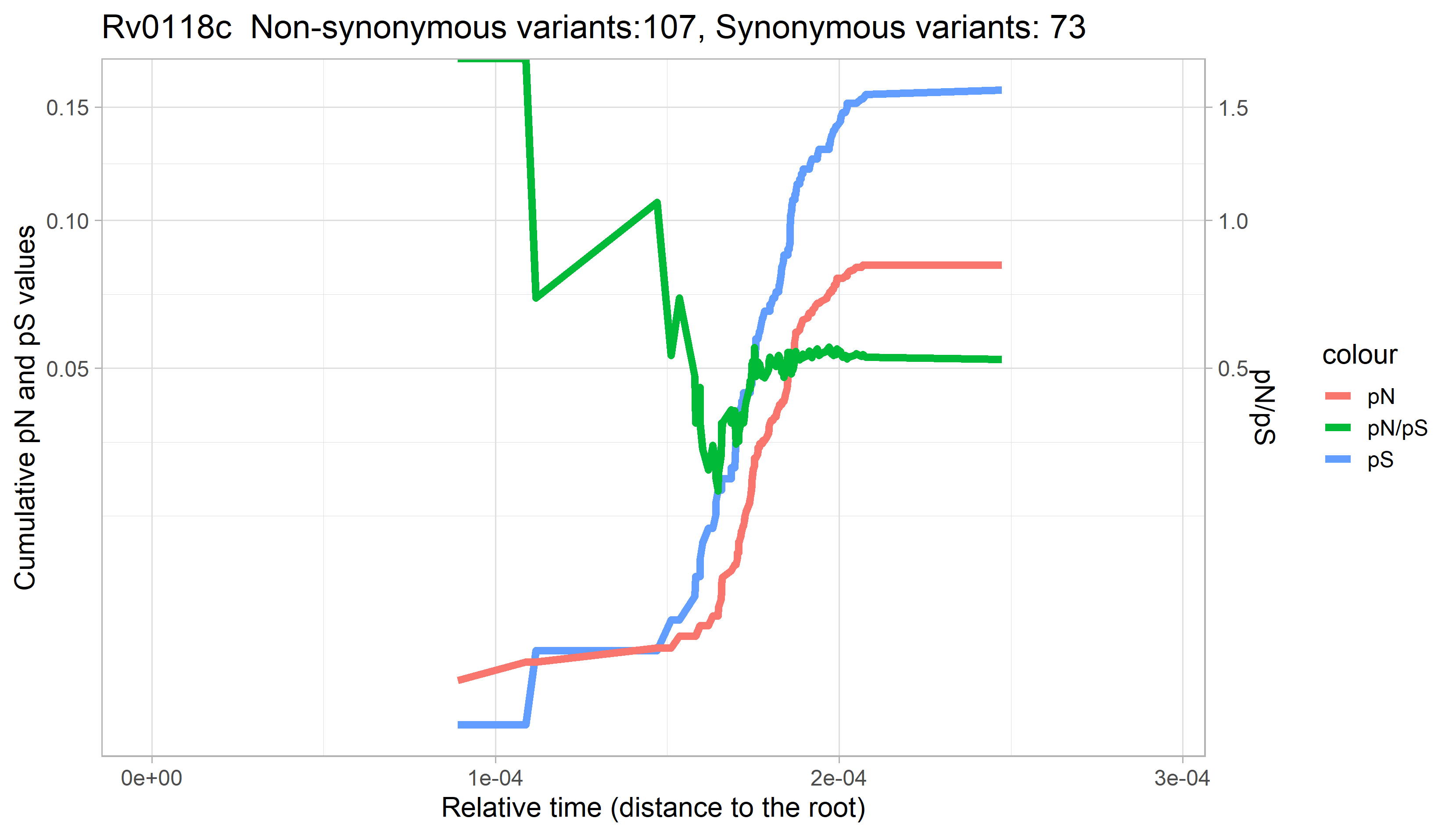

### Rv0119.png

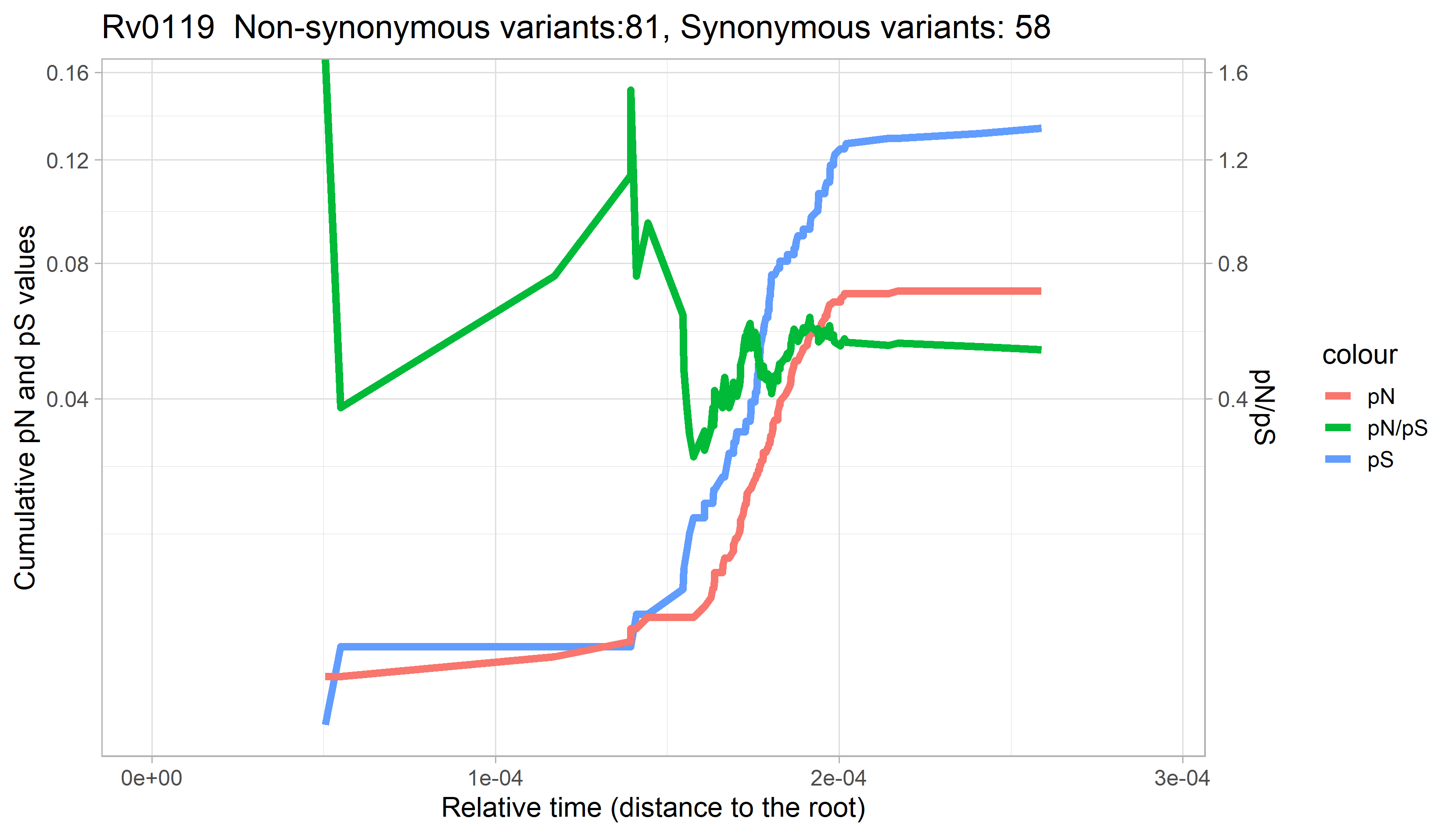

### Rv0120c.png

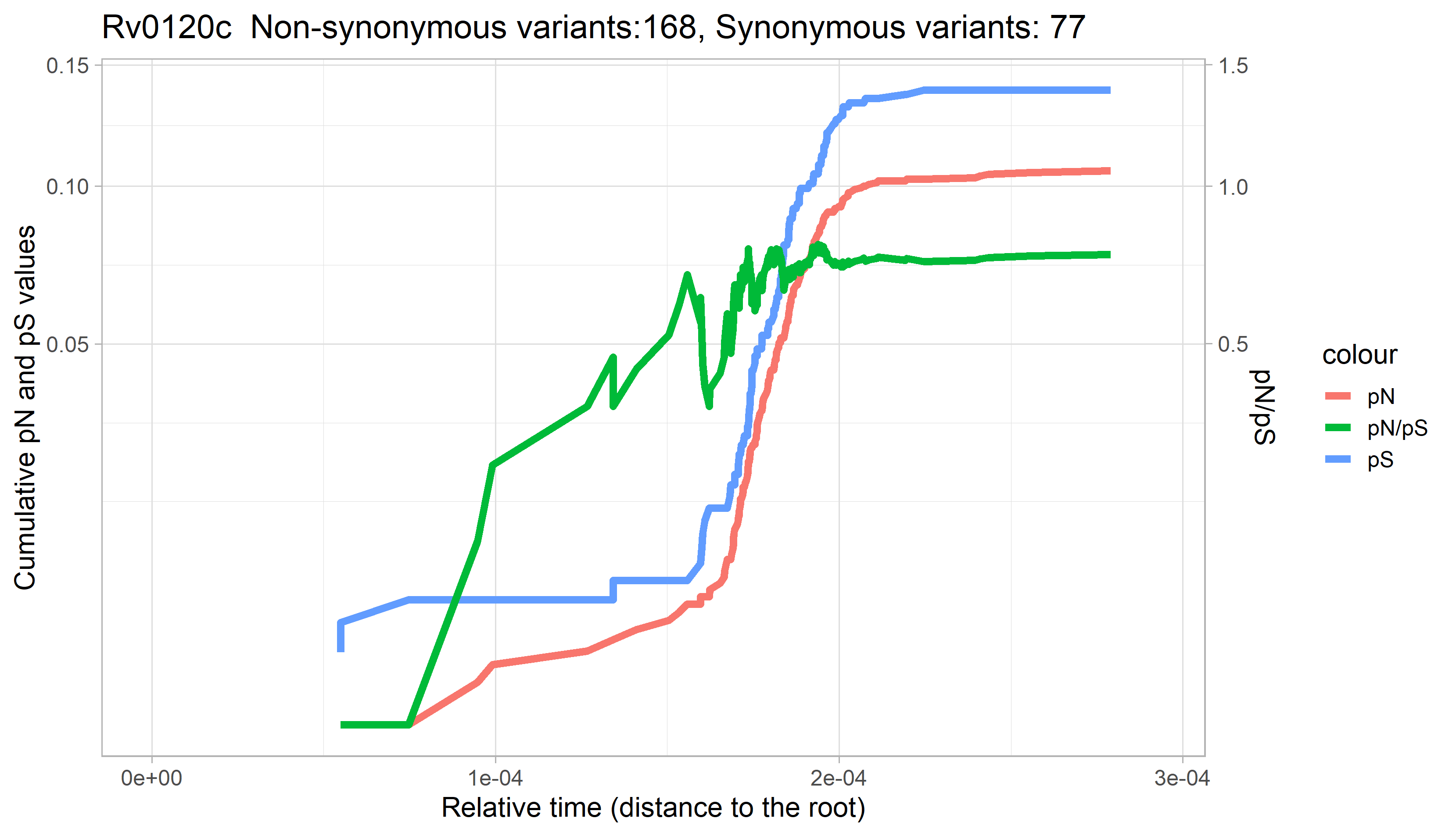

### Rv0121c.png

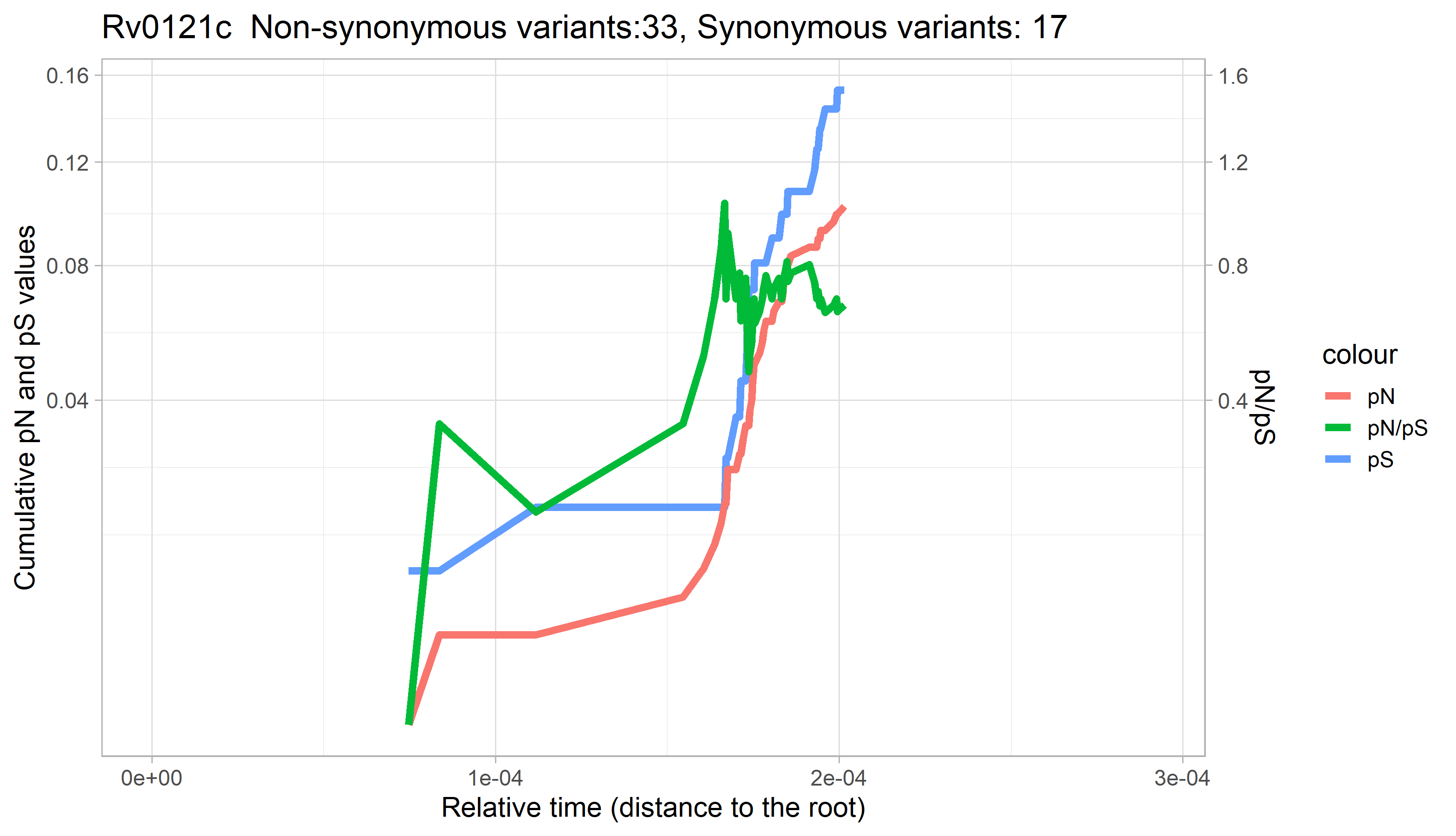

### Rv0122.png

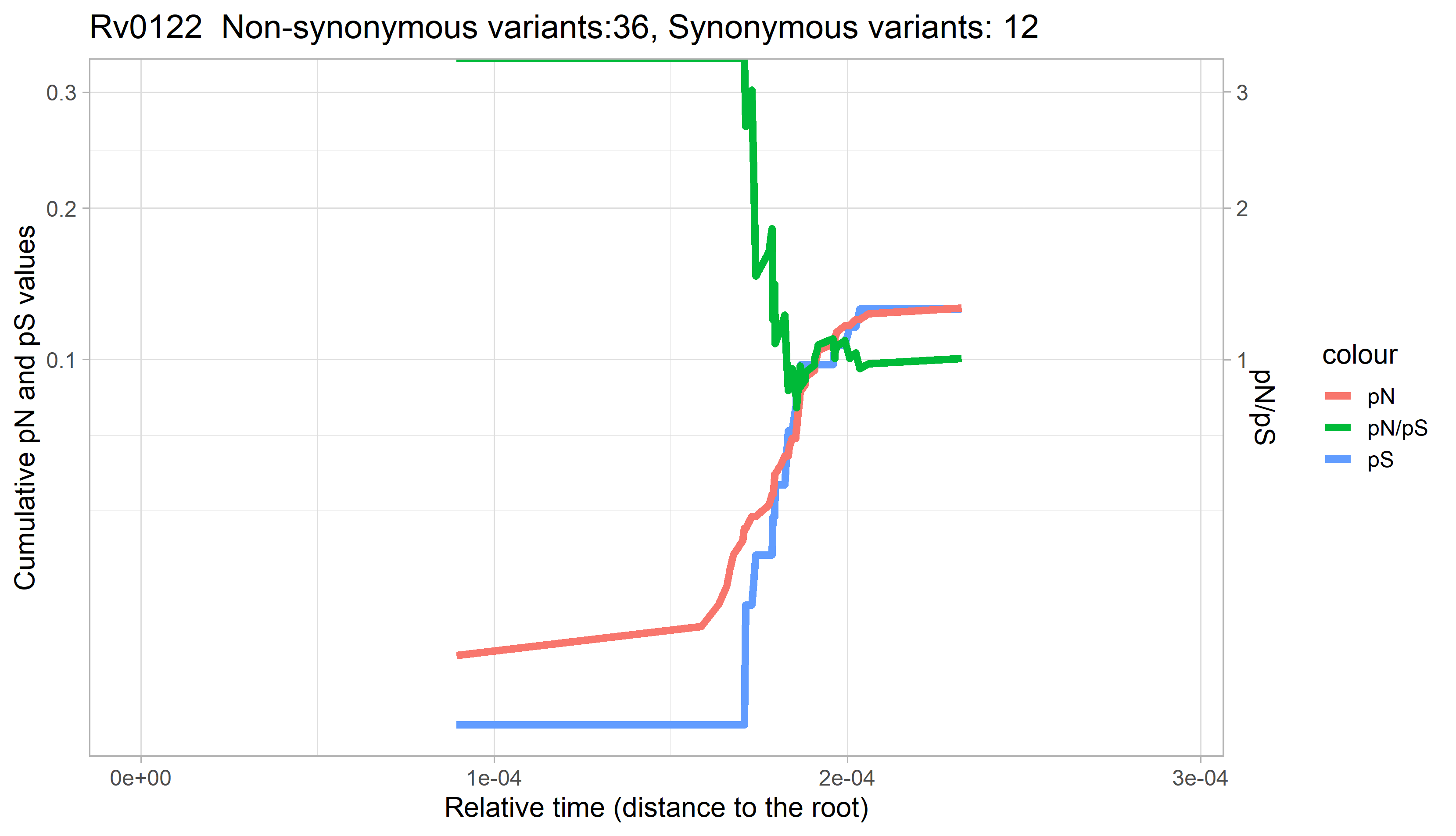

### Rv0123.png

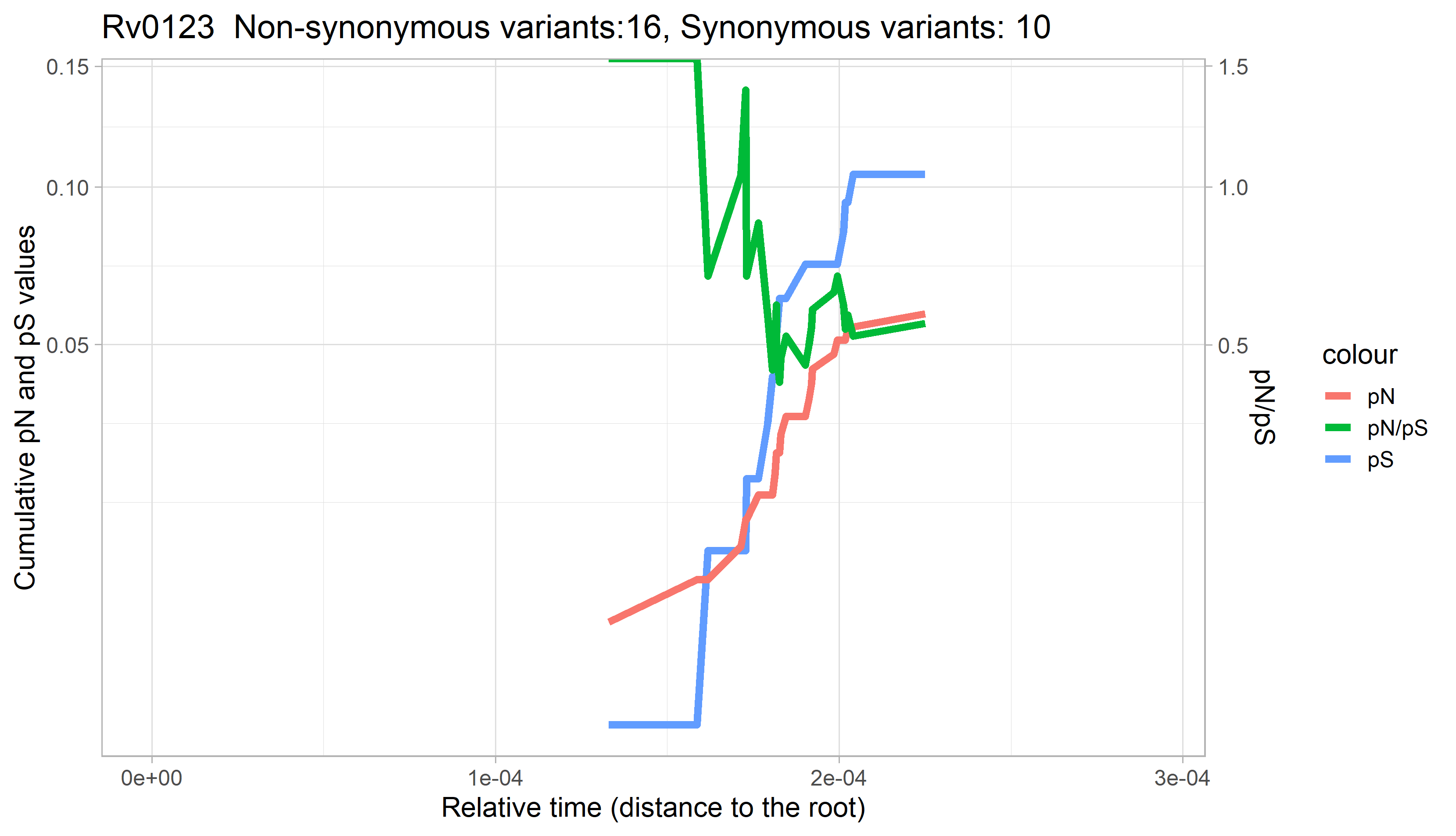

### Rv0125.png

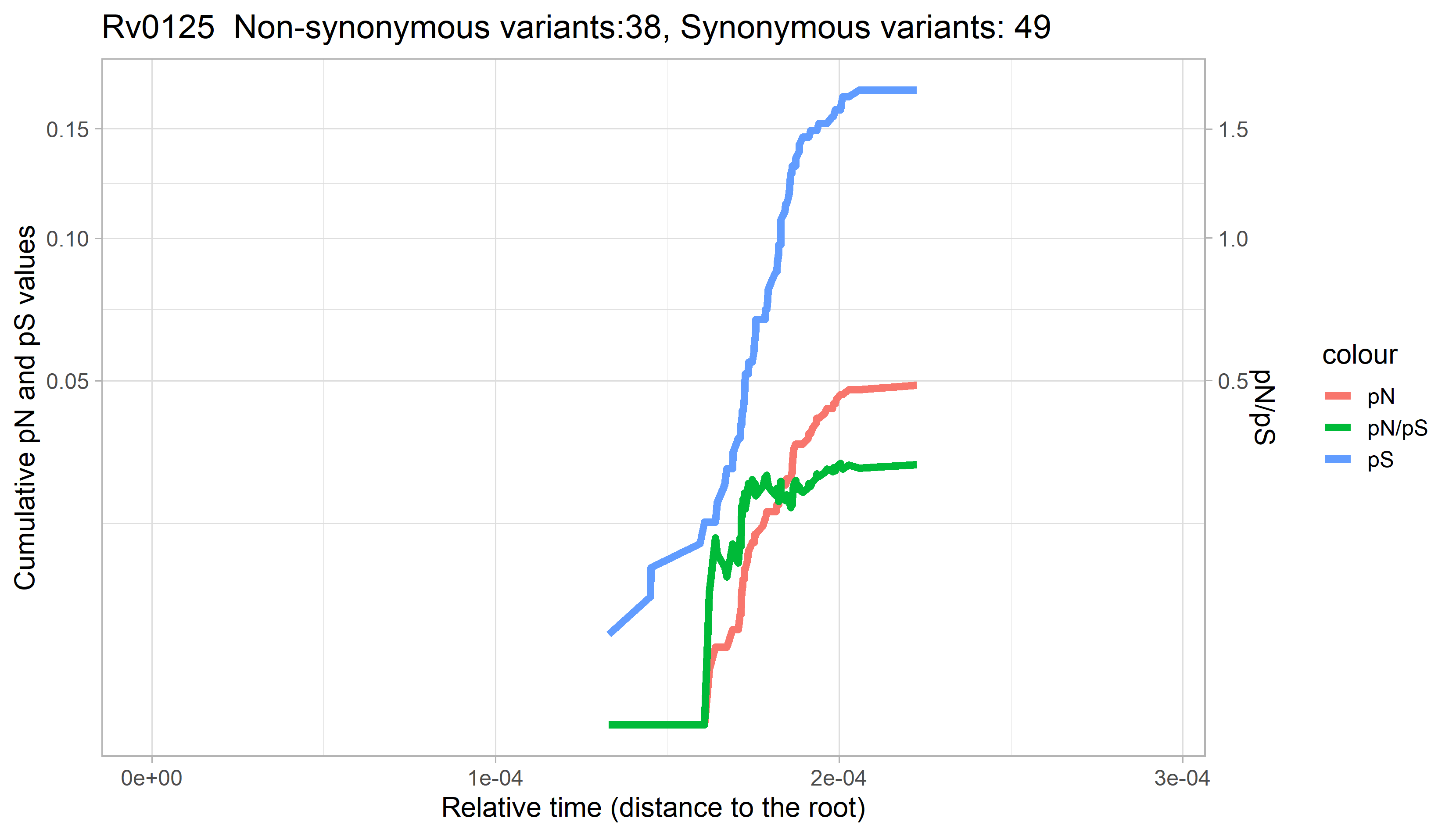

### Rv0126.png

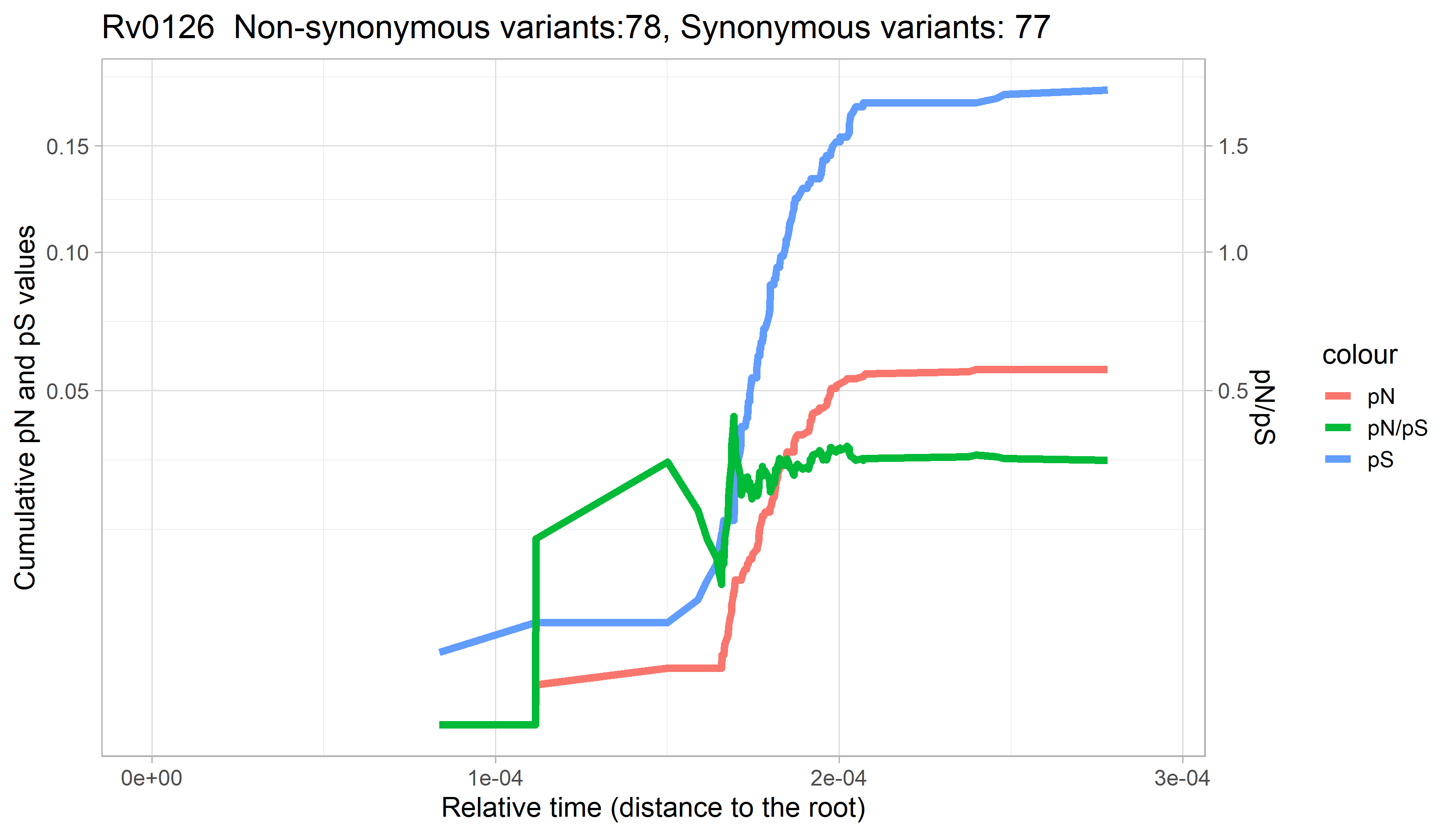

### Rv0127.png

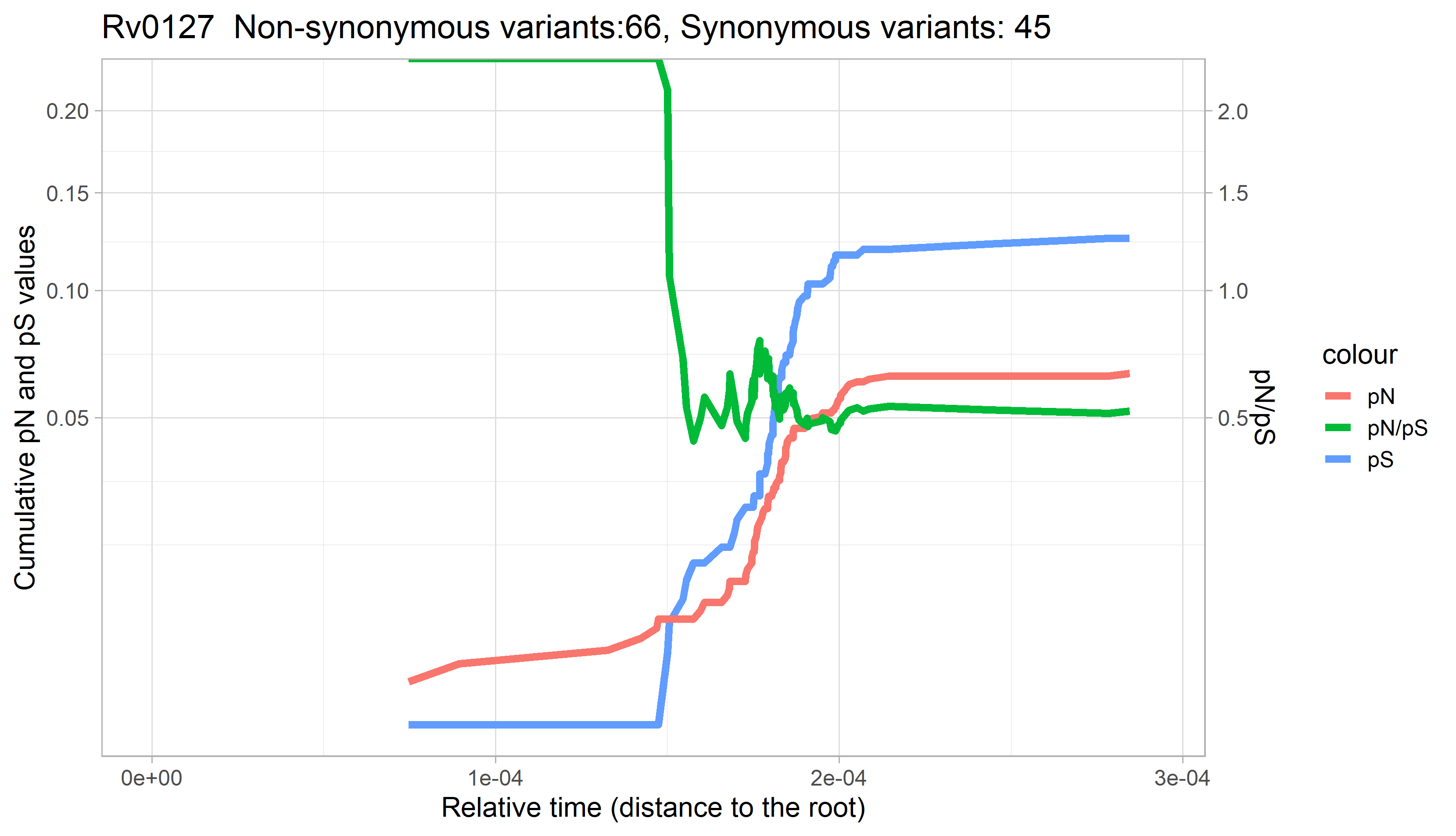

### Rv0128.png

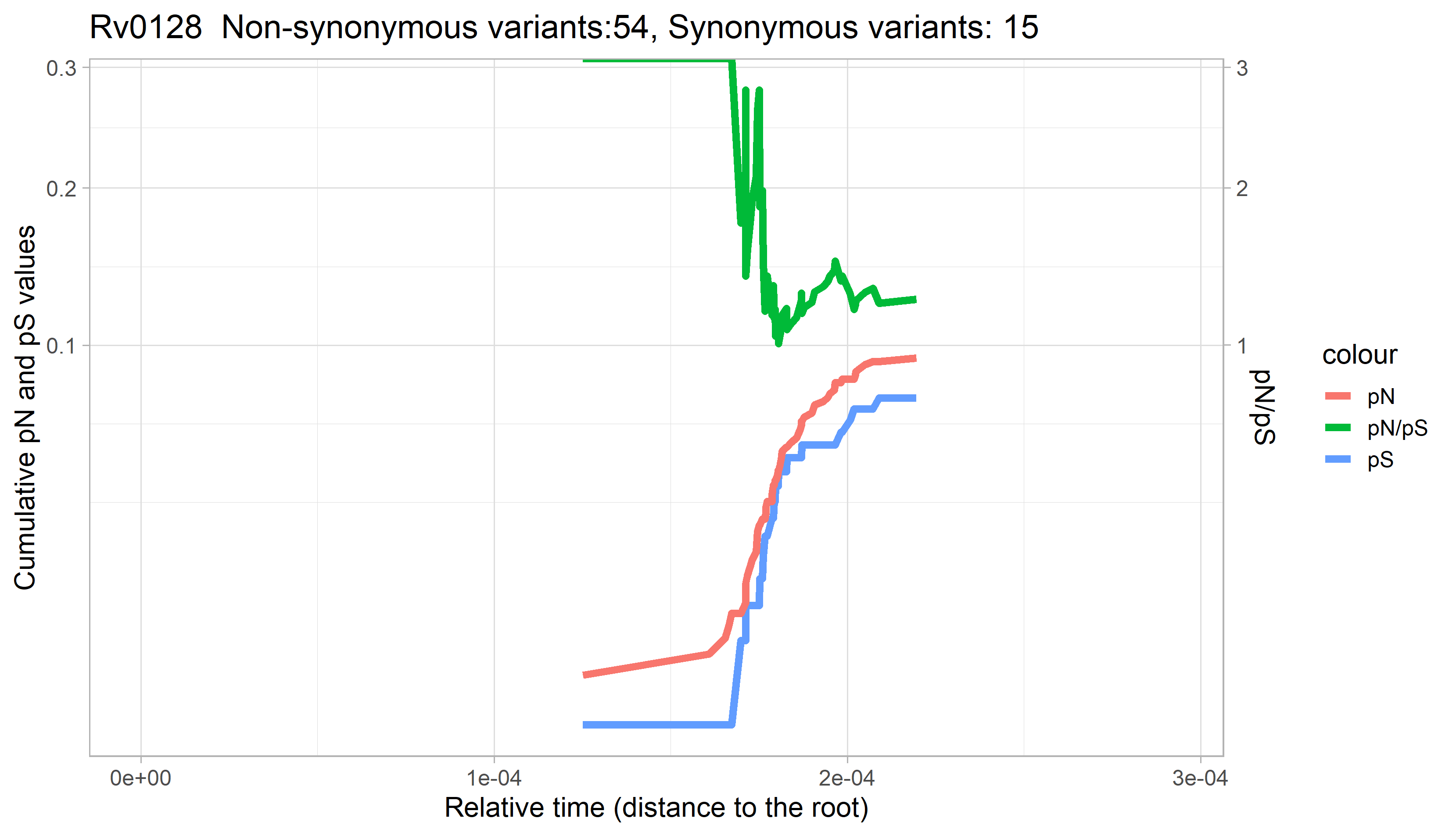

### Rv0129c.png

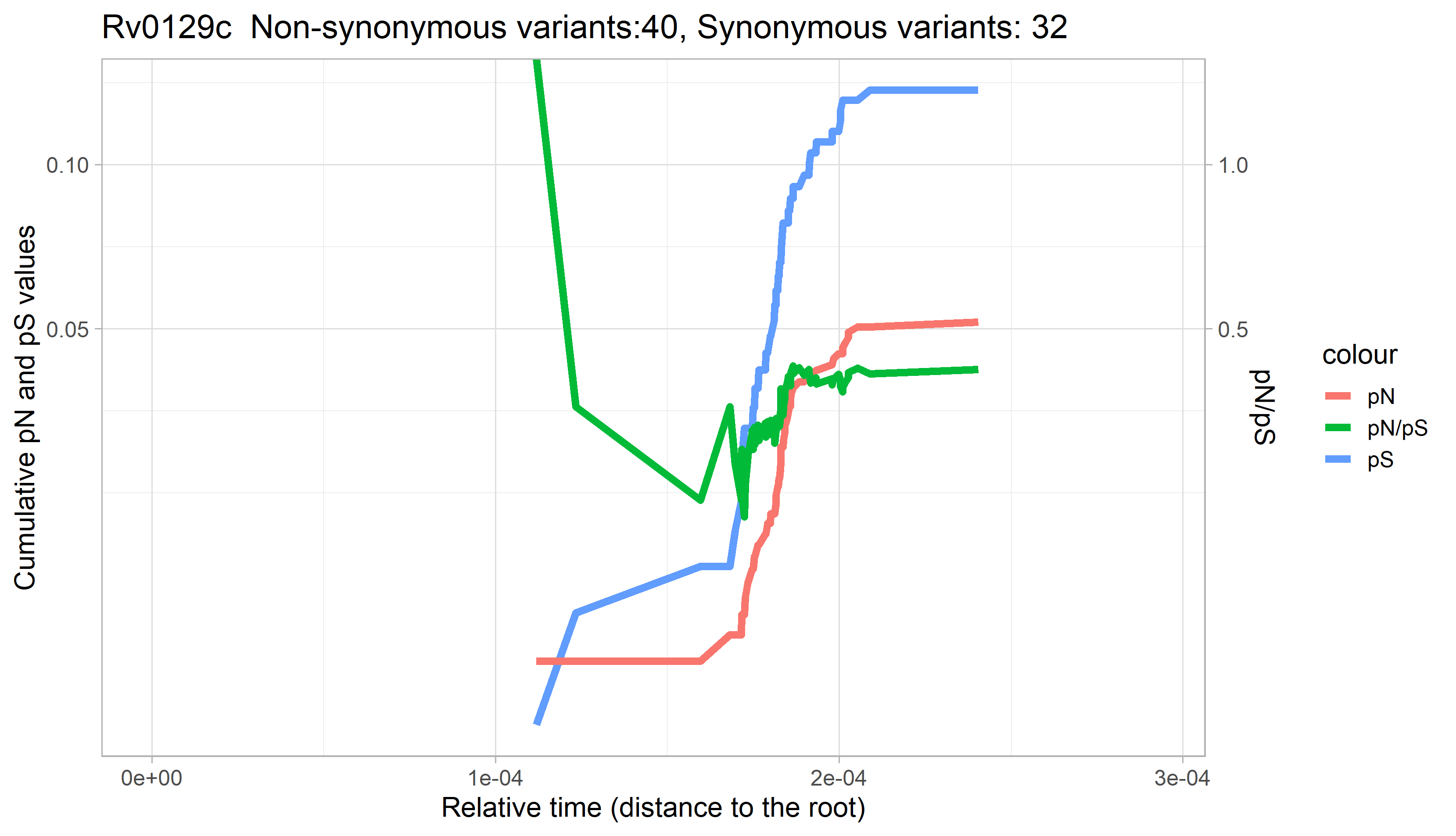

### Rv0130.png

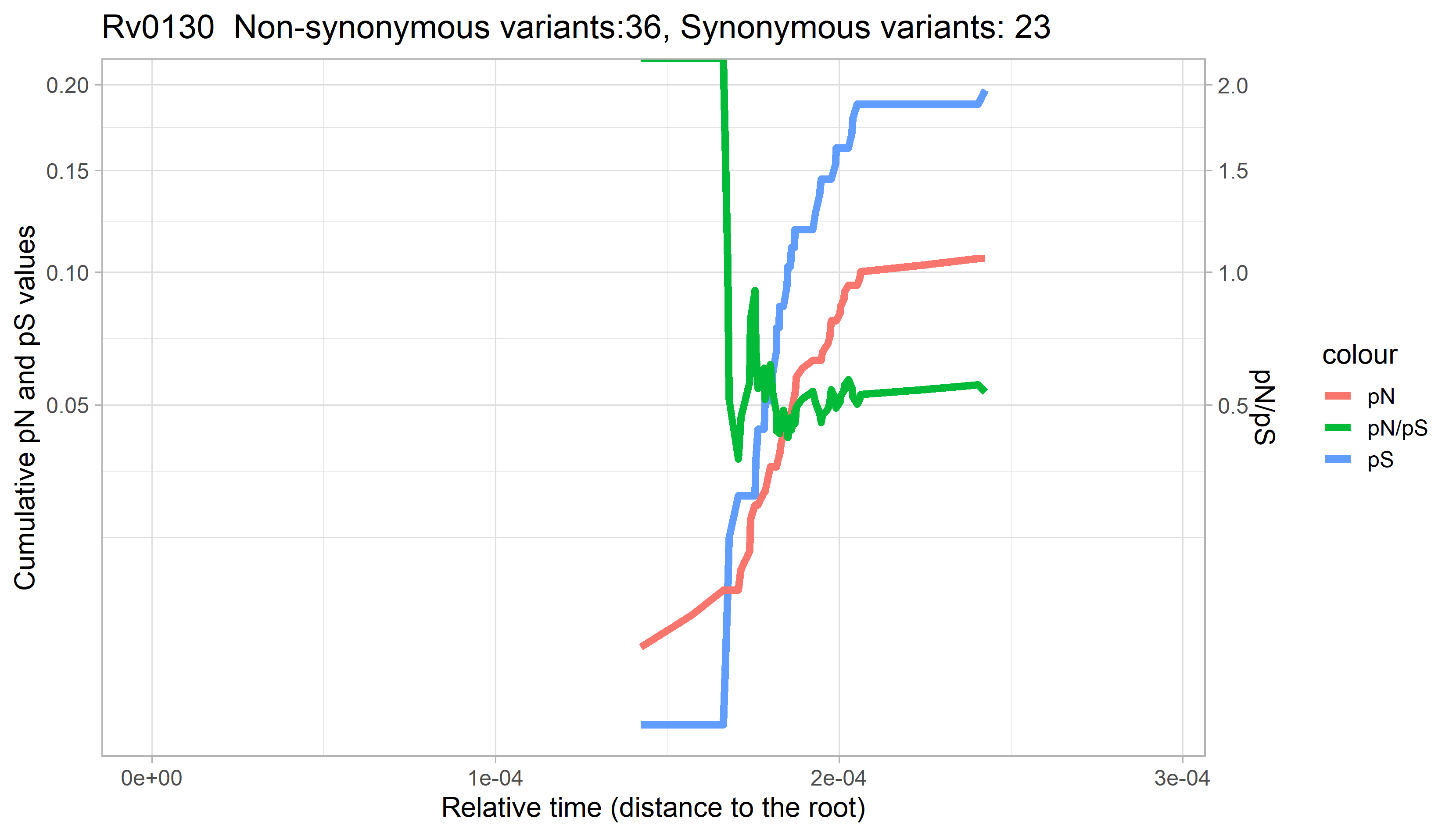

### Rv0131c.png

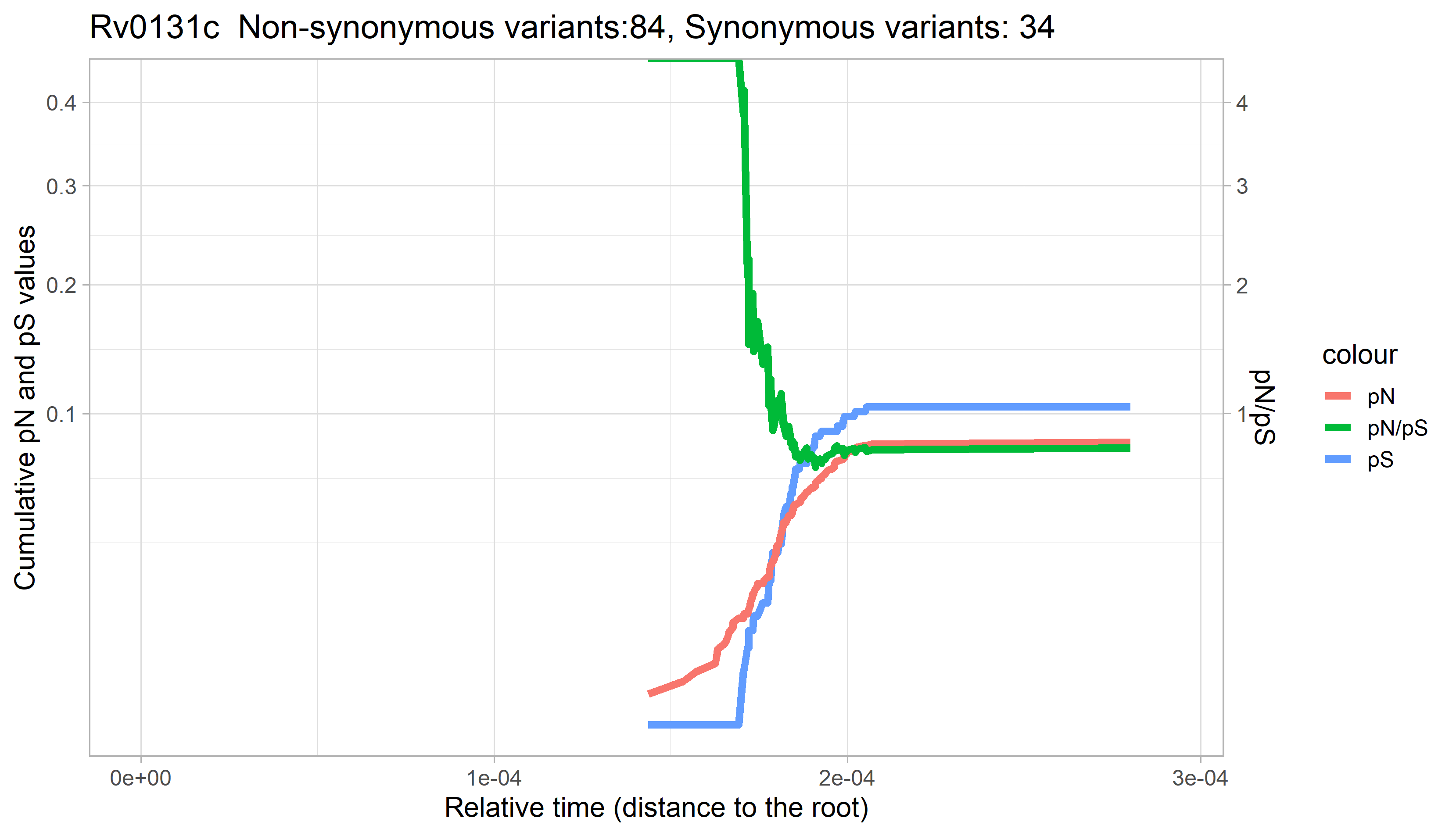

### Rv0132c.png

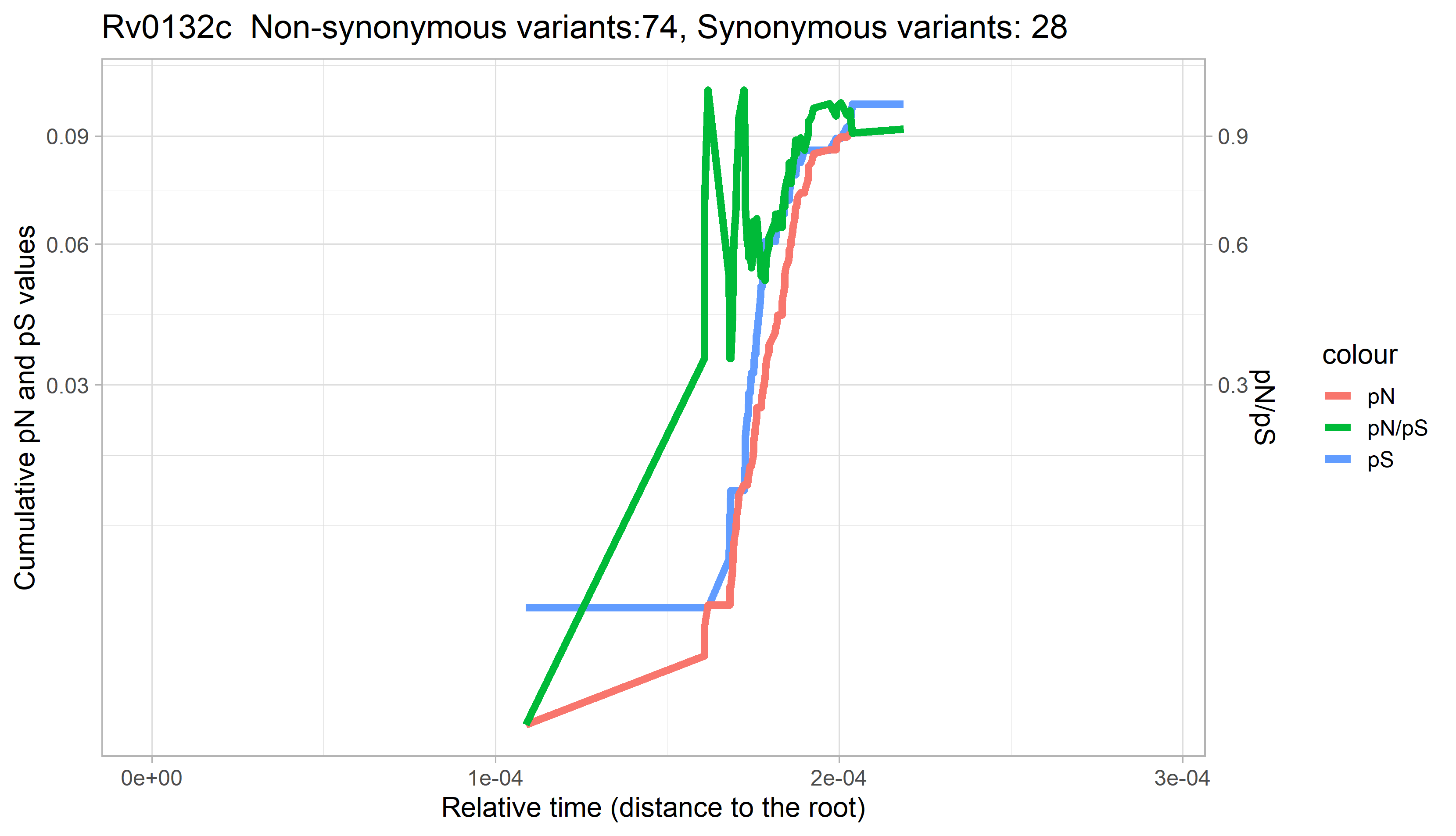

### Rv0133.png

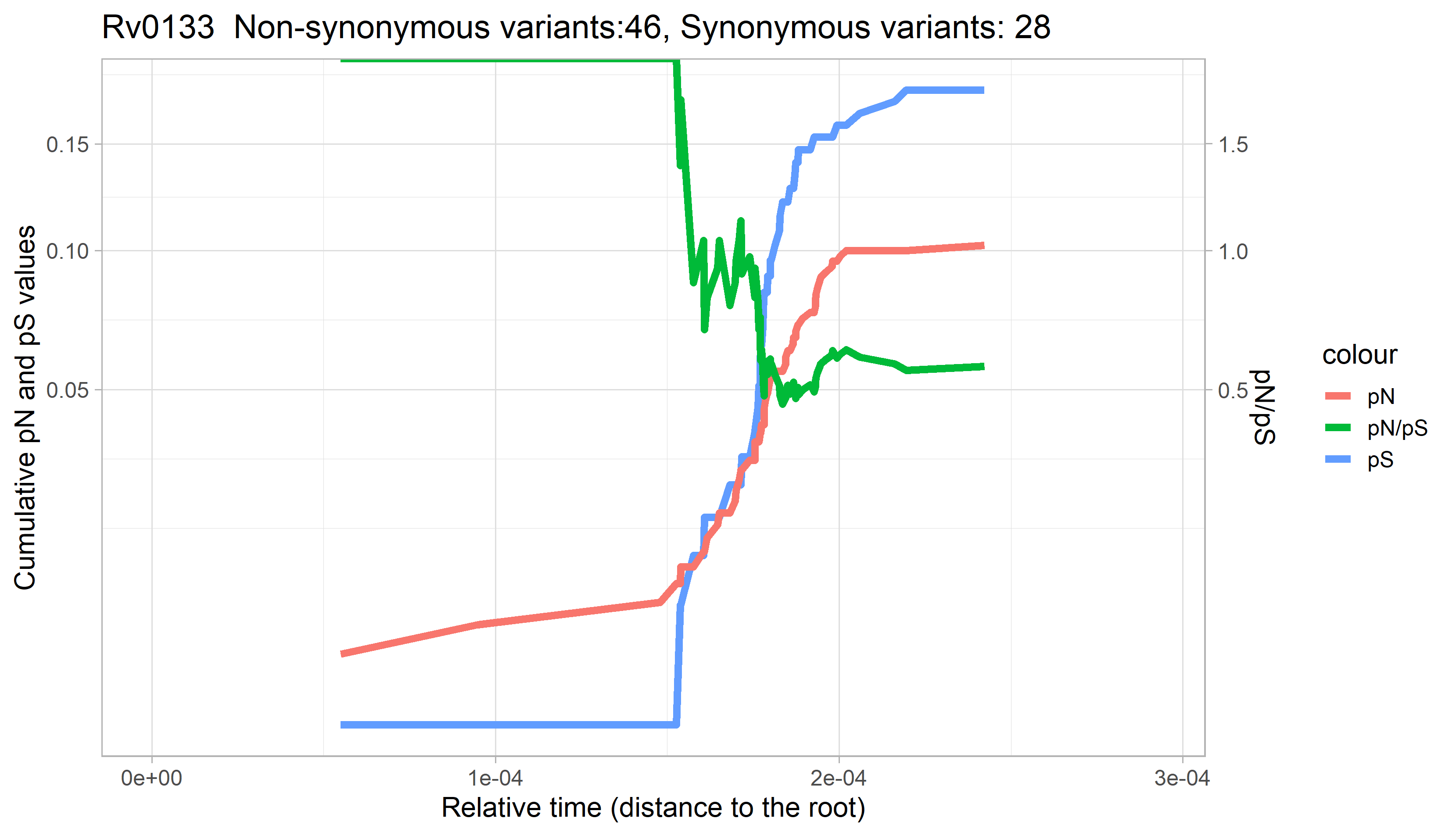

### Rv0134.png

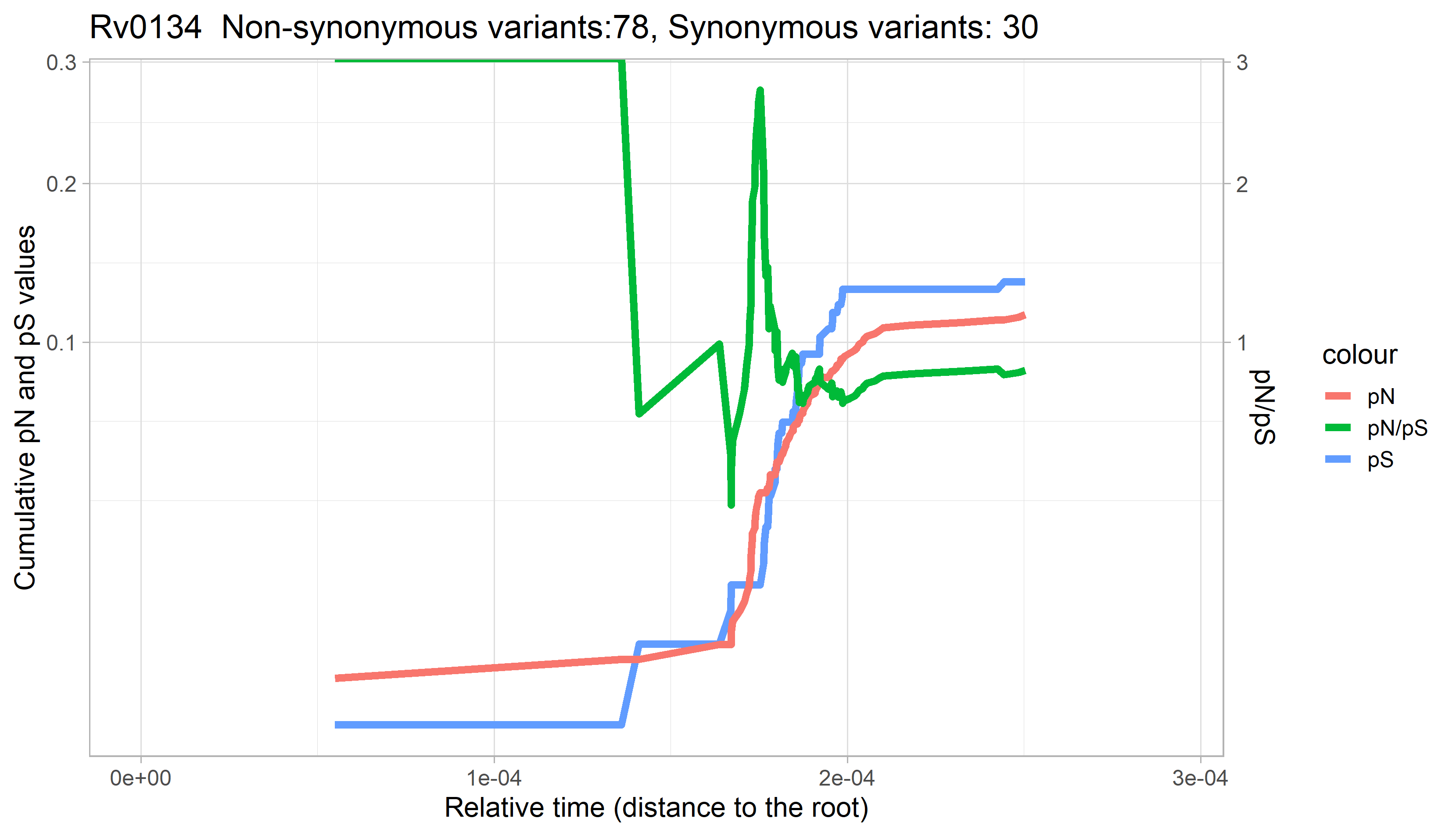

### Rv0135c.png

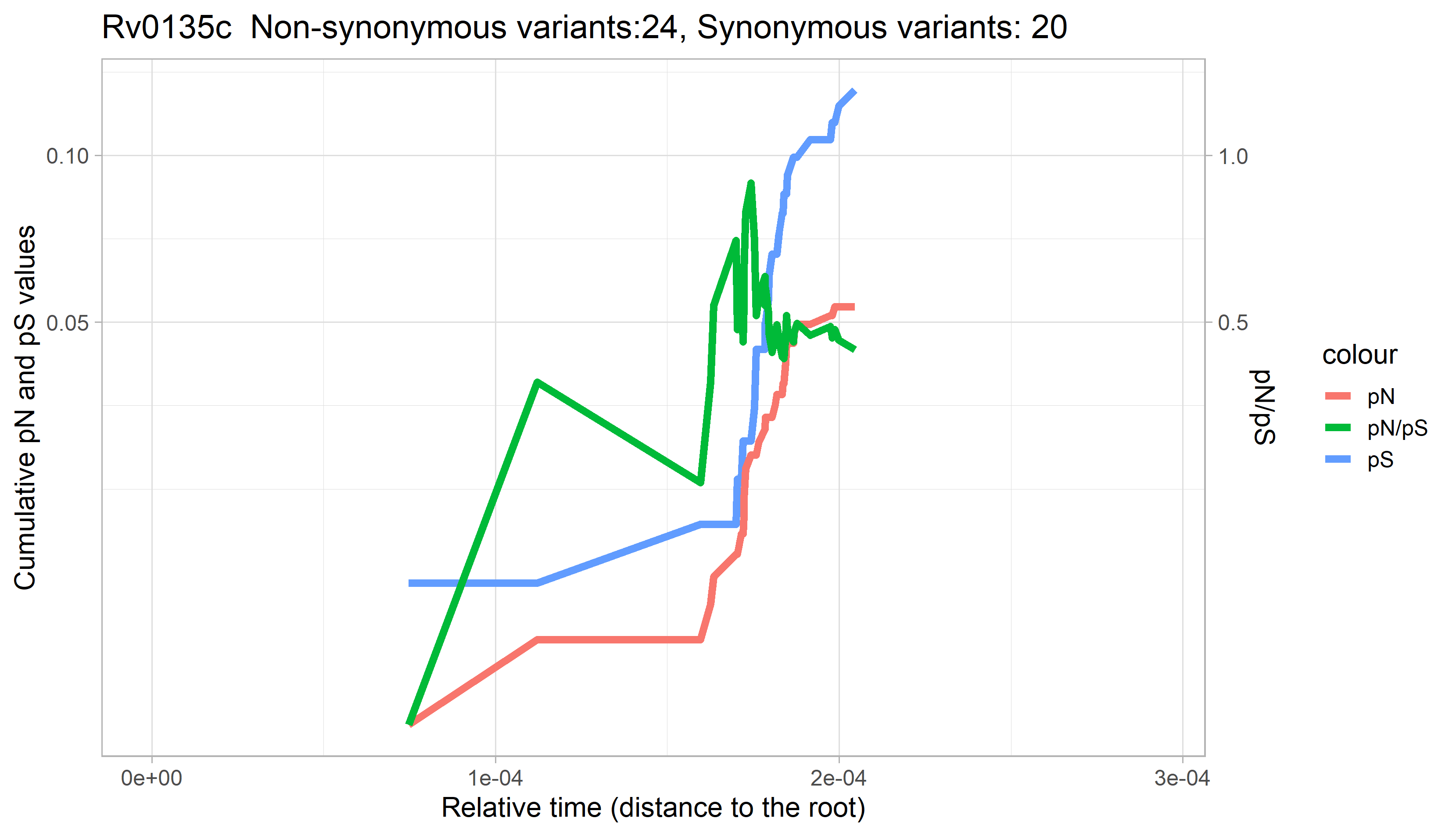

### Rv0136.png

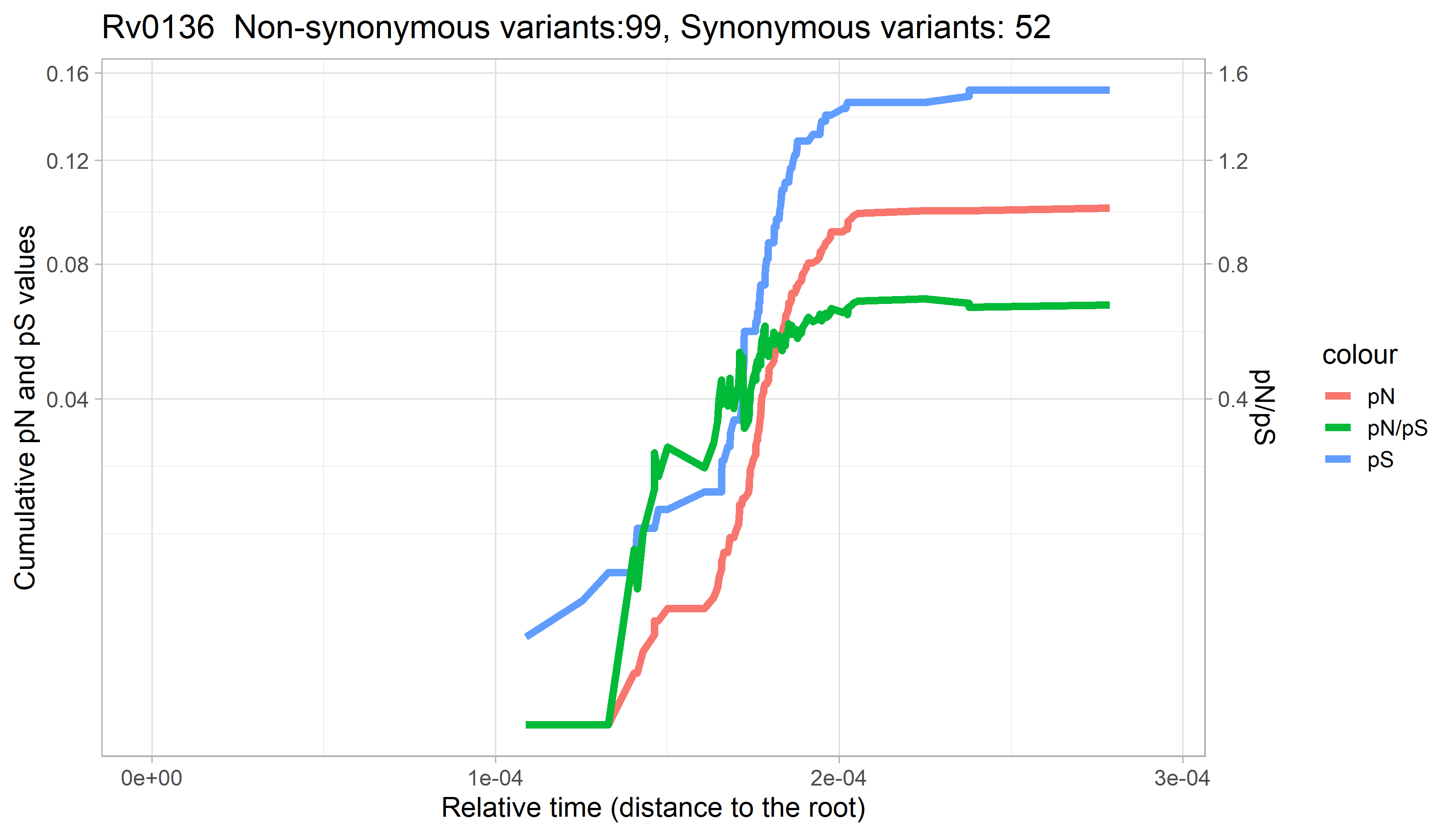

### Rv0137c.png

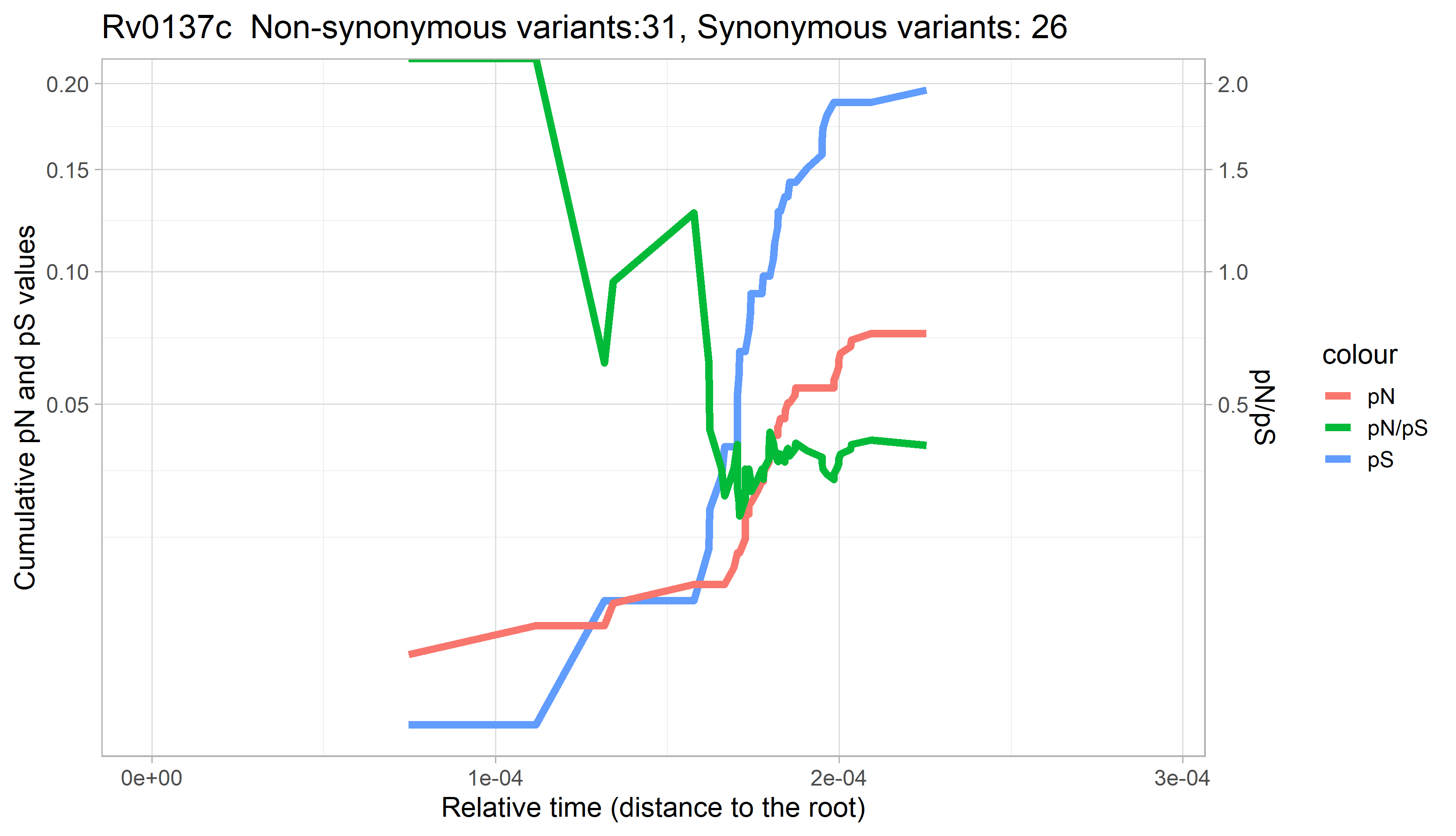

### Rv0138.png

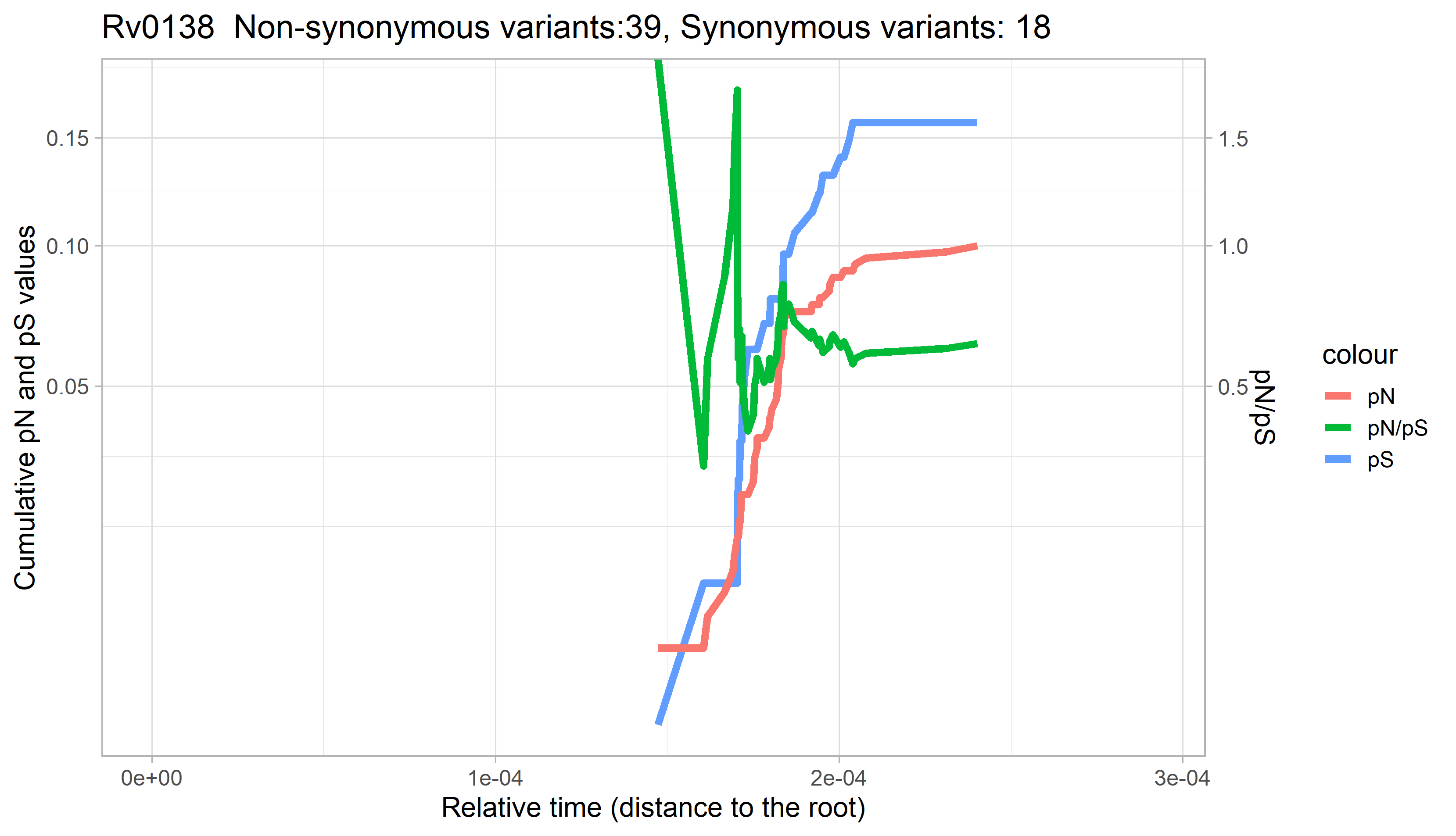

### Rv0139.png

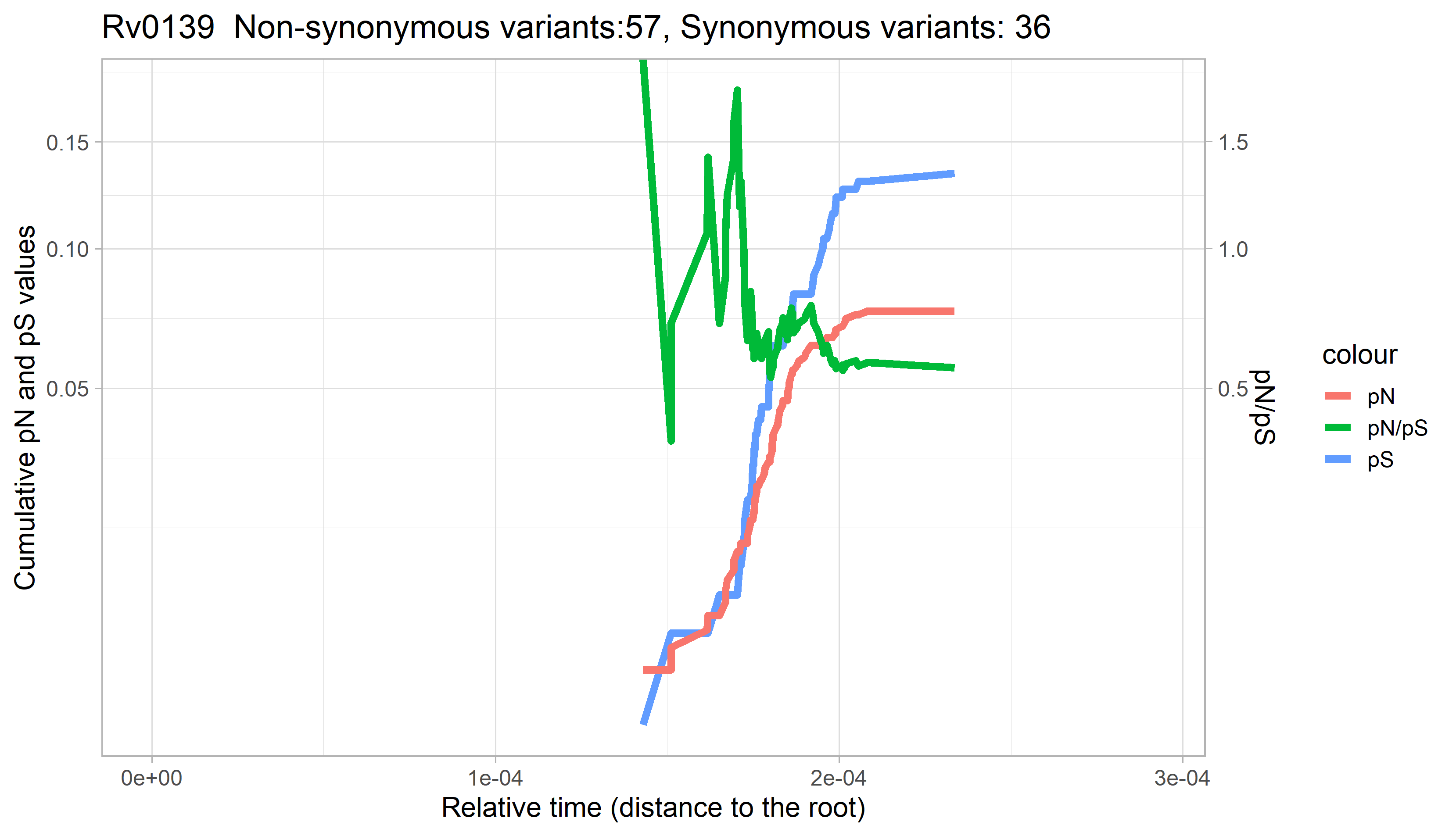

### Rv0140.png

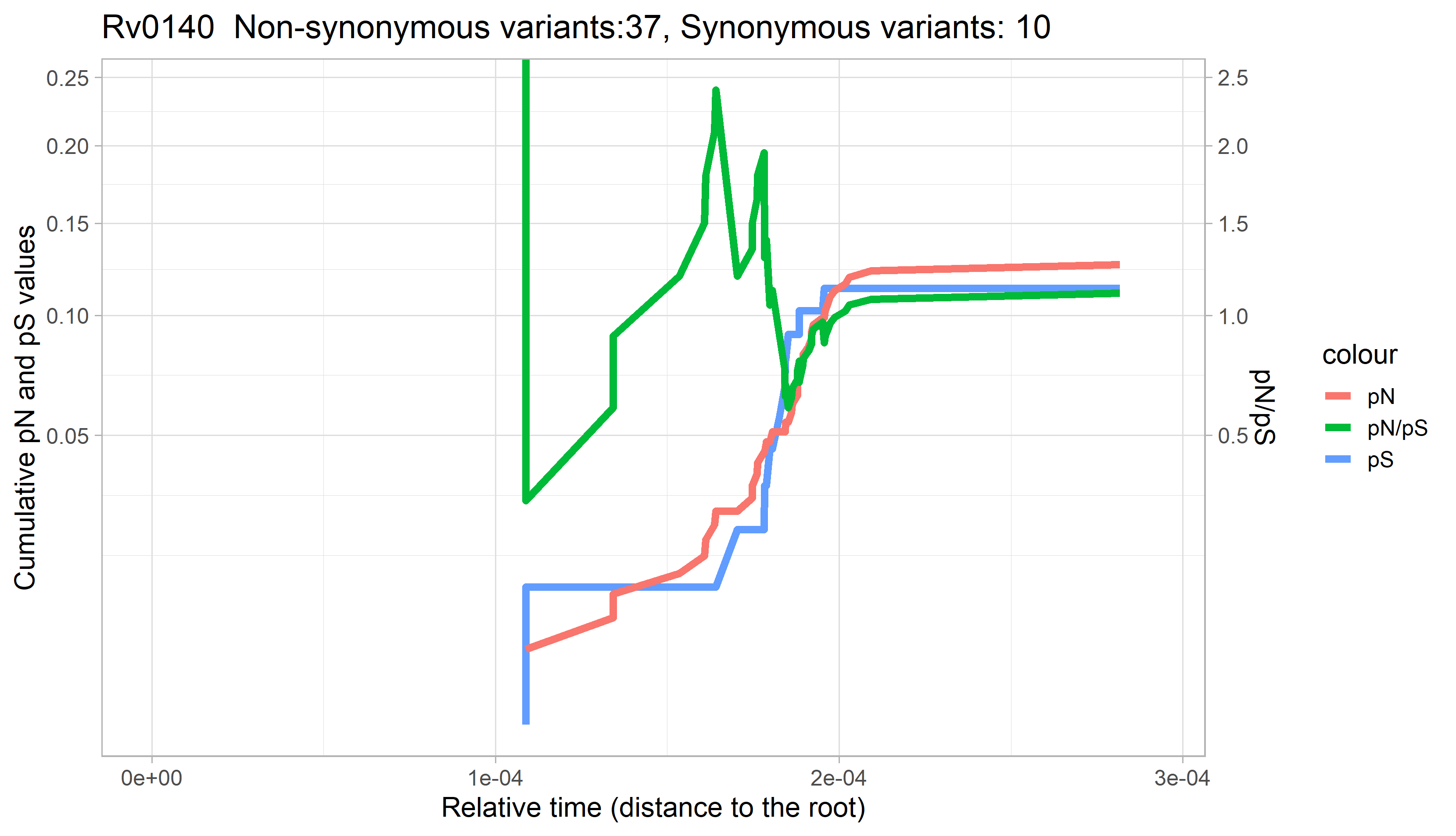

### Rv0141c.png

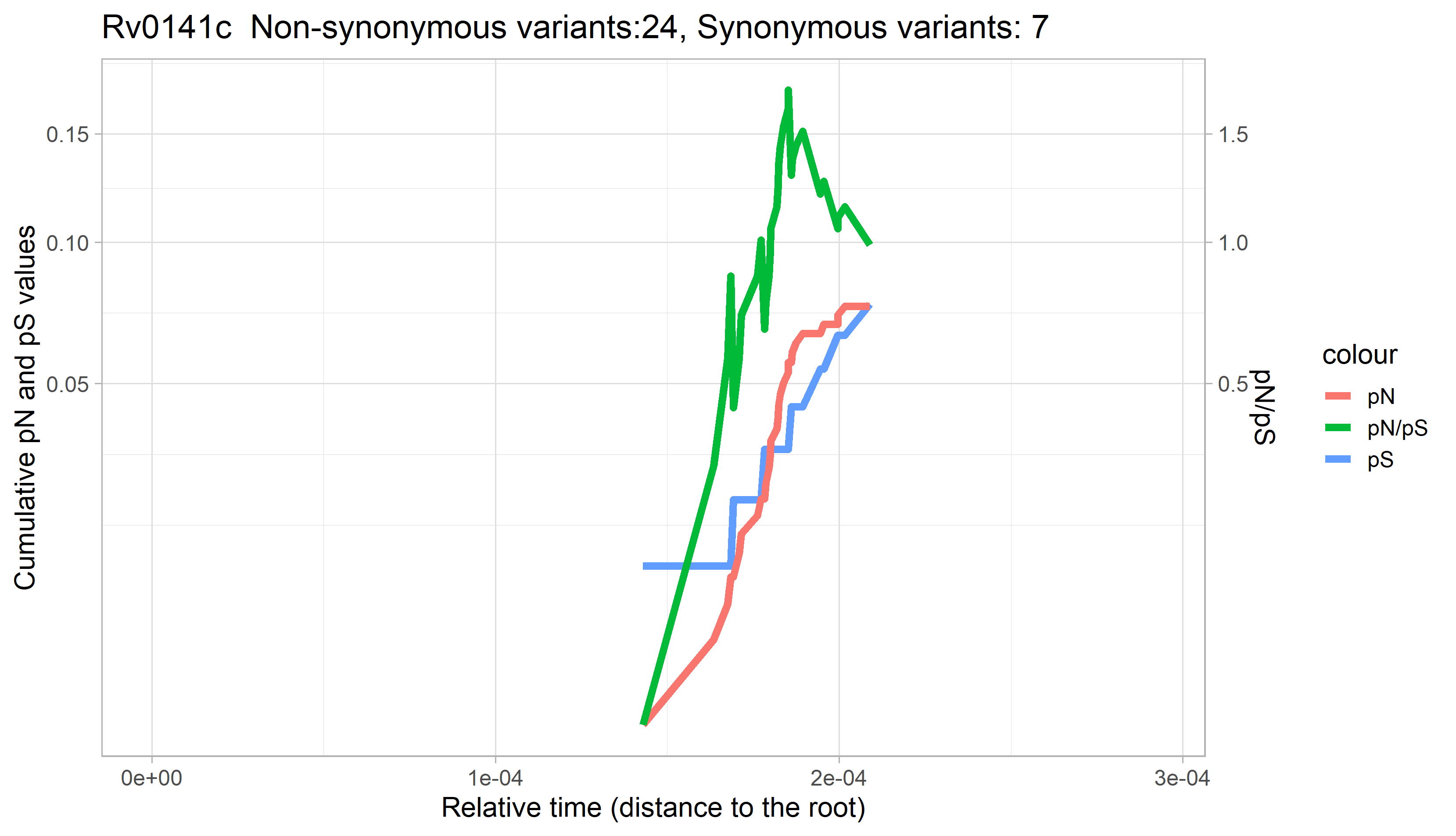

### Rv0142.png

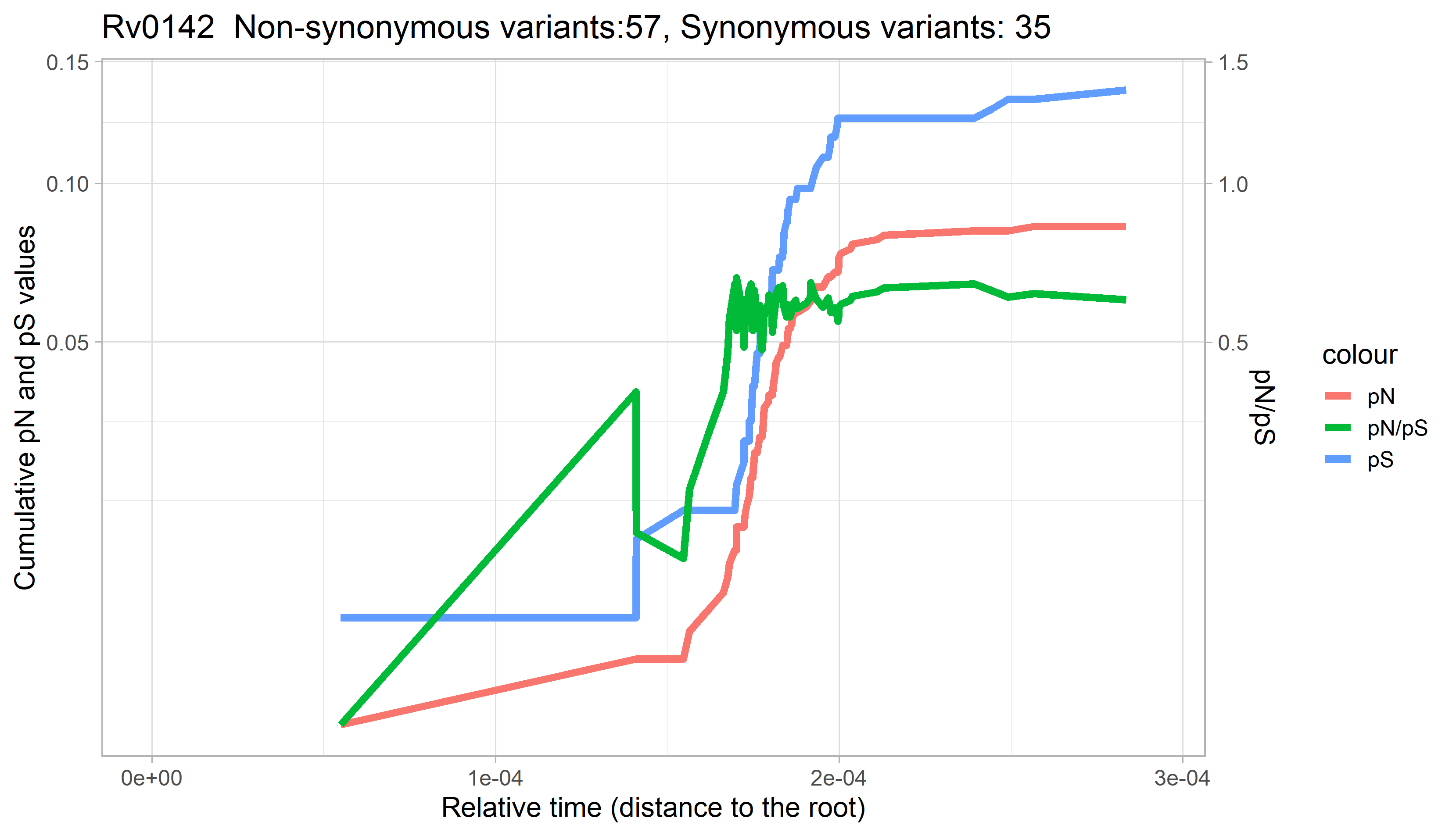

### Rv0143c.png

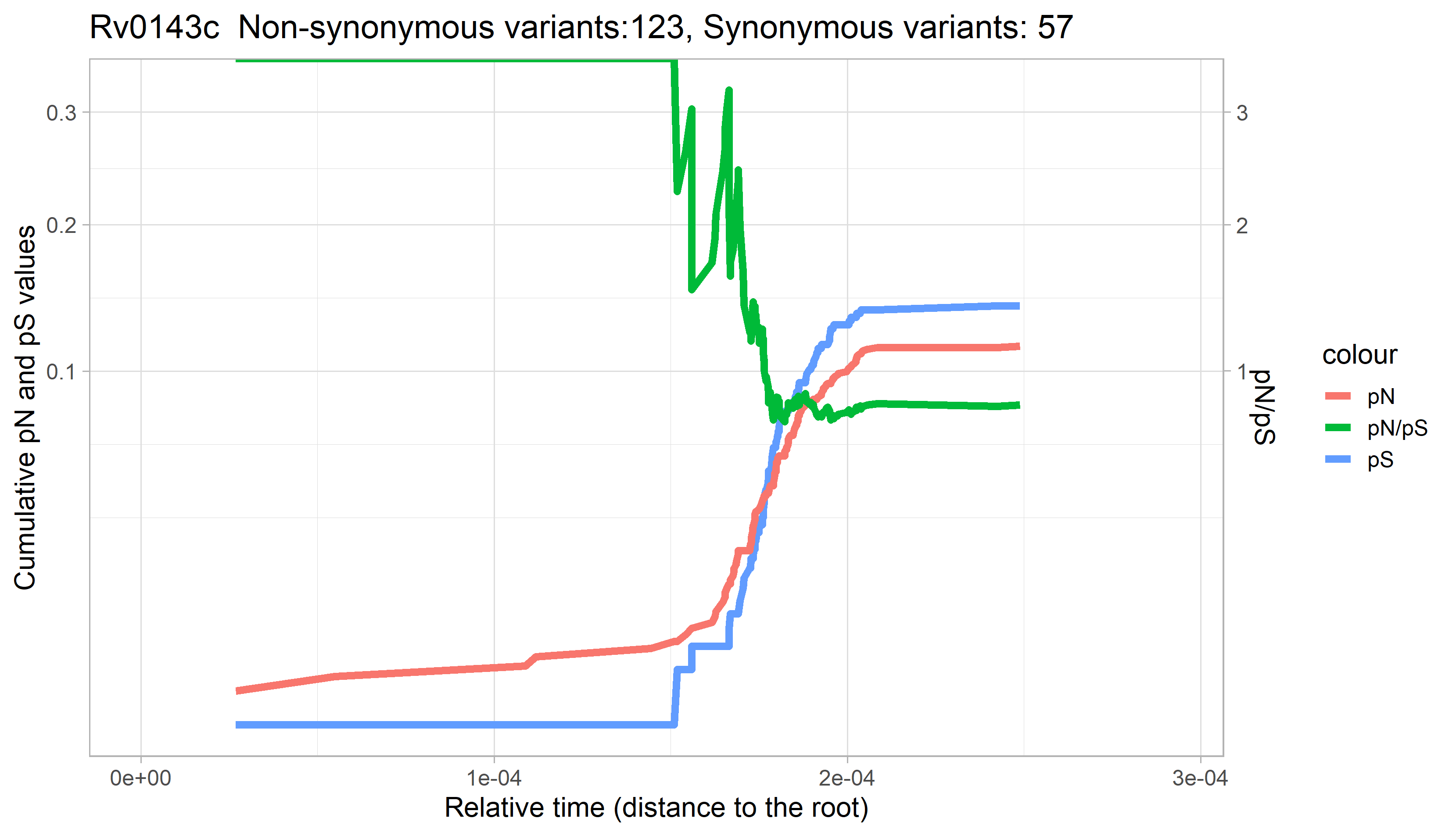

### Rv0144.png

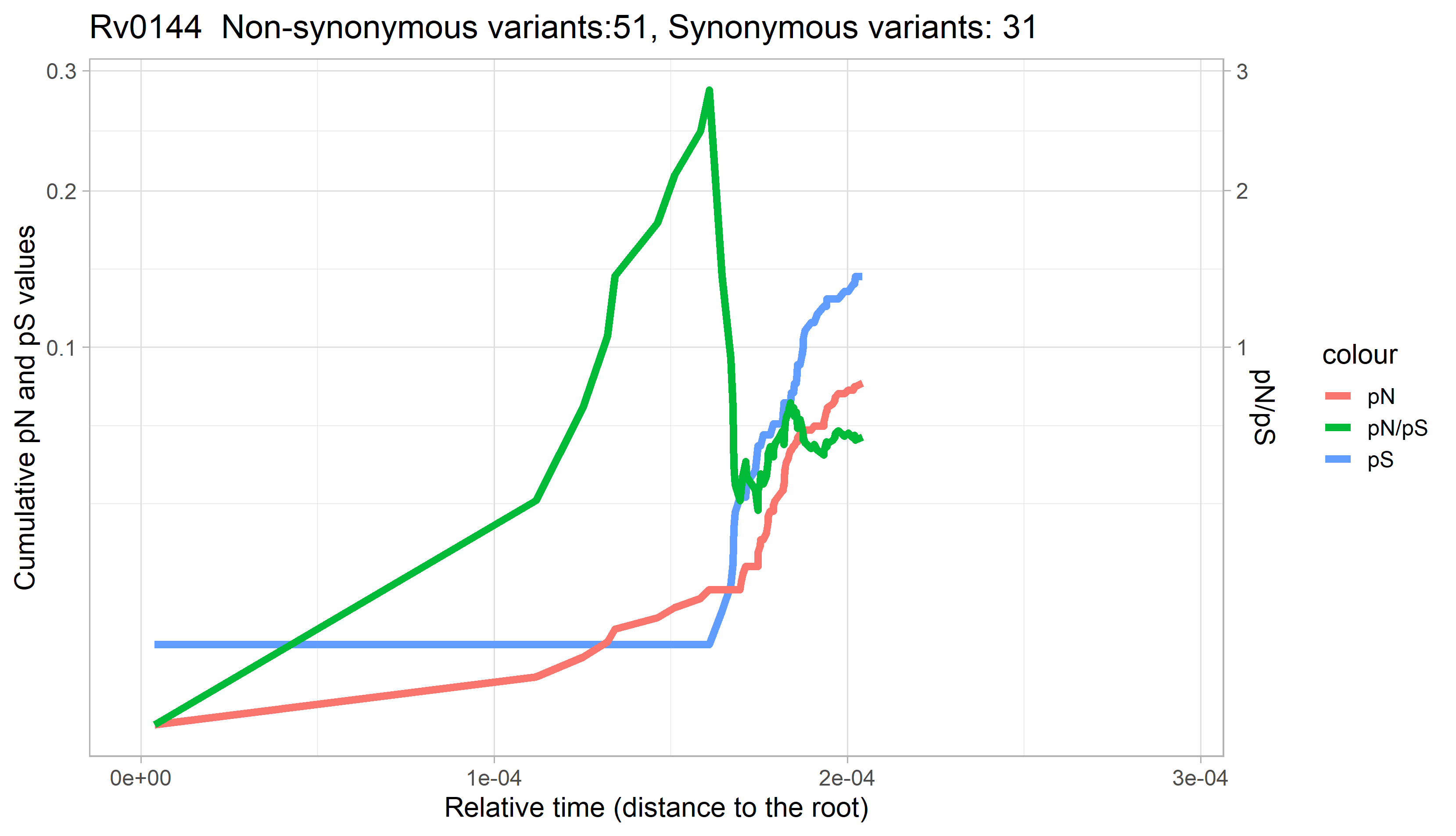

### Rv0145.png

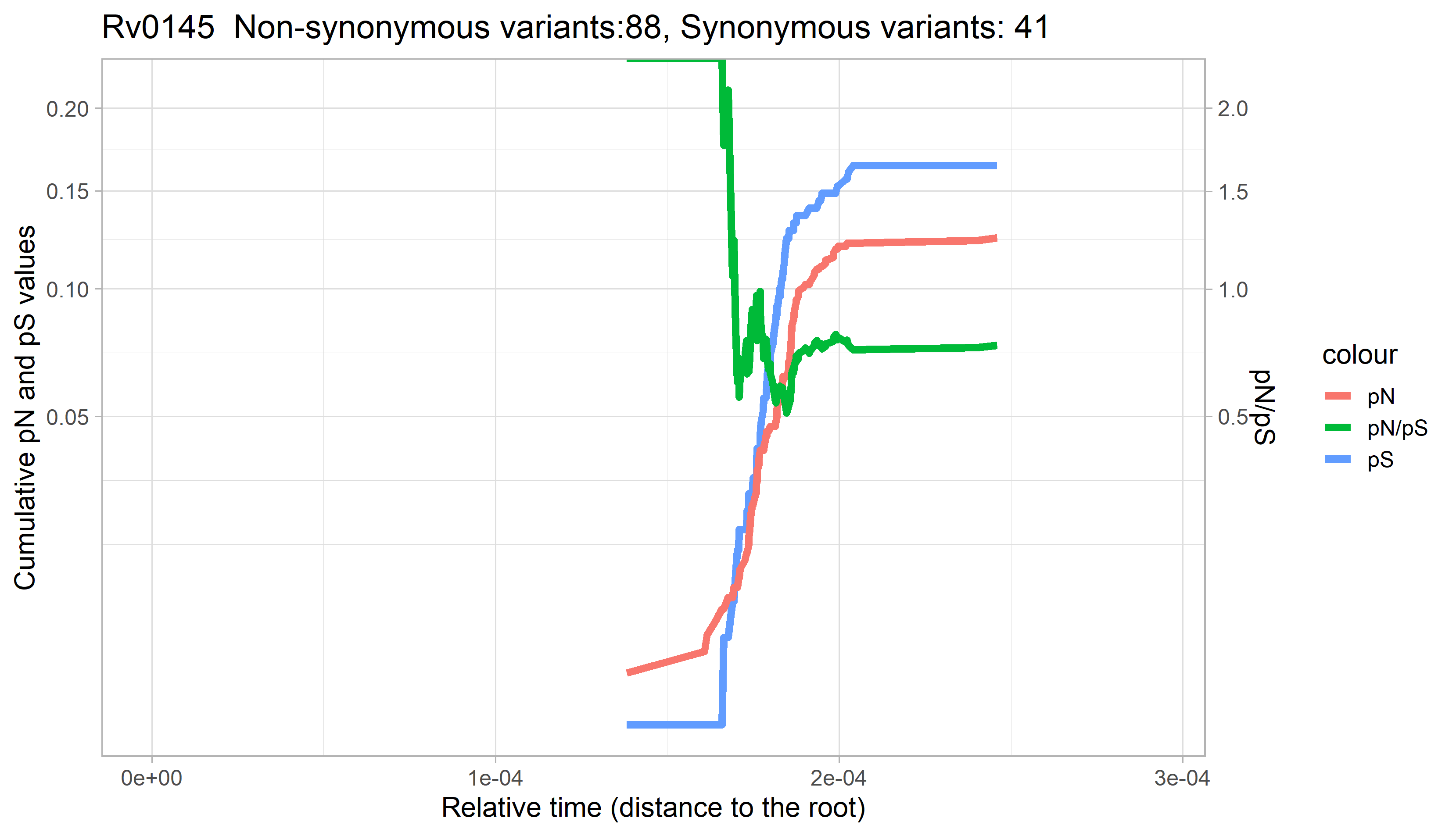

### Rv0146.png

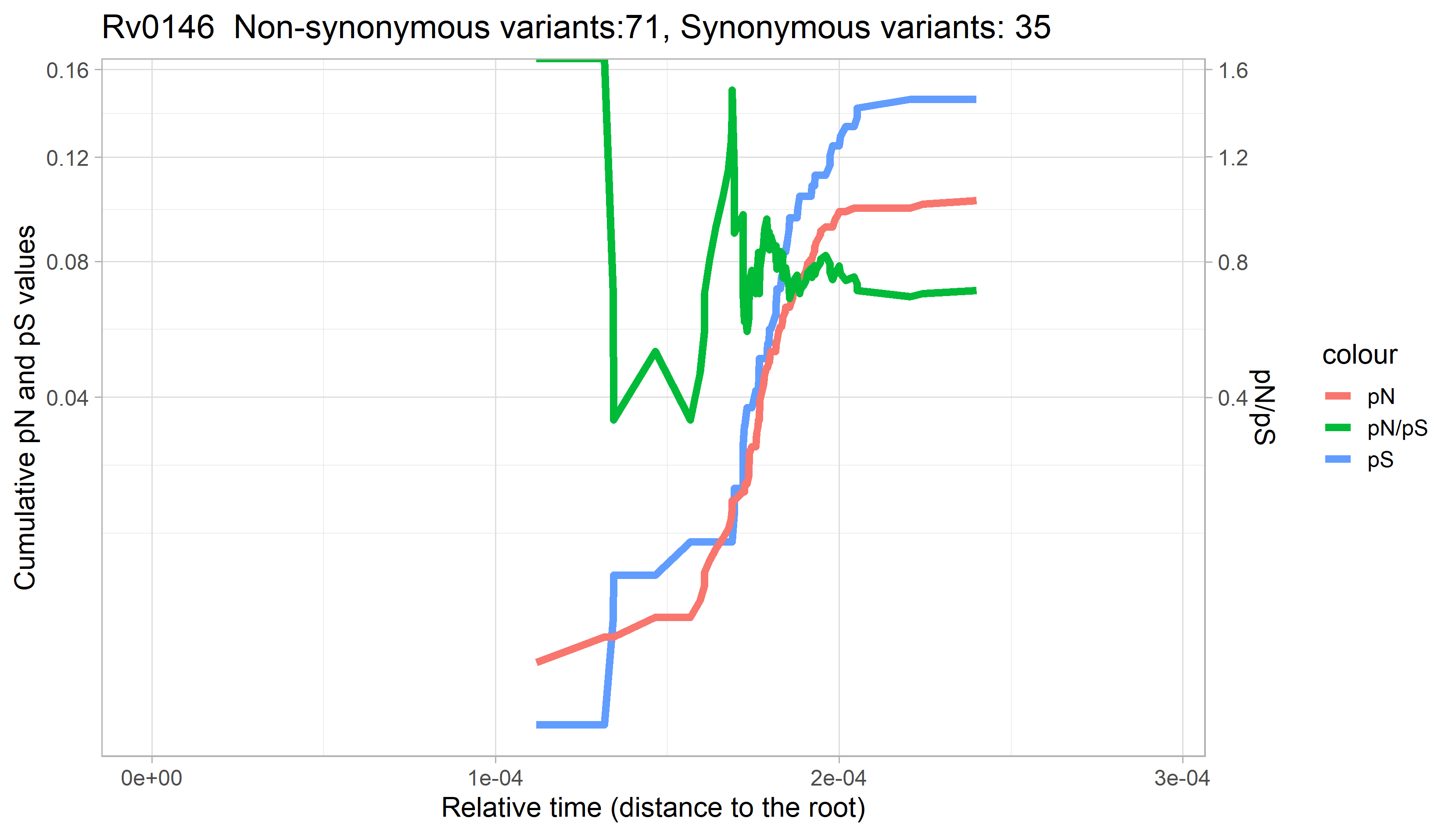

### Rv0147.png

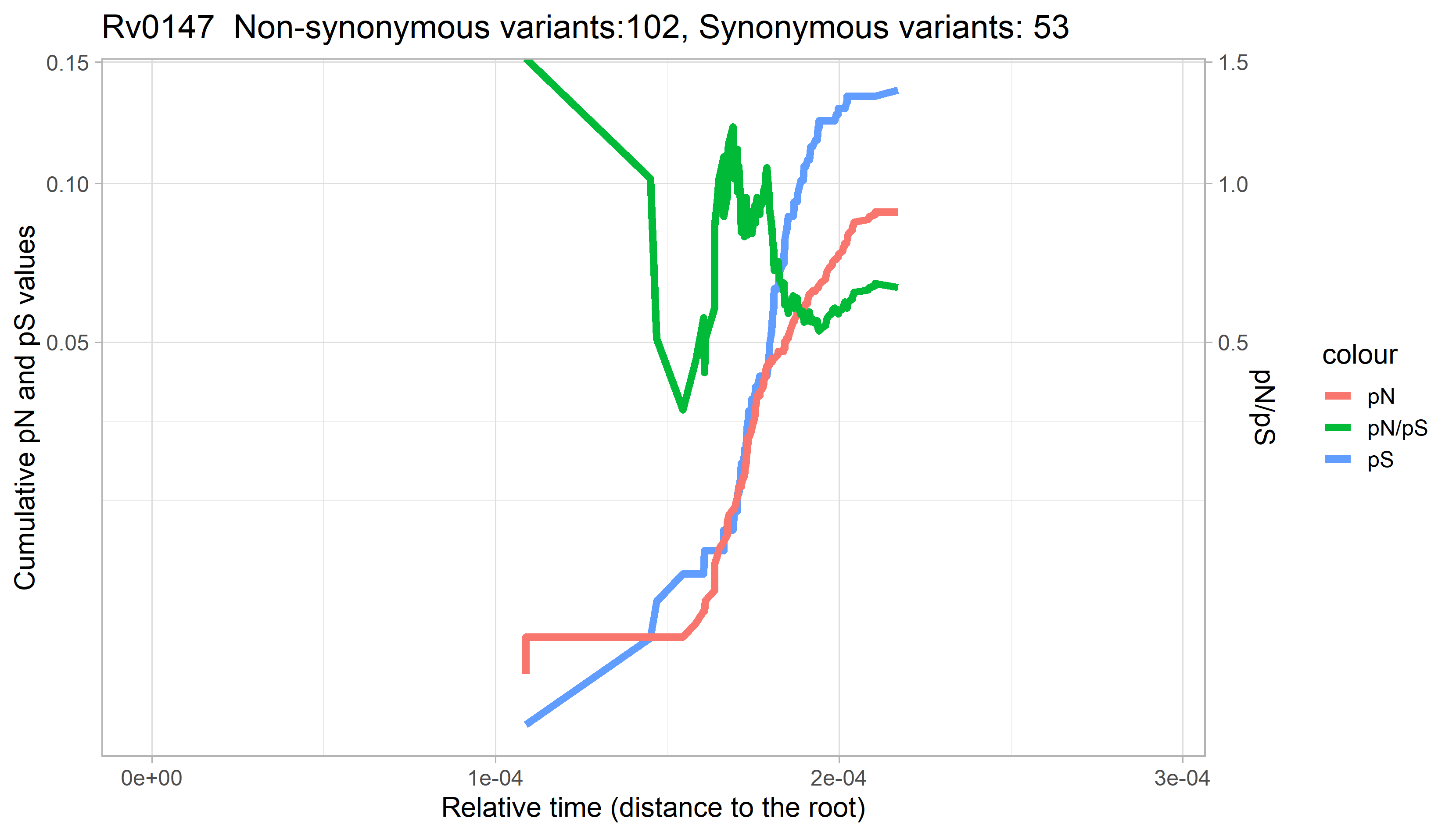

### Rv0148.png

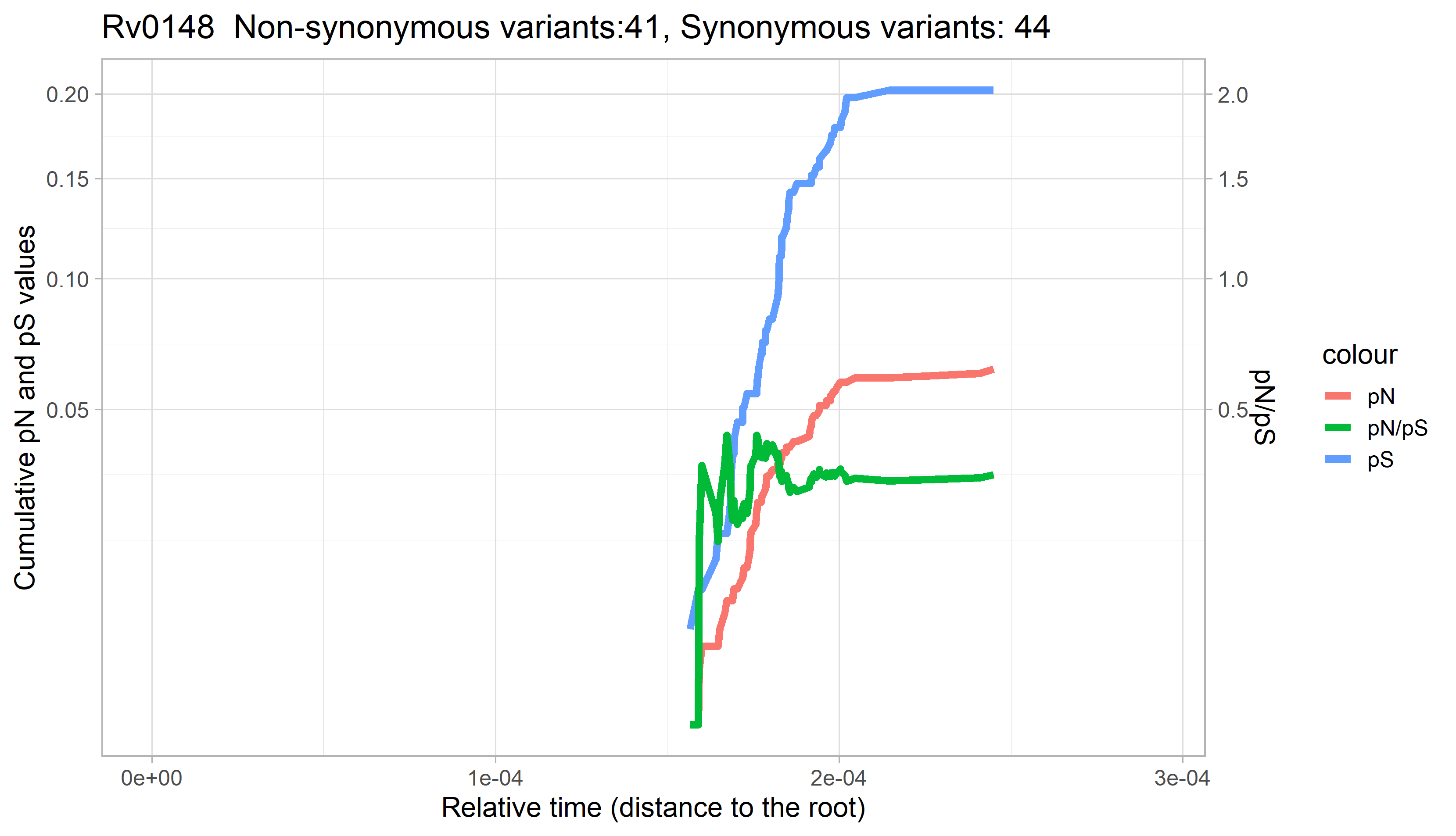

### Rv0149.png

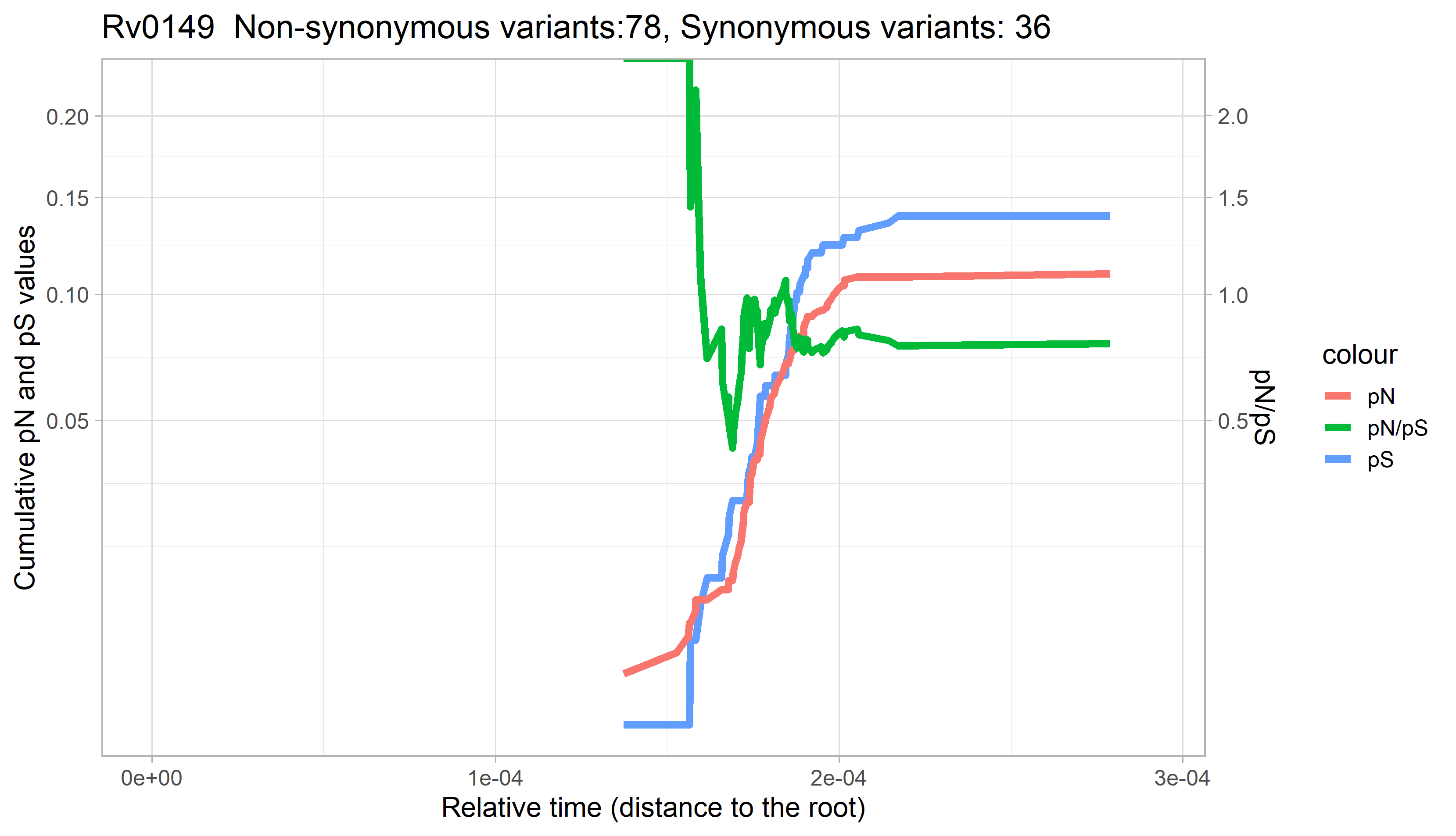

### Rv0150c.png

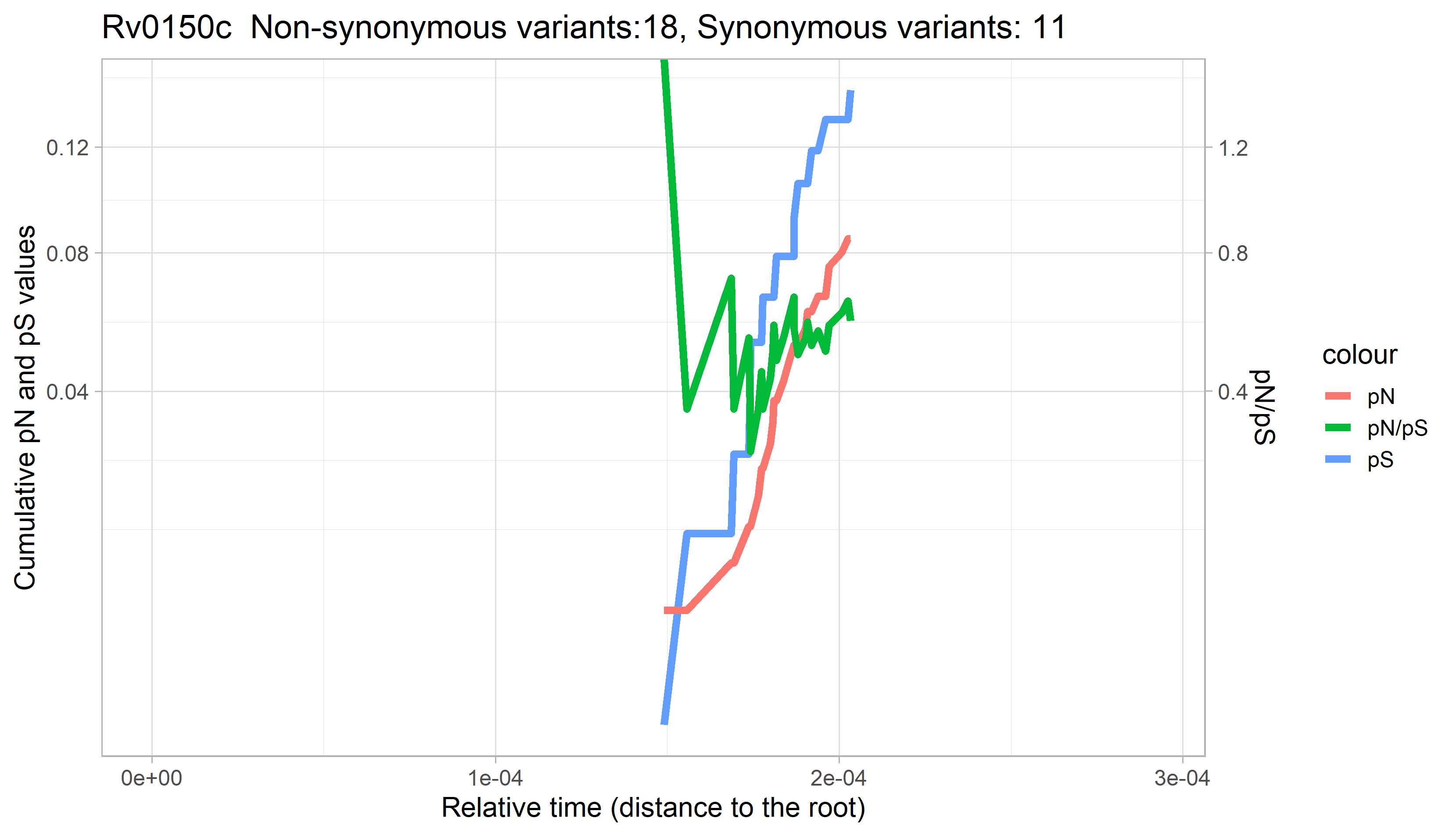

### Rv0153c.png

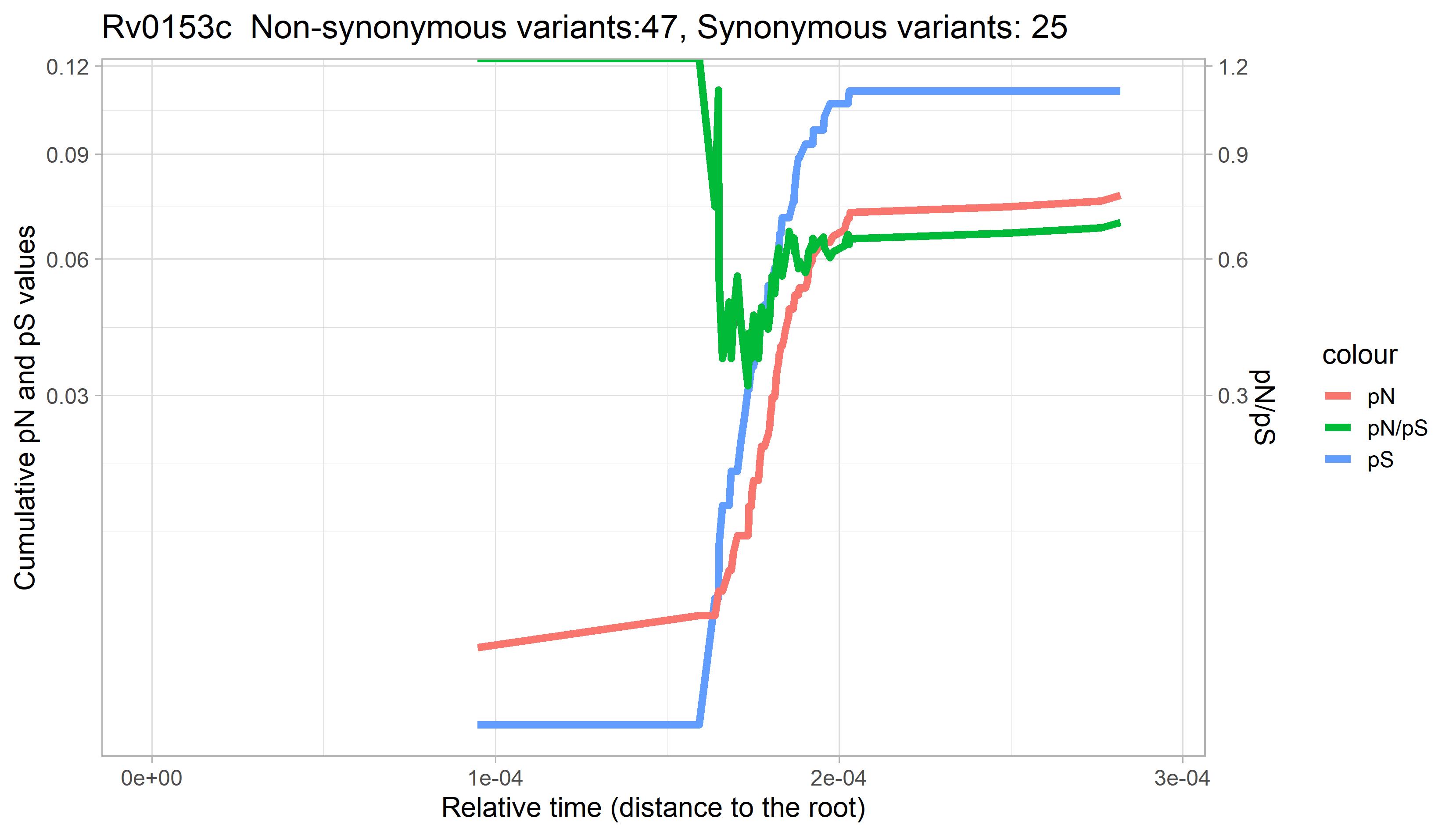

### Rv0154c.png

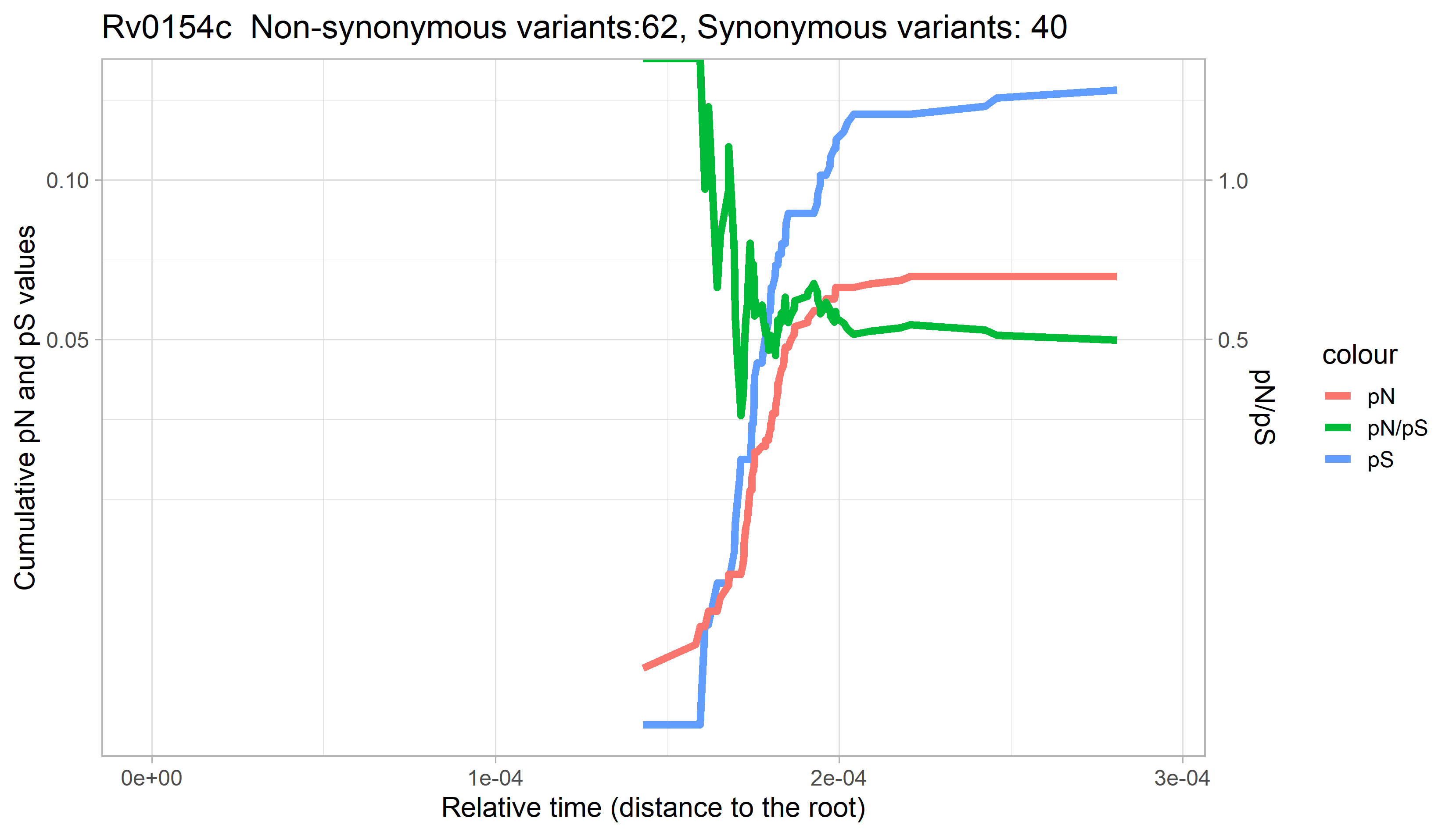

### Rv0155.png

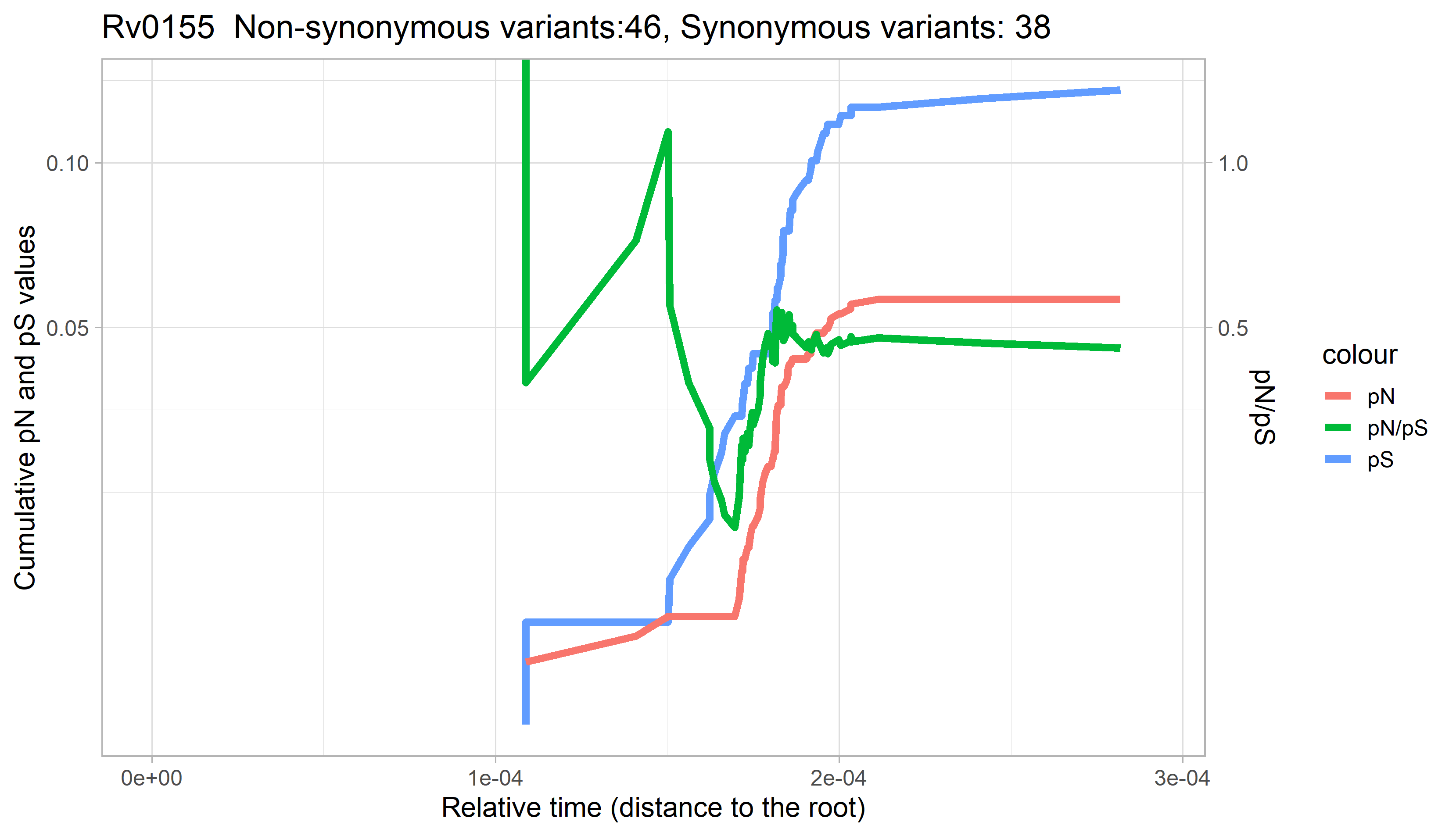

### Rv0156.png

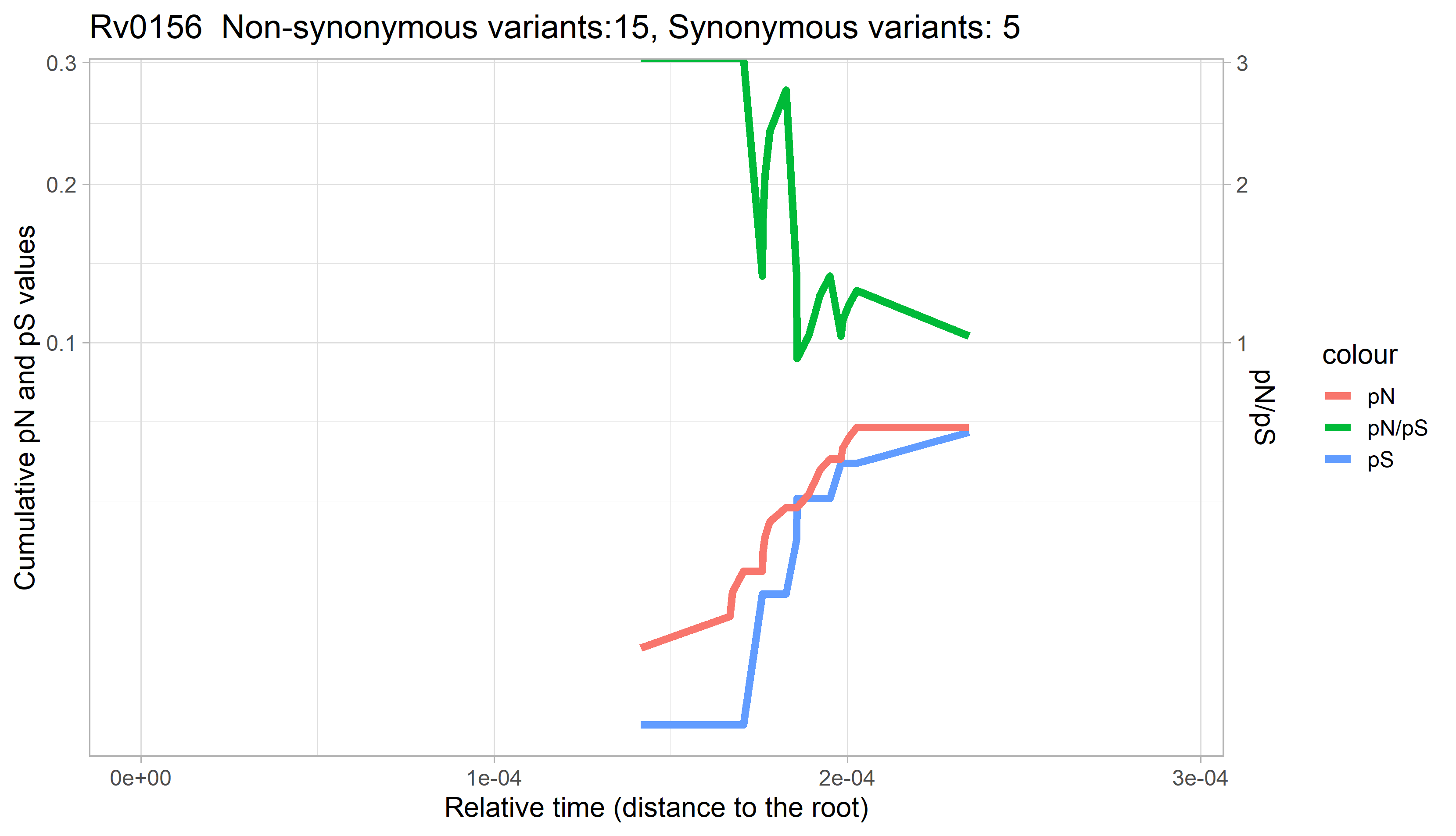

### Rv0157.png

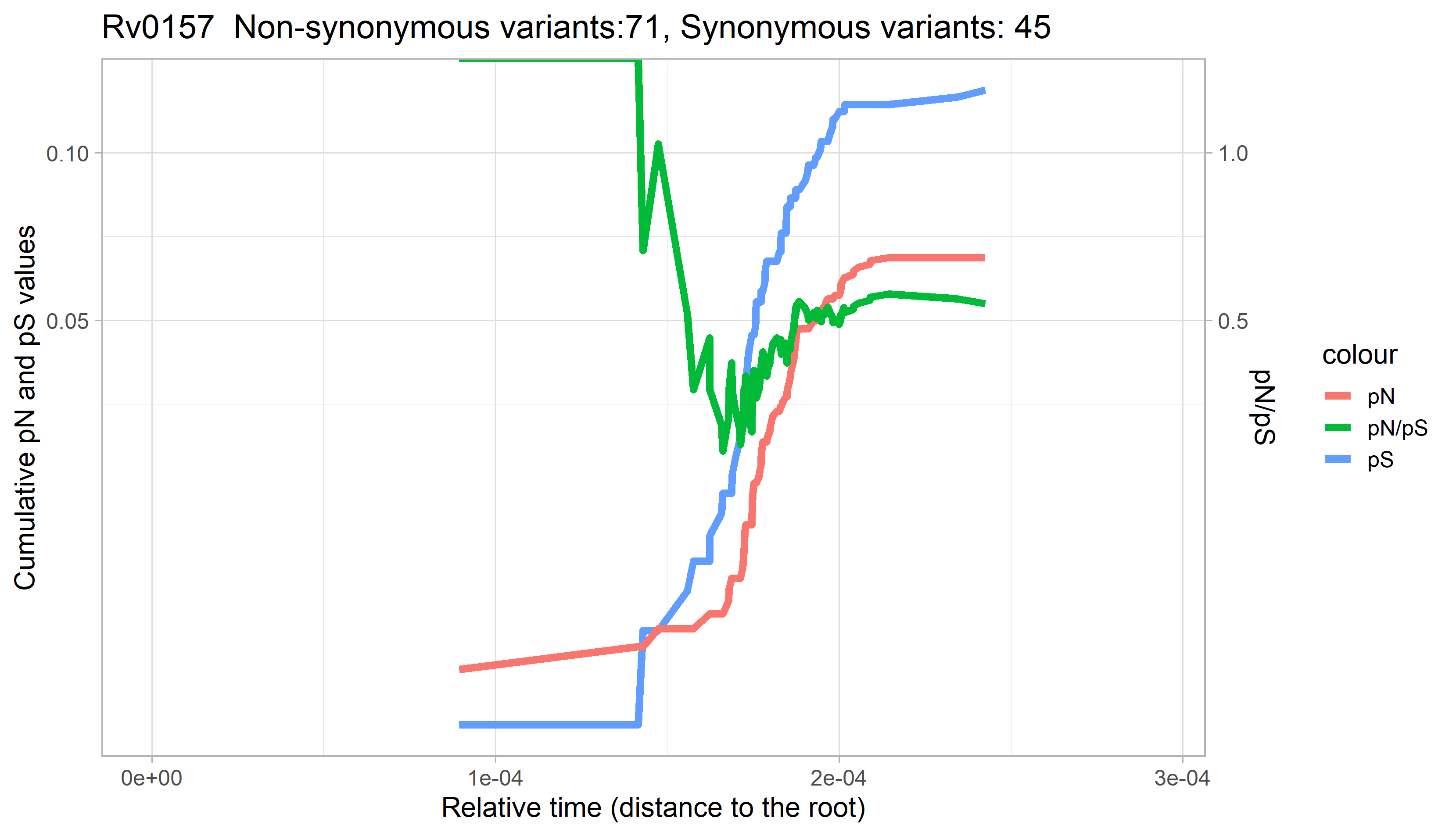

### Rv0158.png

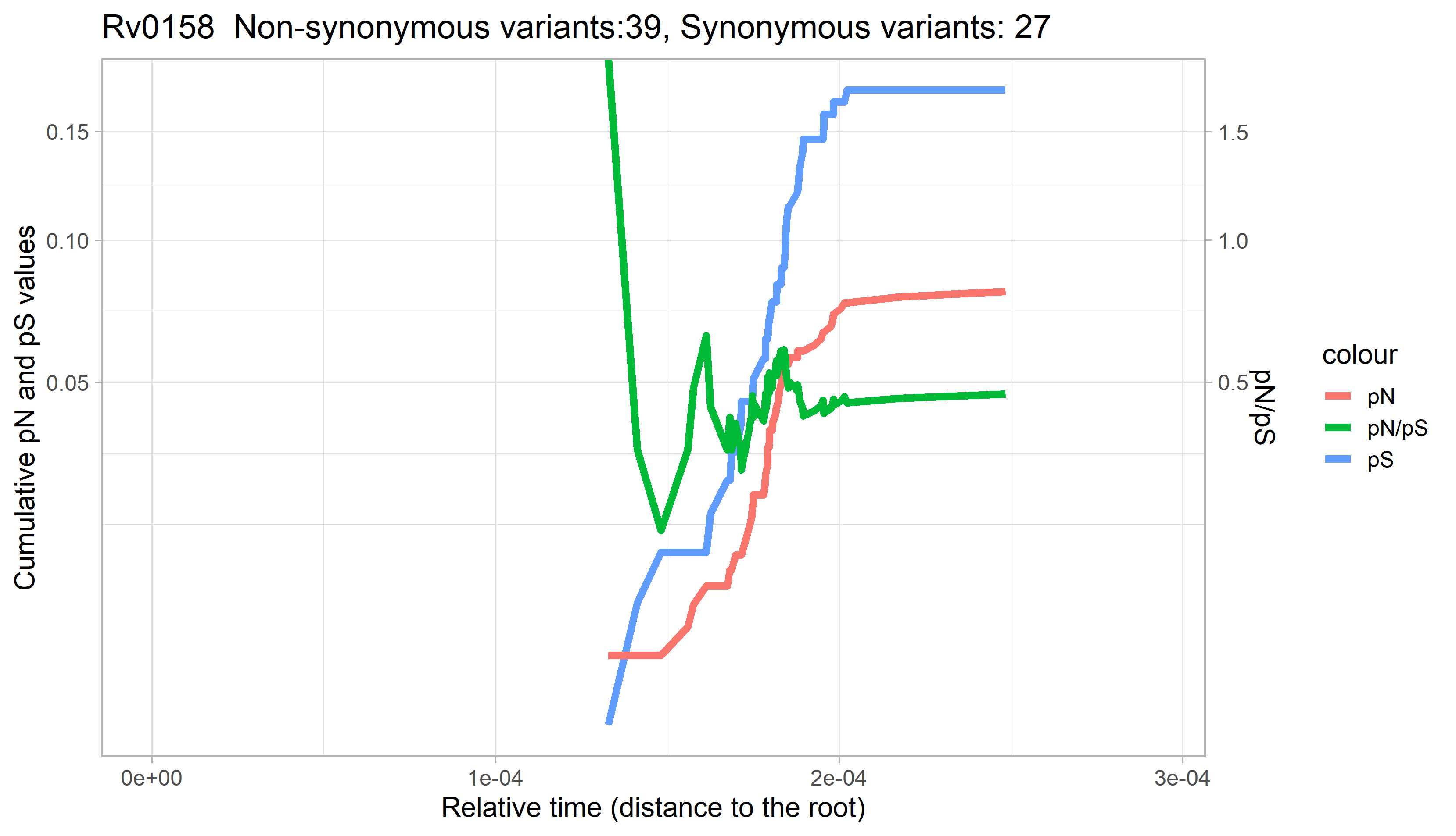

### Rv0161.png

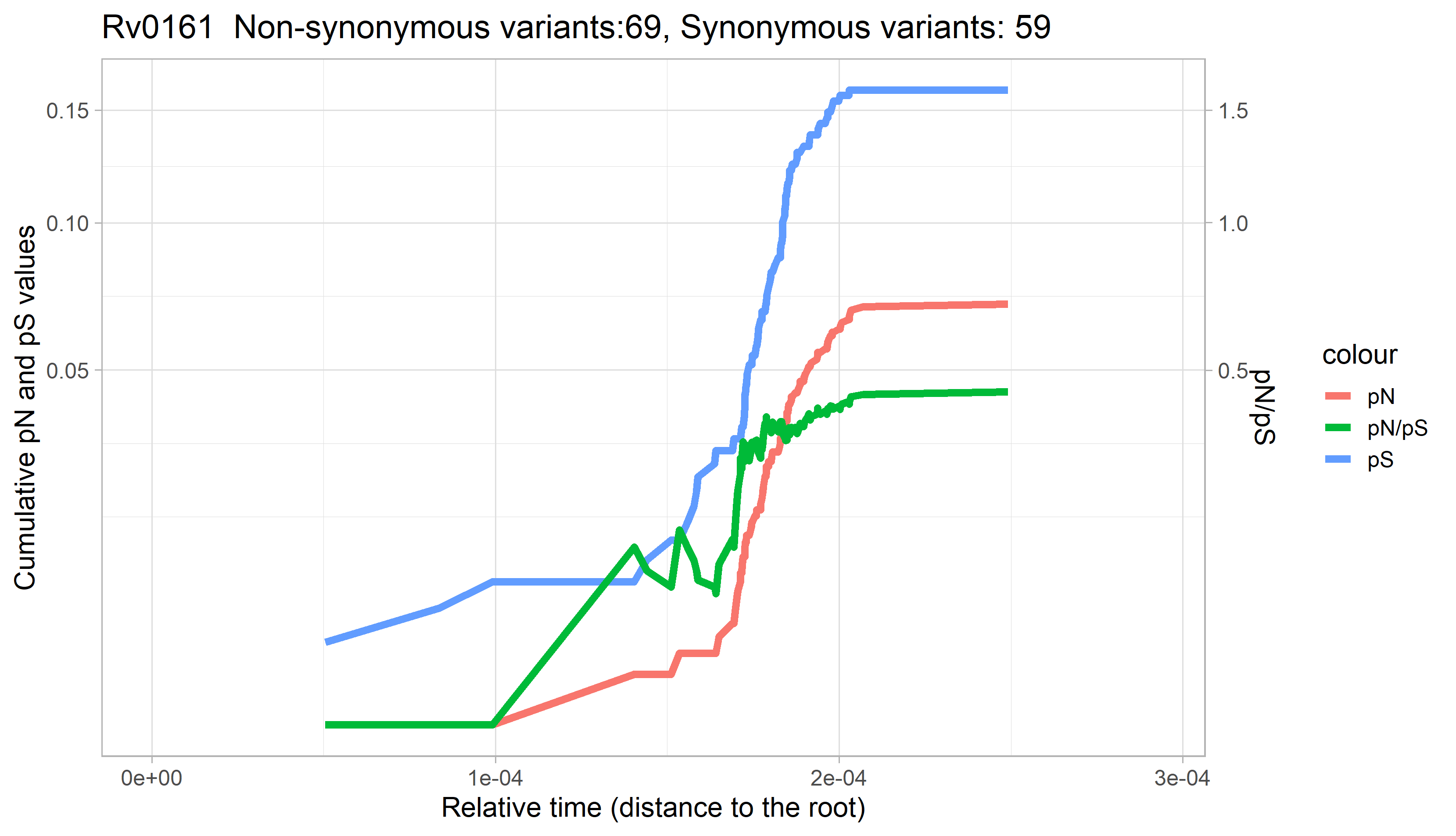

### Rv0162c.png

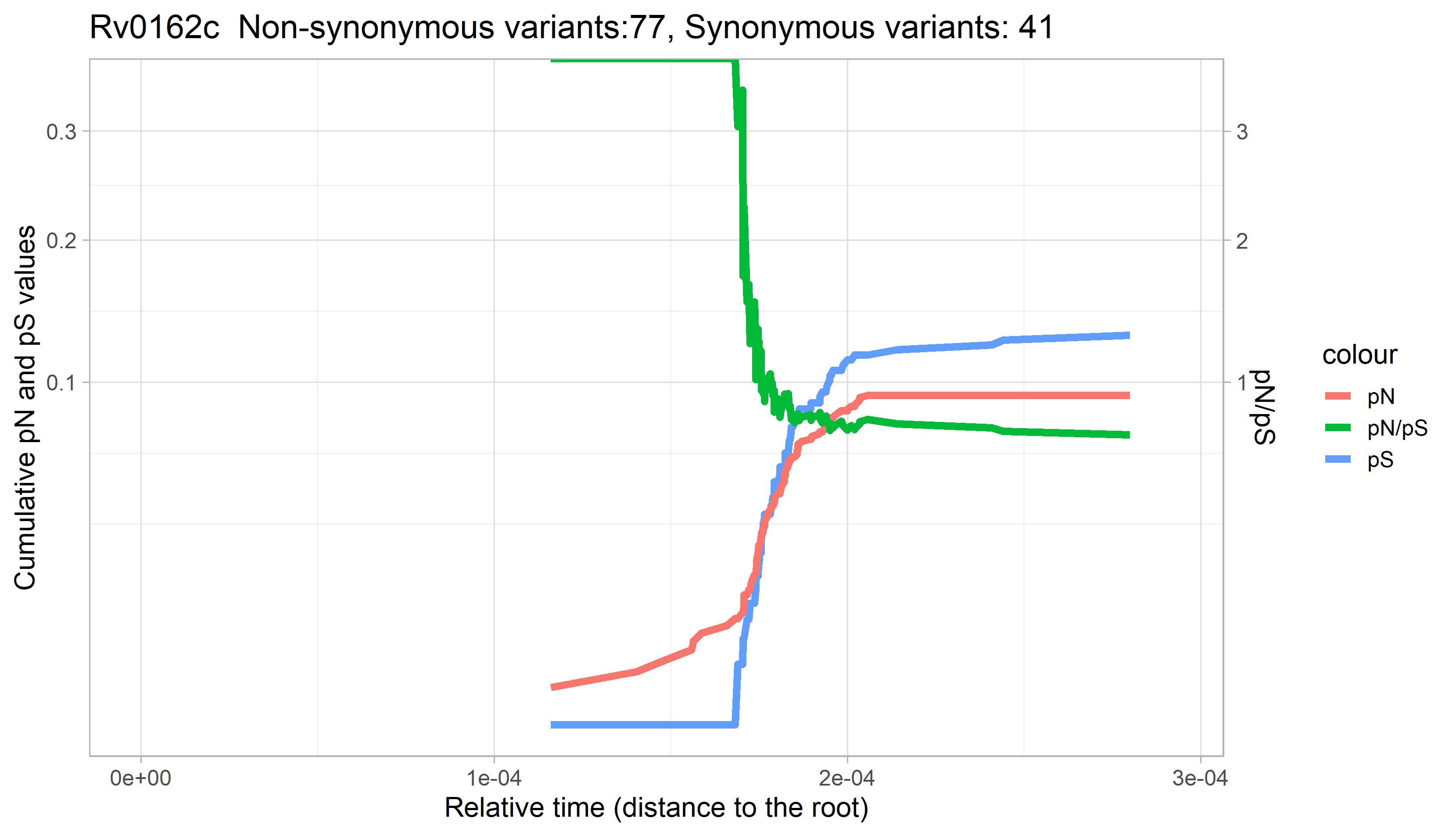

### Rv0163.png

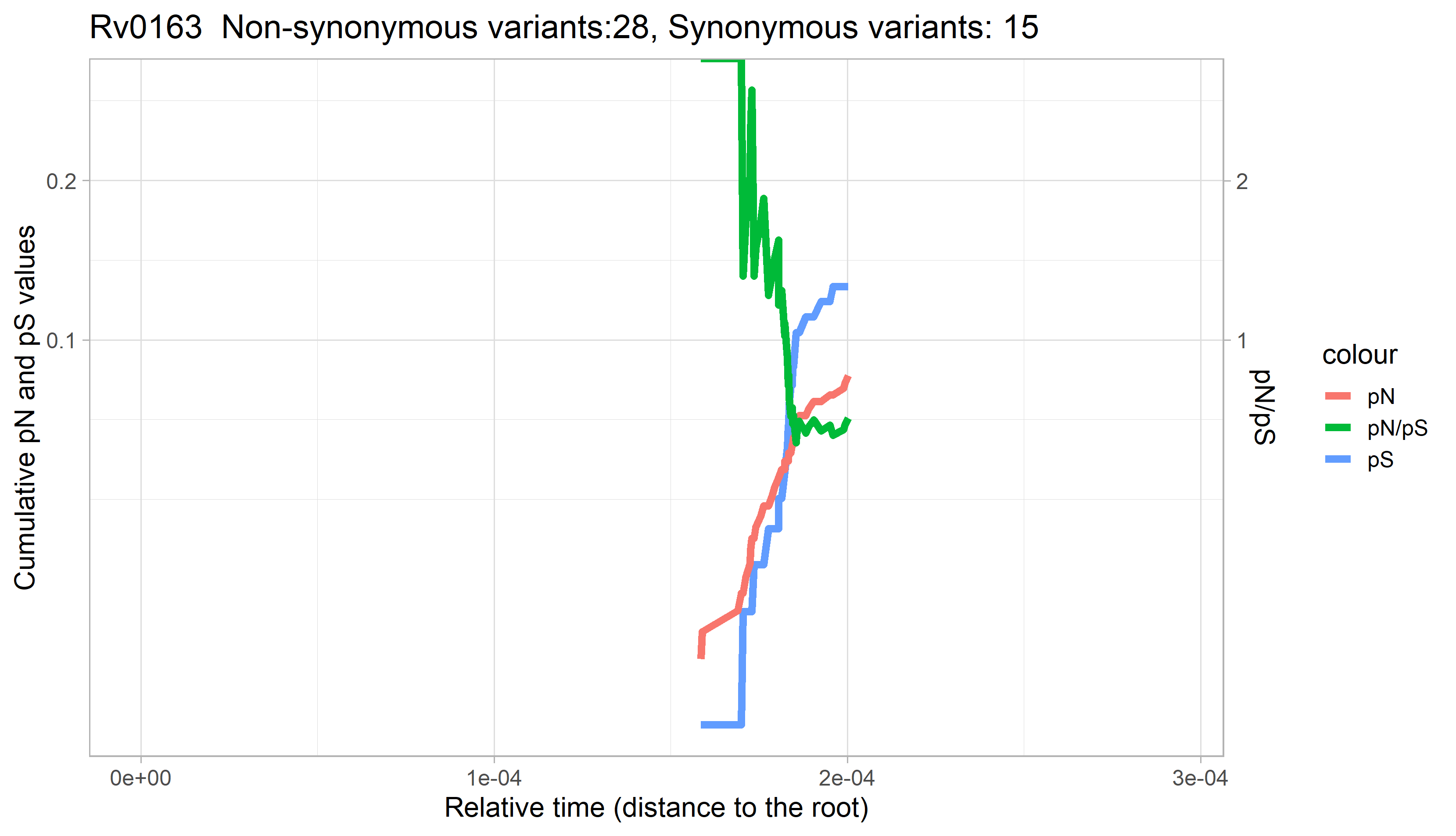

### Rv0164.png

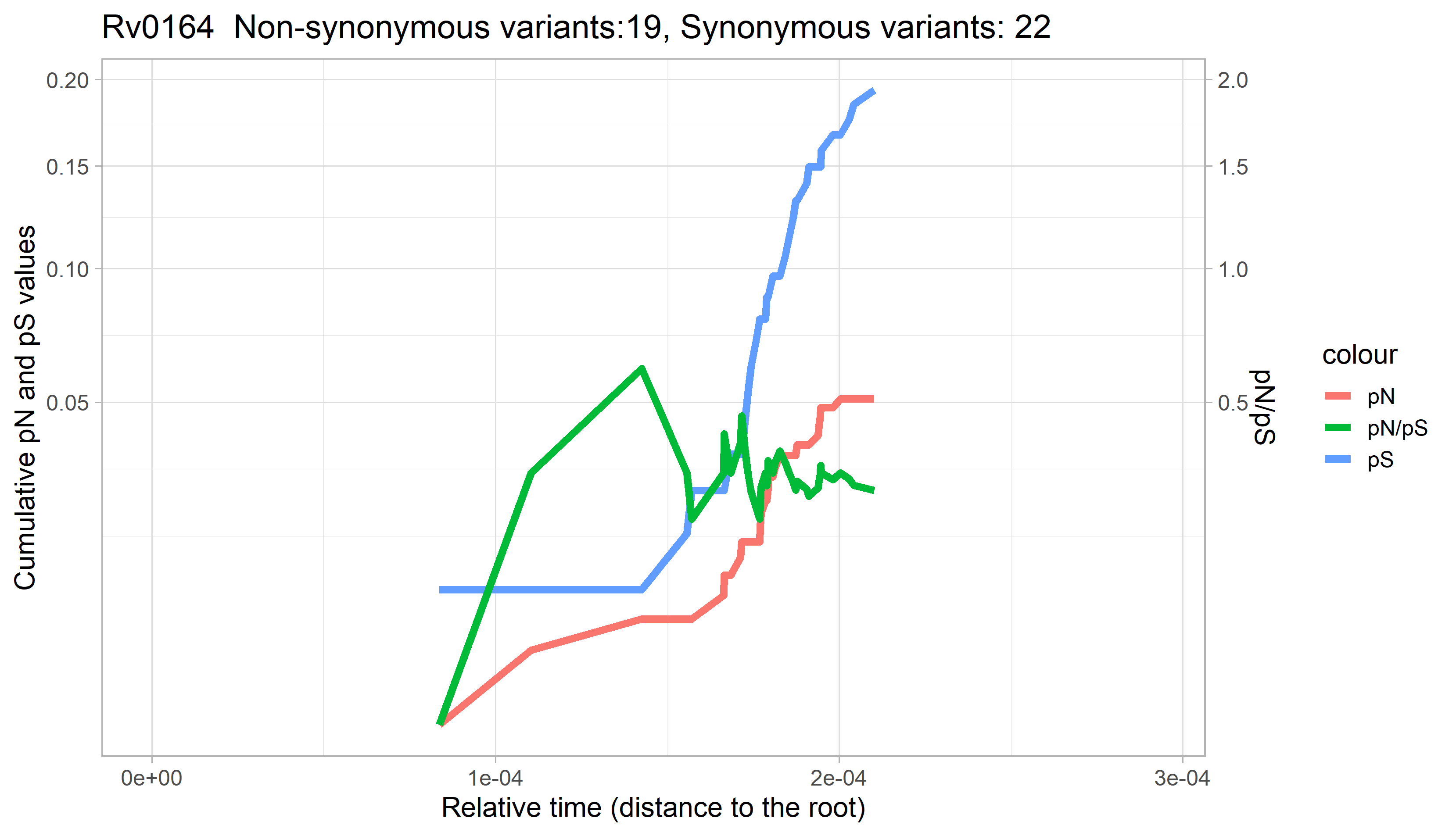

### Rv0165c.png

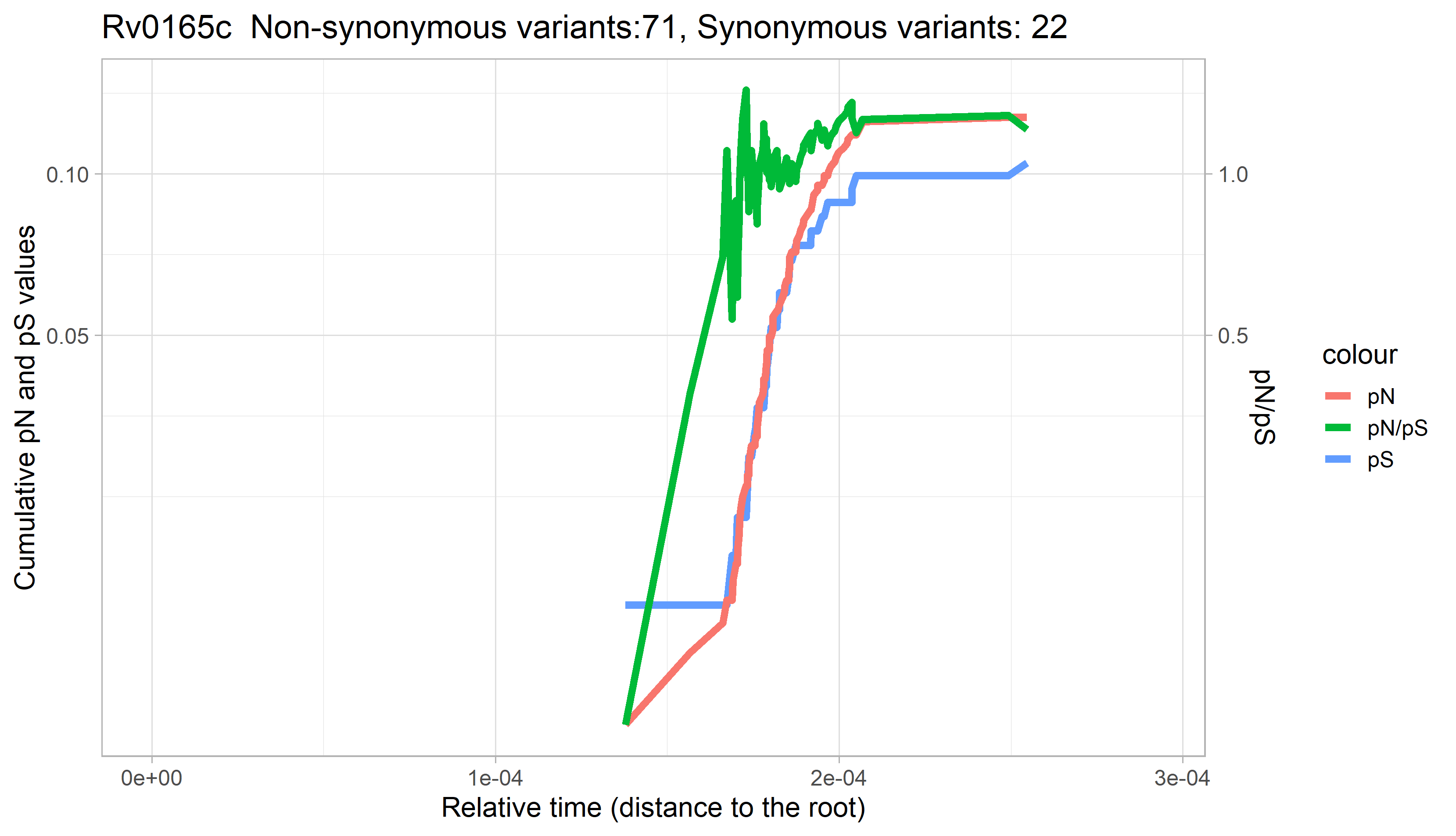

### Rv0166.png

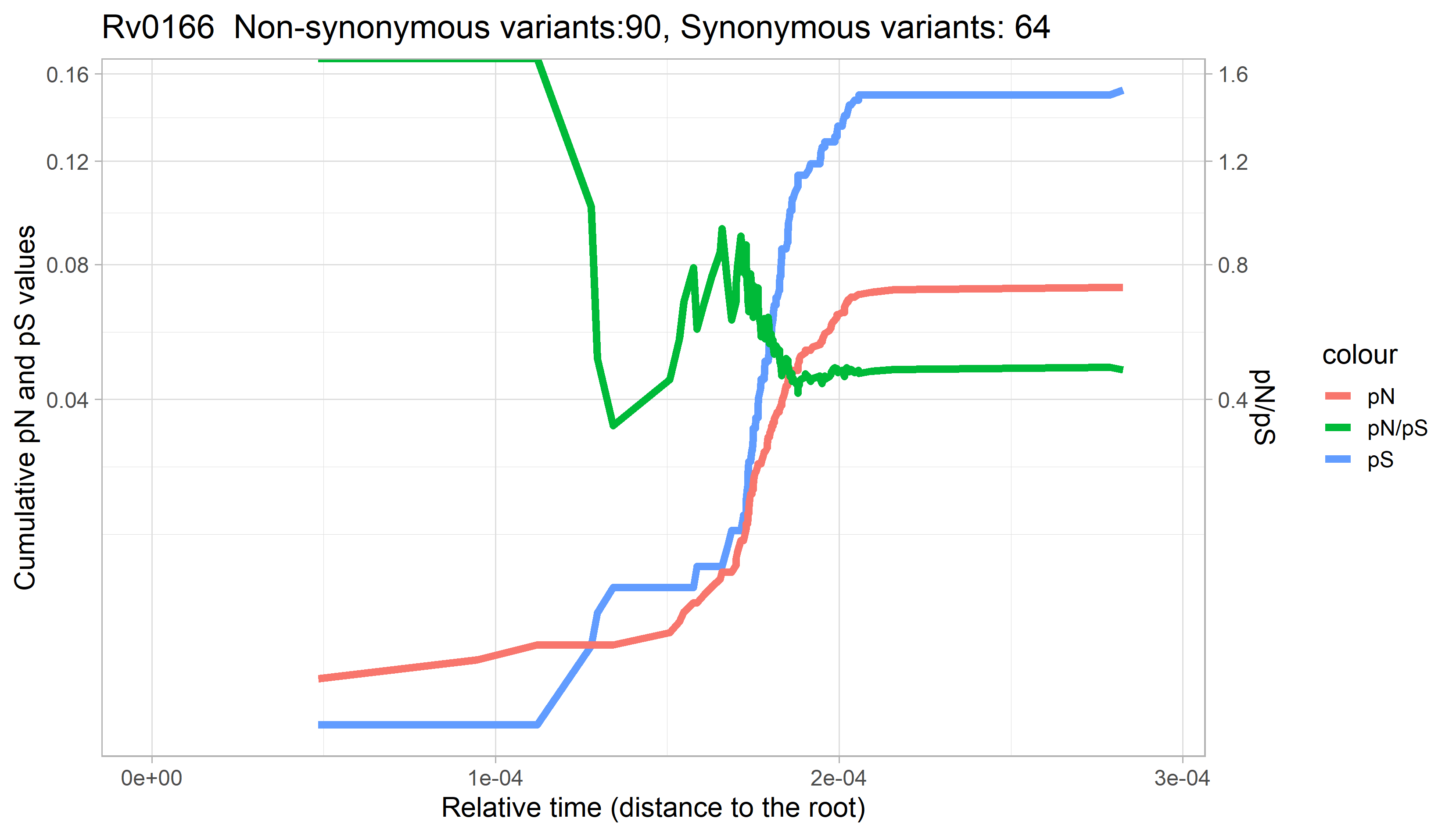

### Rv0167.png

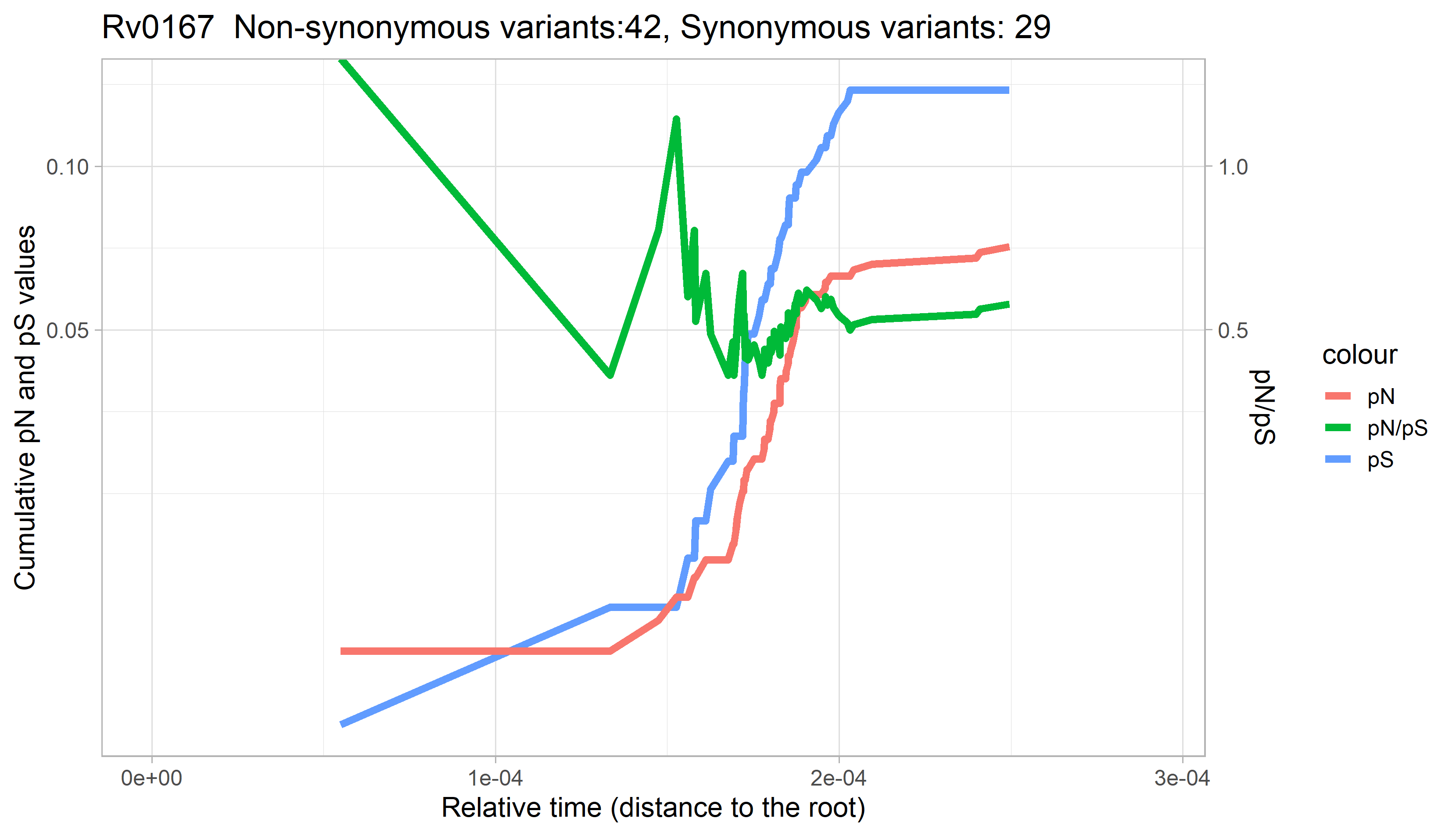

### Rv0168.png

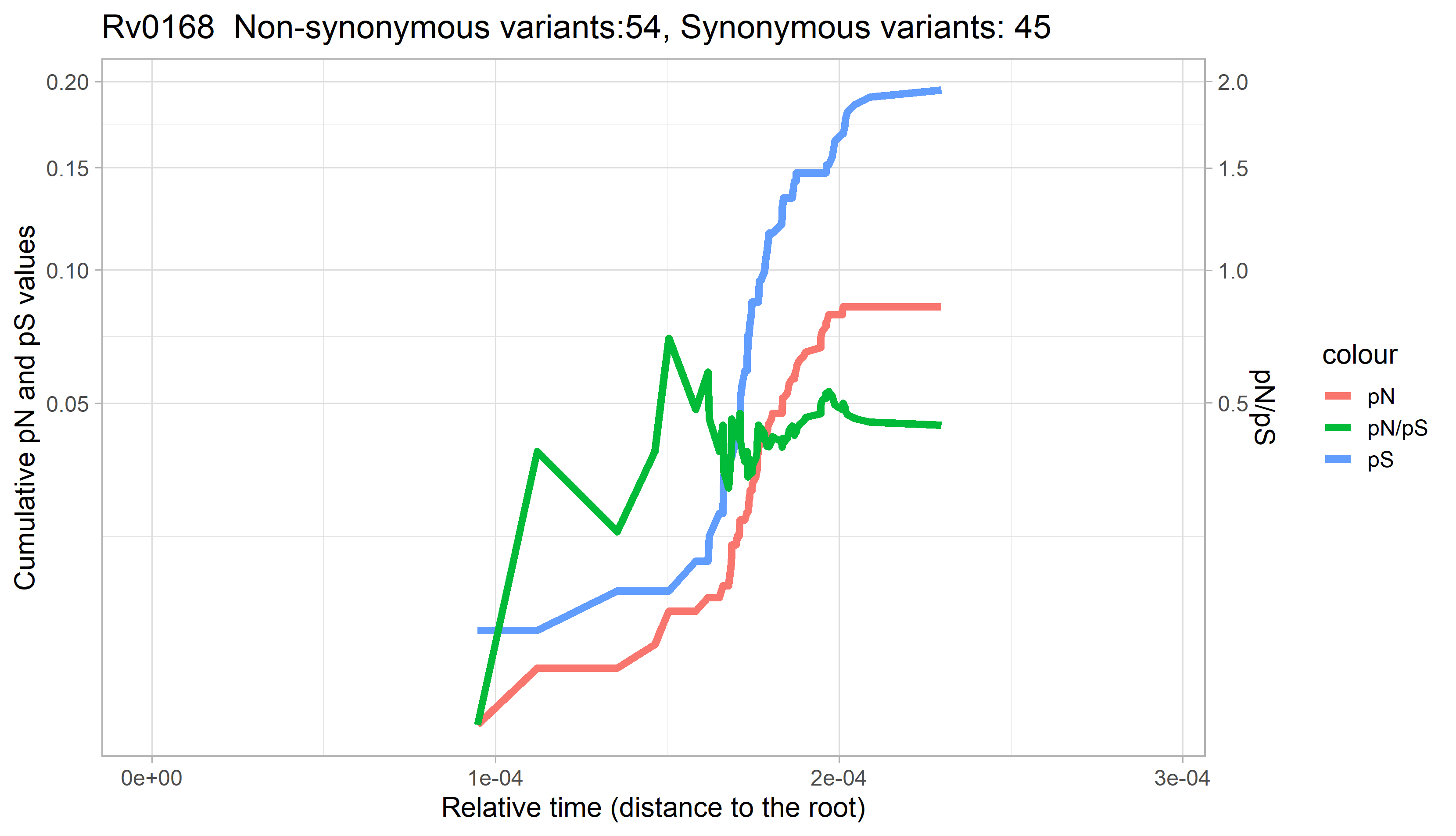

### Rv0169.png

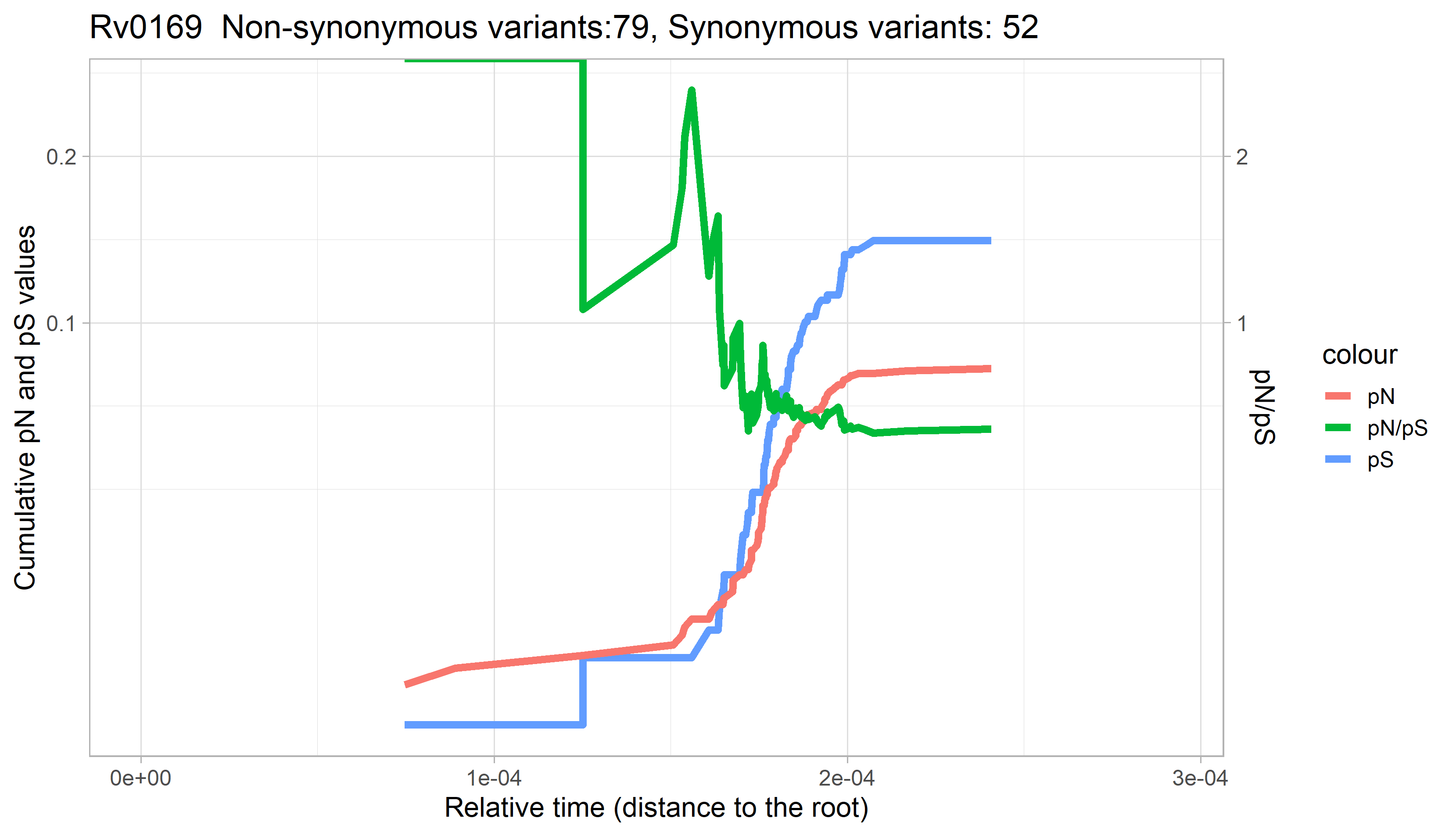

### Rv0170.png

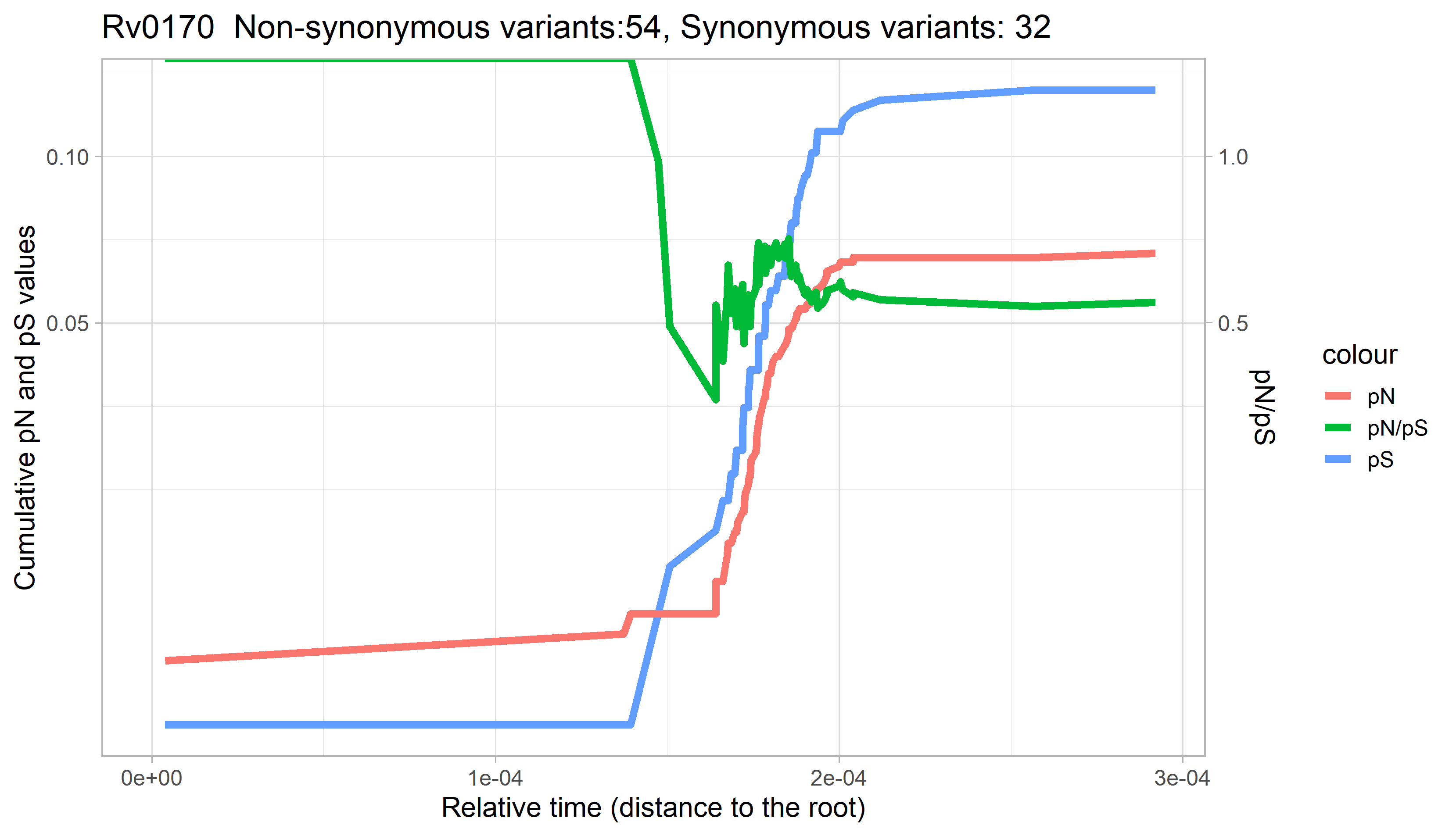

### Rv0171.png

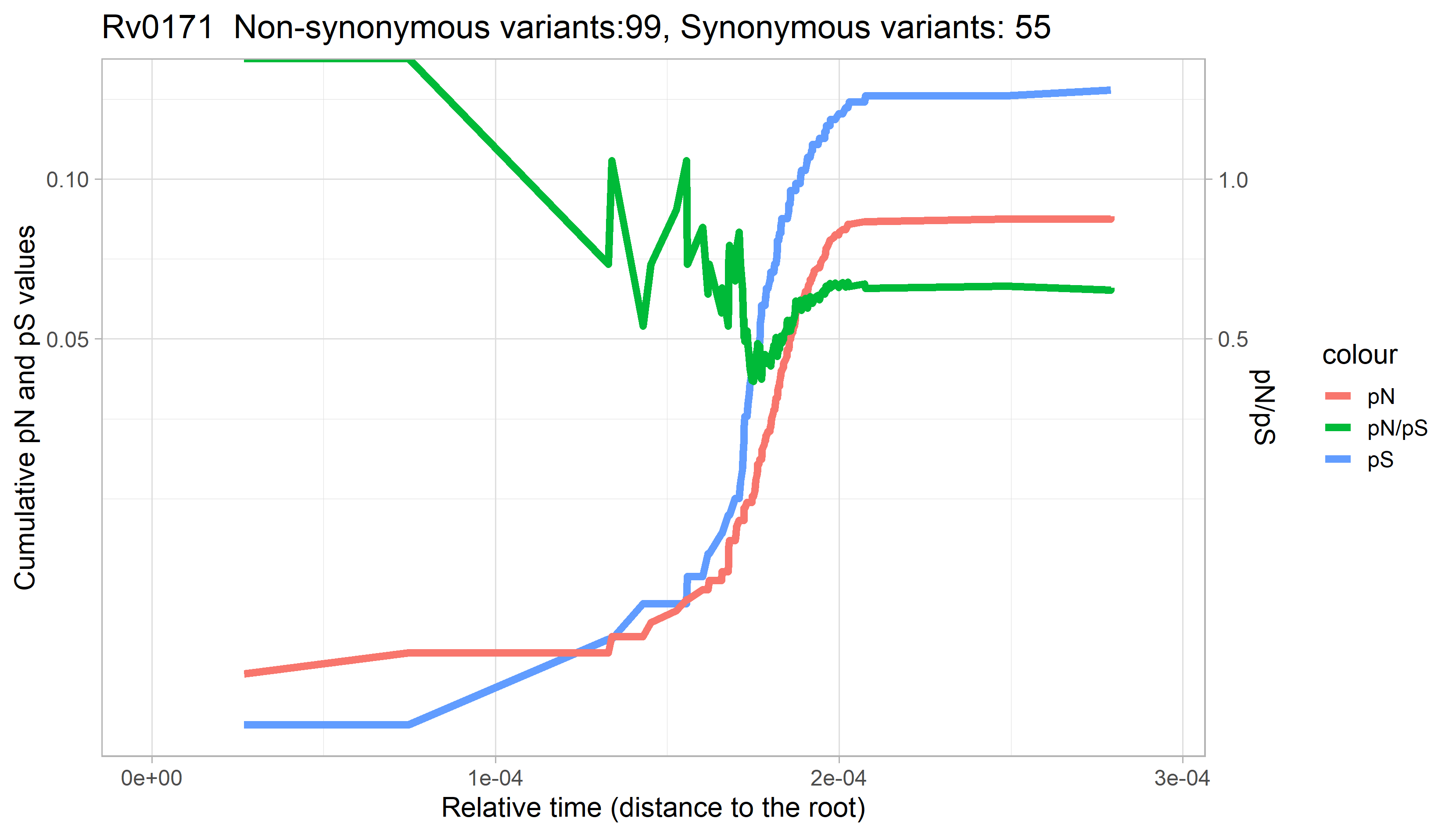

### Rv0172.png

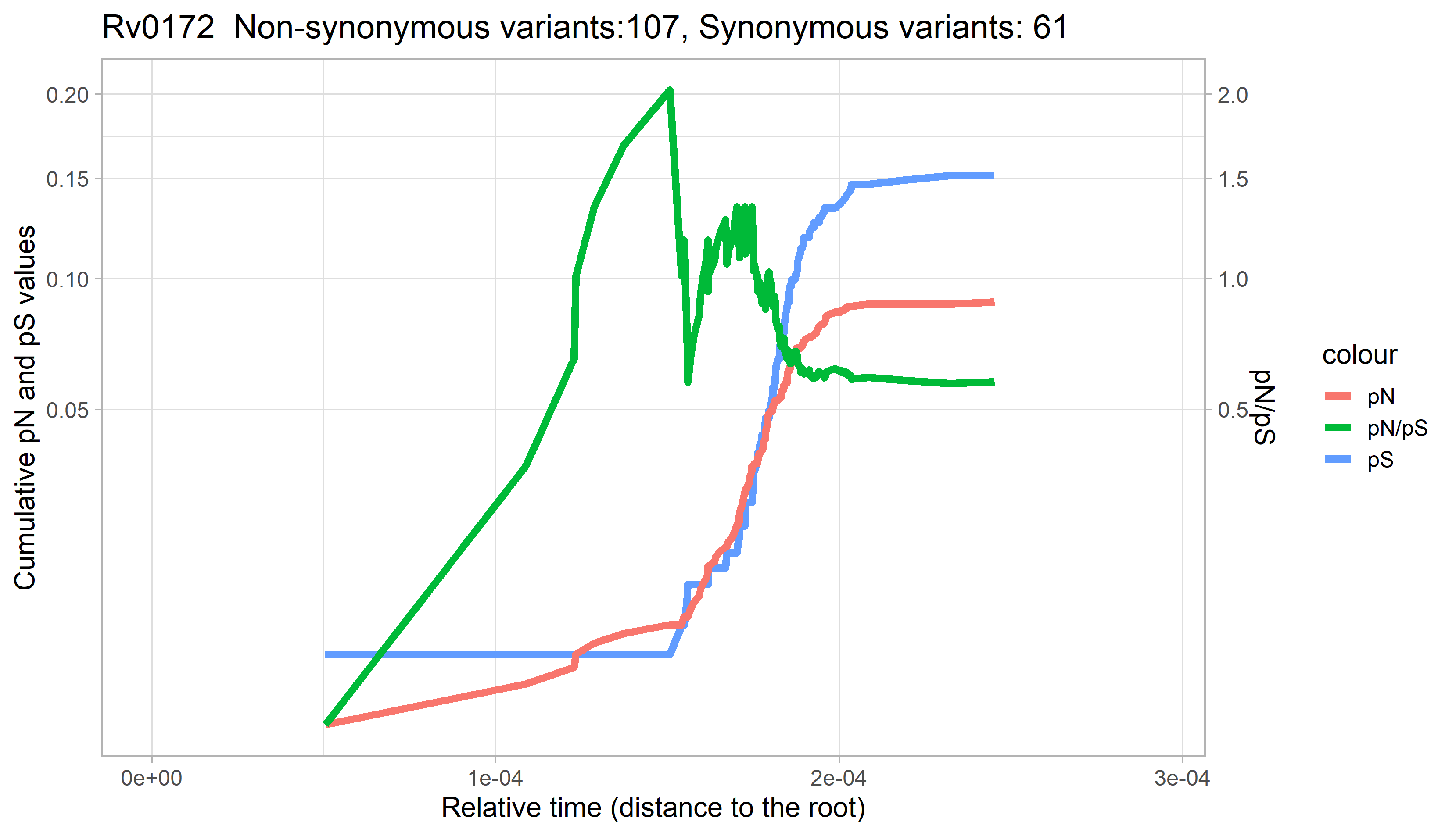

### Rv0173.png

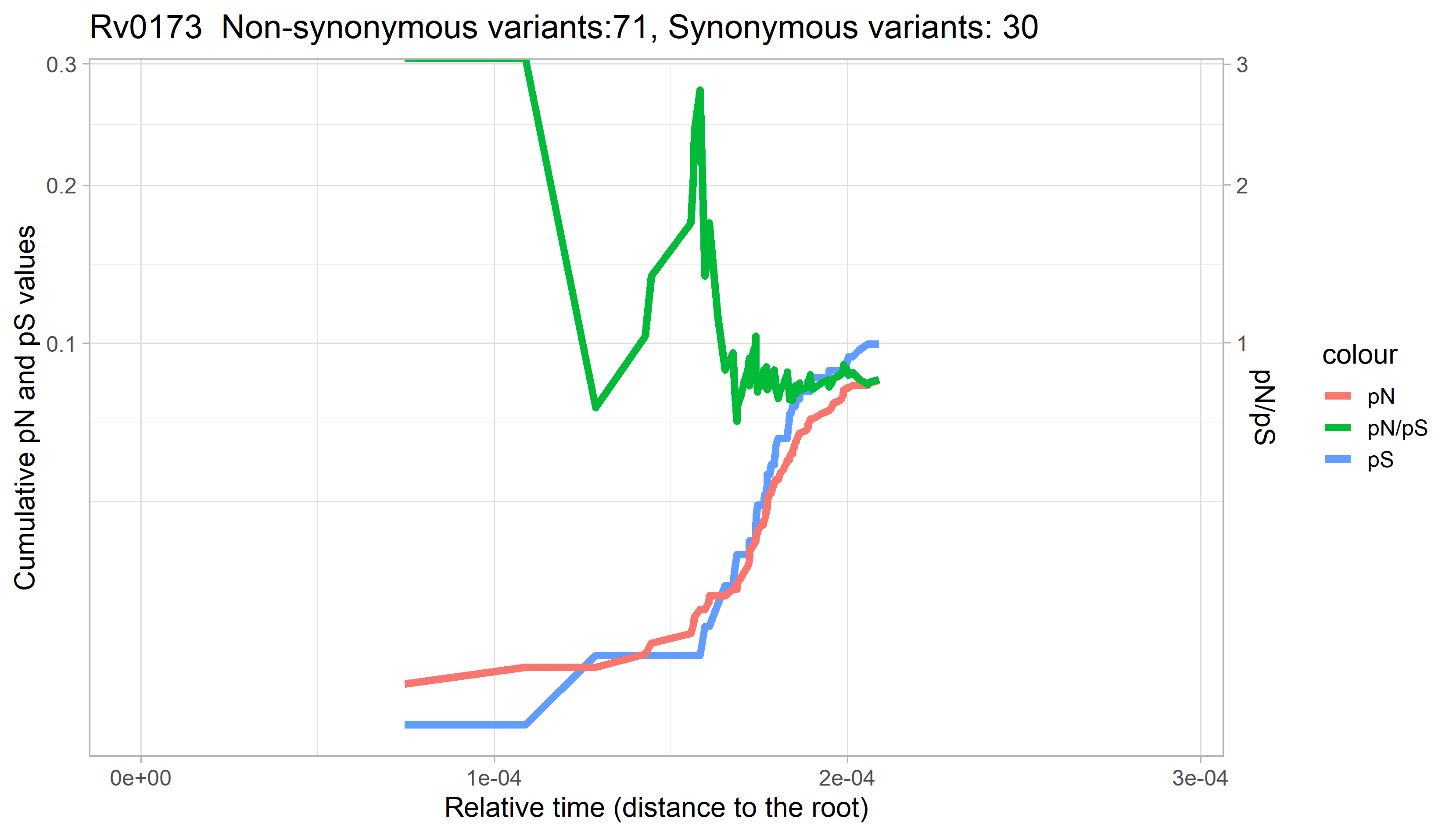

### Rv0174.png

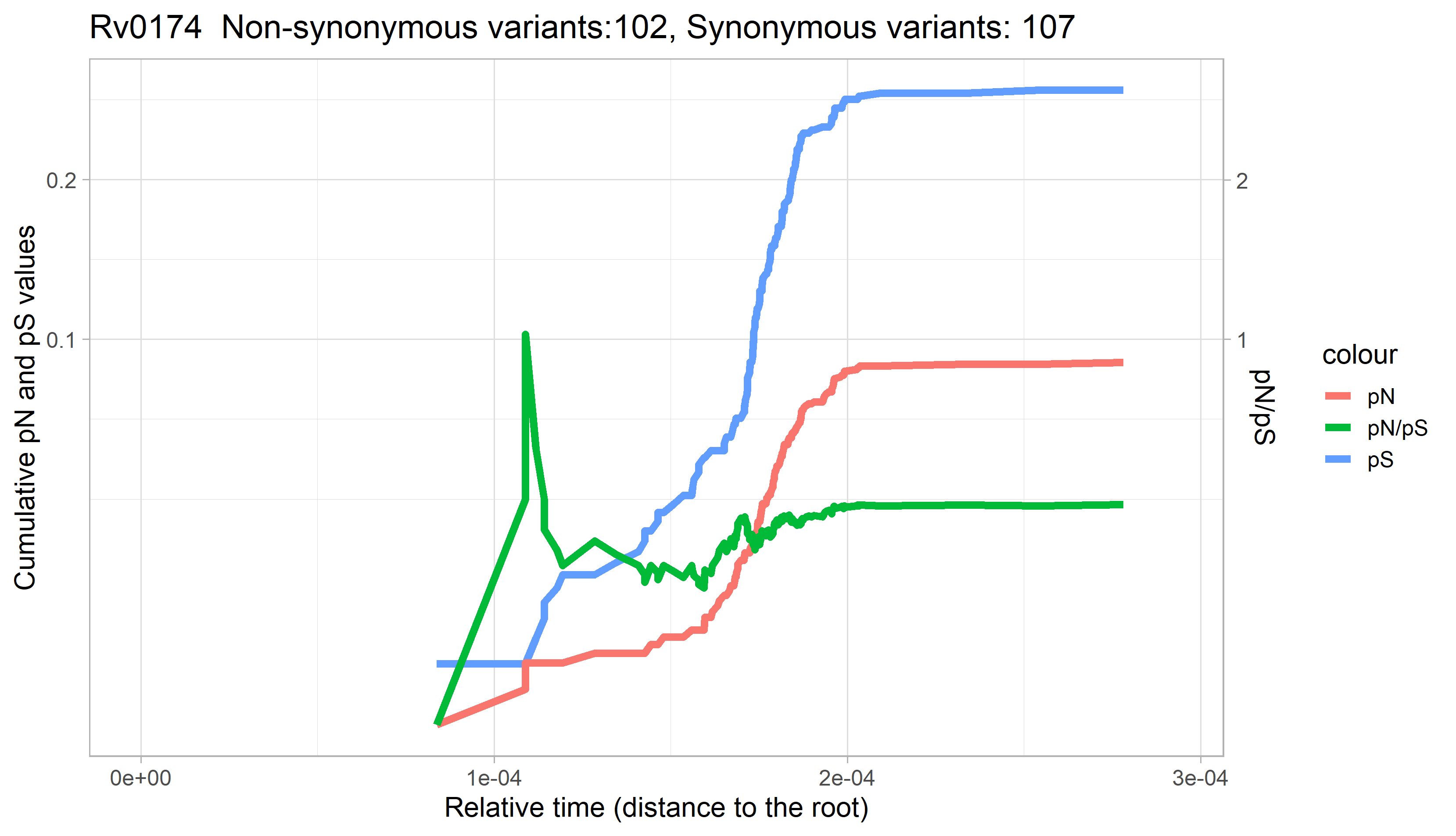

### Rv0175.png

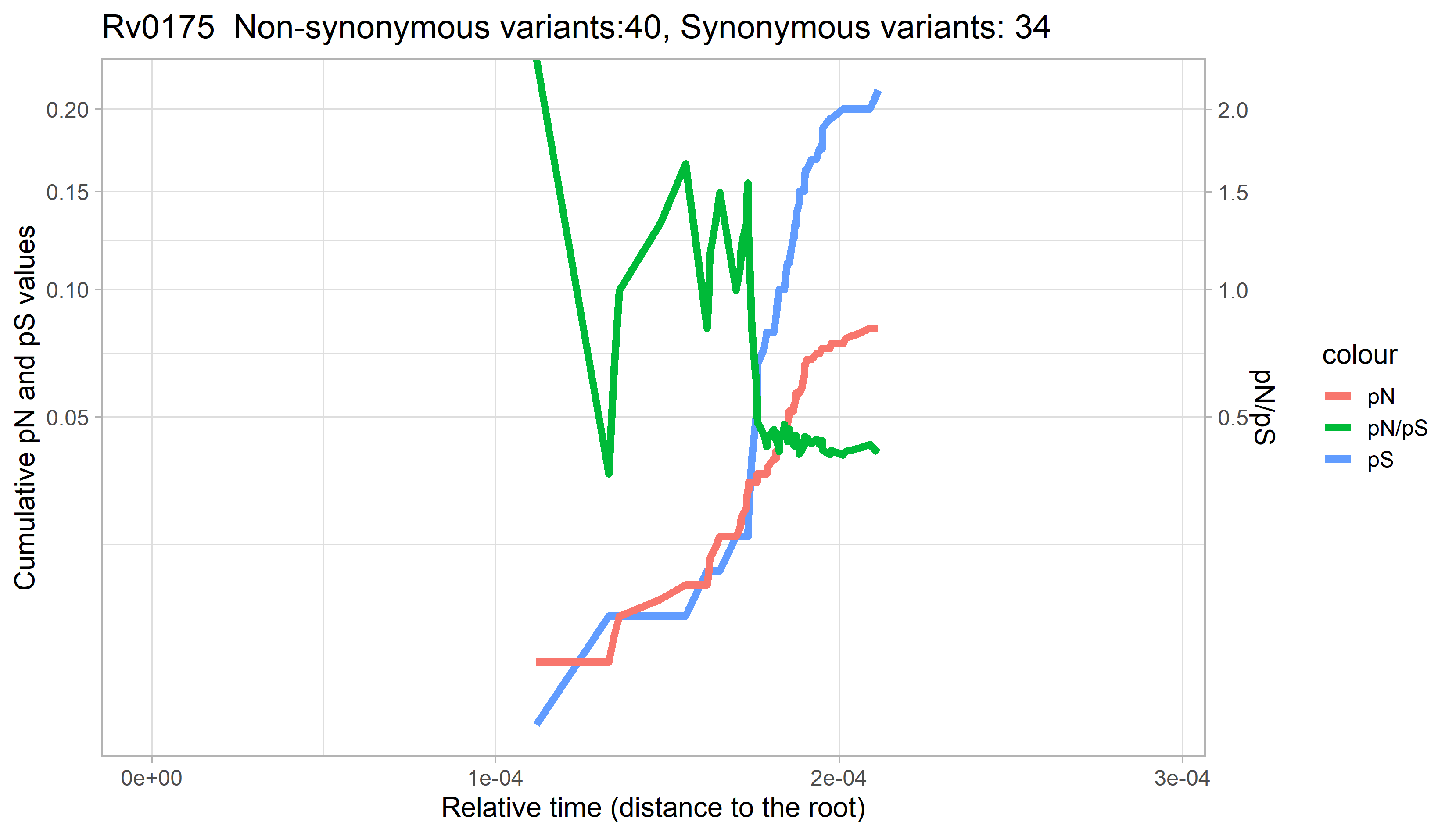

### Rv0176.png

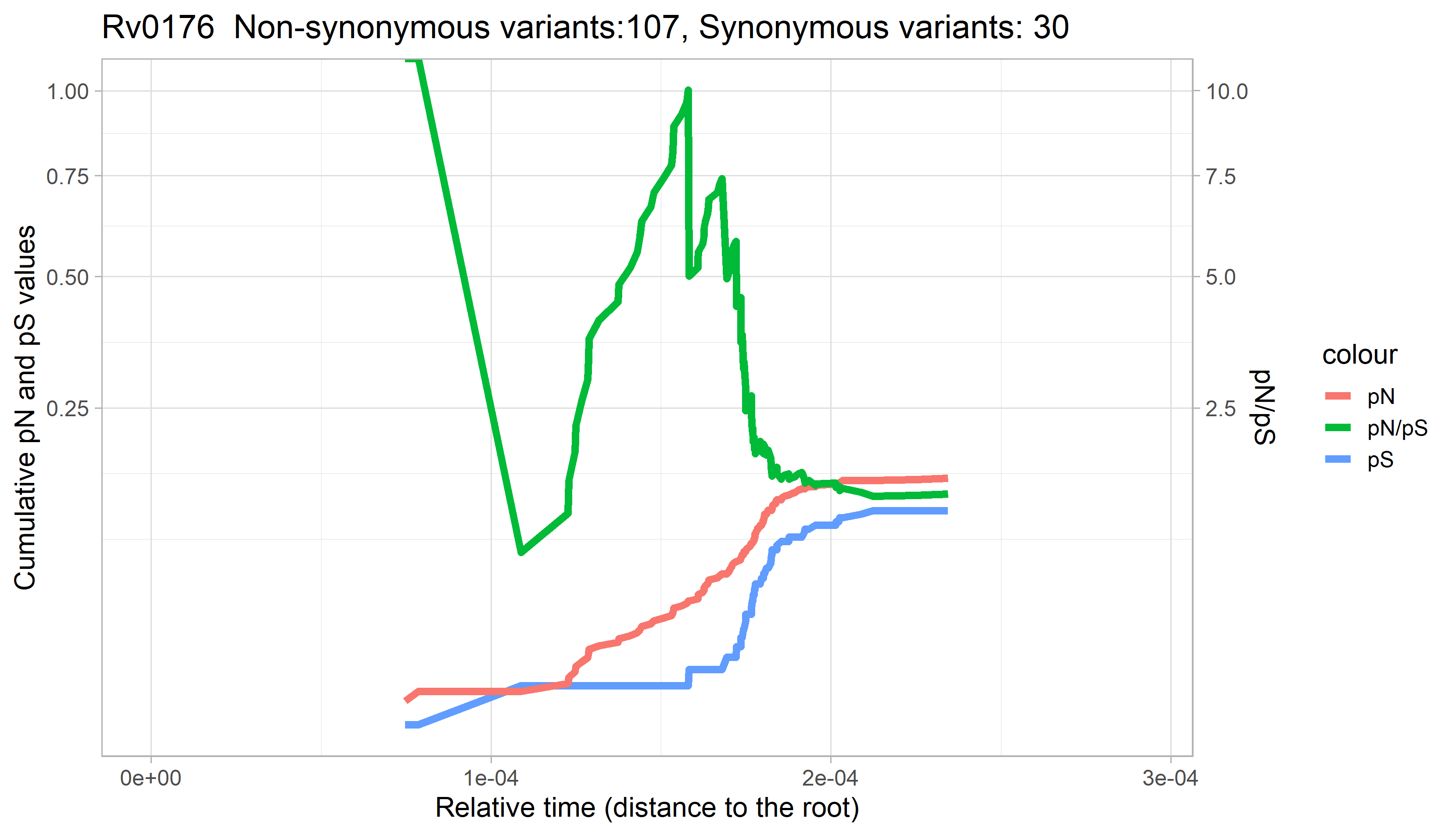

### Rv0177.png

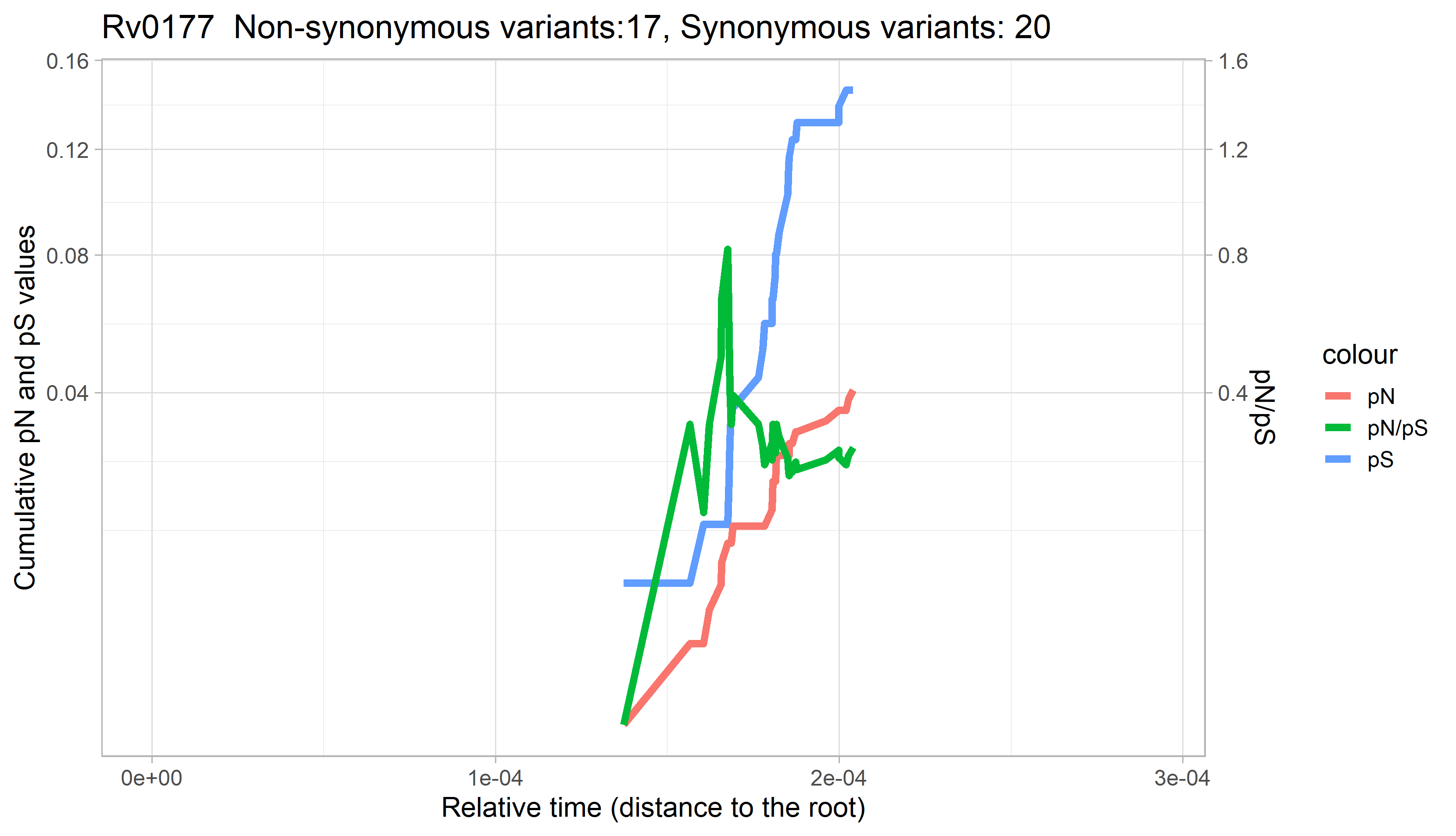

### Rv0178.png

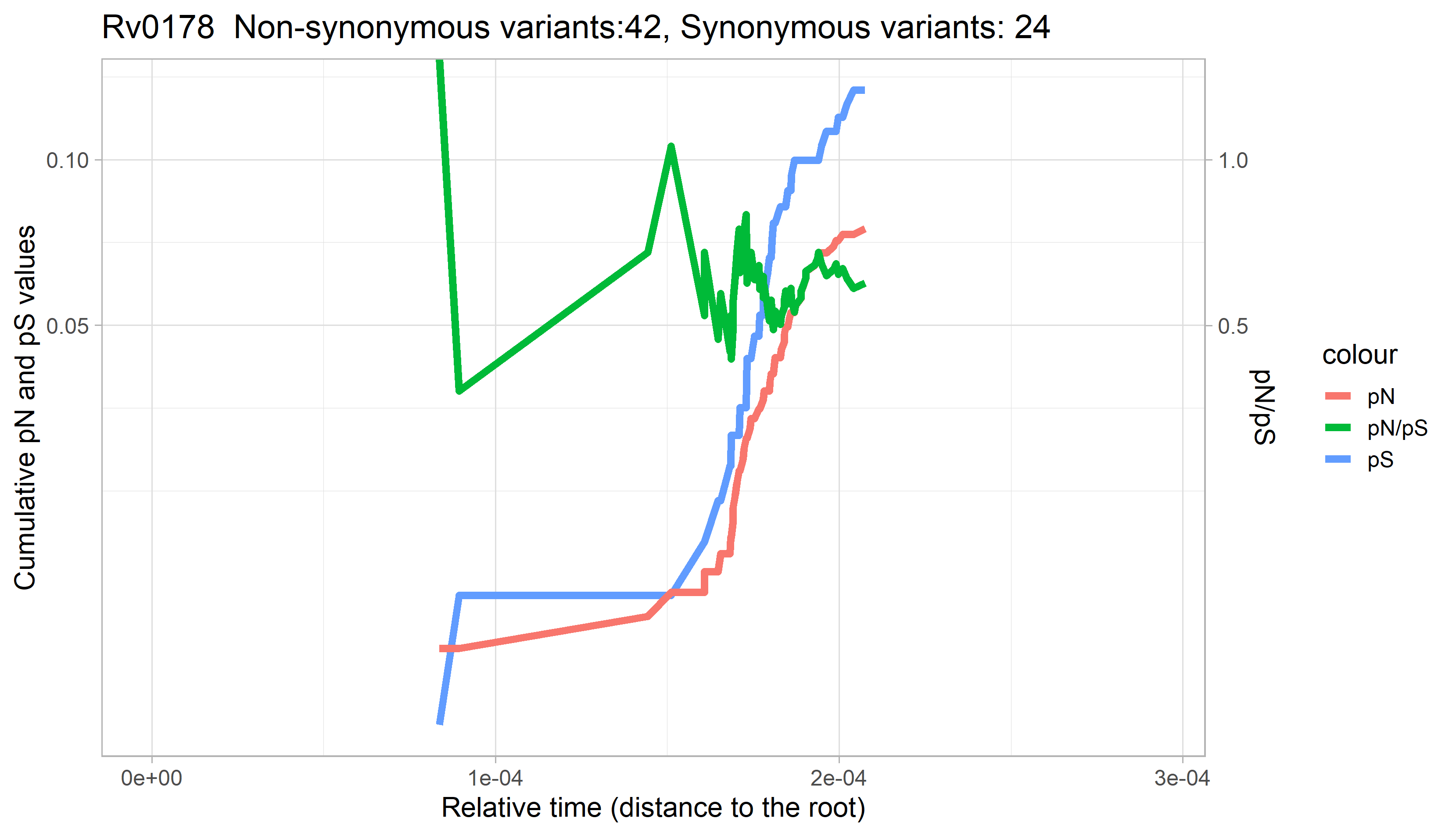

### Rv0179c.png

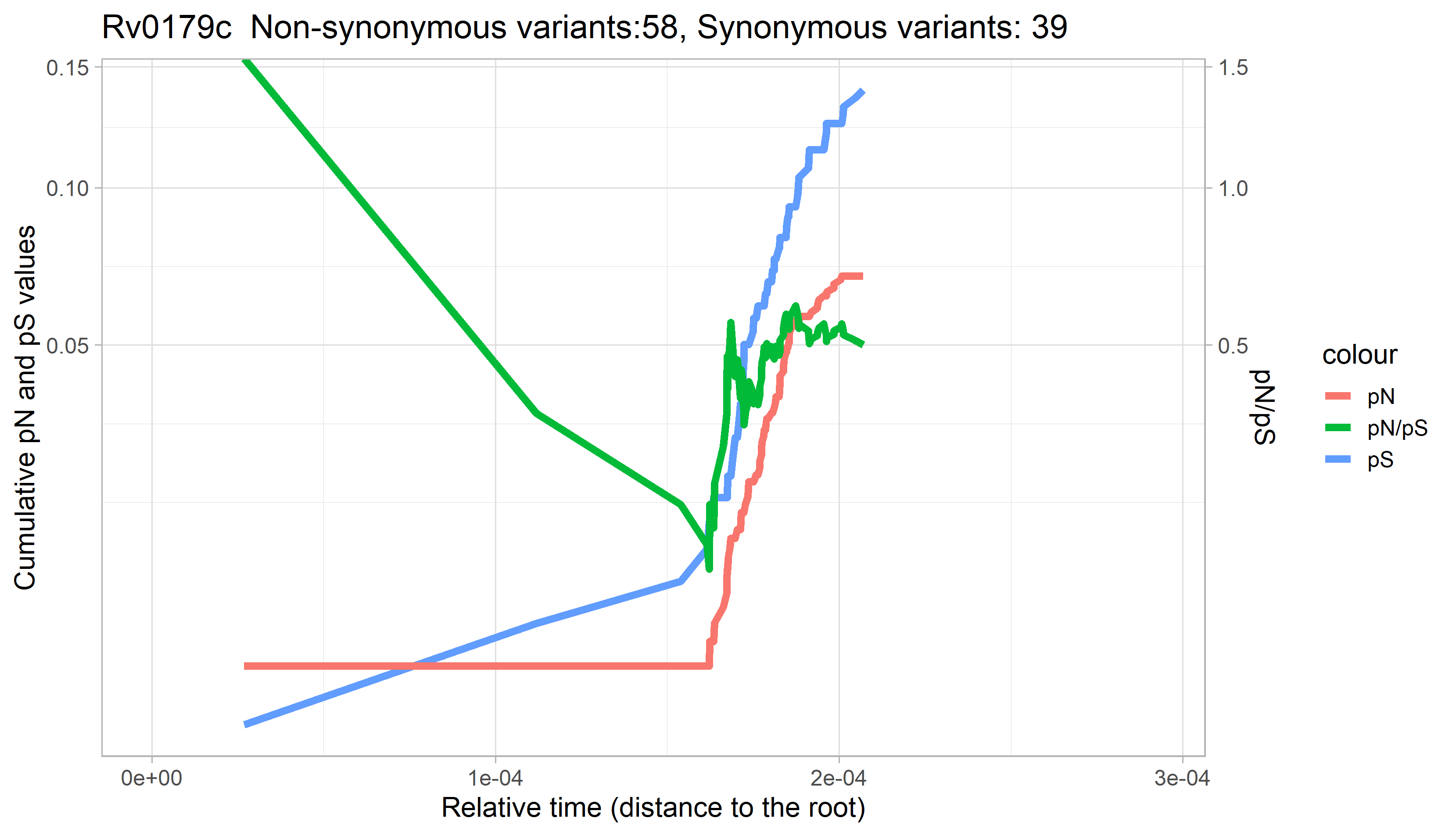

### Rv0180c.png

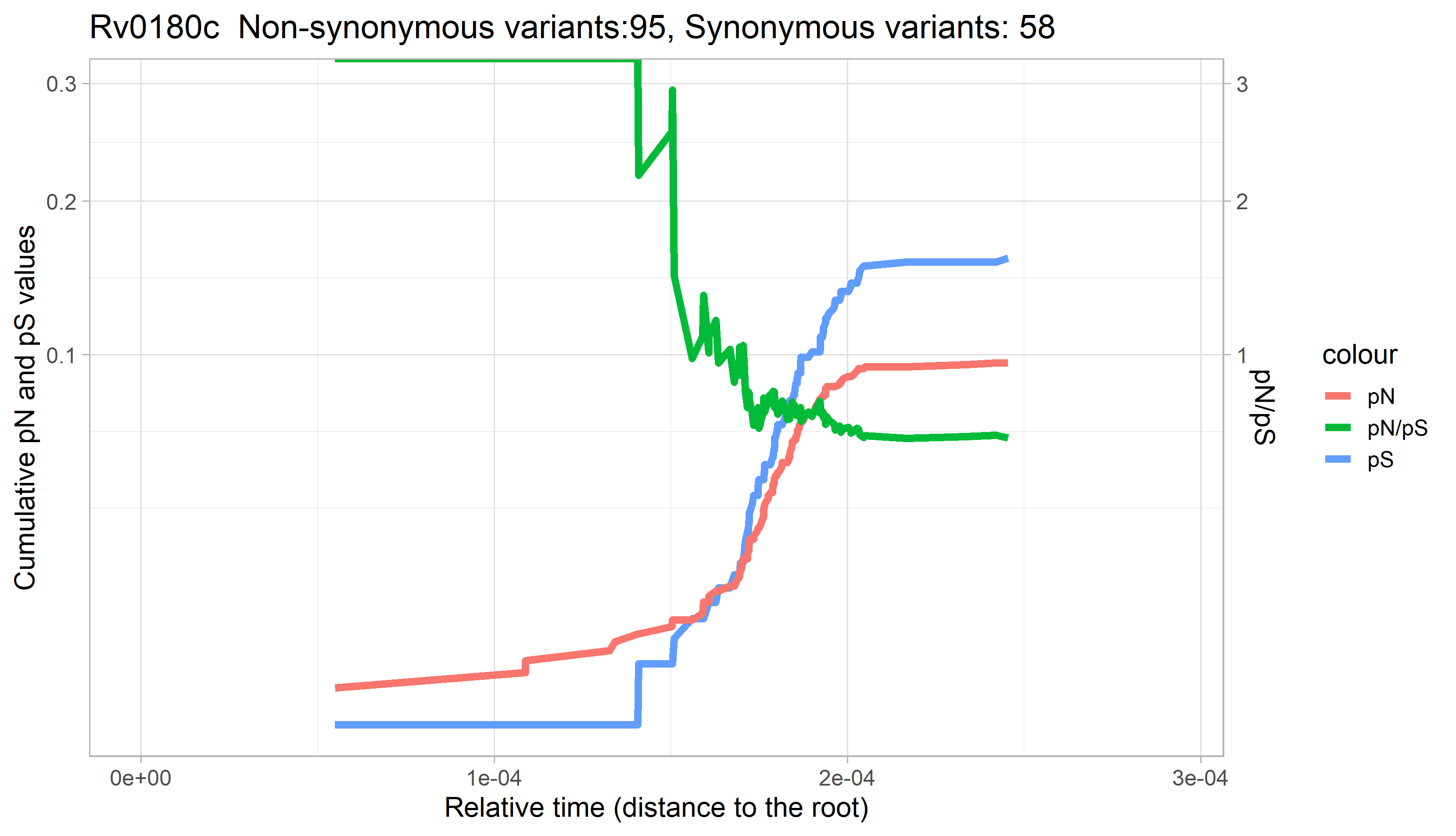

### Rv0181c.png

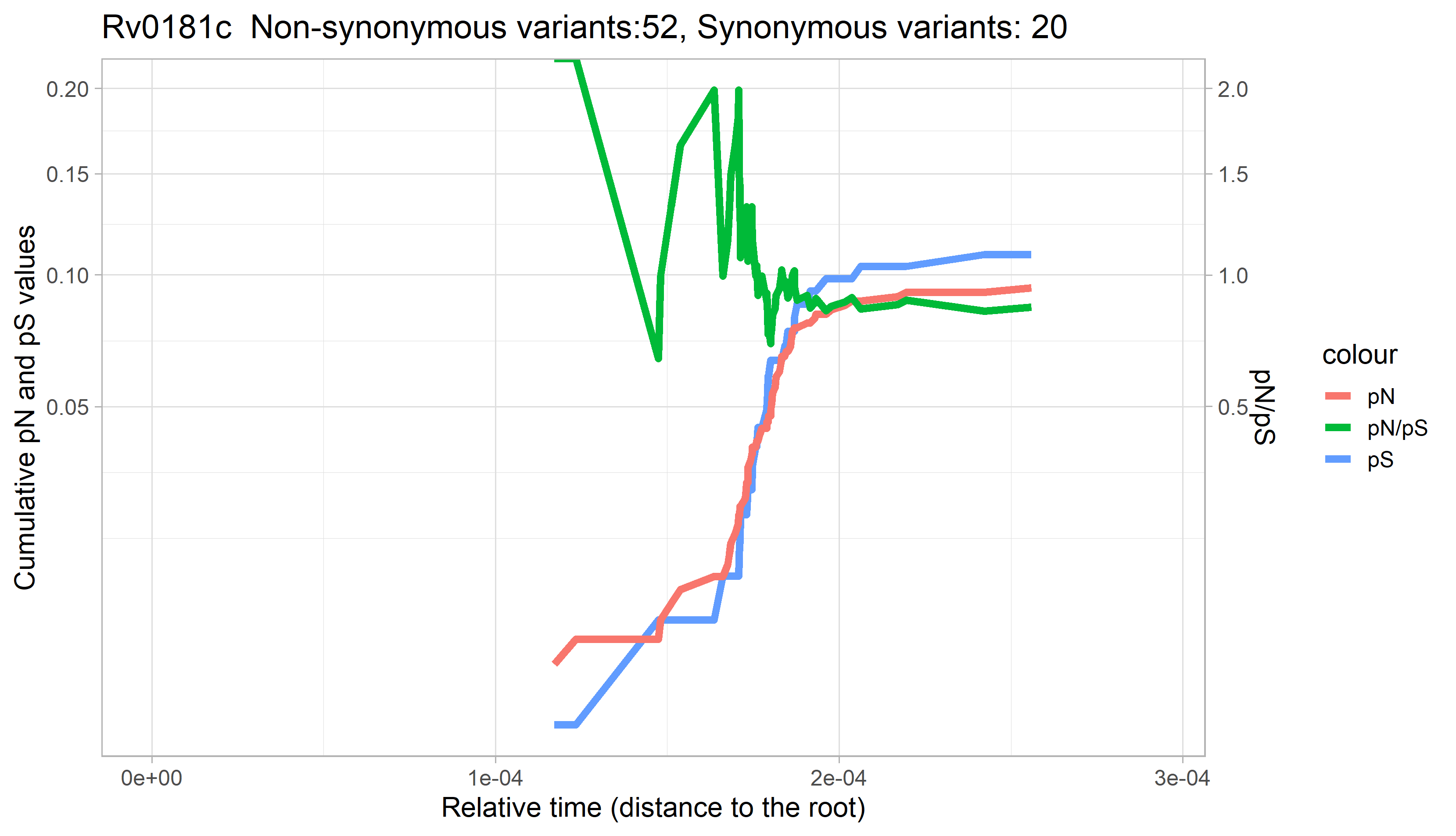

### Rv0182c.png

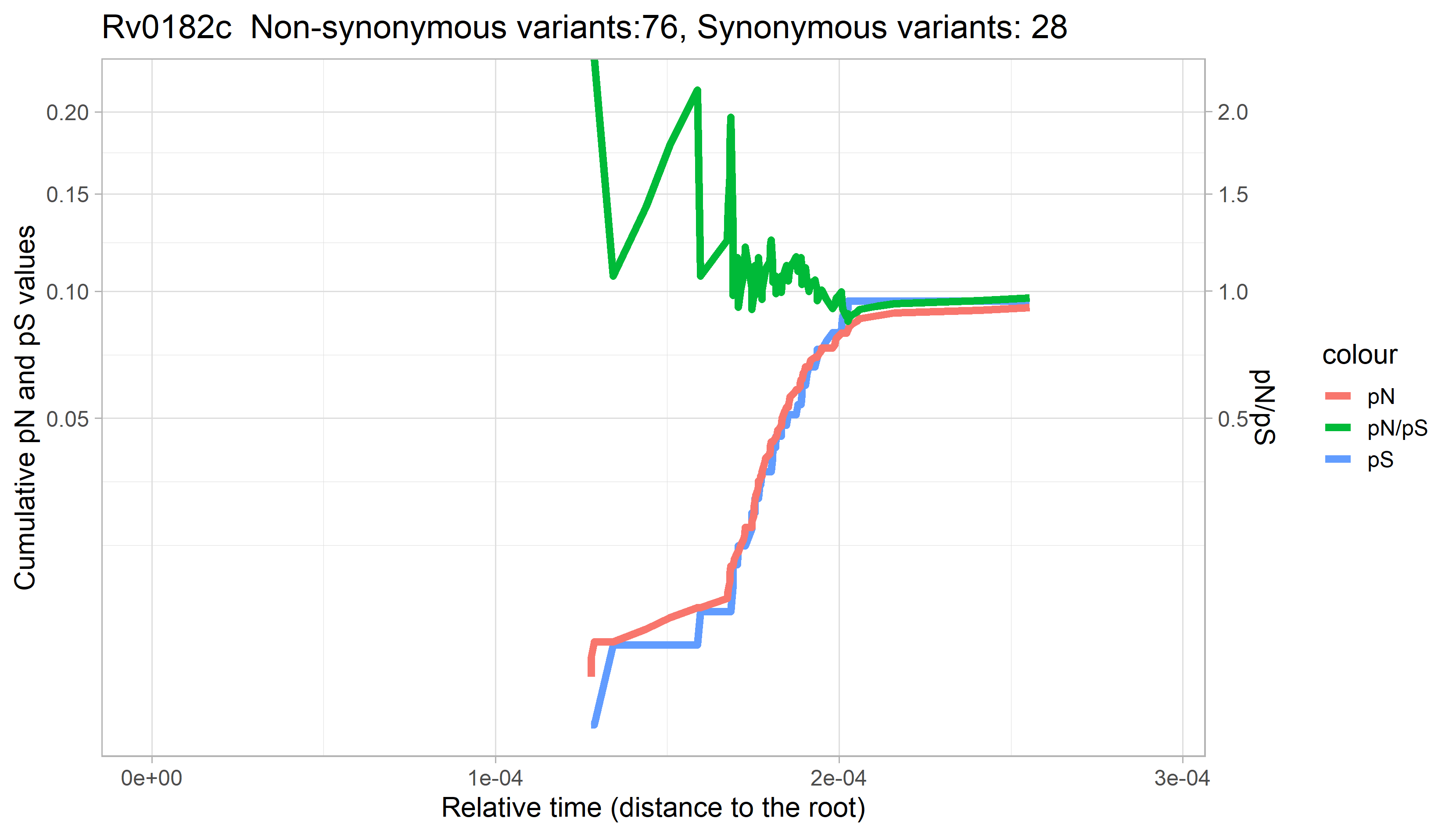

### Rv0183.png

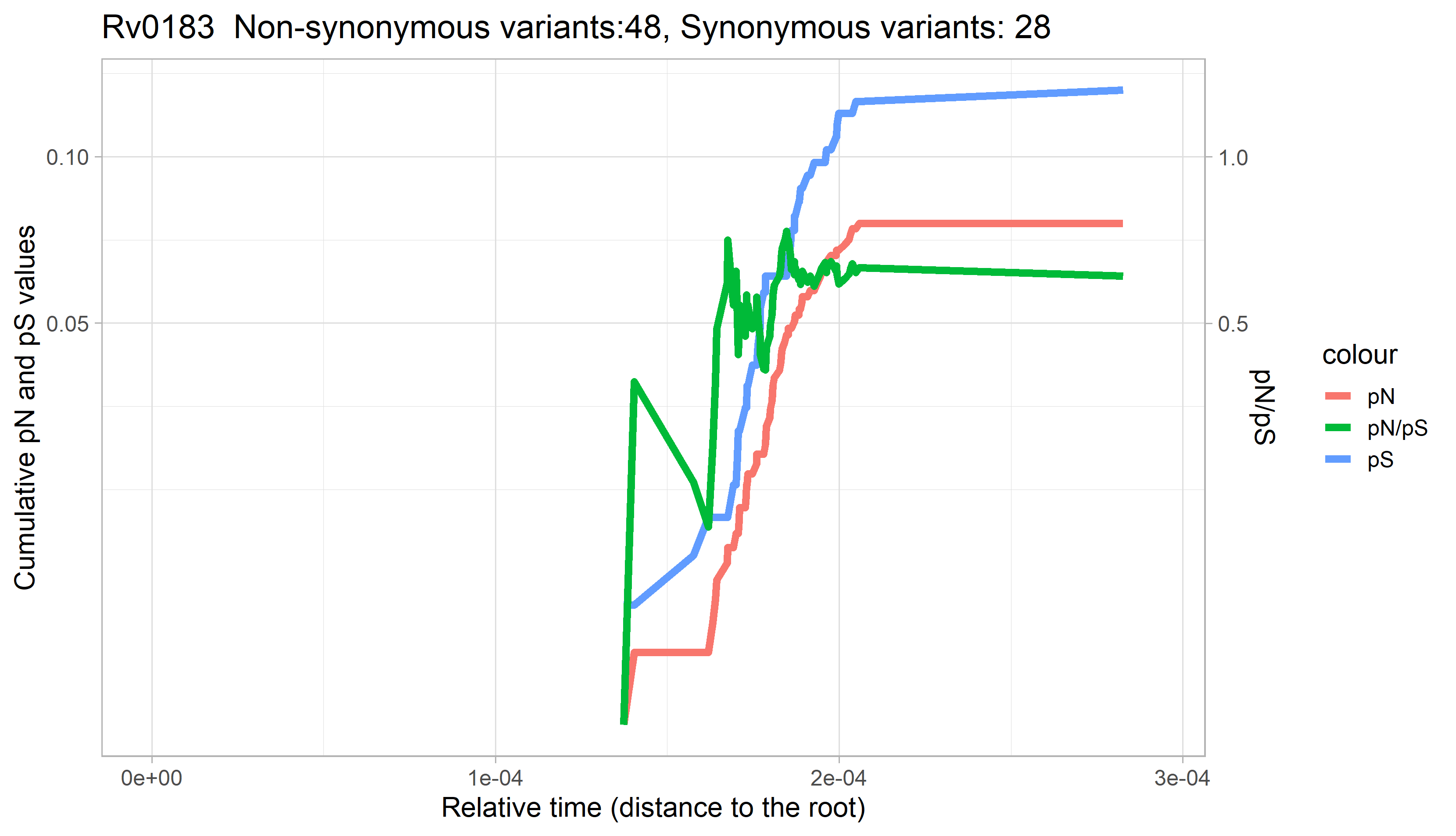

### Rv0184.png

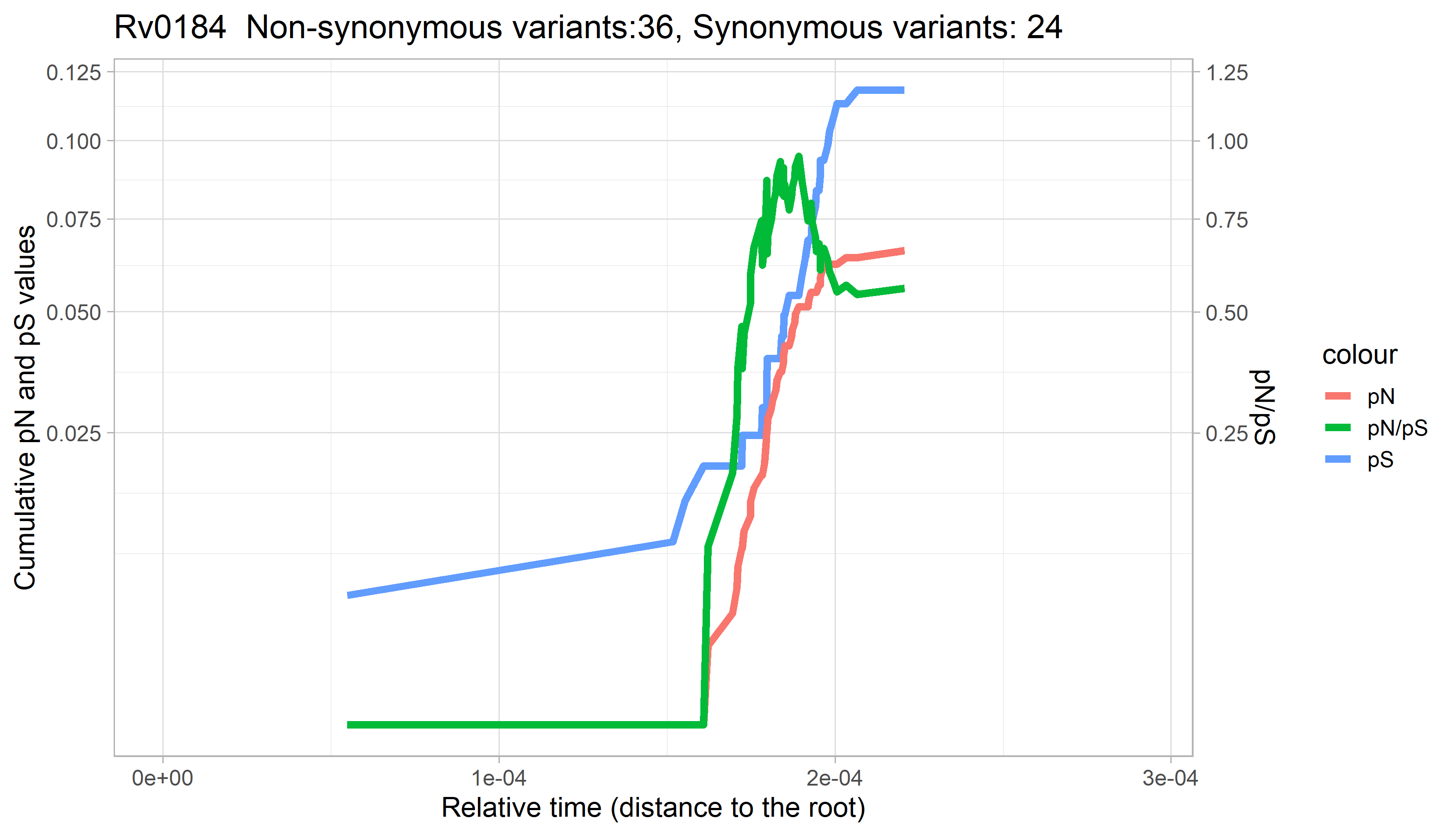

### Rv0185.png

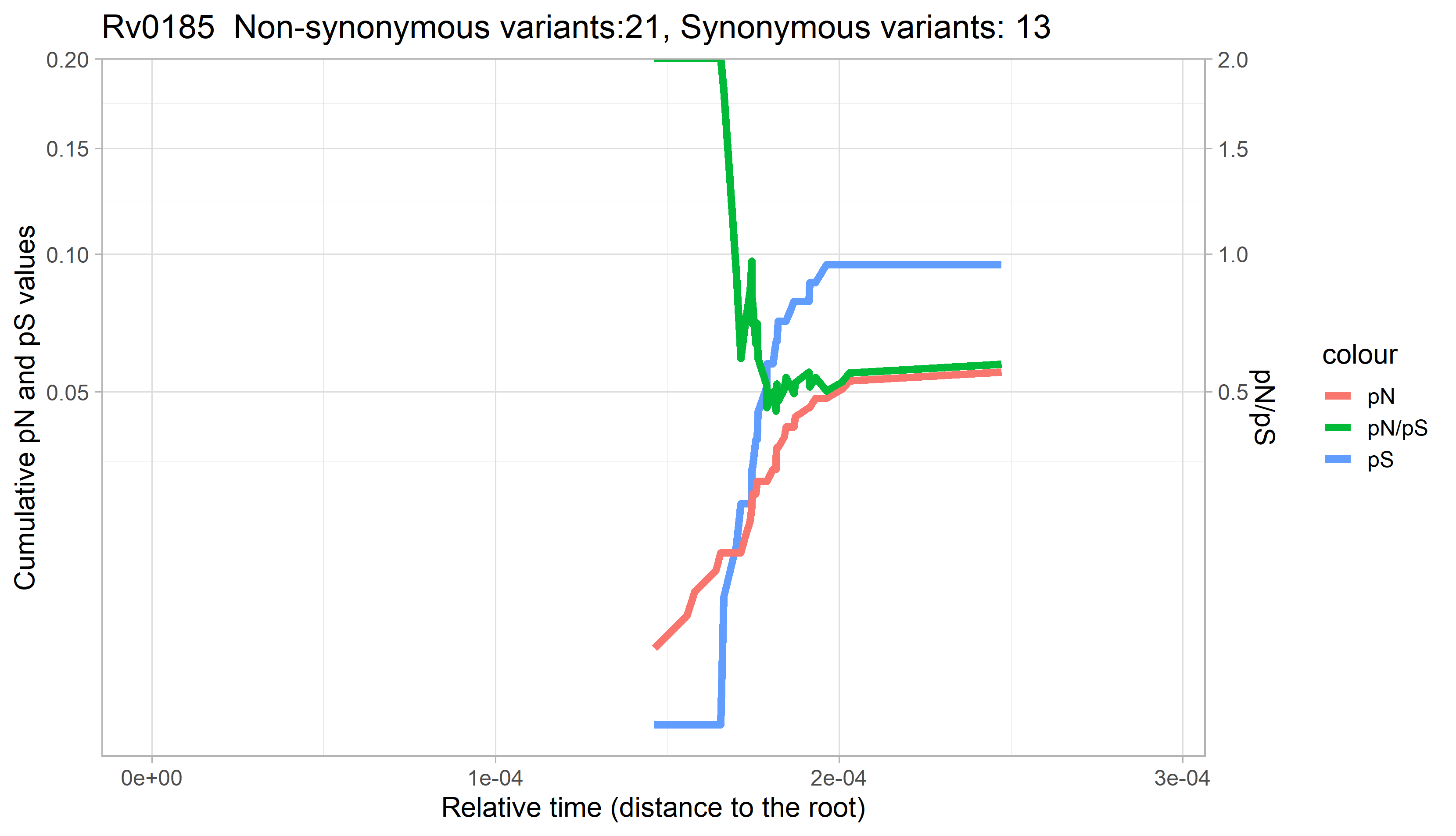

### Rv0186.png

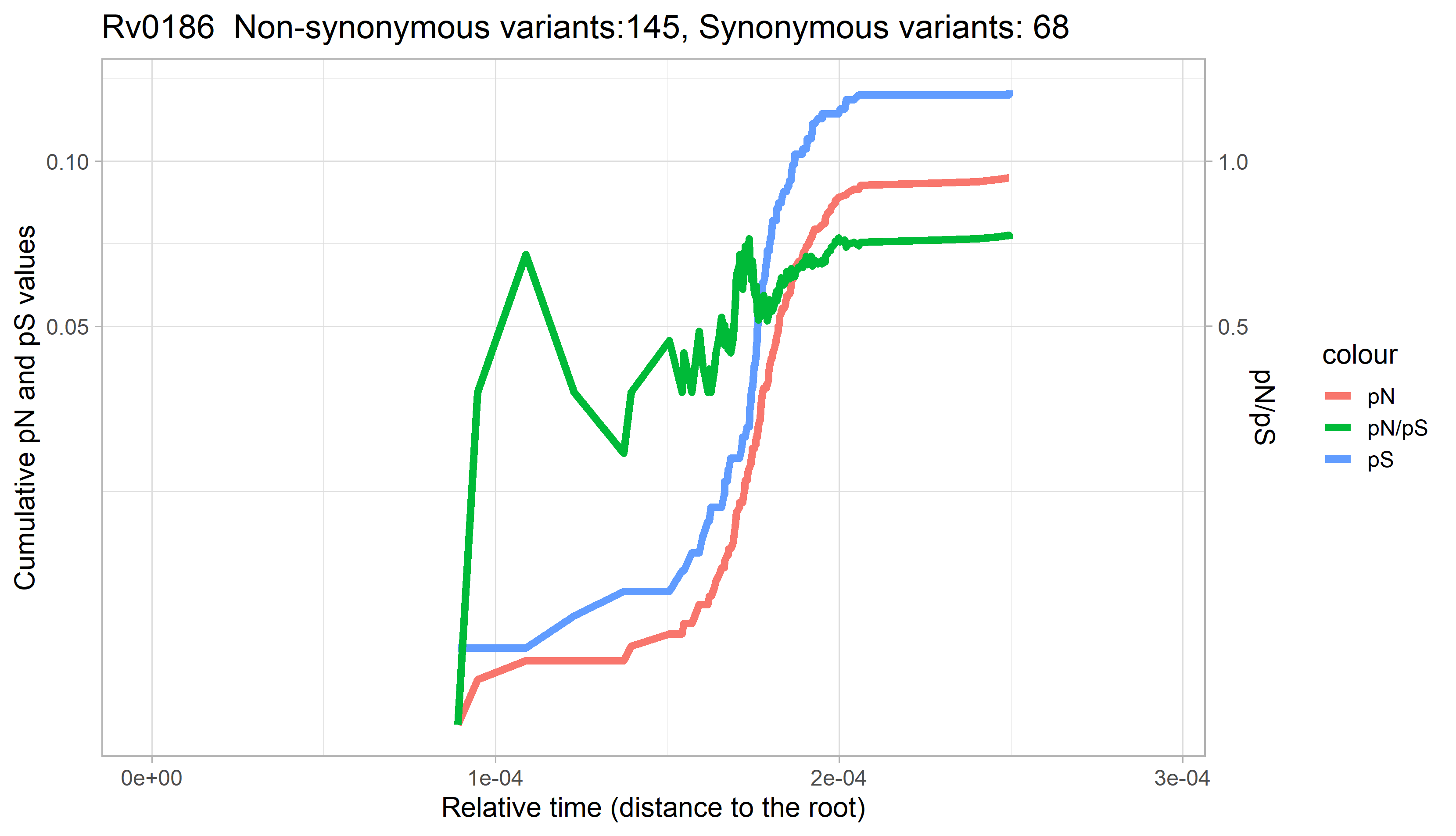

### Rv0187.png

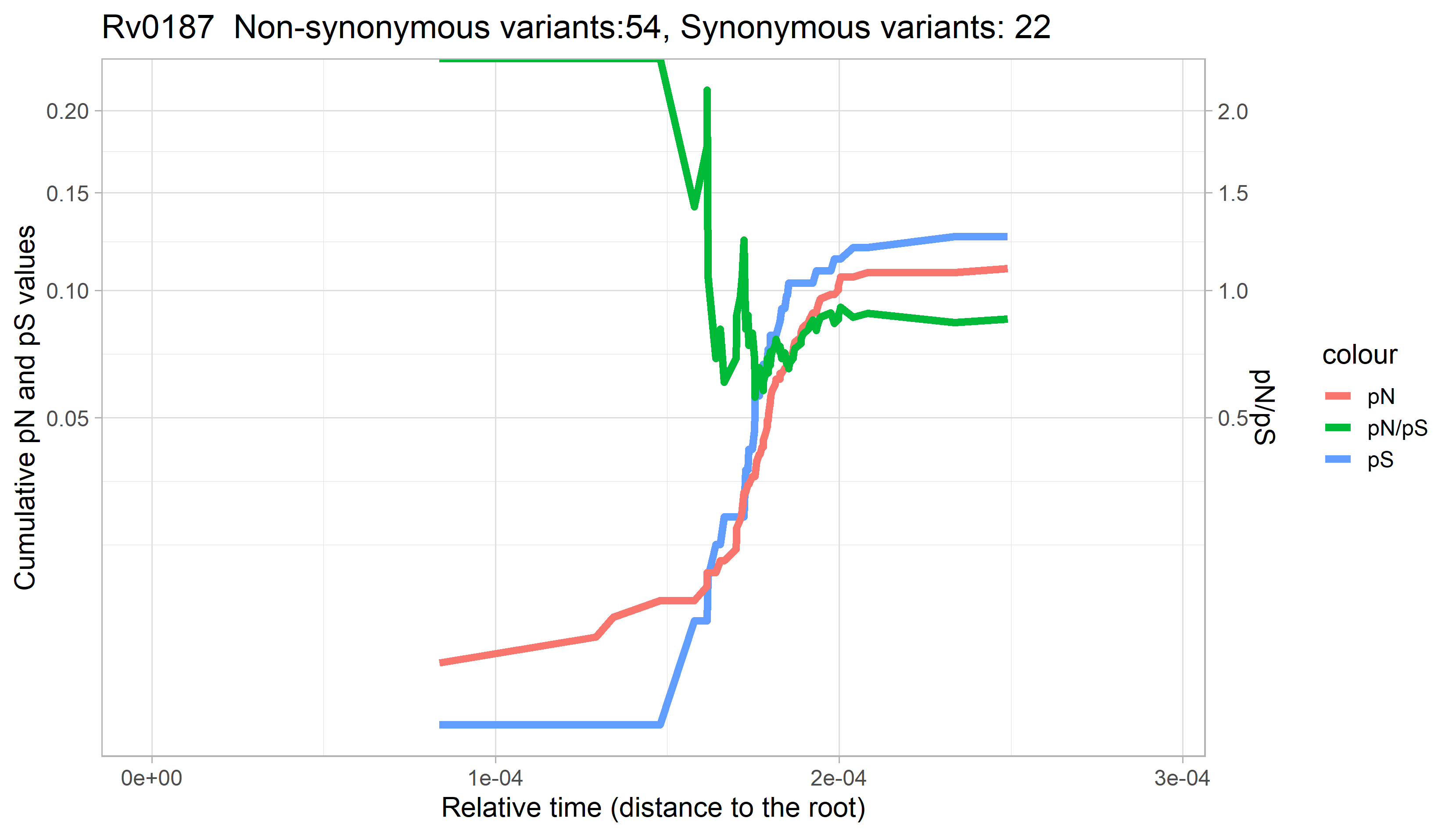

### Rv0188.png

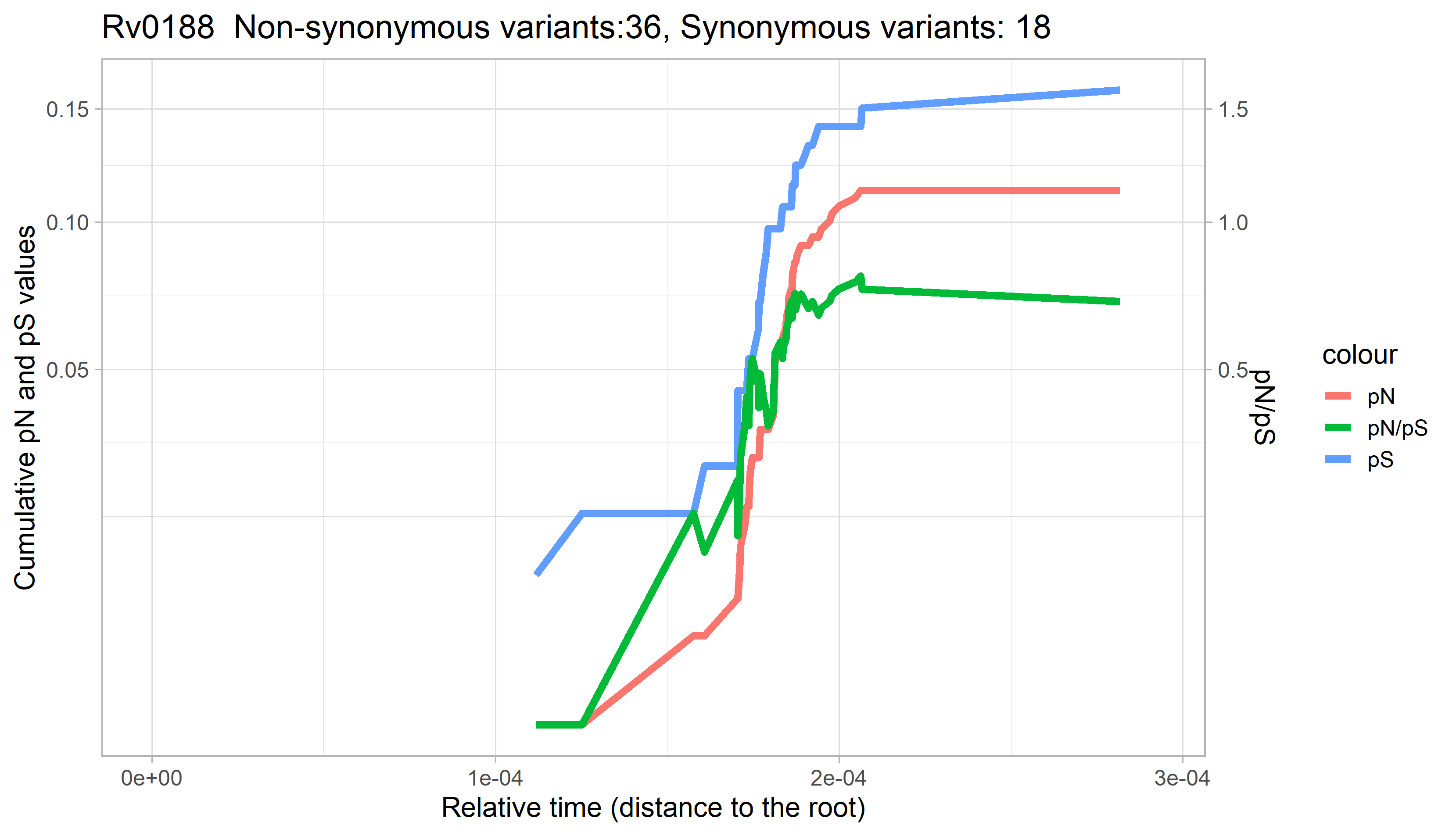

### Rv0189c.png

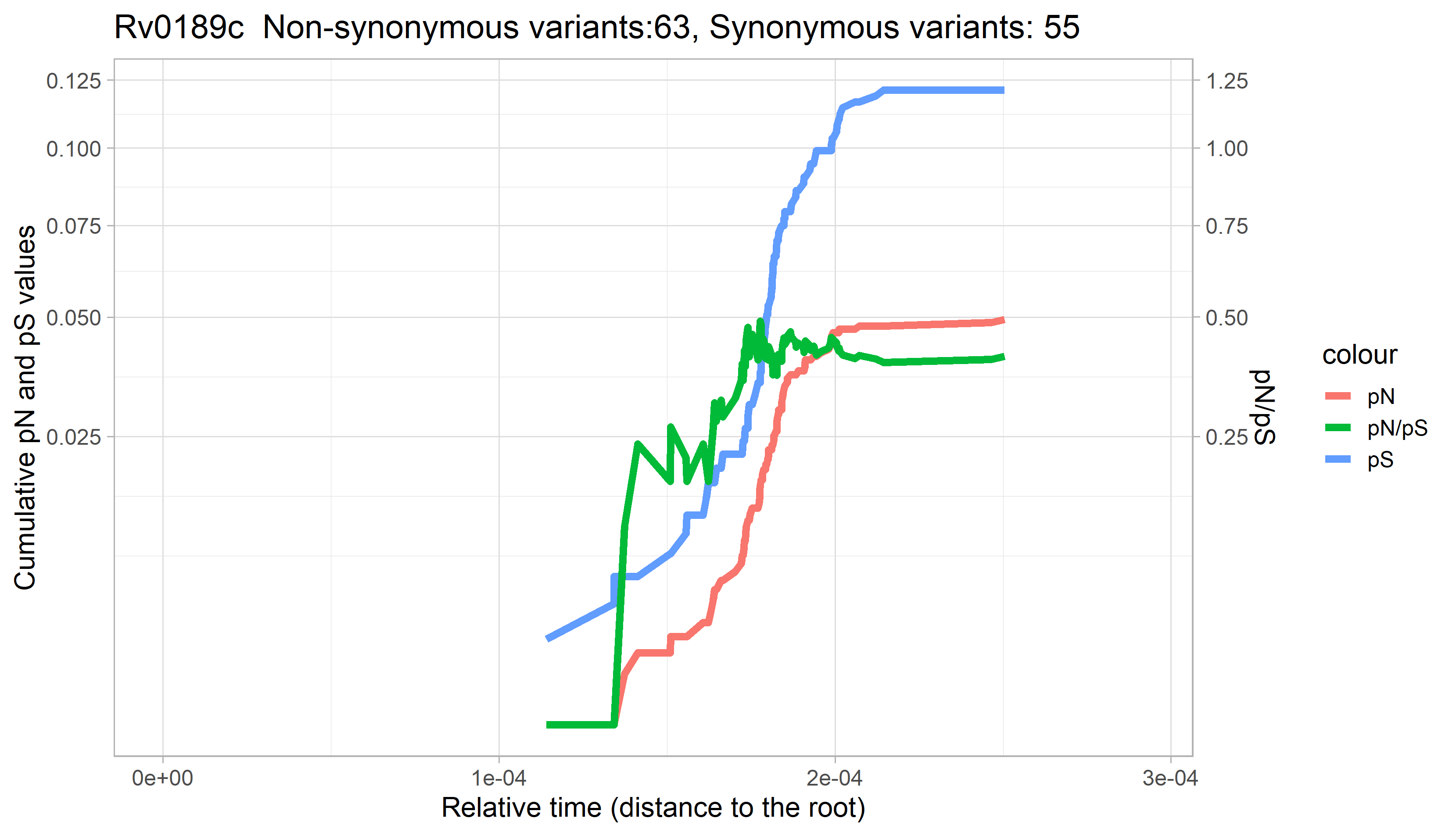

### Rv0190.png

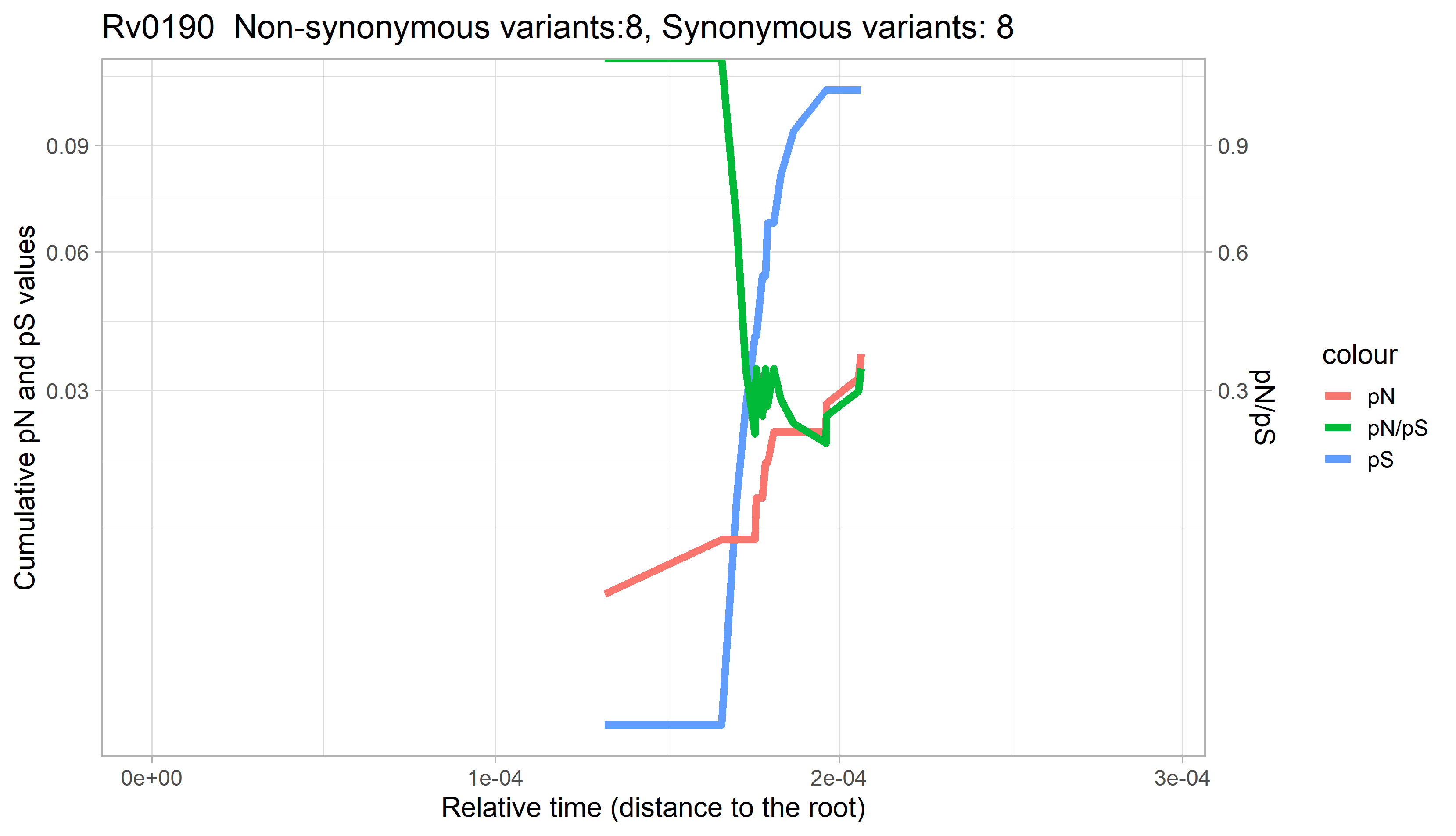

### Rv0191.png

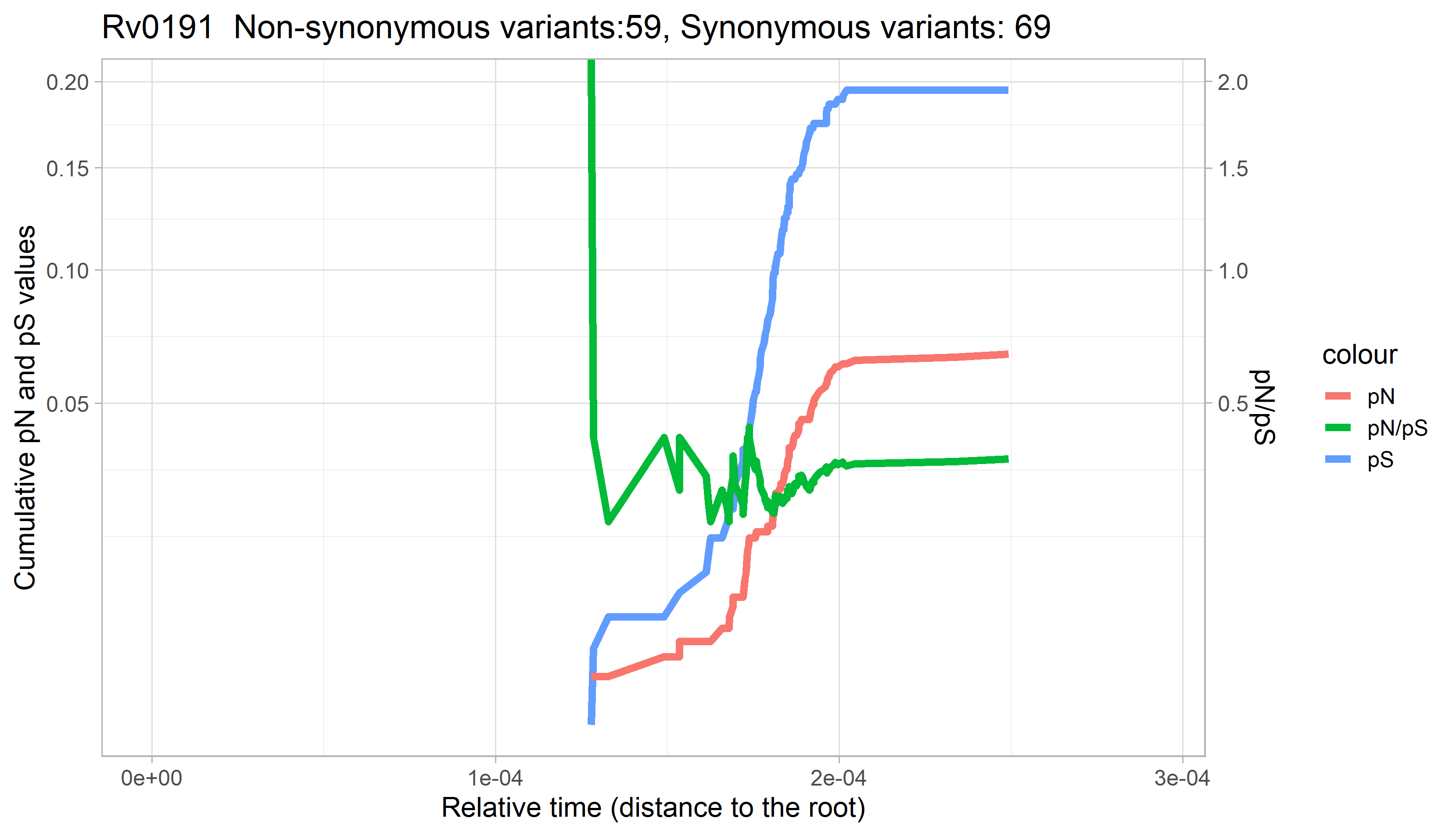

### Rv0192.png

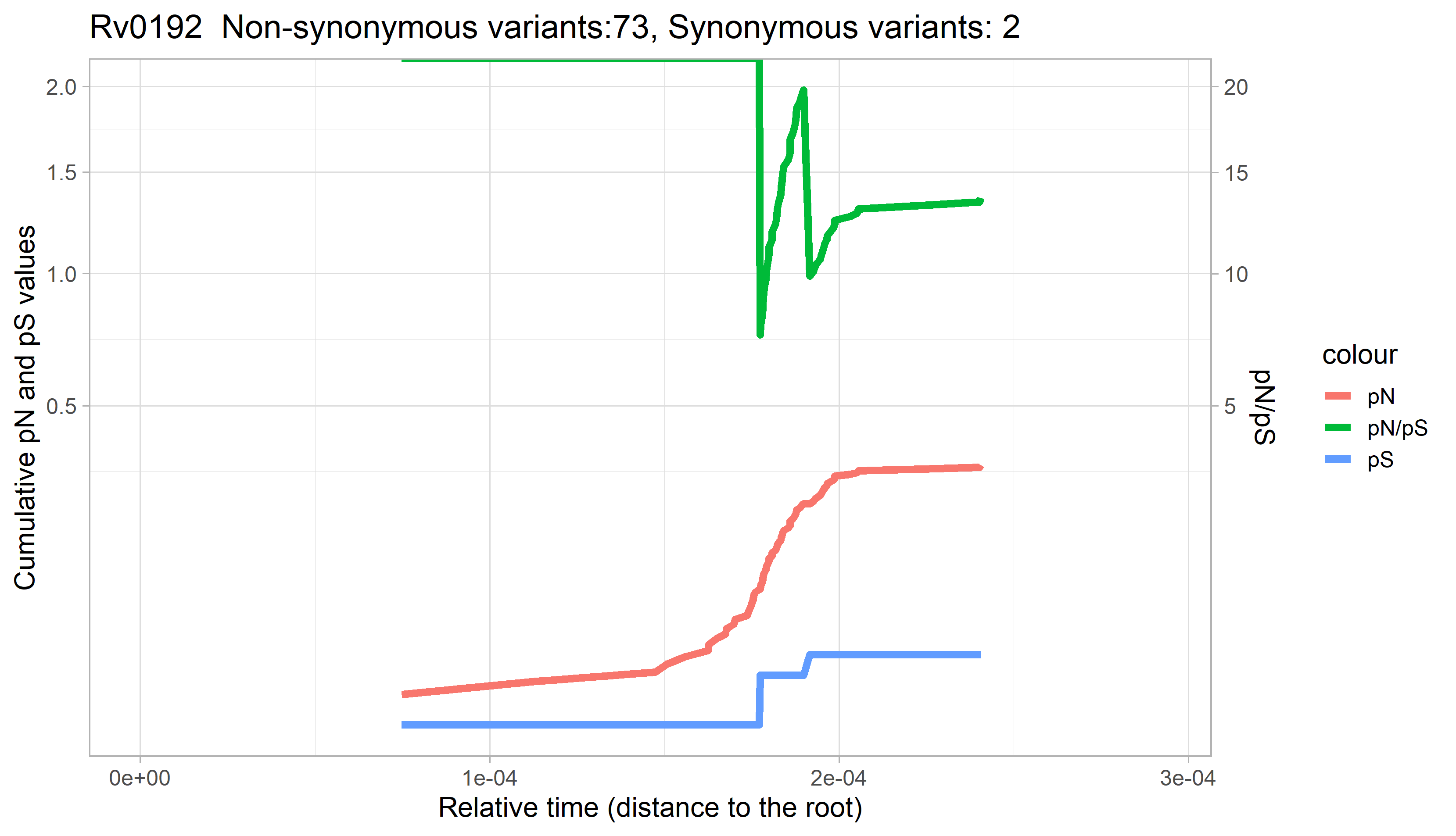

### Rv0192A.png

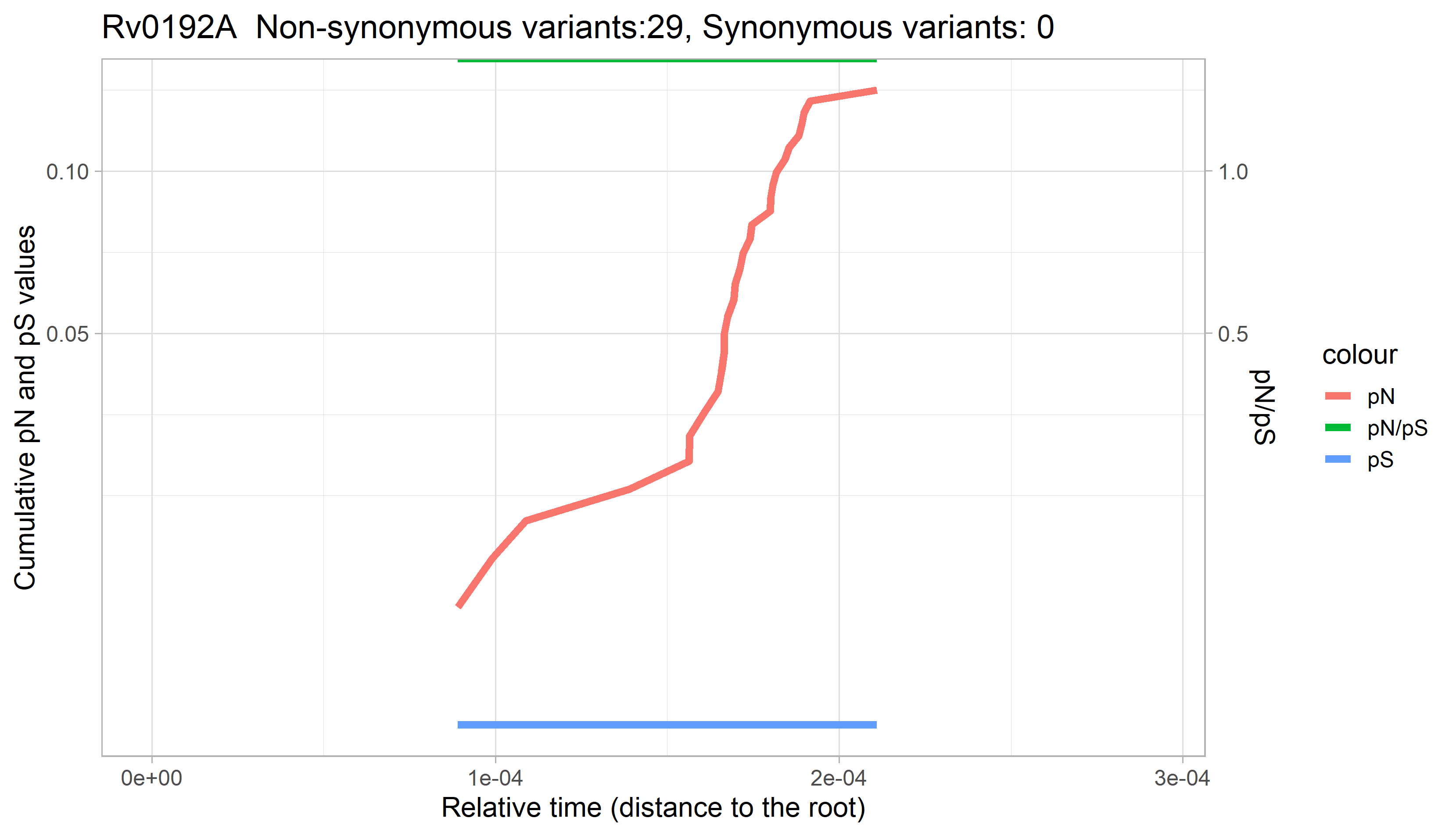

### Rv0193c.png

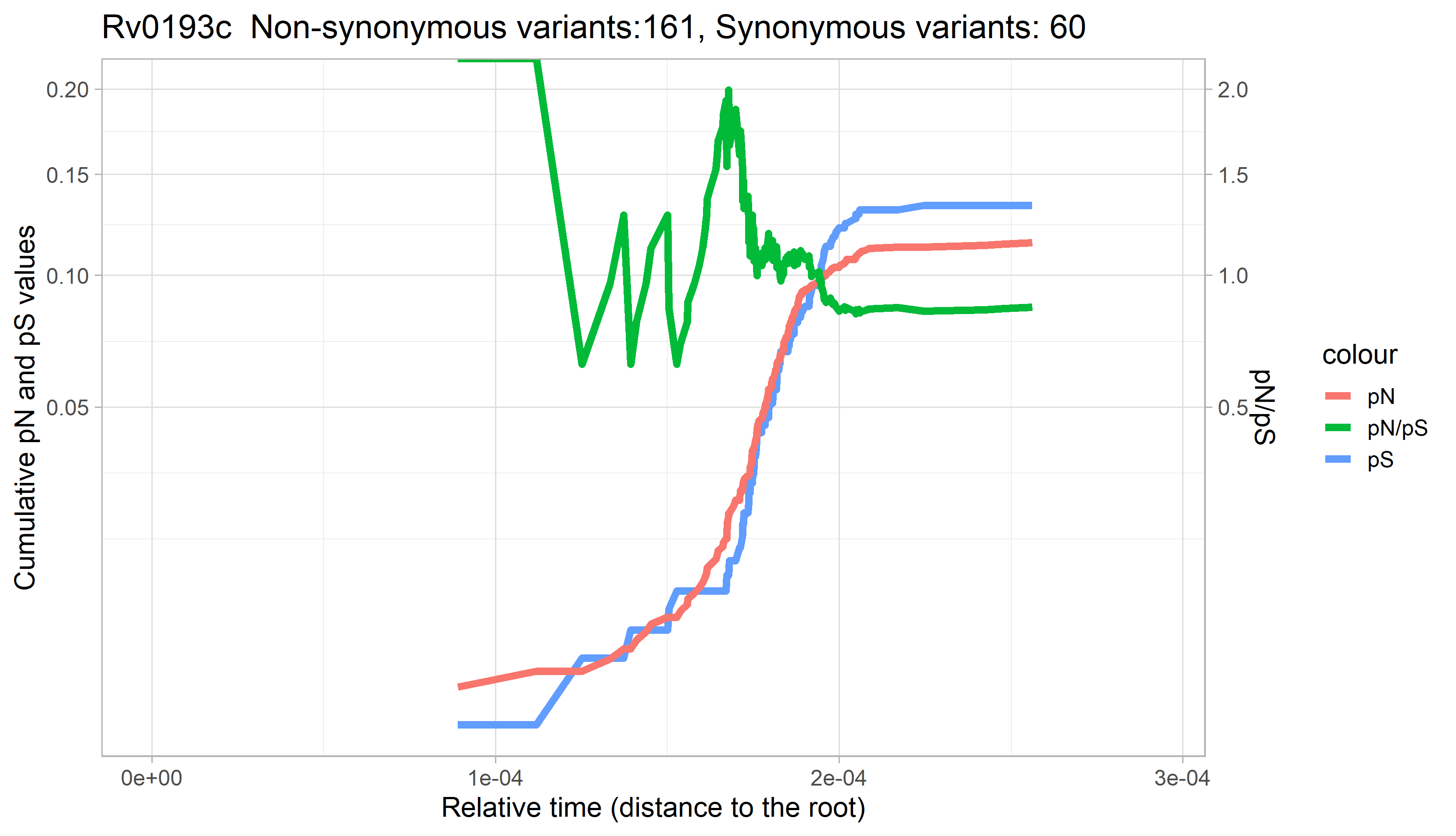

### Rv0194.png

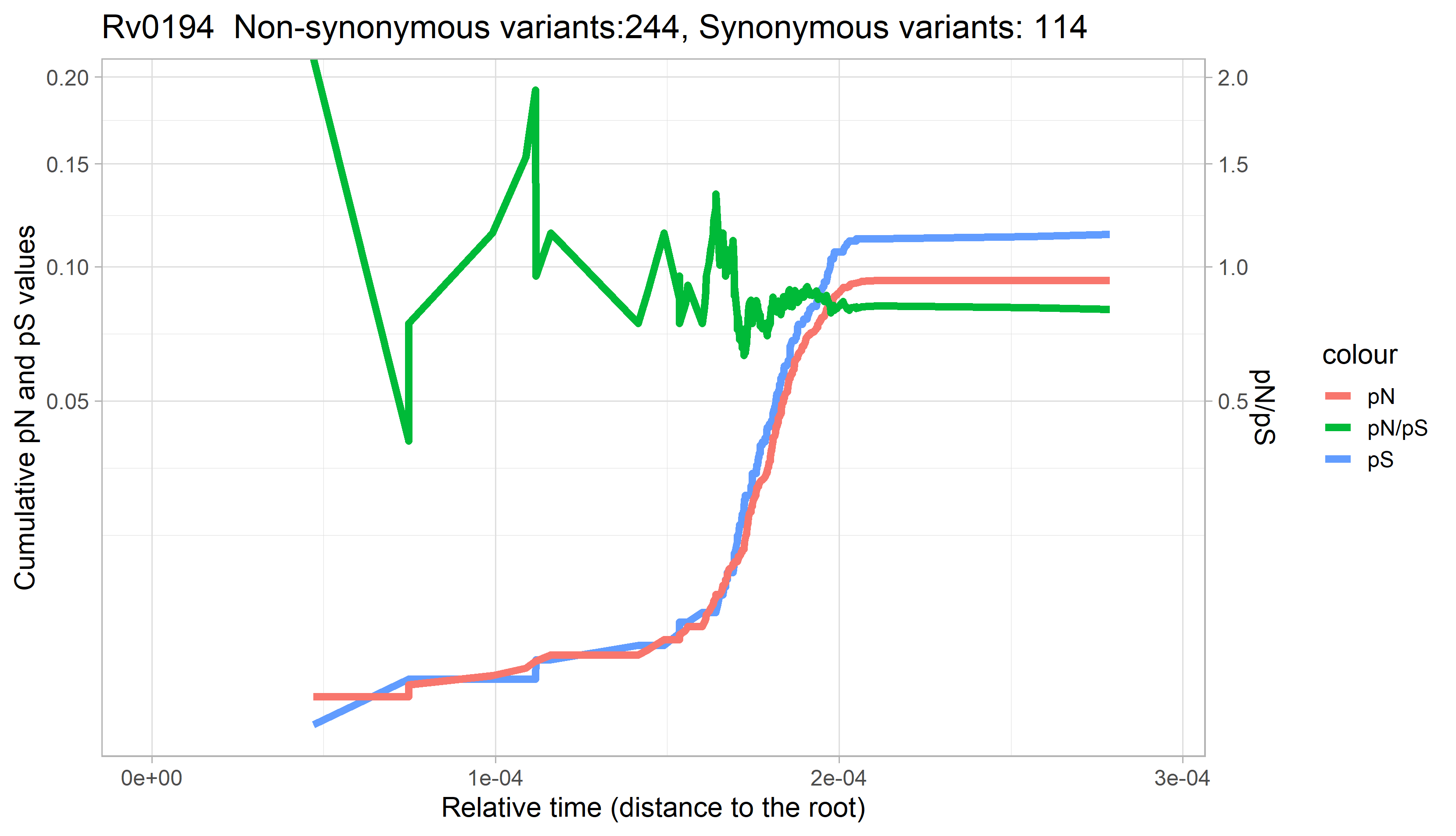

### Rv0195.png

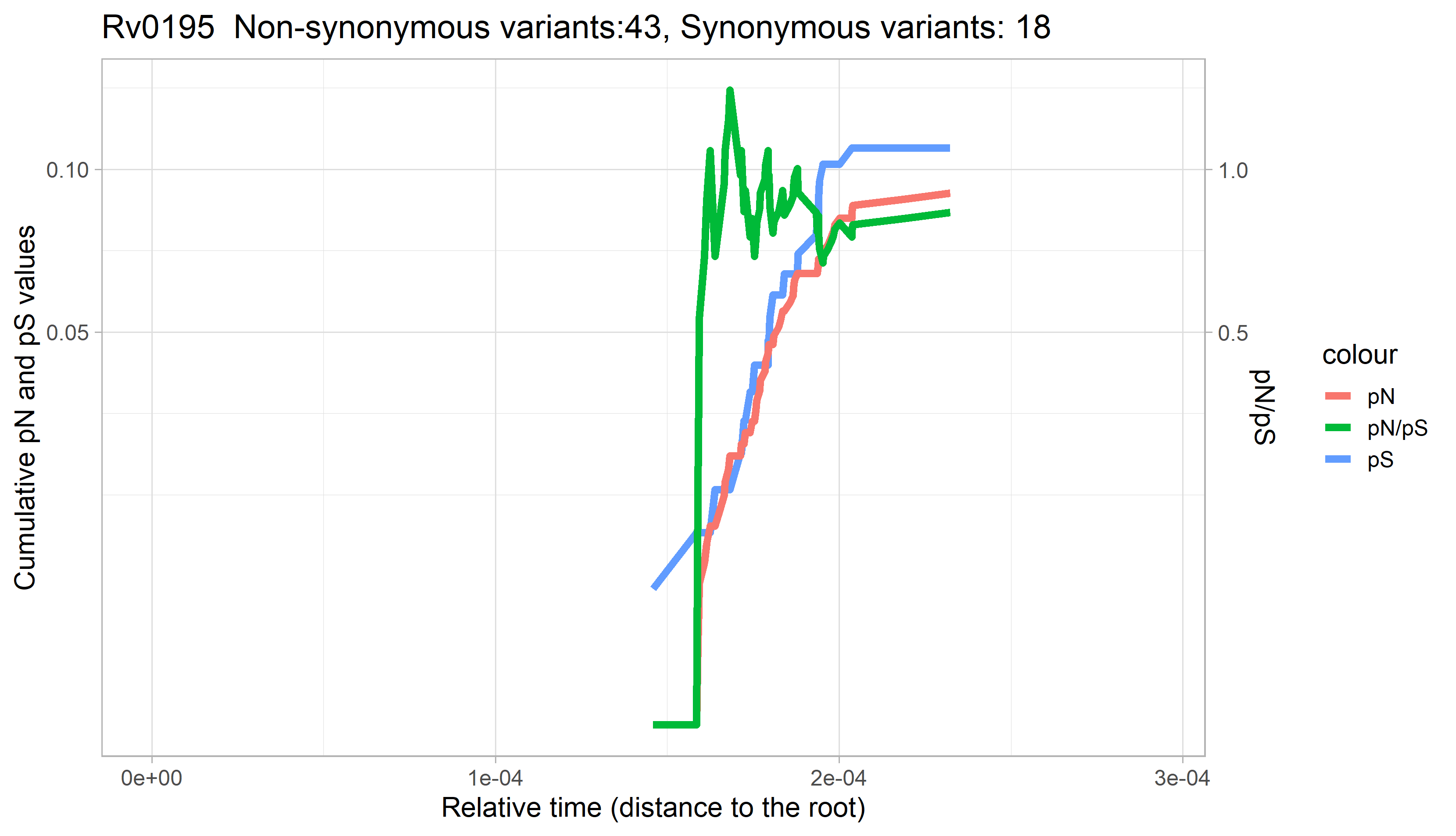

### Rv0196.png

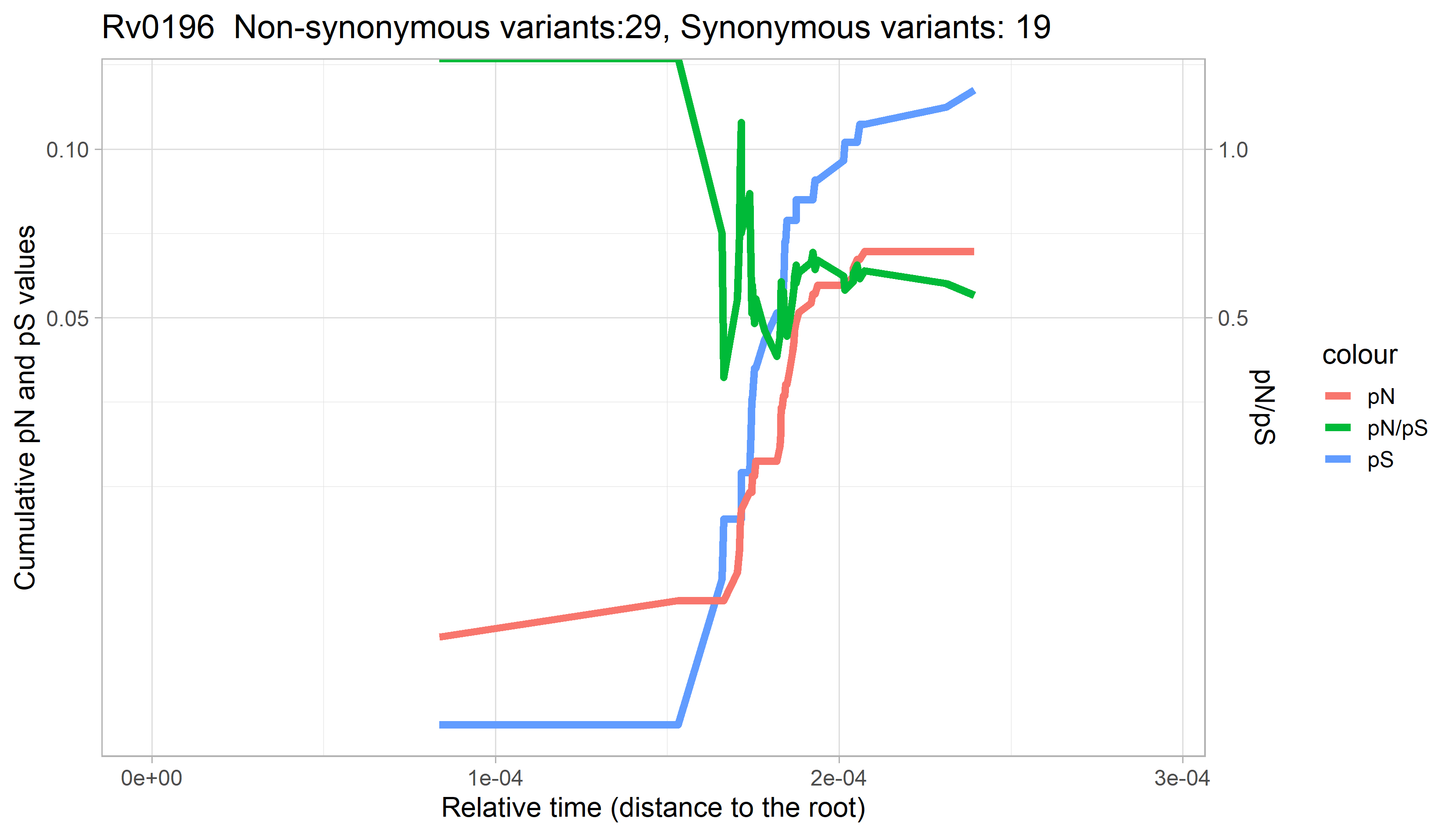

### Rv0197.png

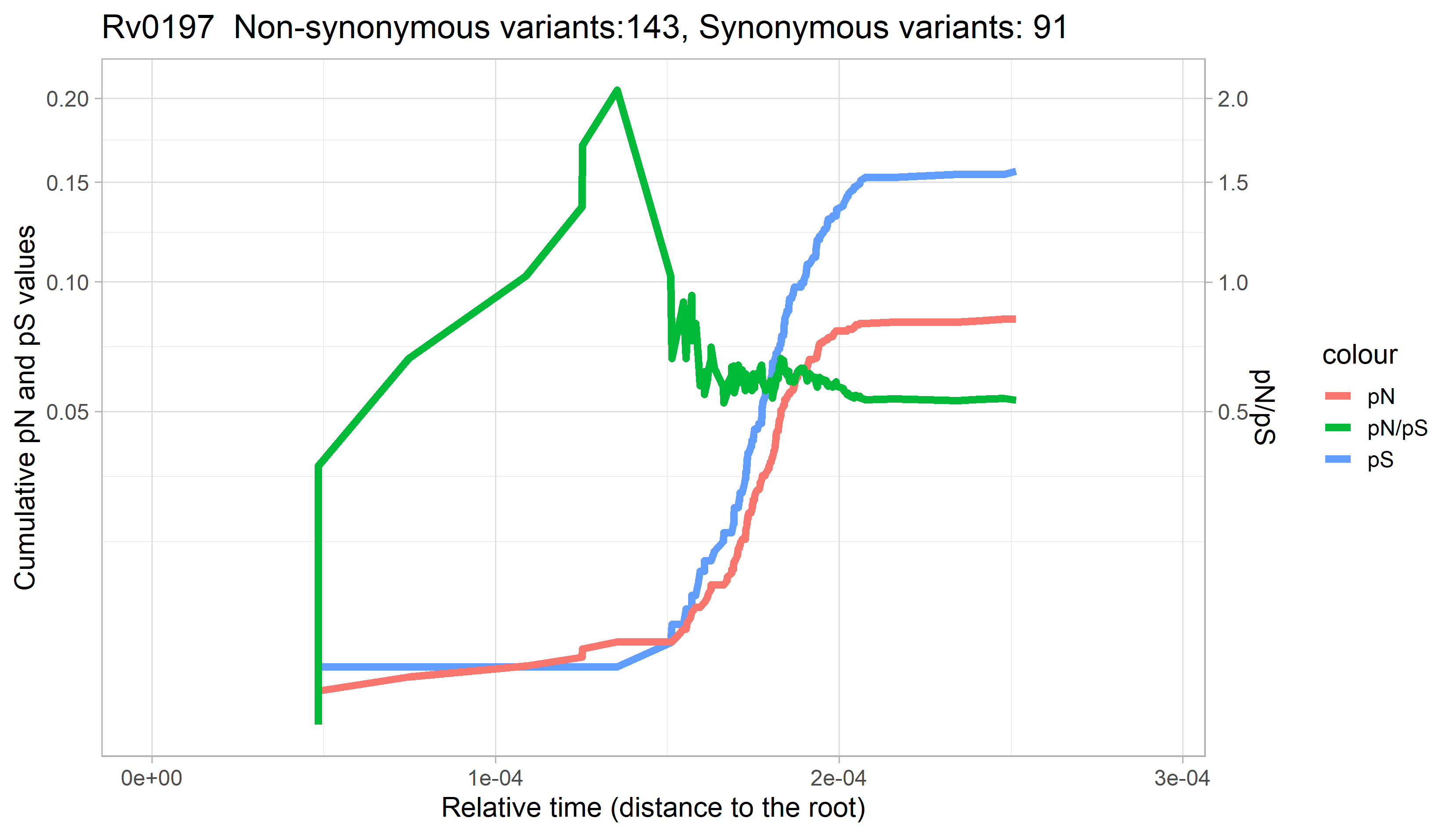

### Rv0198c.png

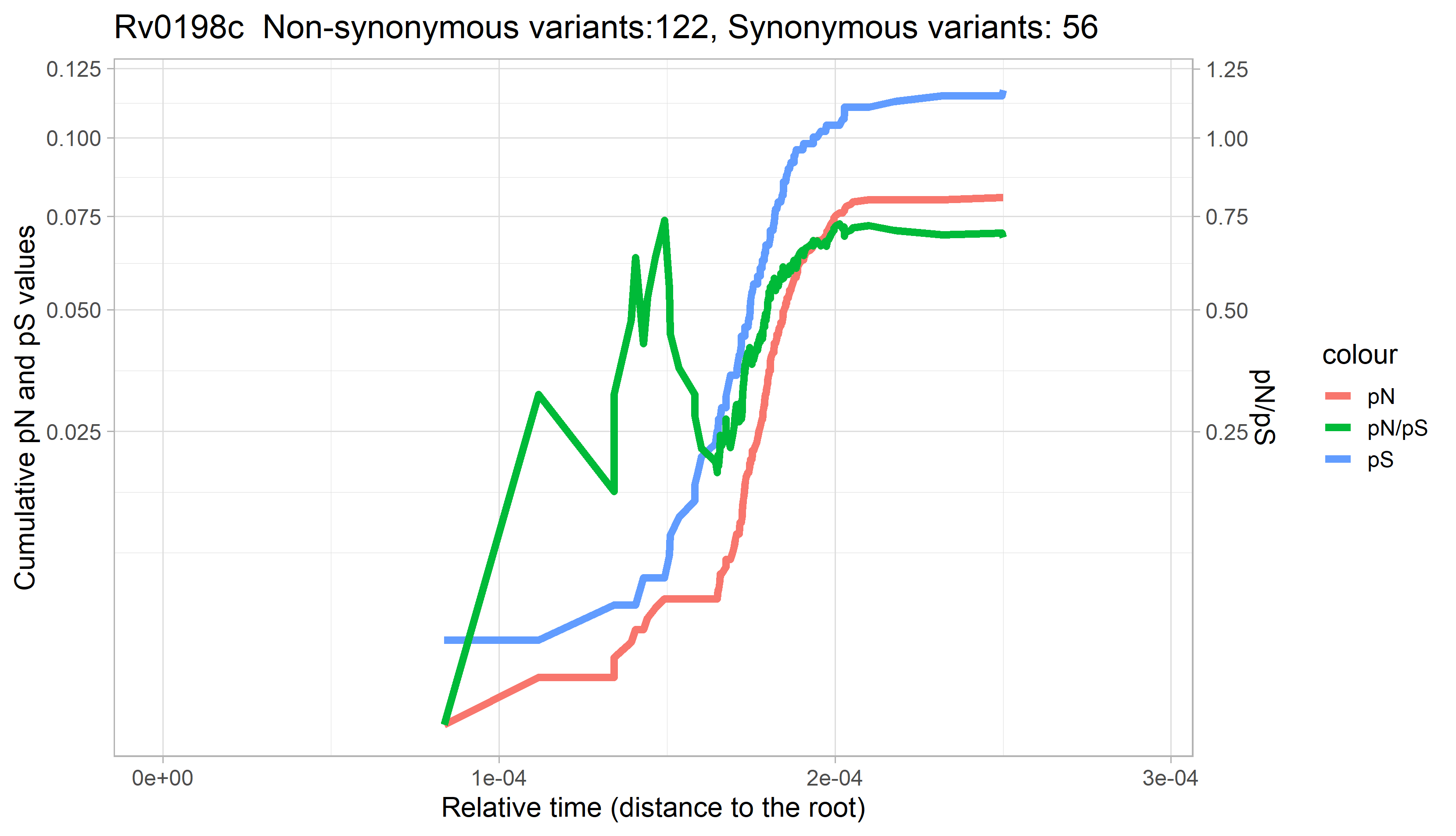

### Rv0199.png

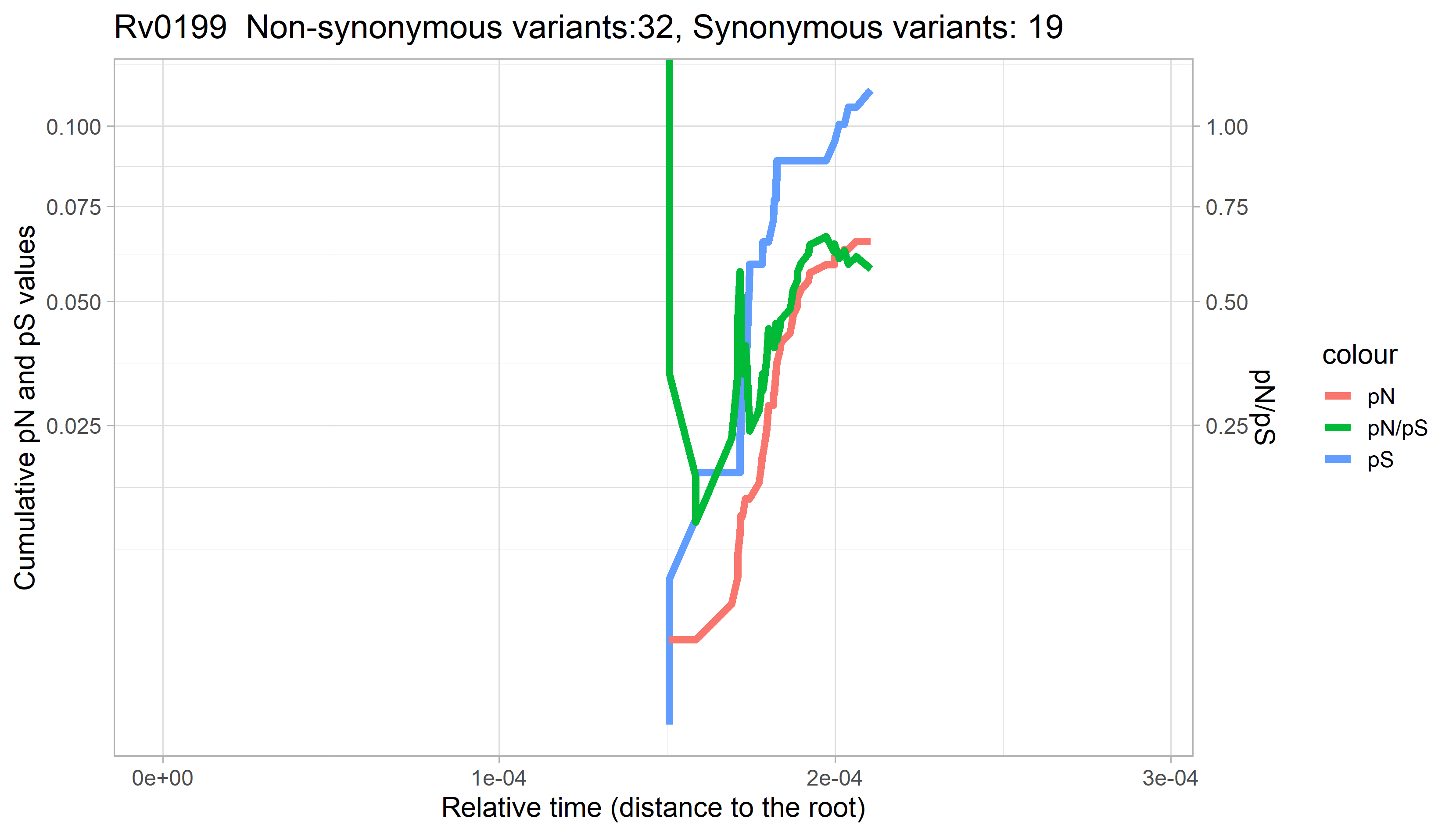

### Rv0200.png

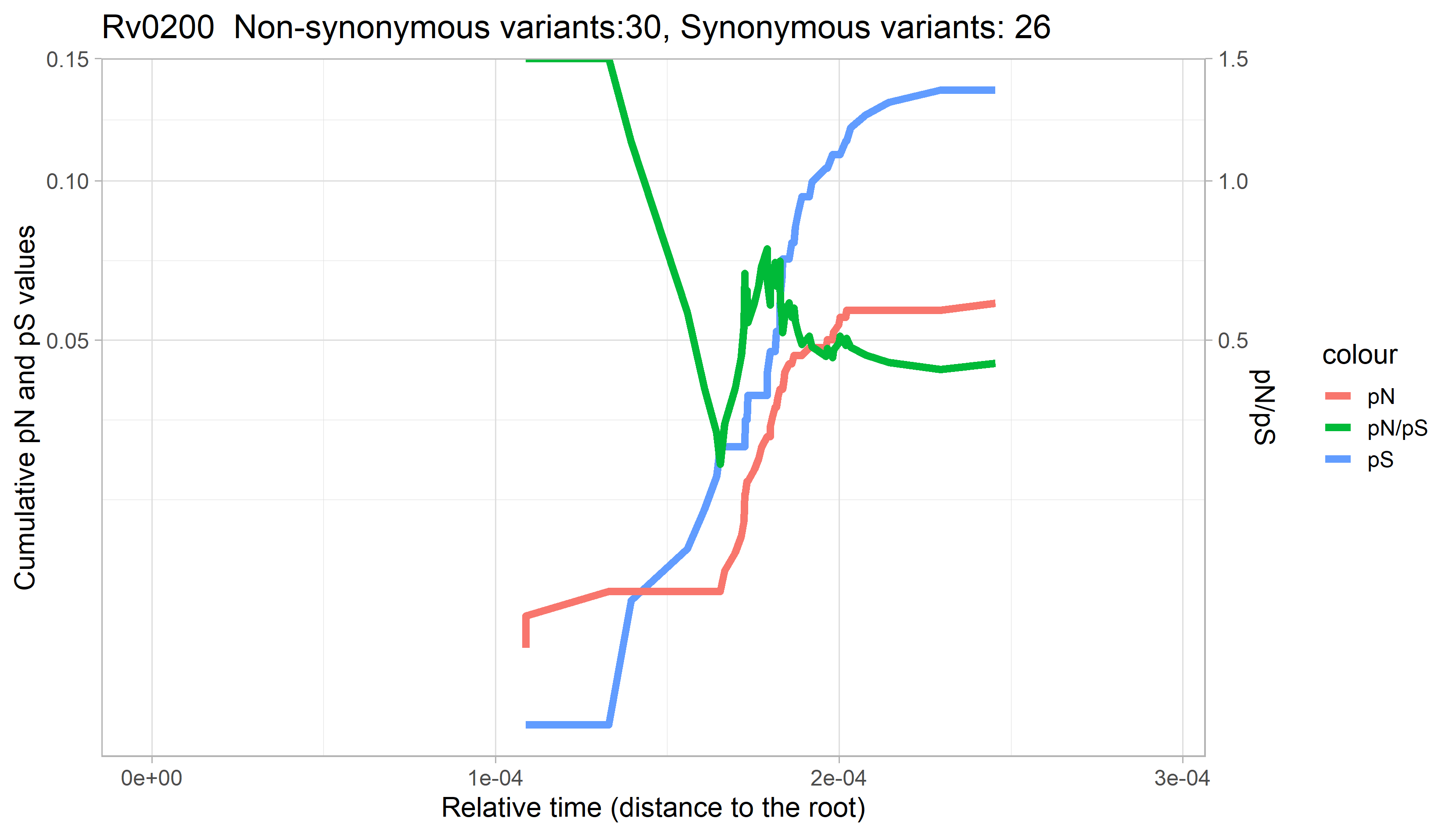

### Rv0201c.png

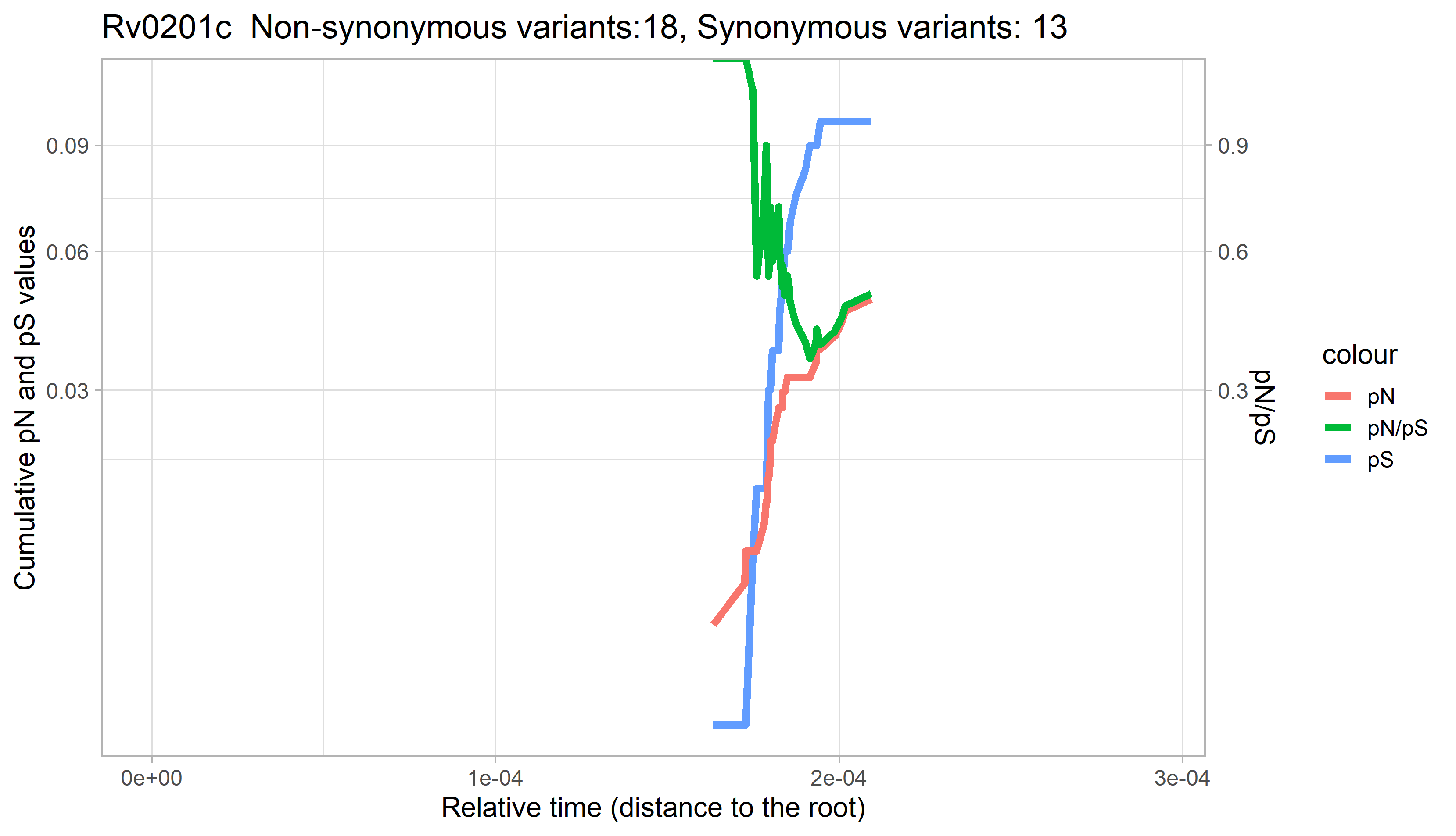

### Rv0202c.png

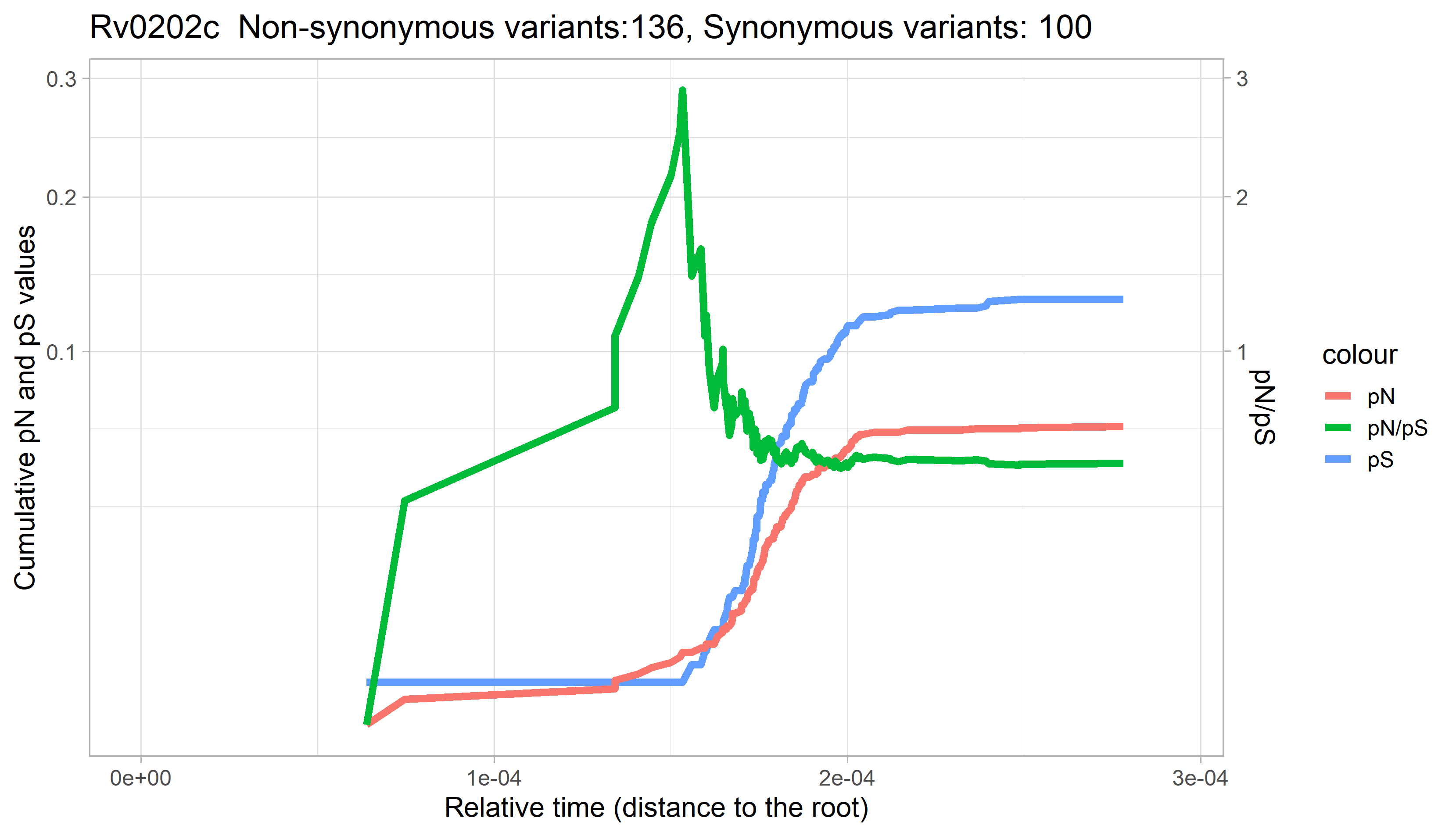

### Rv0203.png

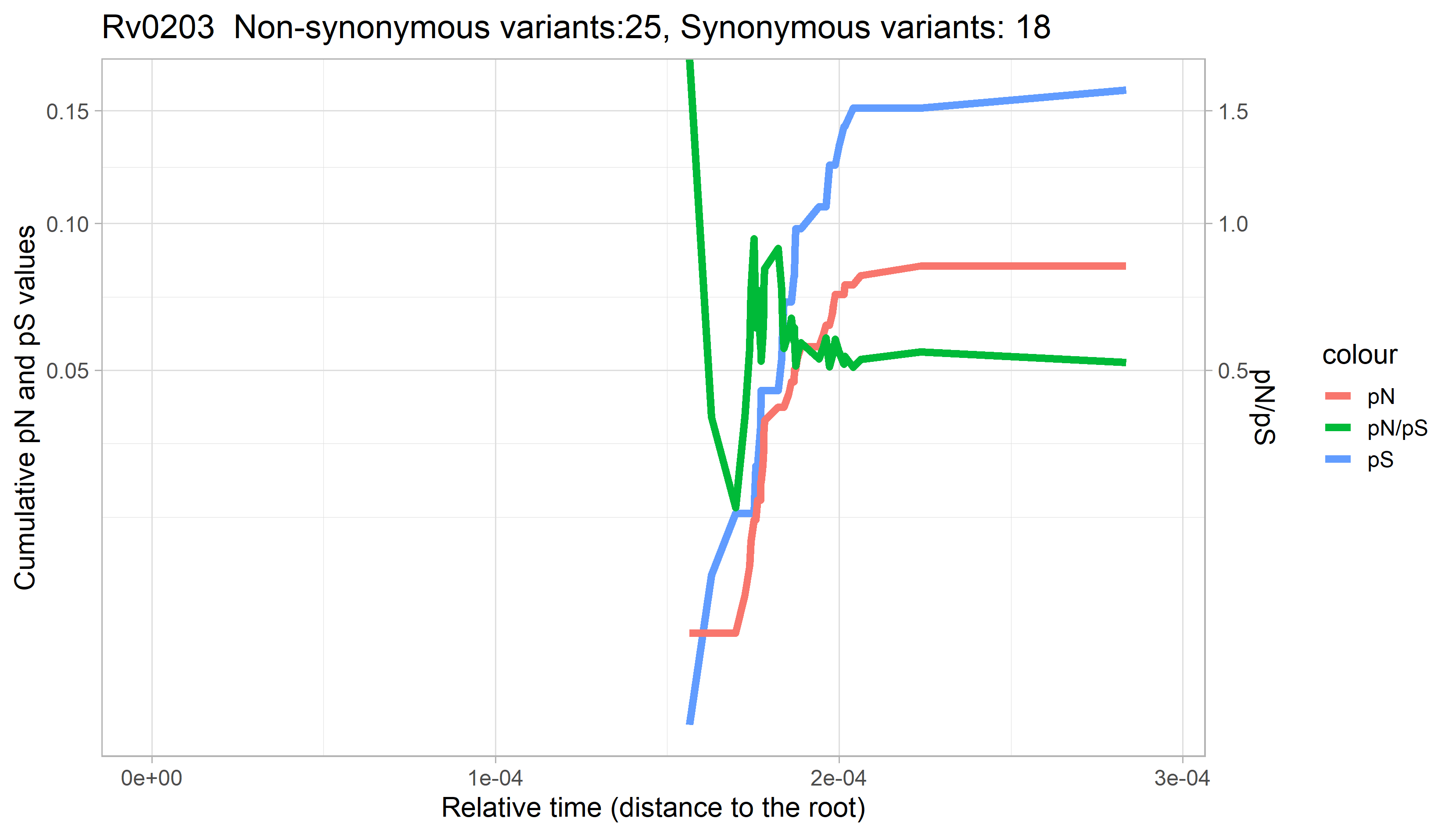

### Rv0204c.png

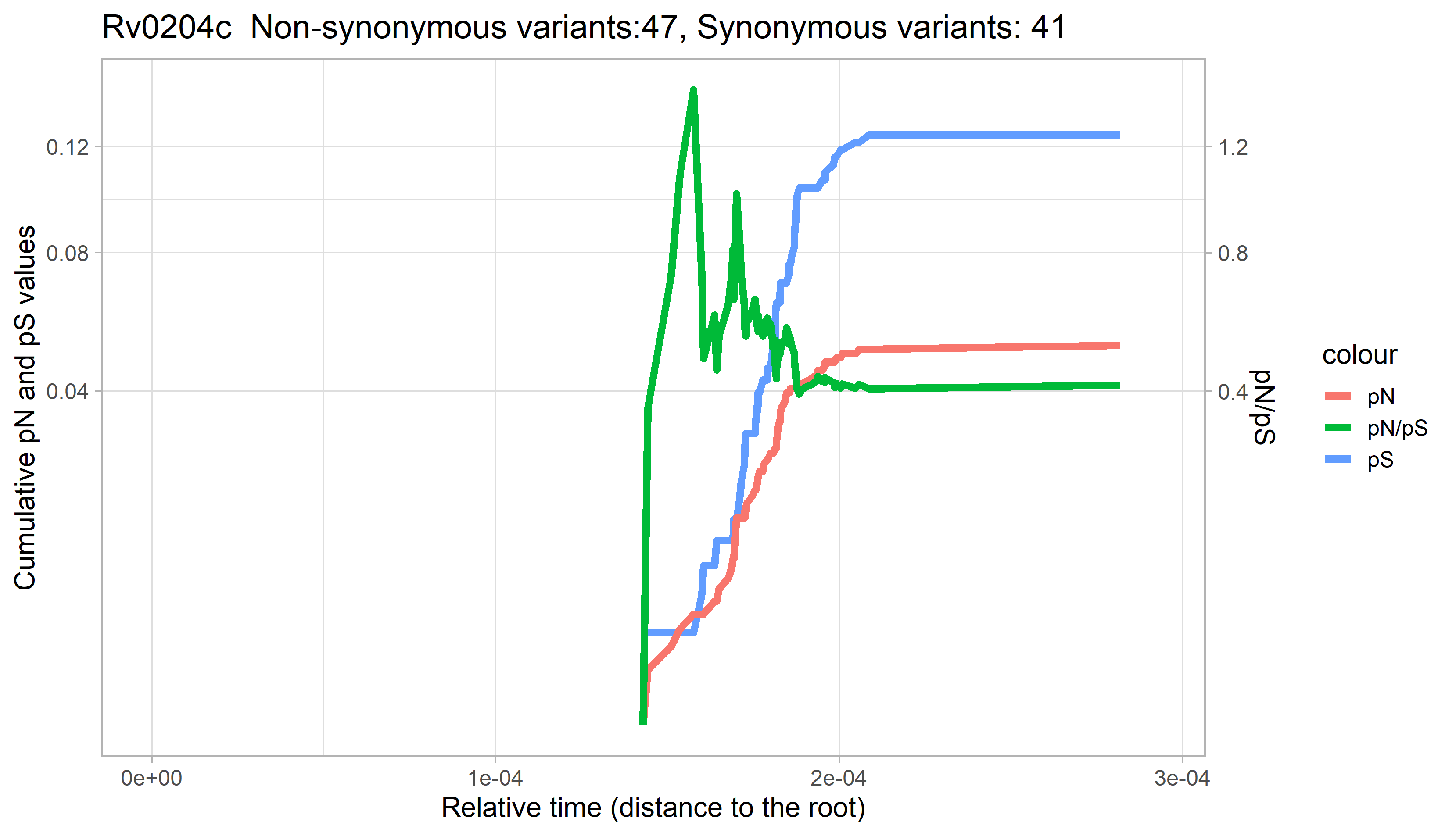

### Rv0205.png

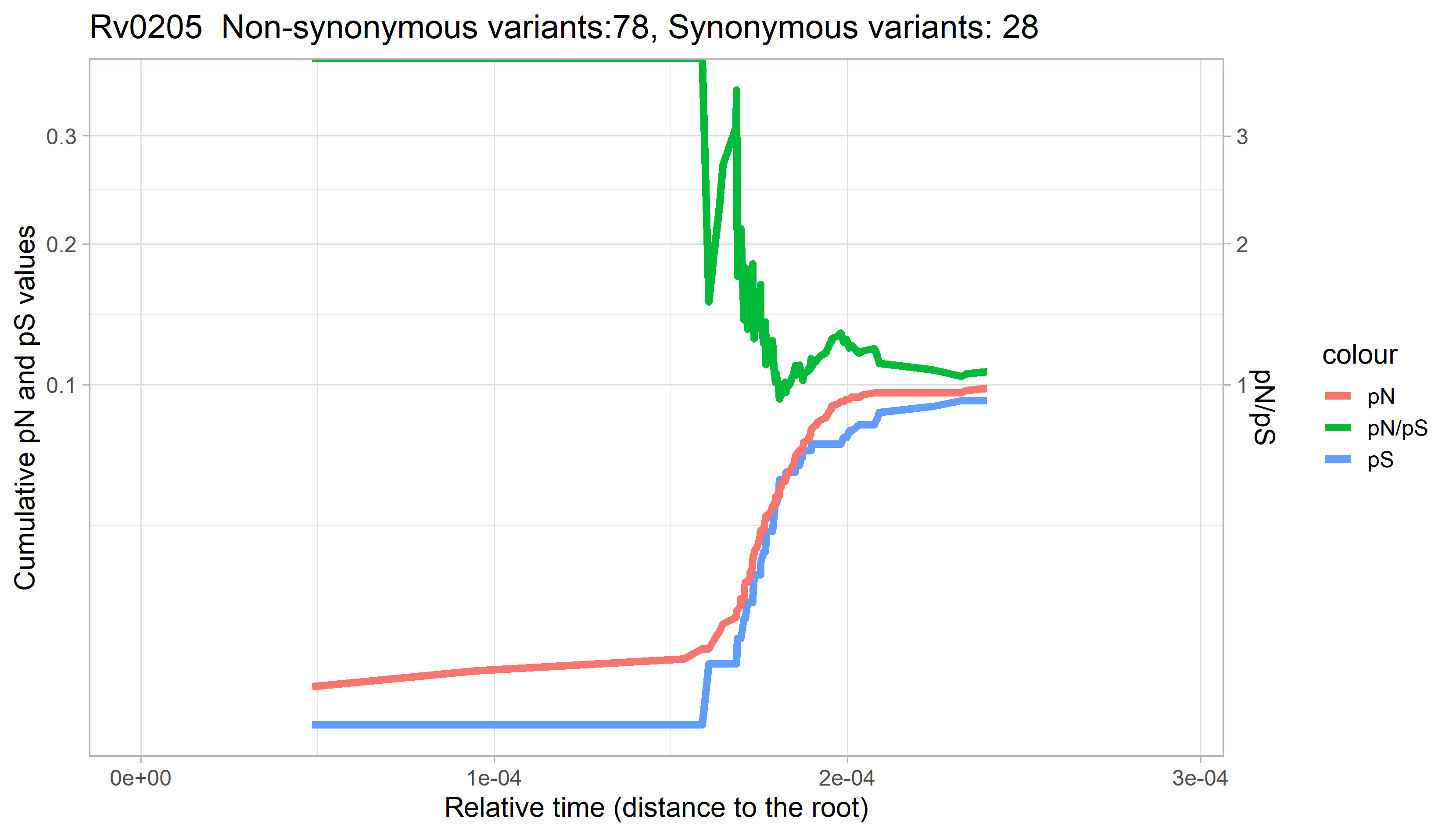

### Rv0206c.png

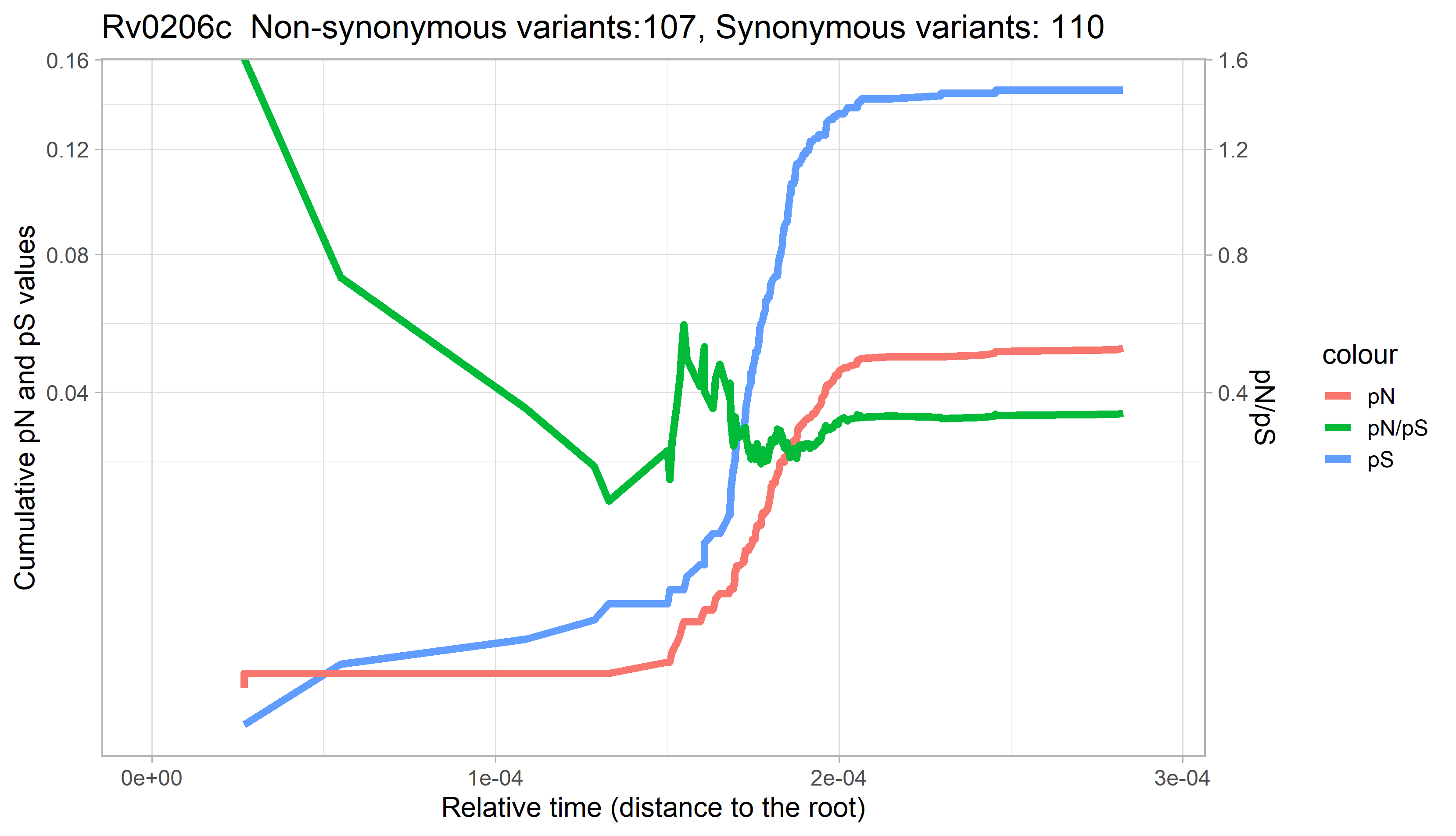

### Rv0207c.png

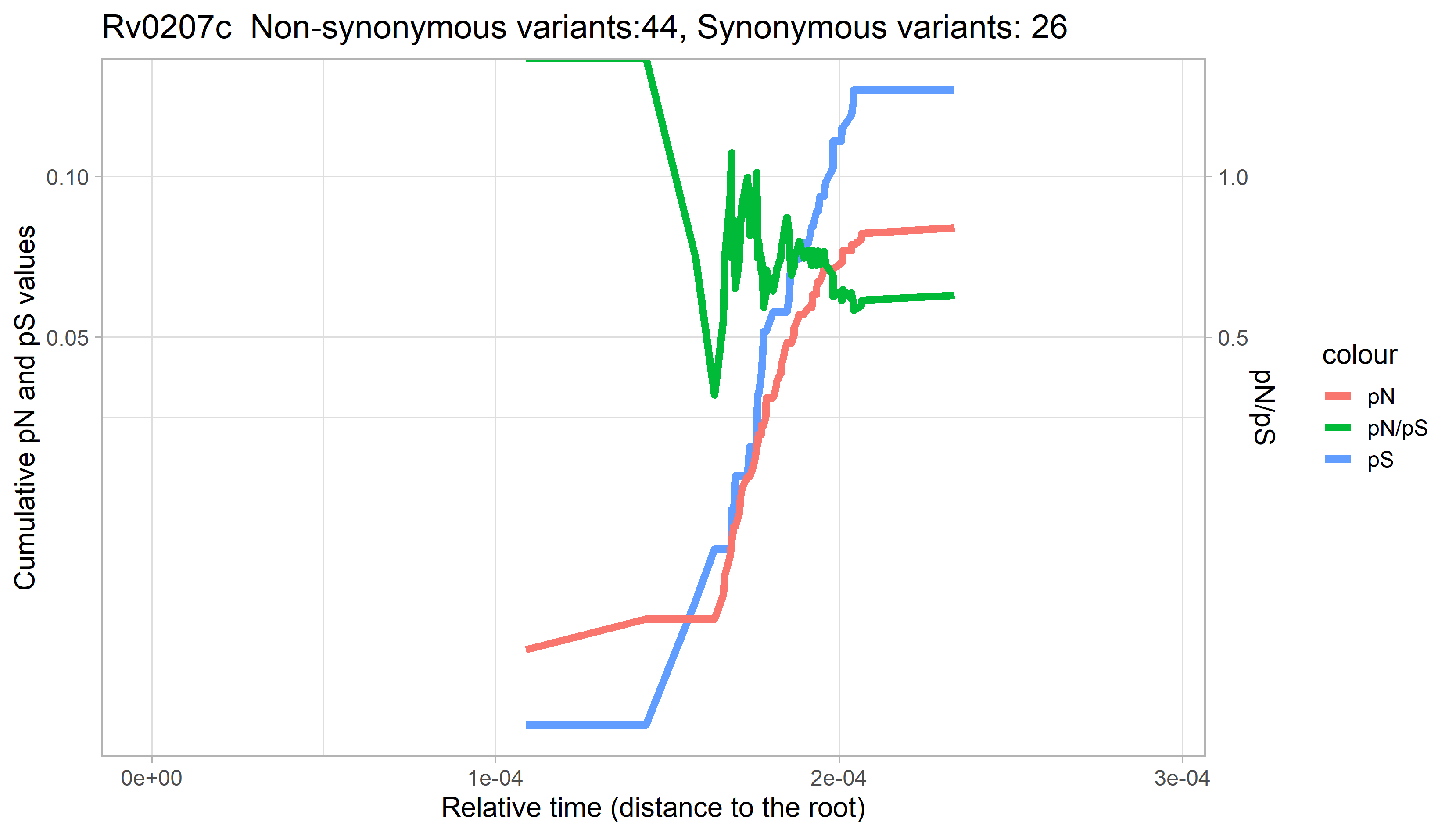

### Rv0208c.png

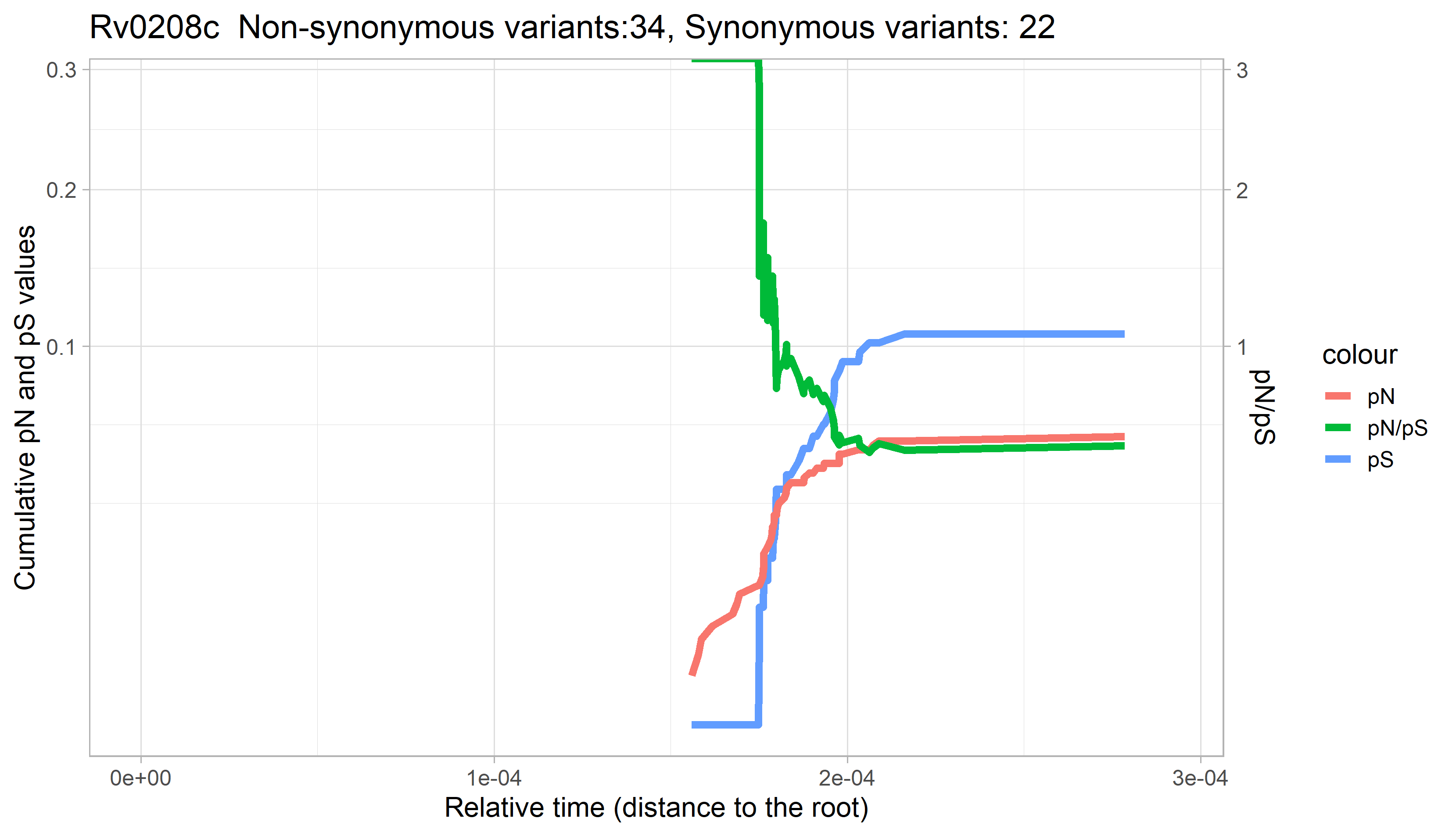
