## Supplementary figures and images for "Gene Evolutionary Trajectories in *M. tuberculosis* Reveal Temporal Signs of Selection"

### Rv0001.png

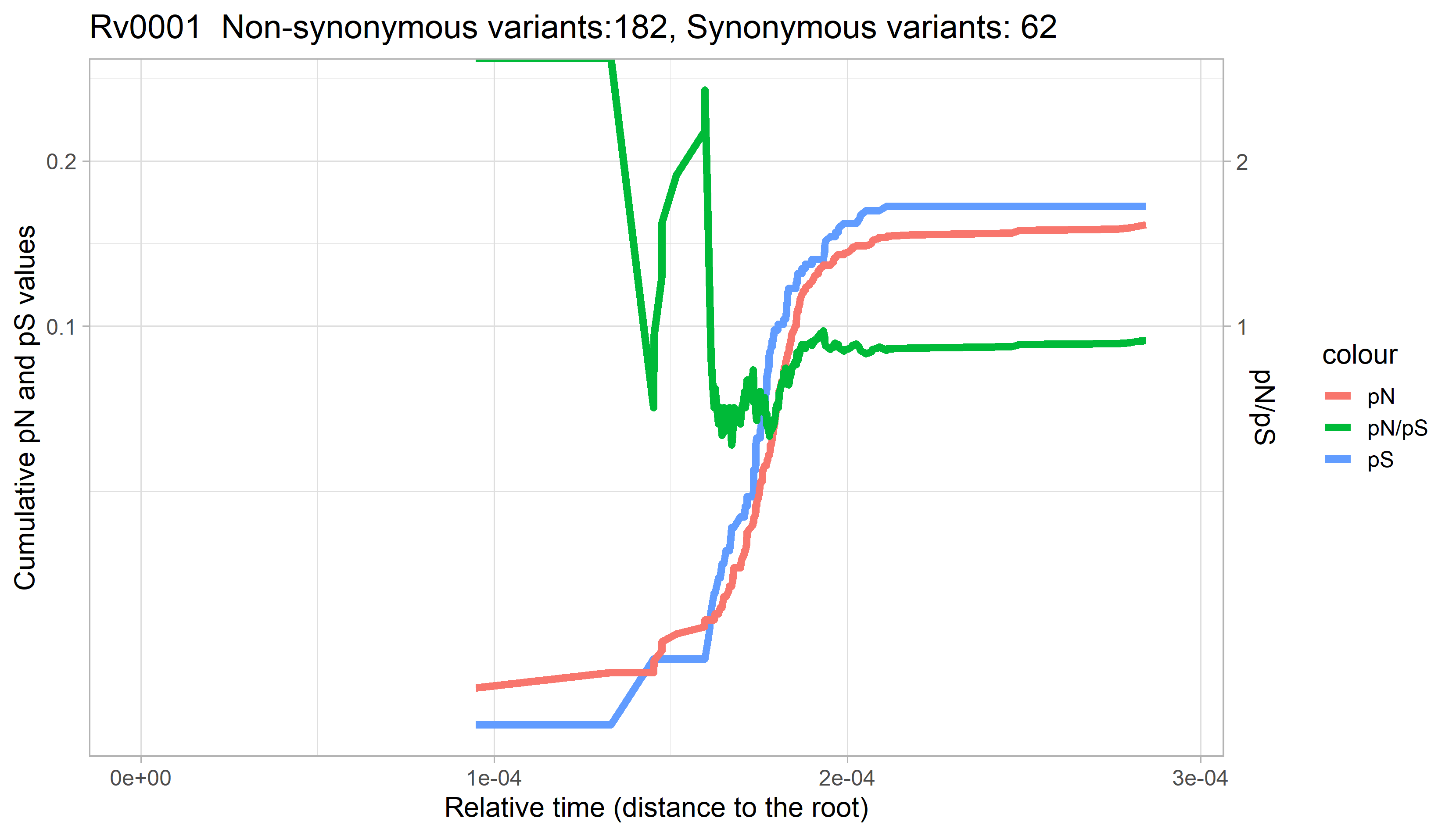

### Rv0002.png

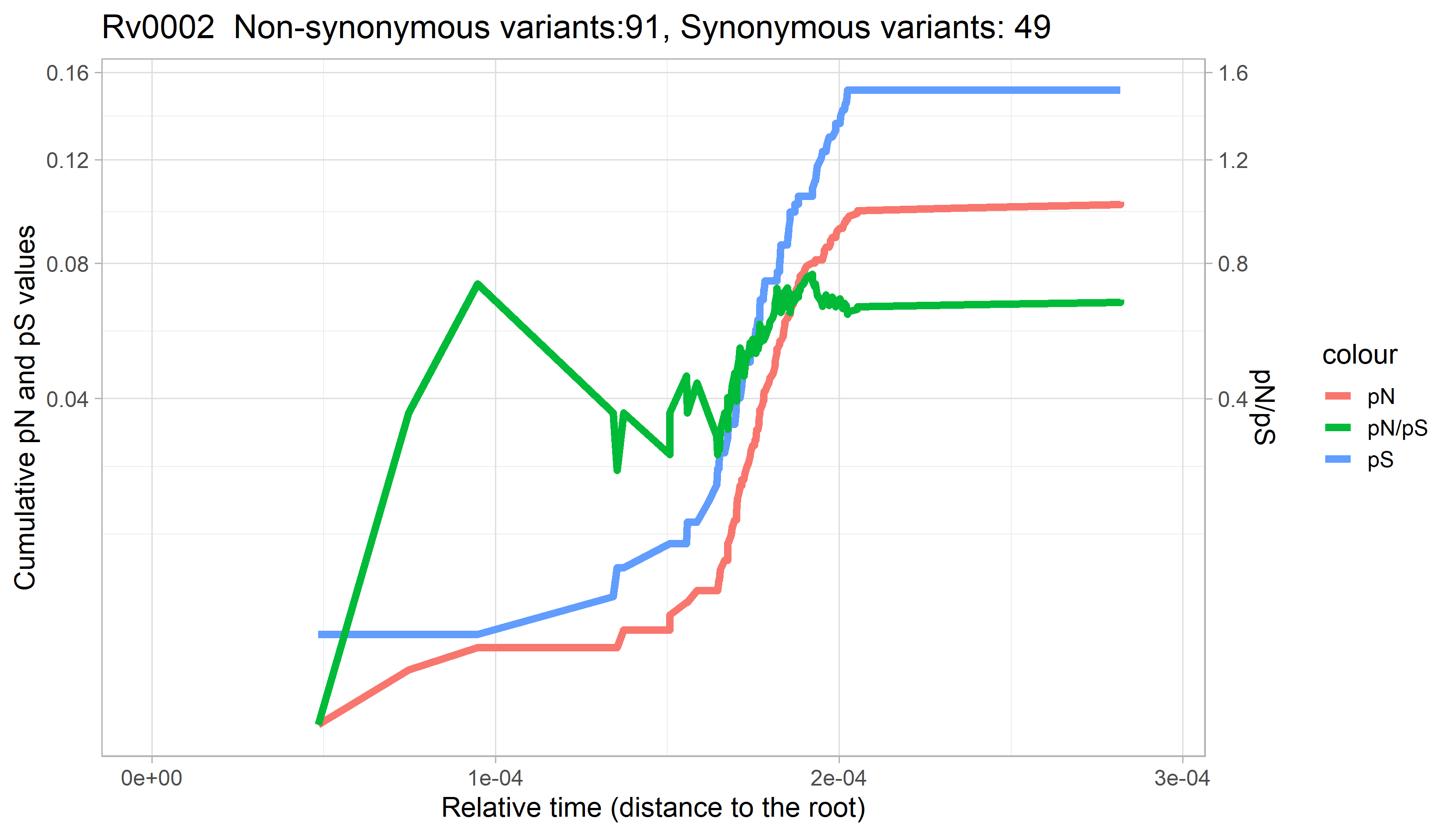

### Rv0003.png

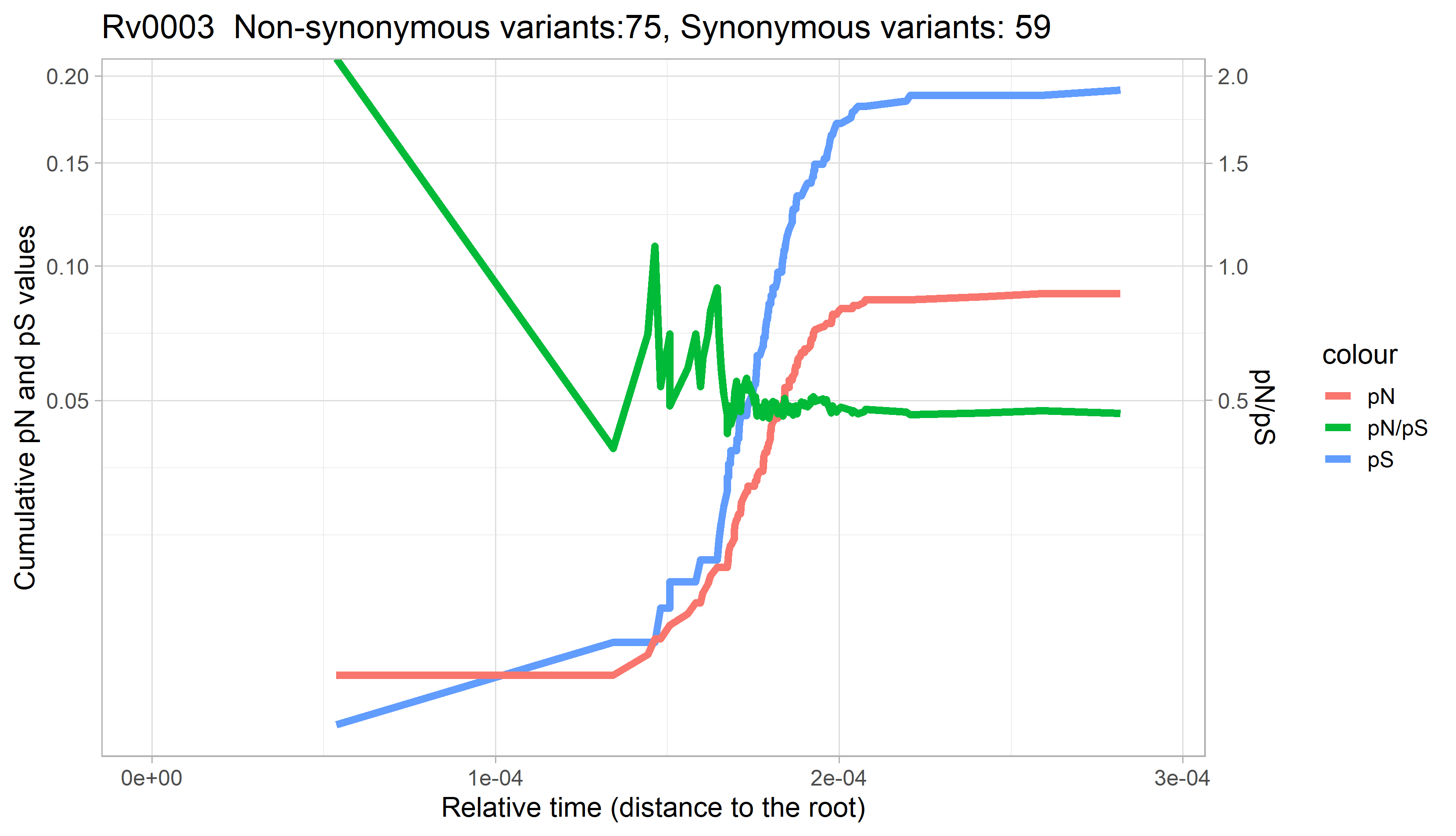

### Rv0004.png

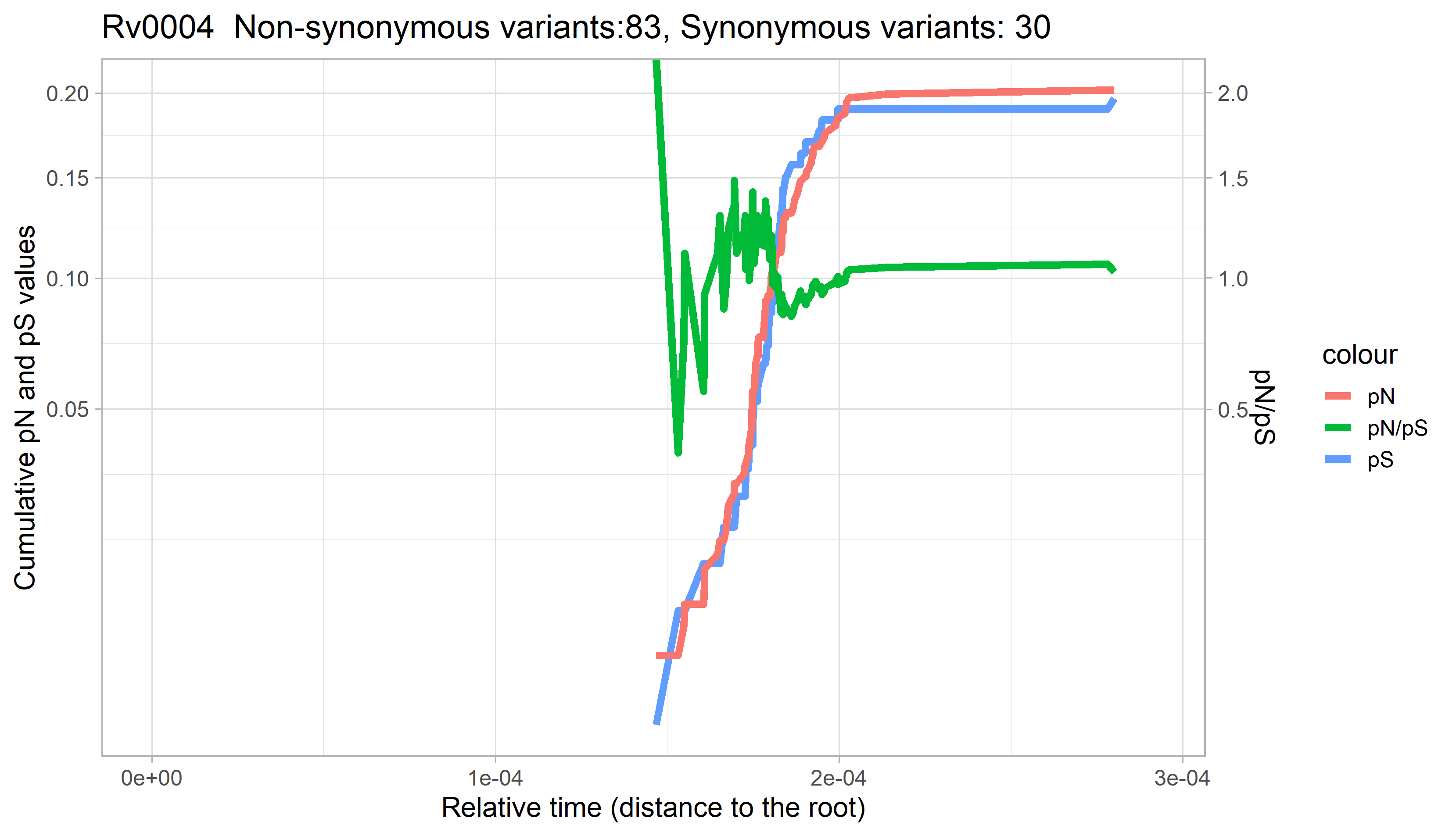

### Rv0005.png

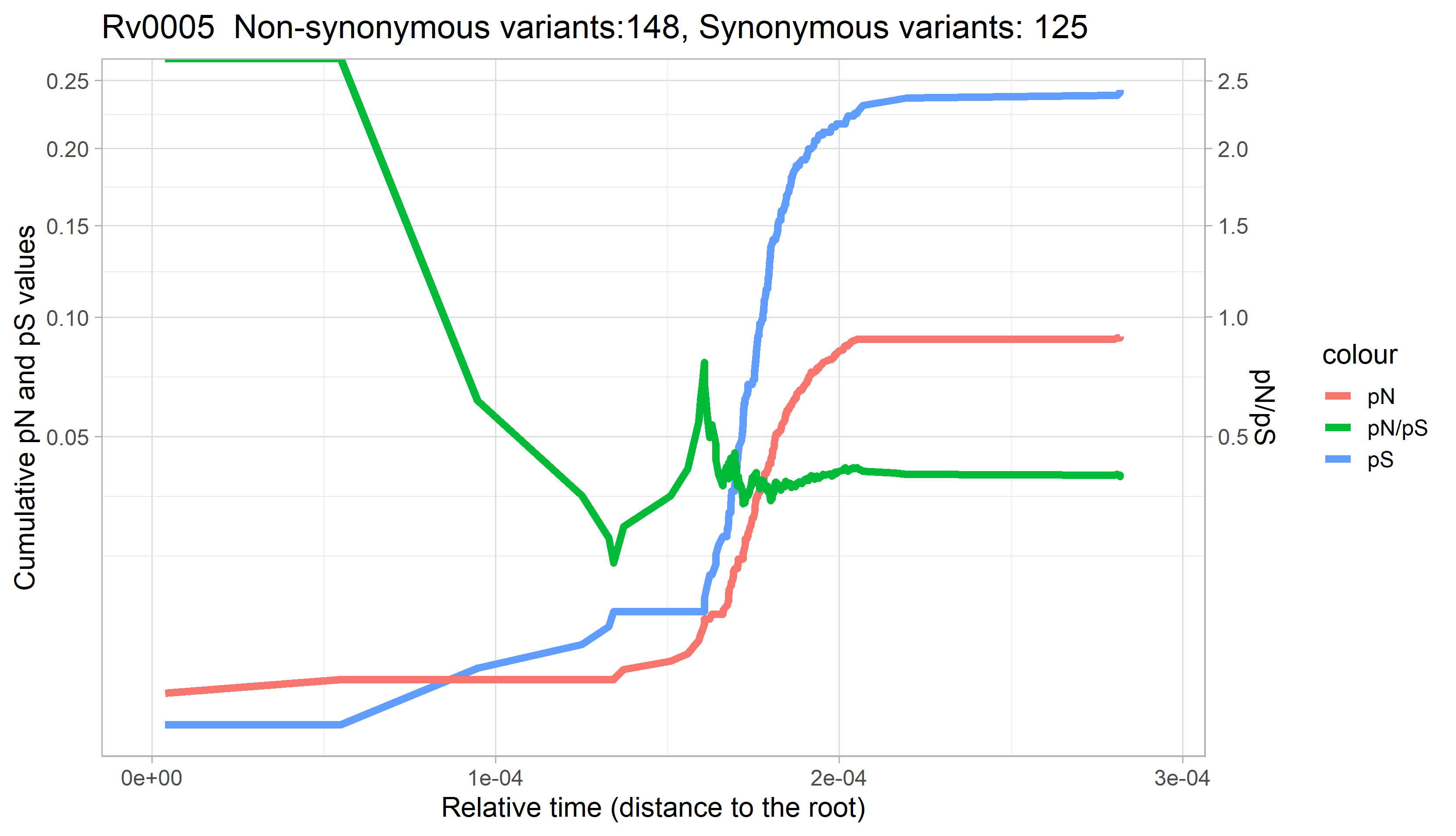

### Rv0006.png

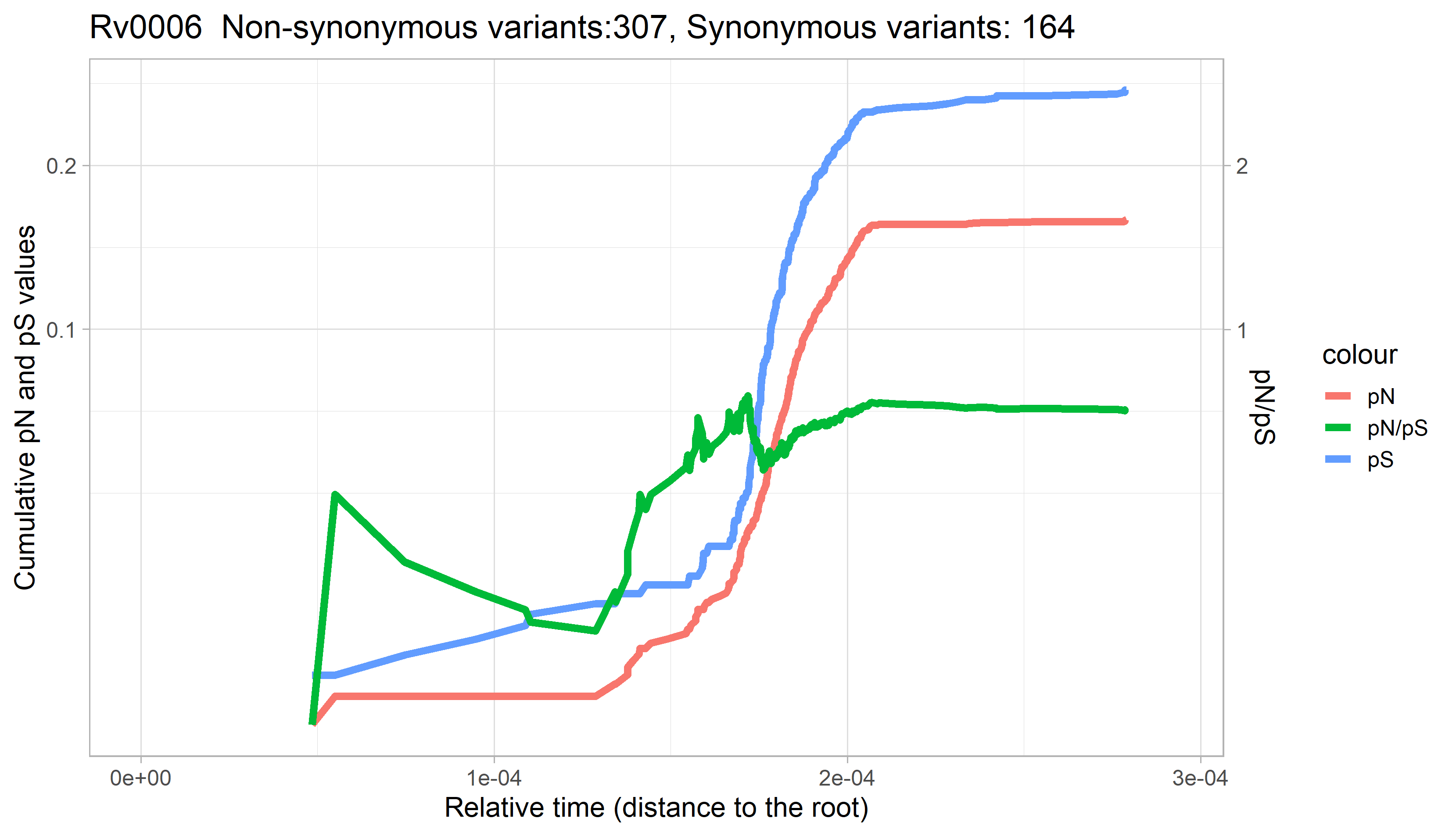

### Rv0007.png

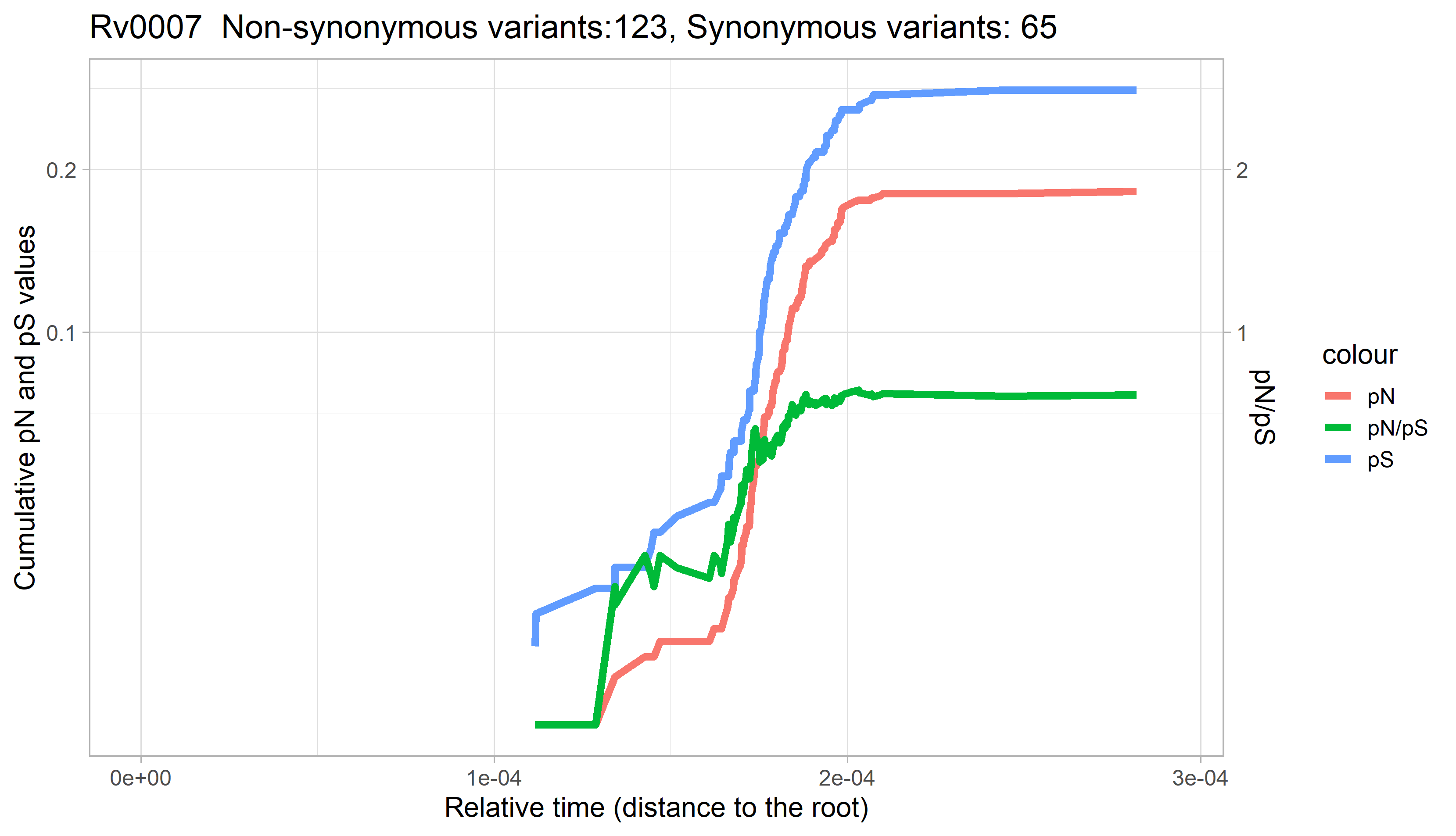

### Rv0008c.png

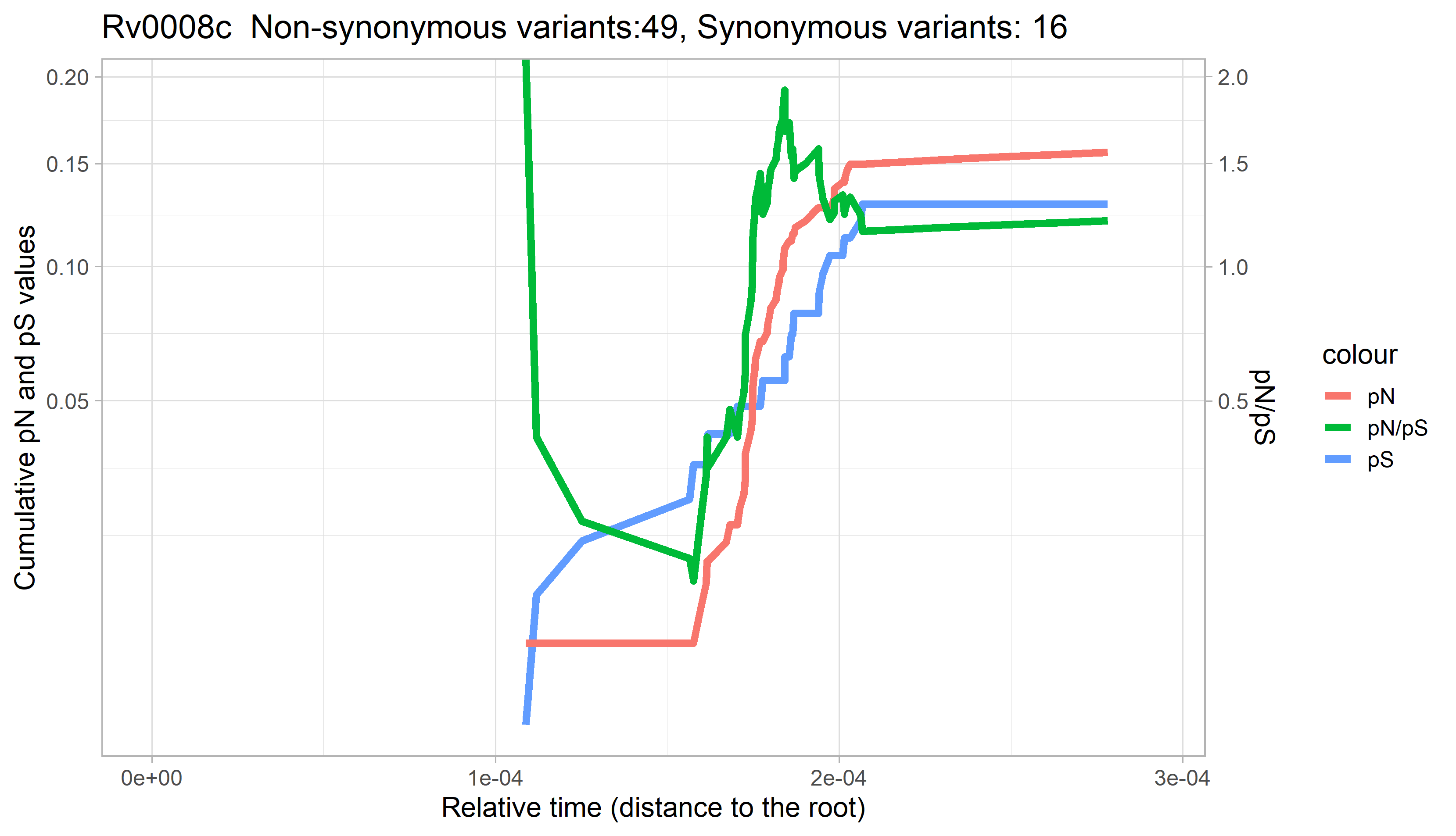

### Rv0009.png

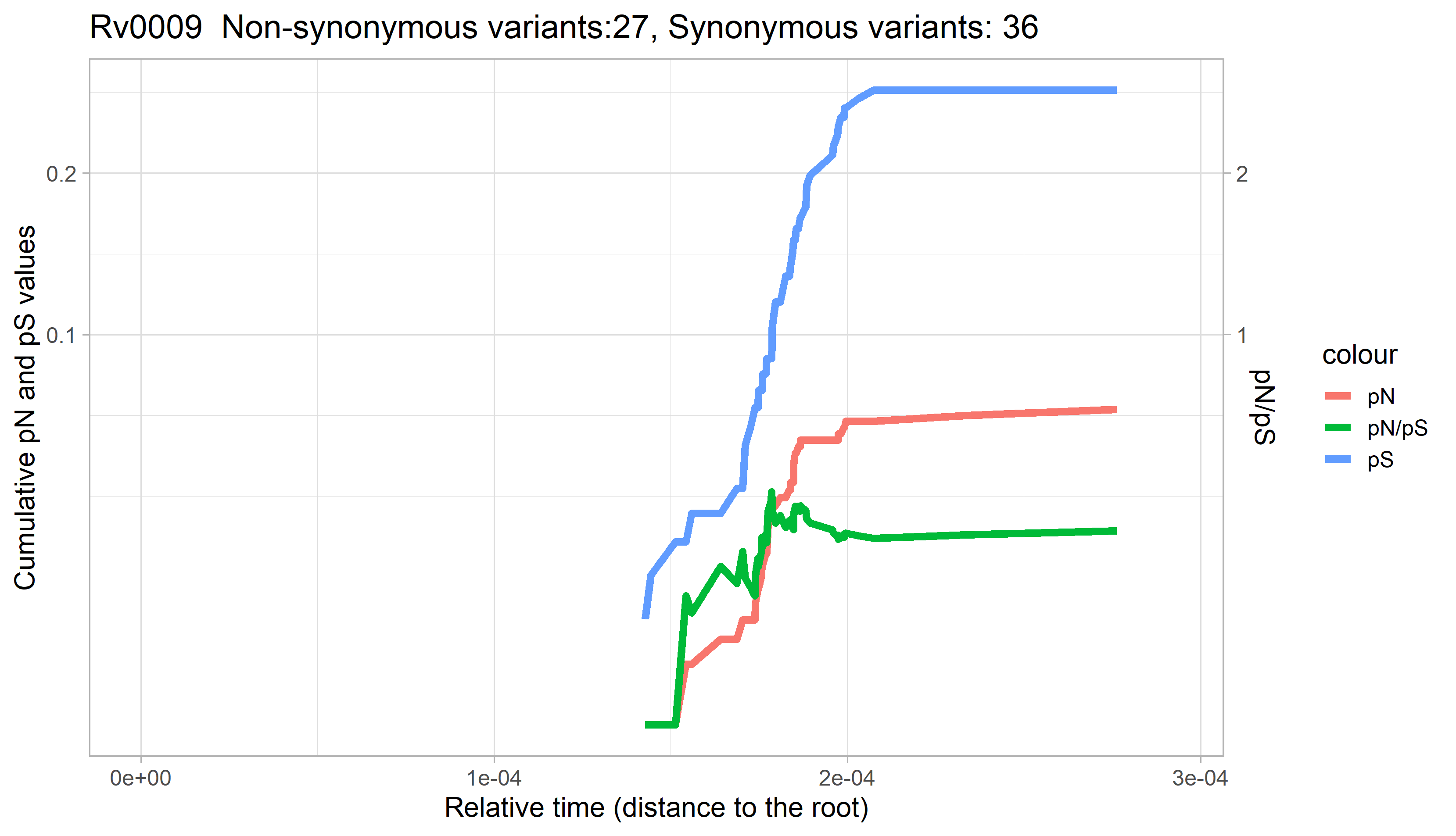

### Rv0010c.png

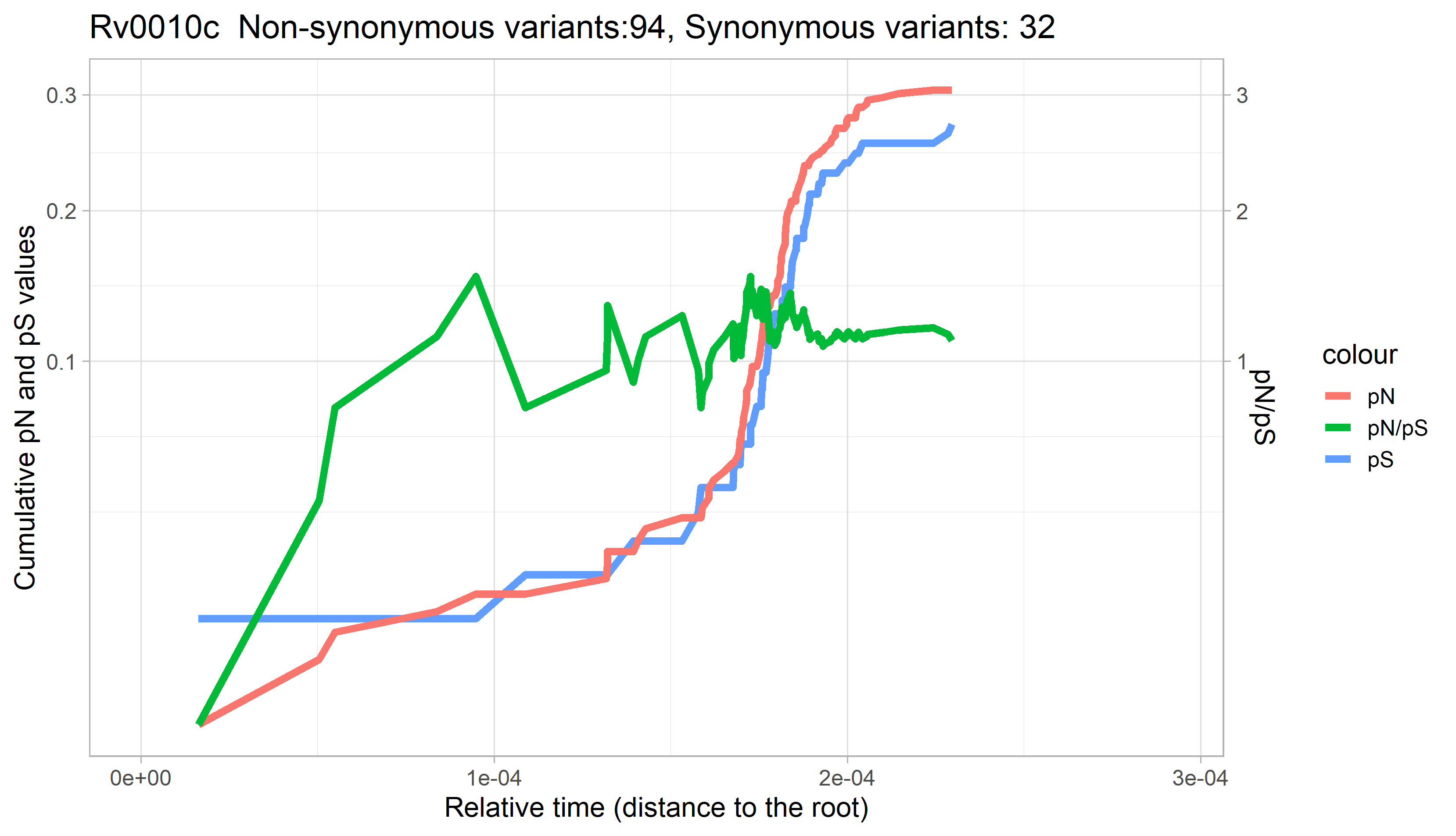

### Rv0011c.png

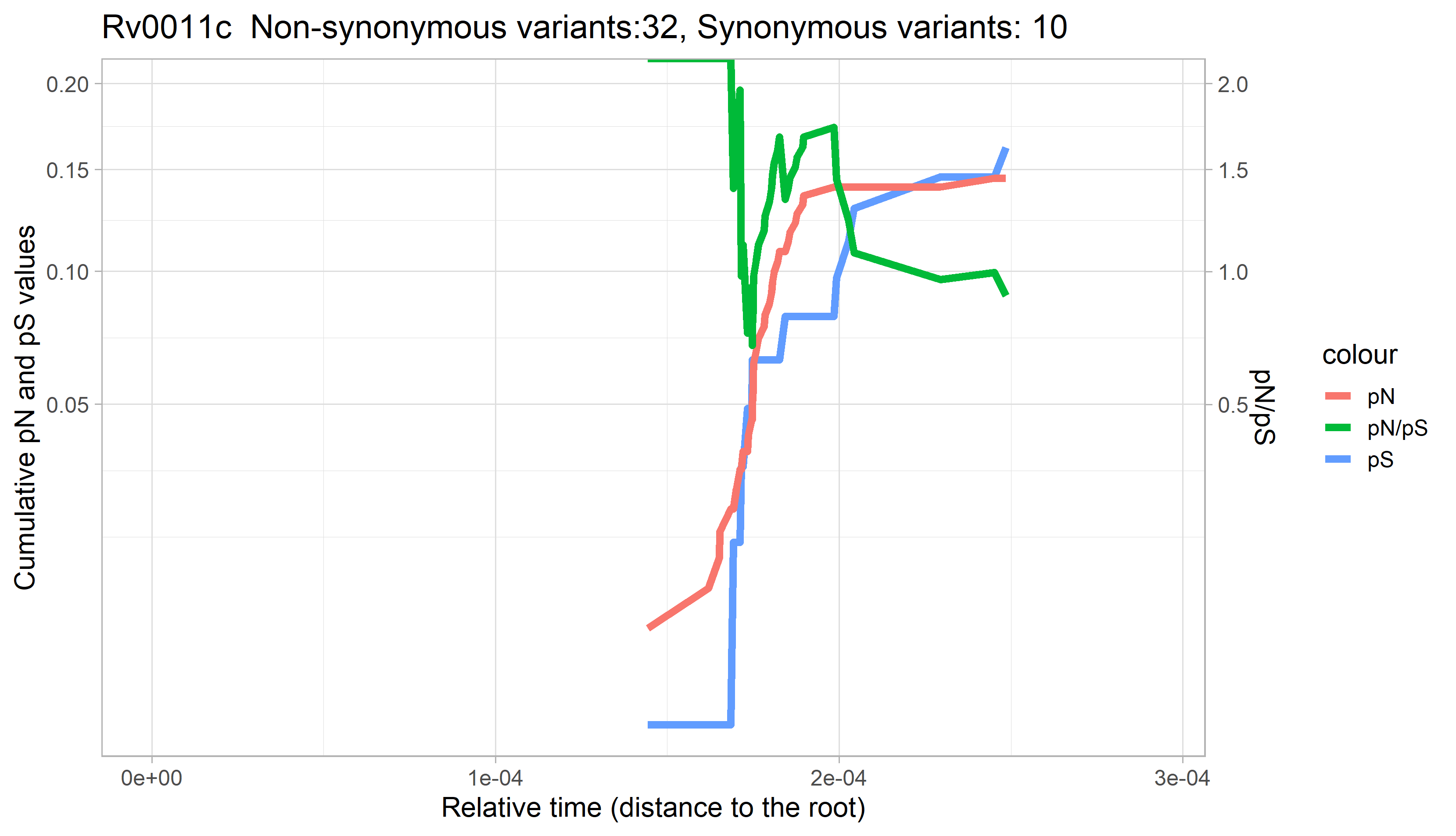

### Rv0012.png

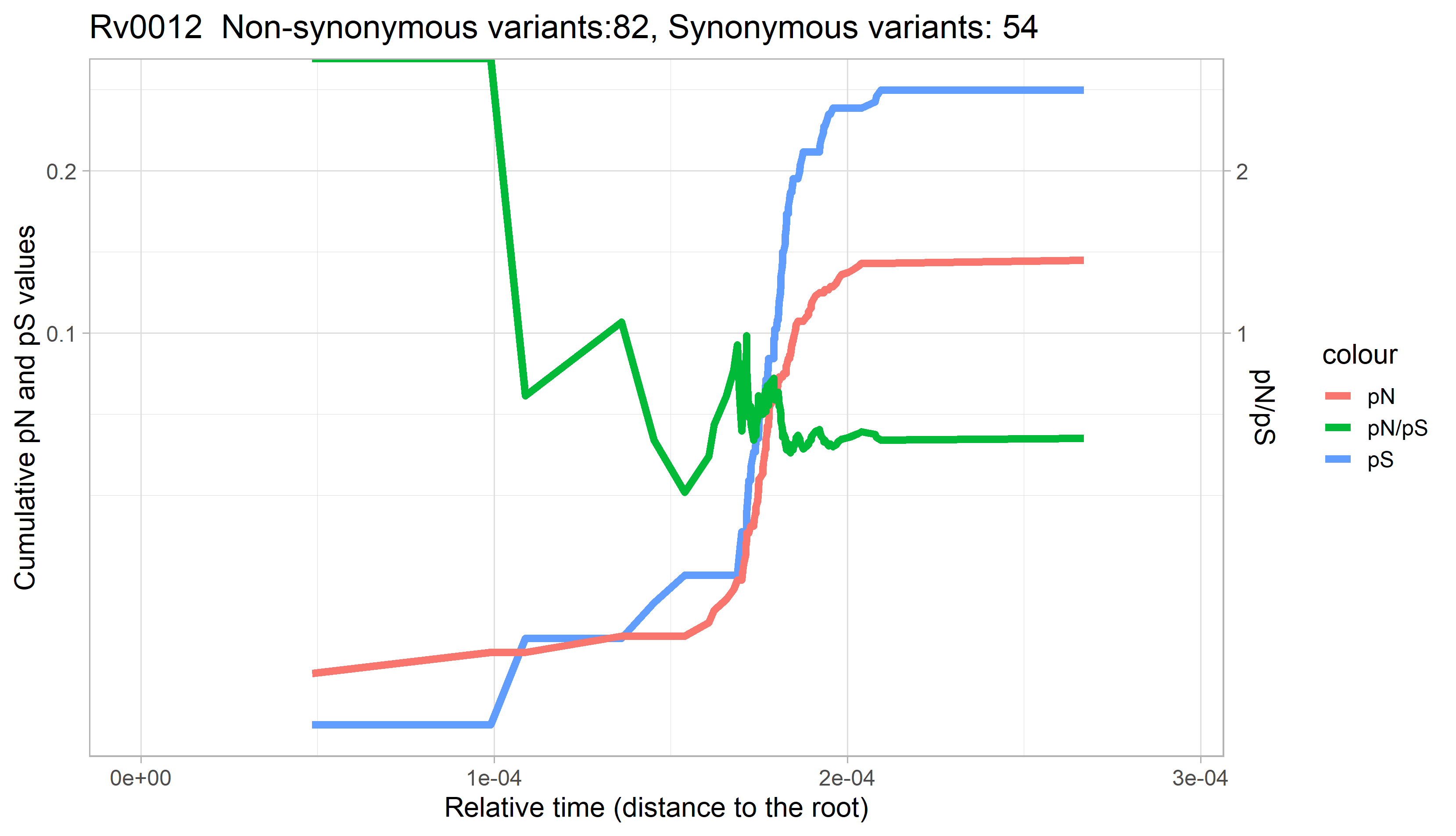

### Rv0013.png

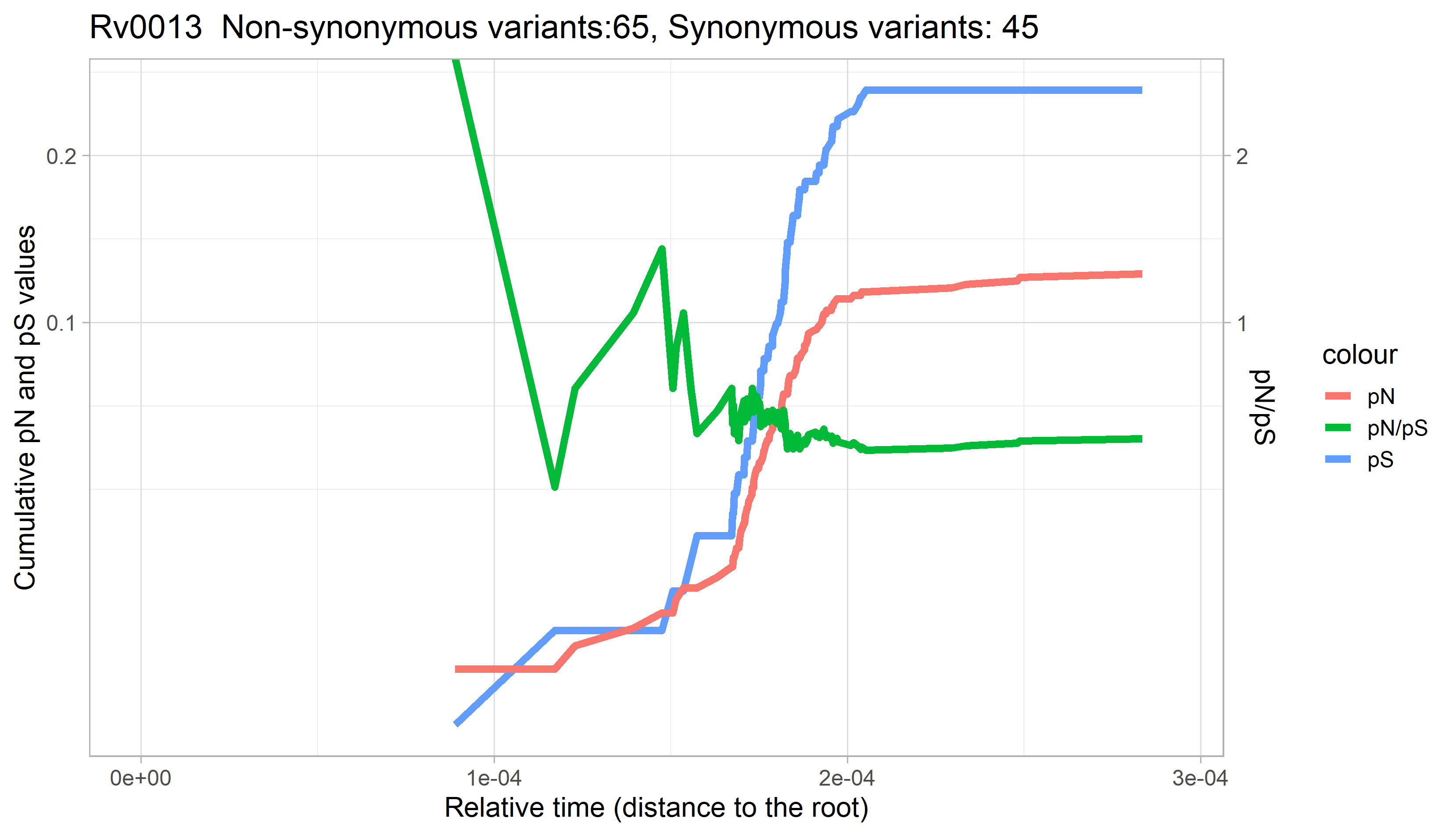

### Rv0014c.png

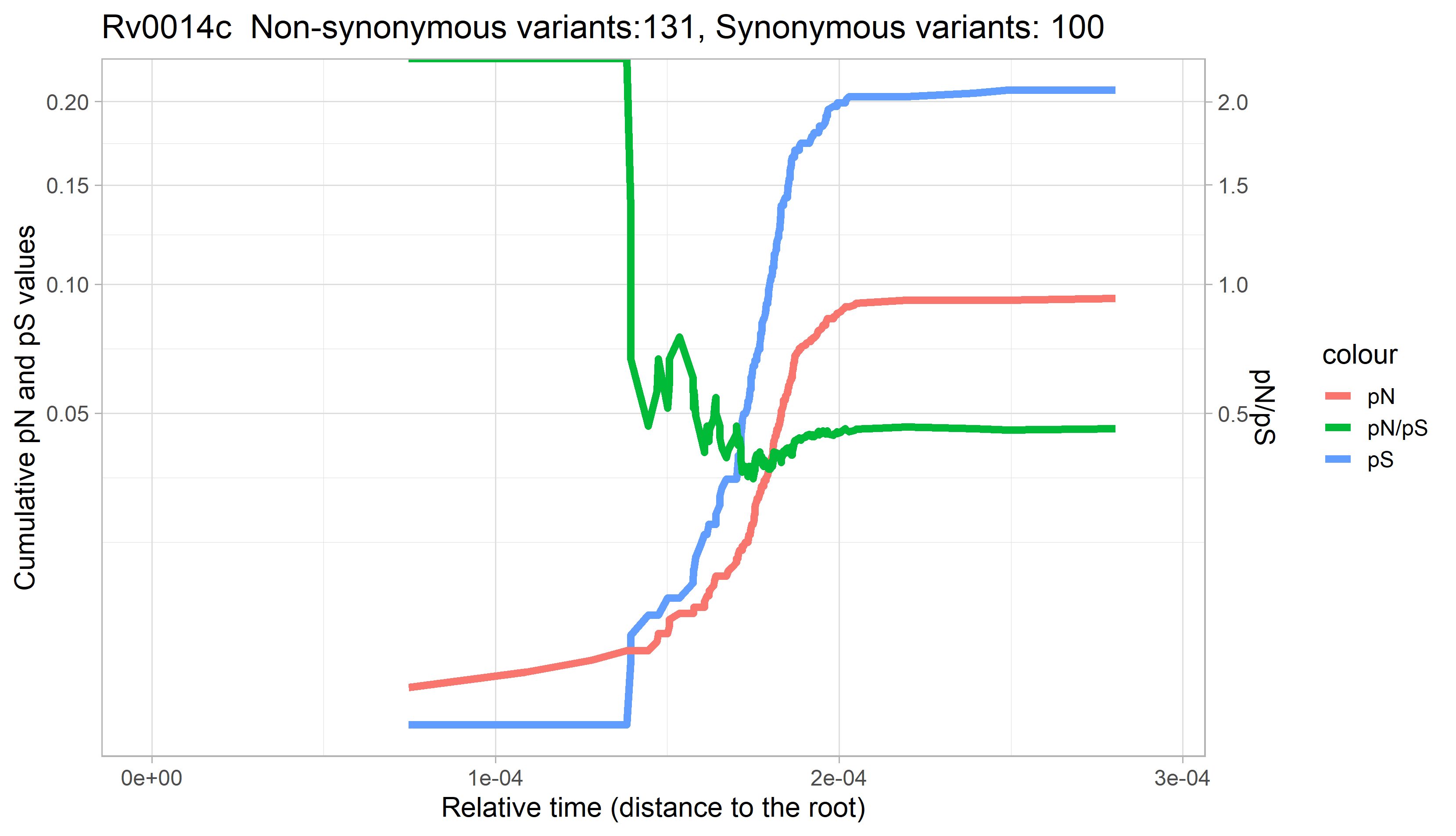

### Rv0015c.png

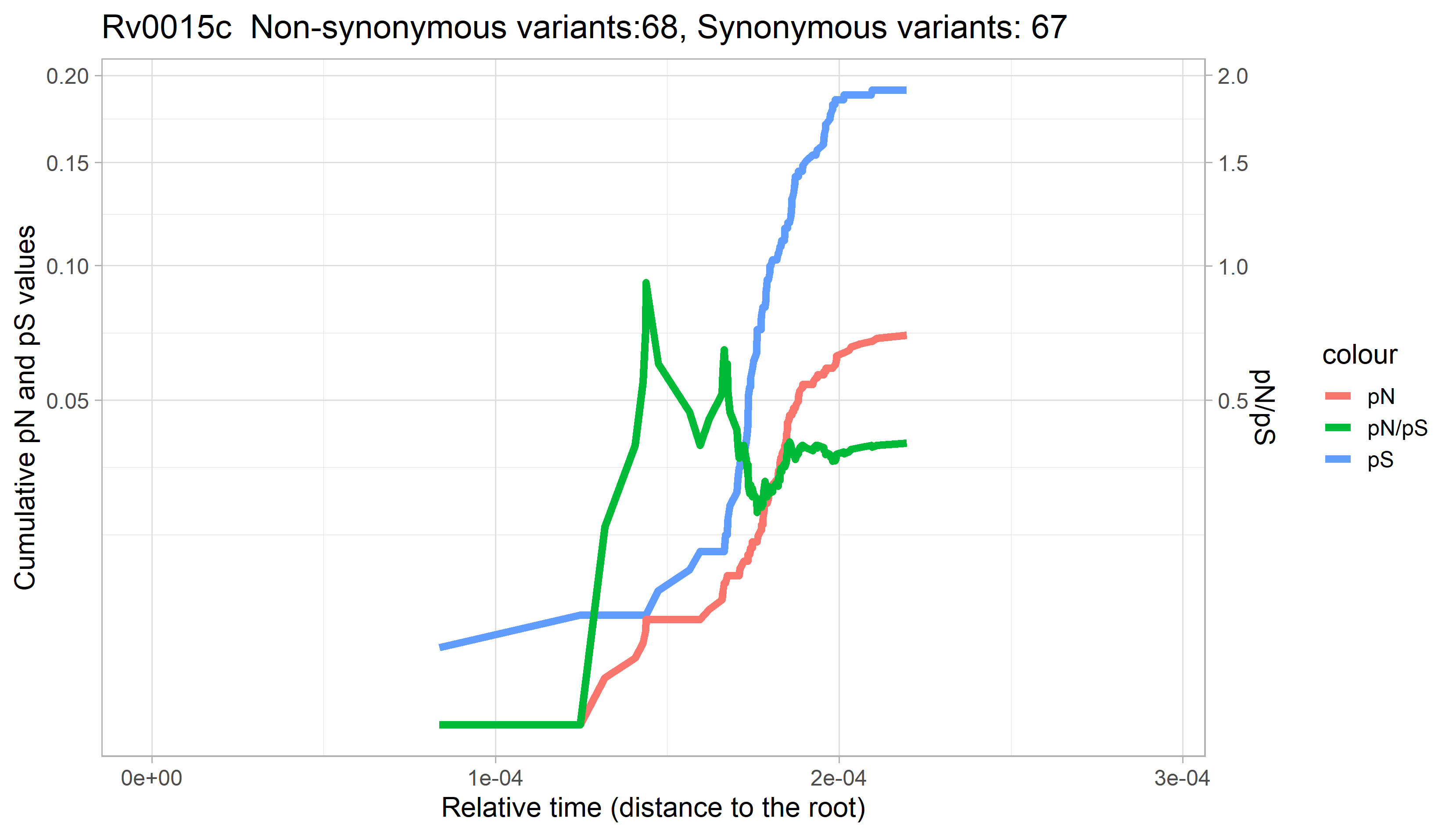

### Rv0016c.png

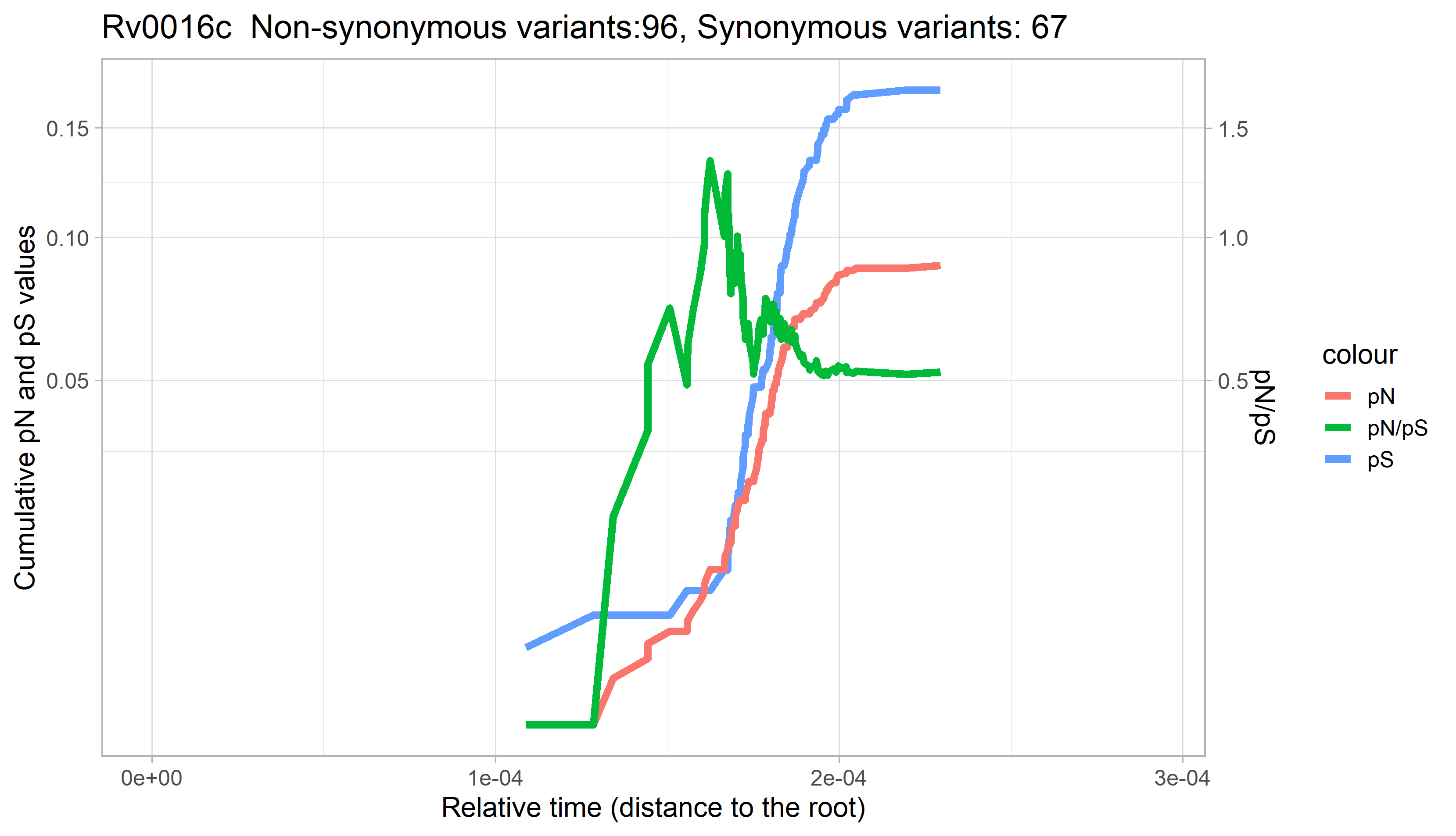

### Rv0017c.png

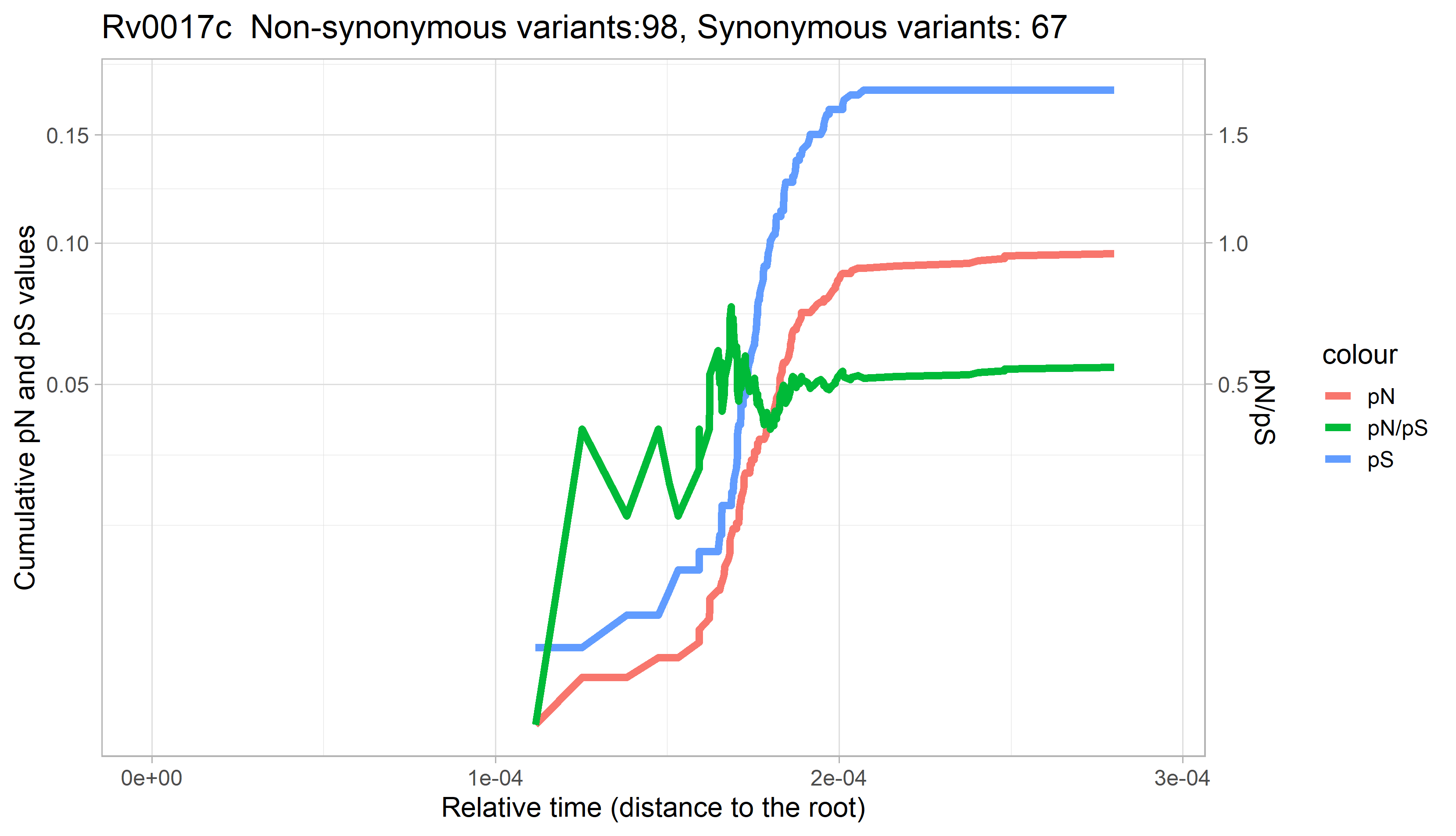

### Rv0018c.png

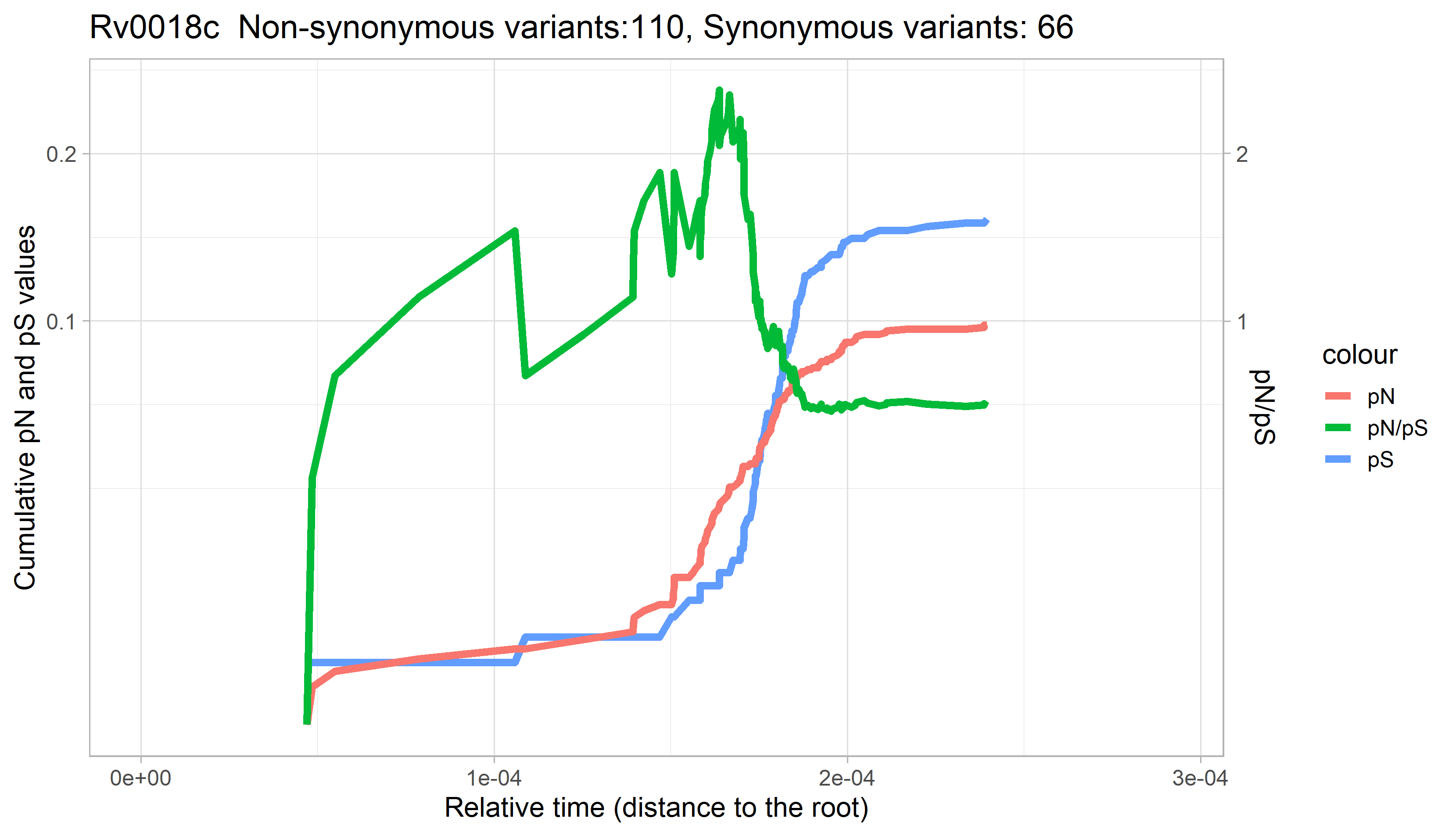

### Rv0019c.png

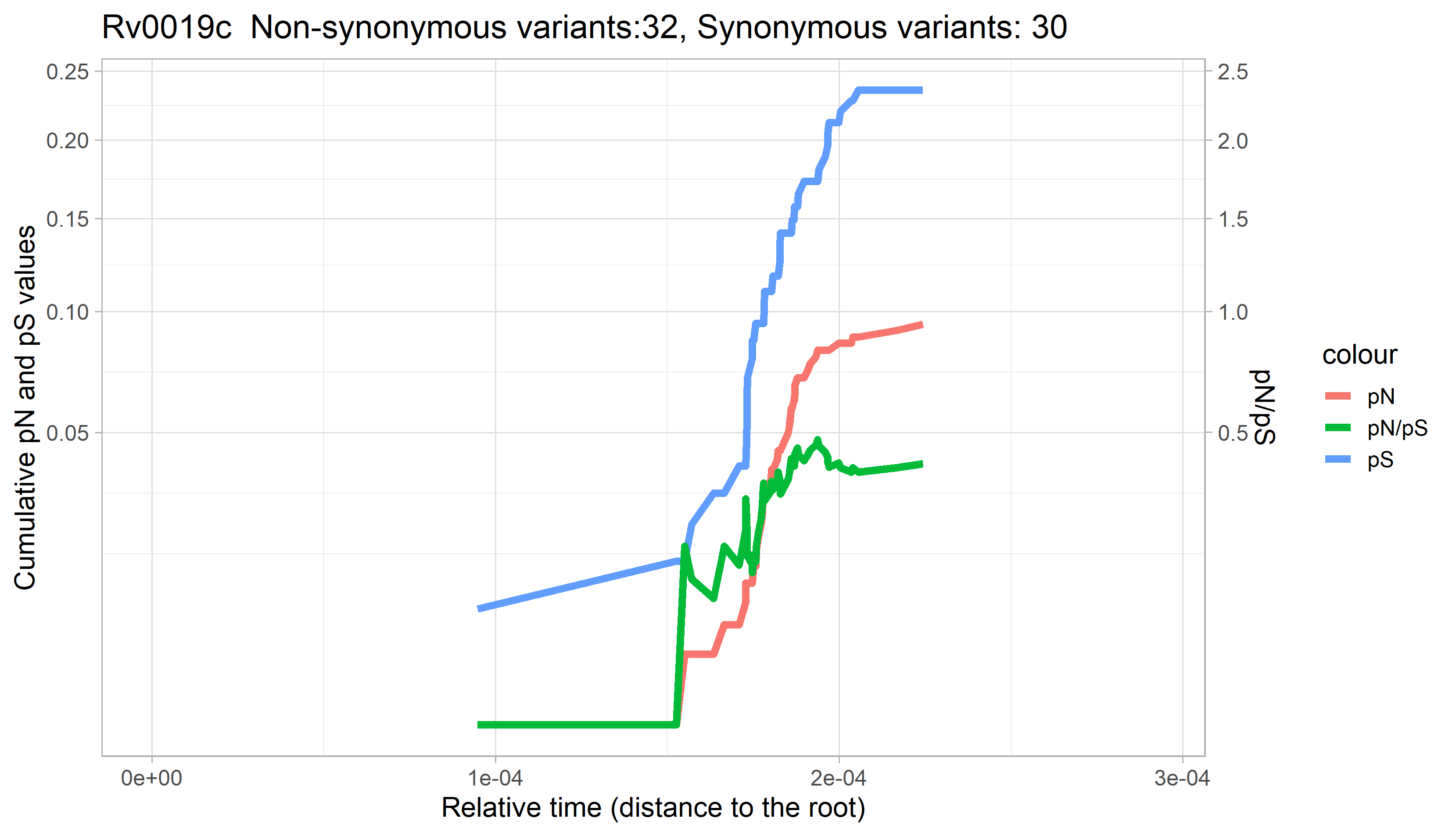

### Rv0020c.png

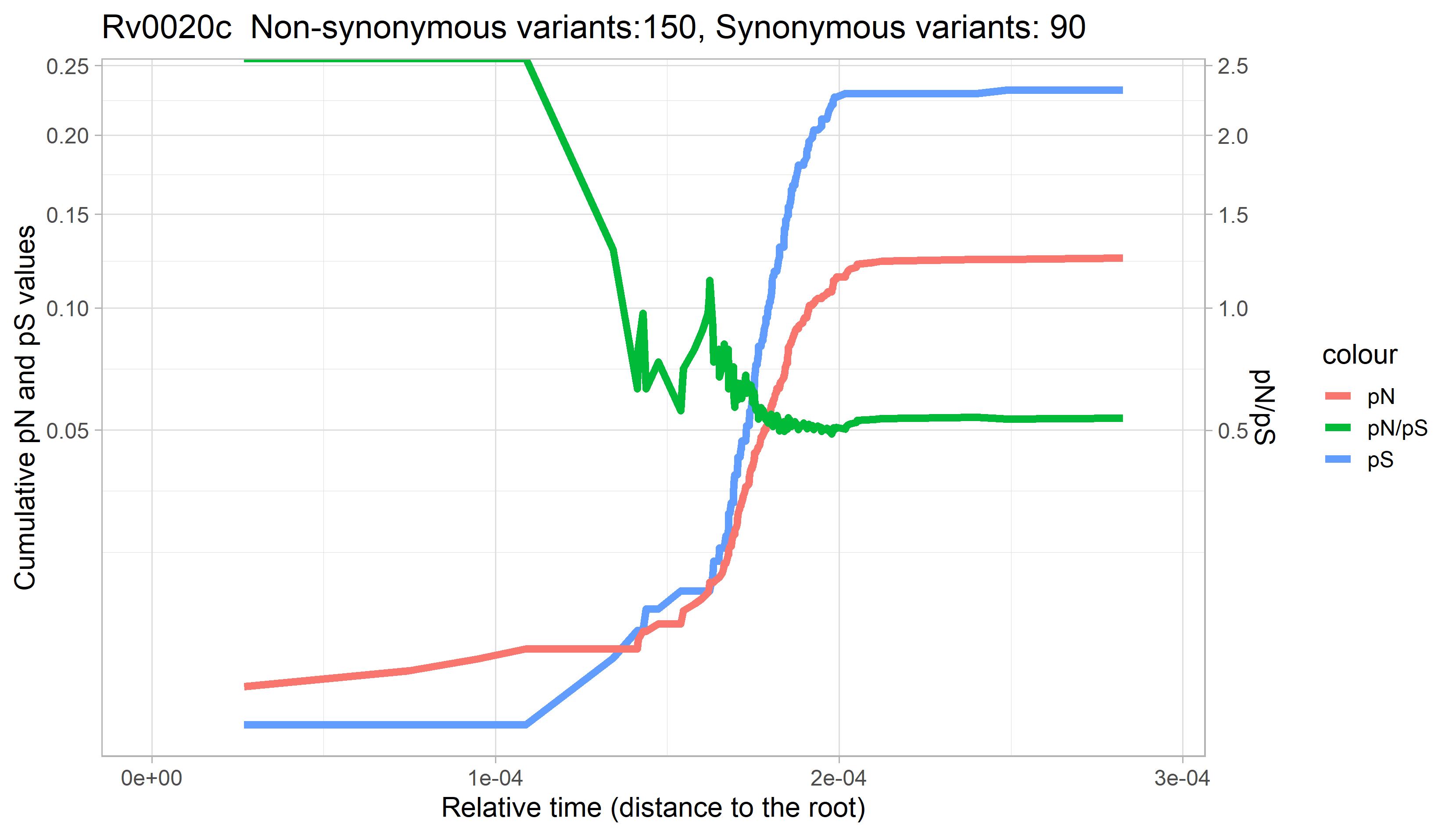

### Rv0021c.png

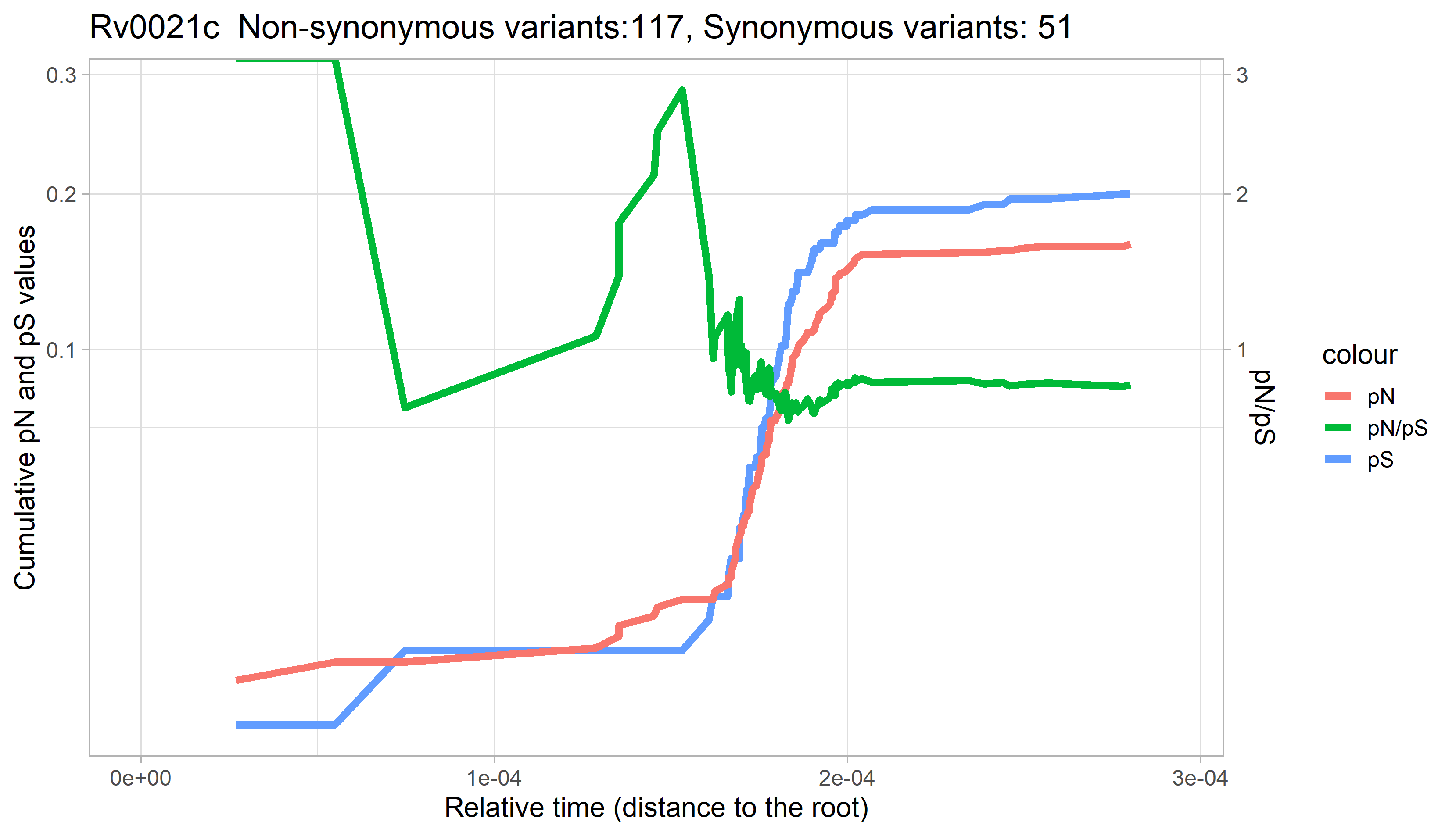

### Rv0022c.png

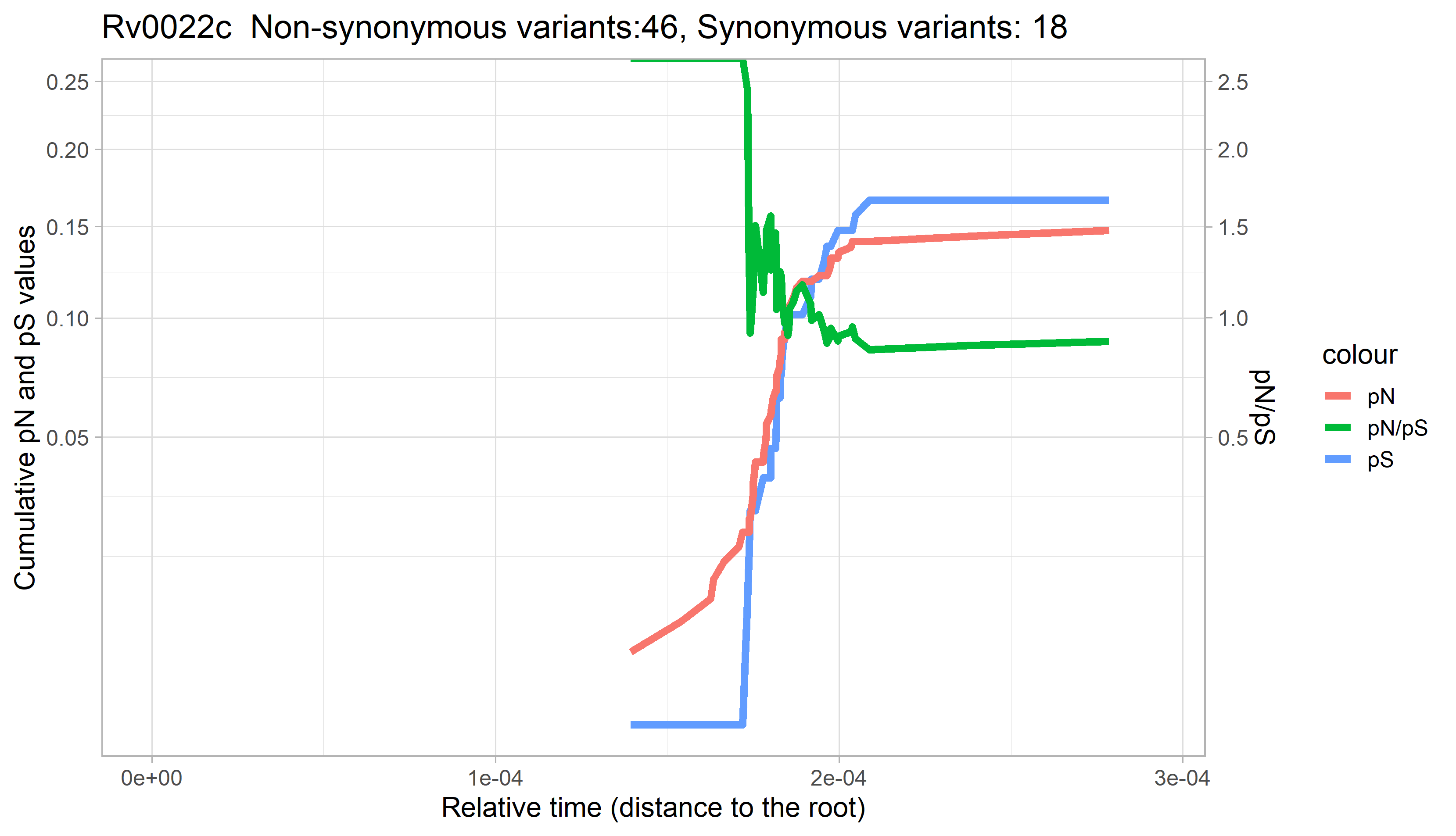

### Rv0023.png

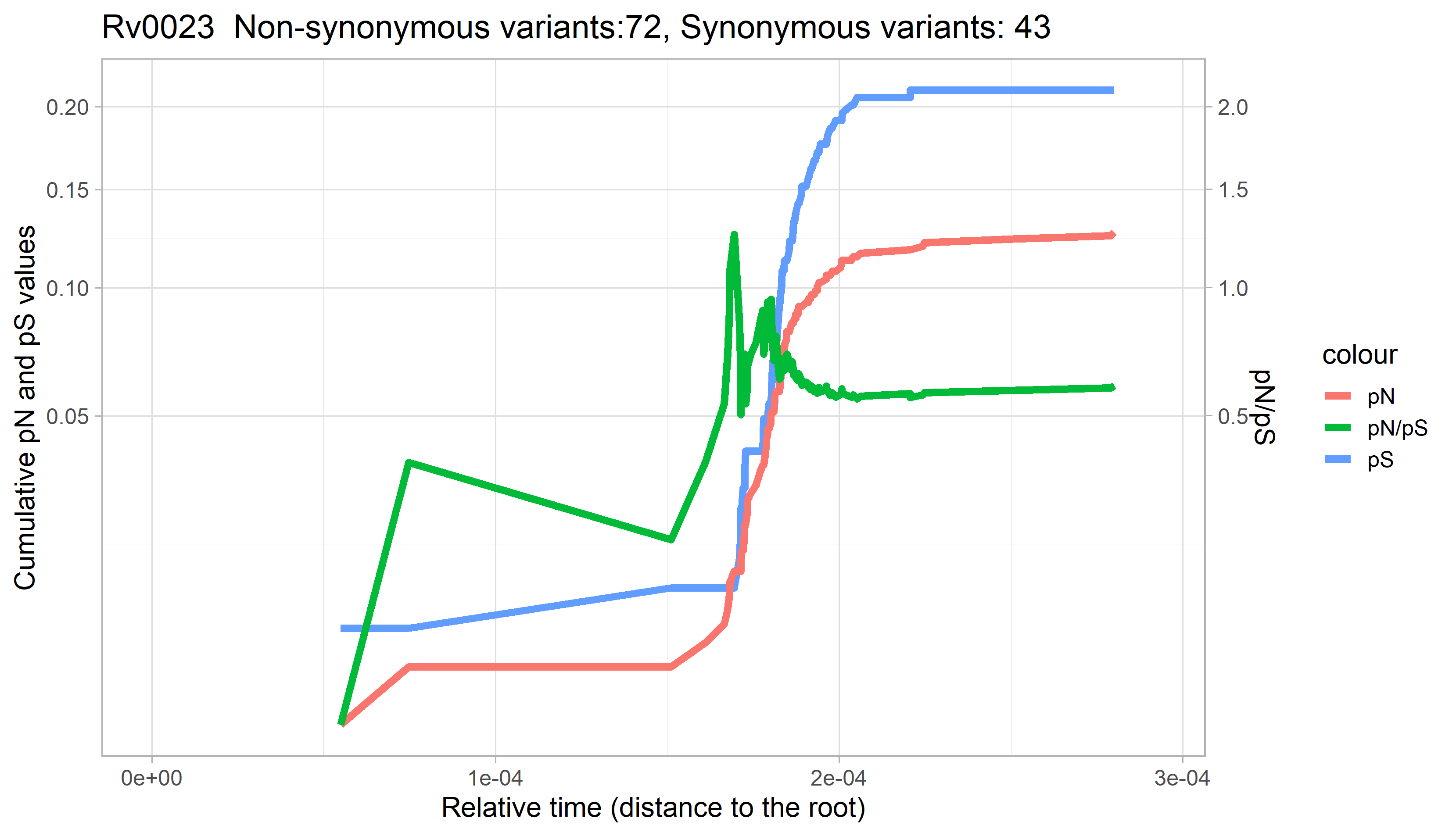

### Rv0024.png

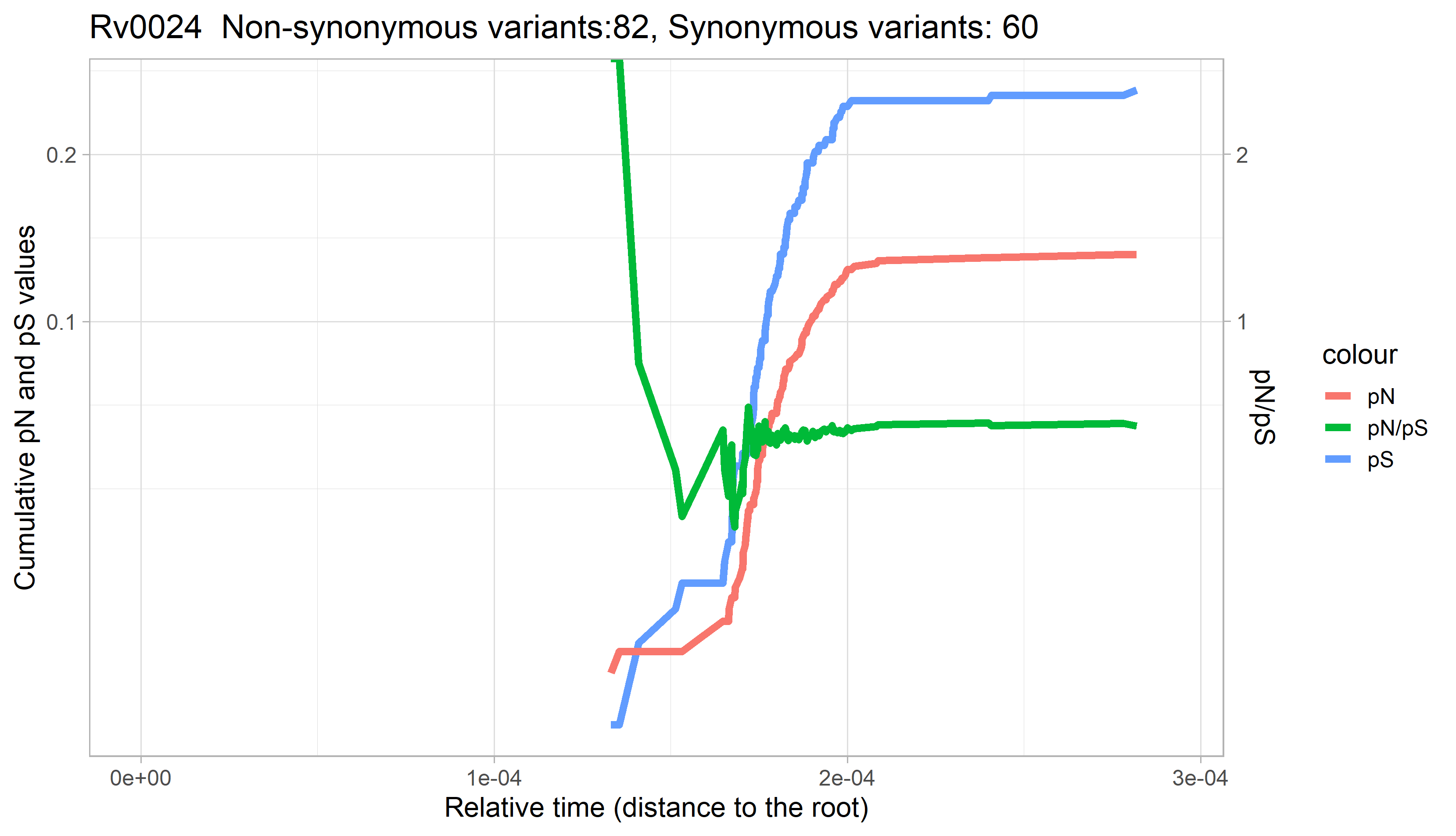

### Rv0025.png

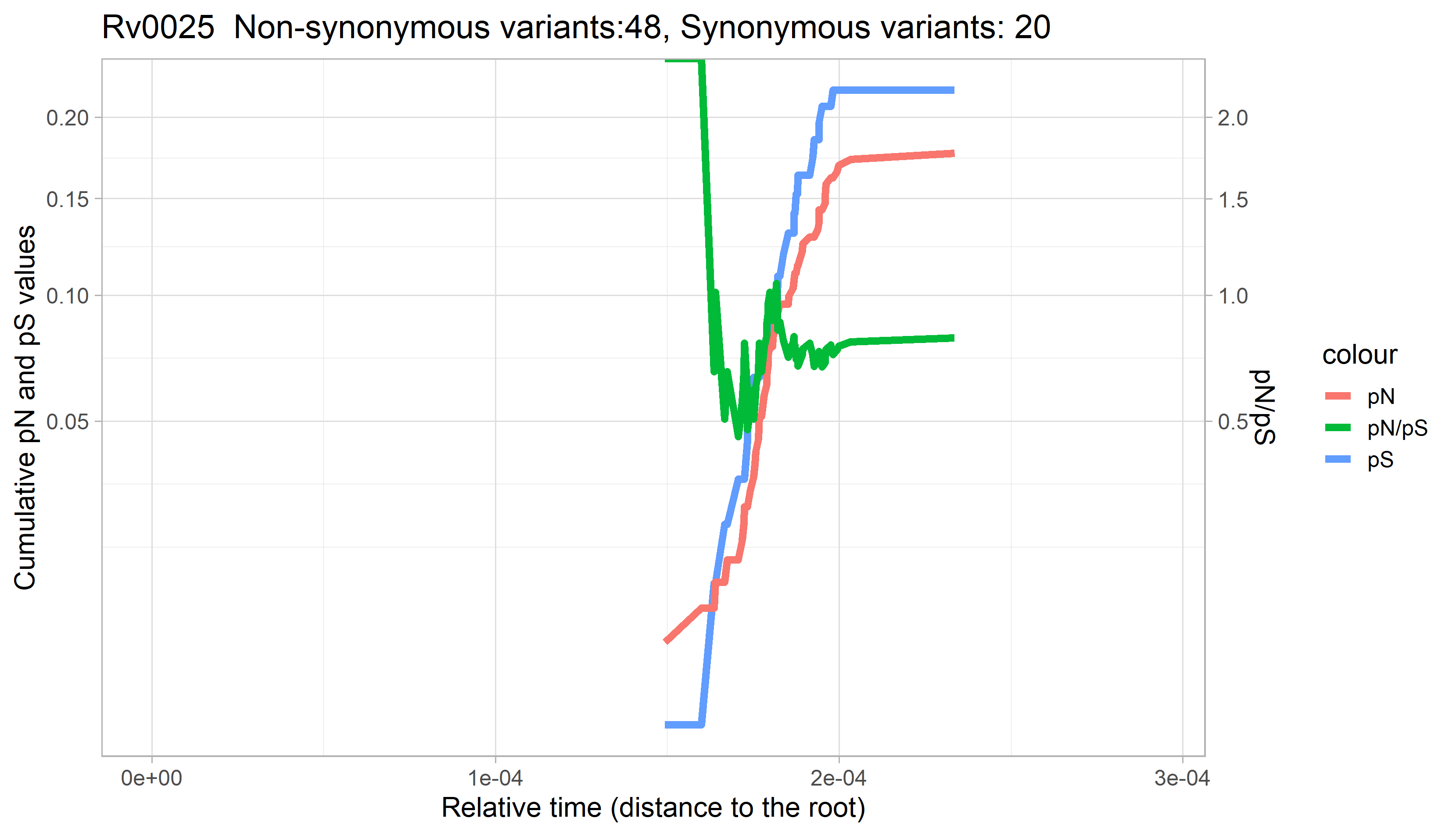

### Rv0026.png

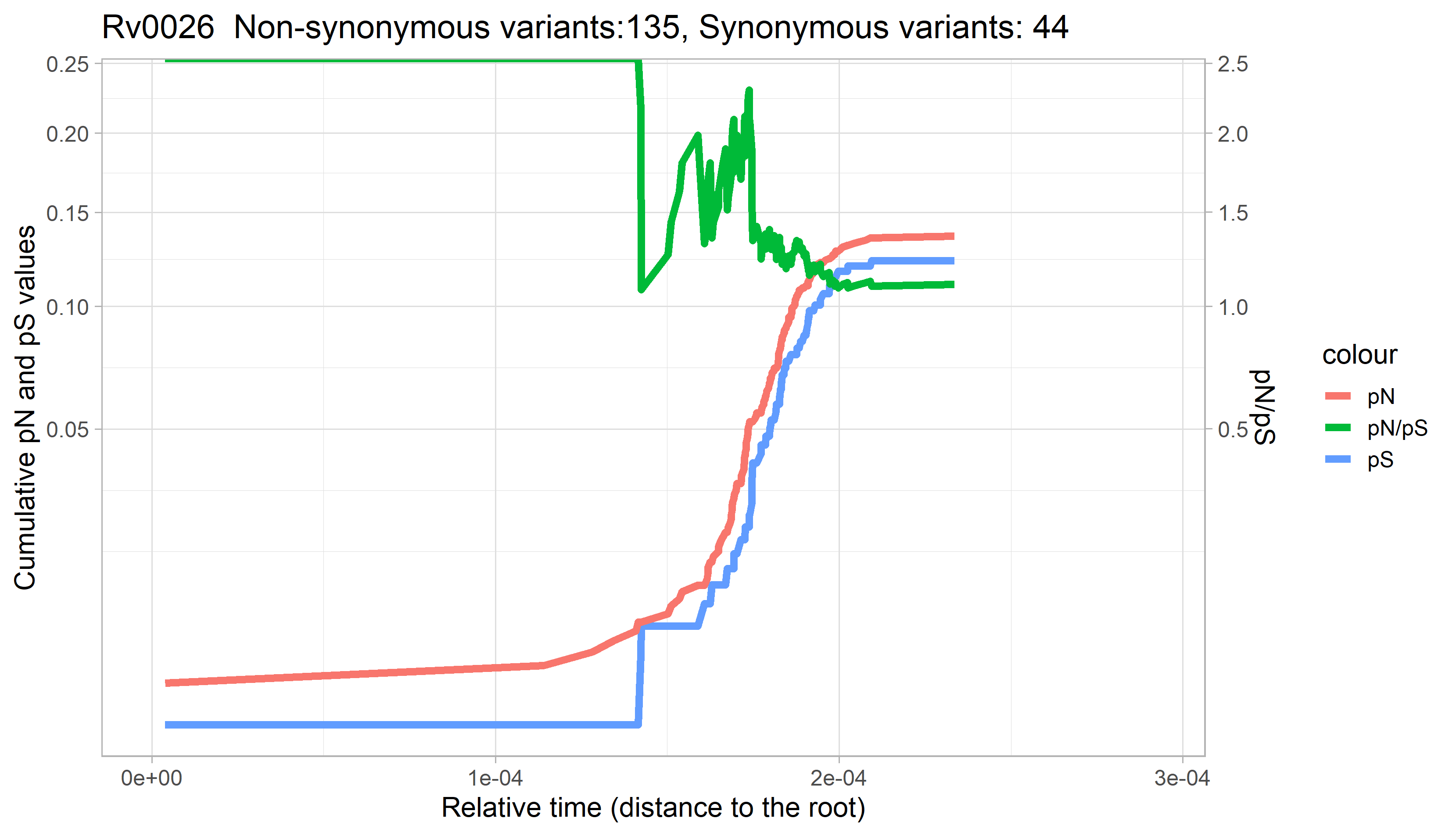

### Rv0027.png

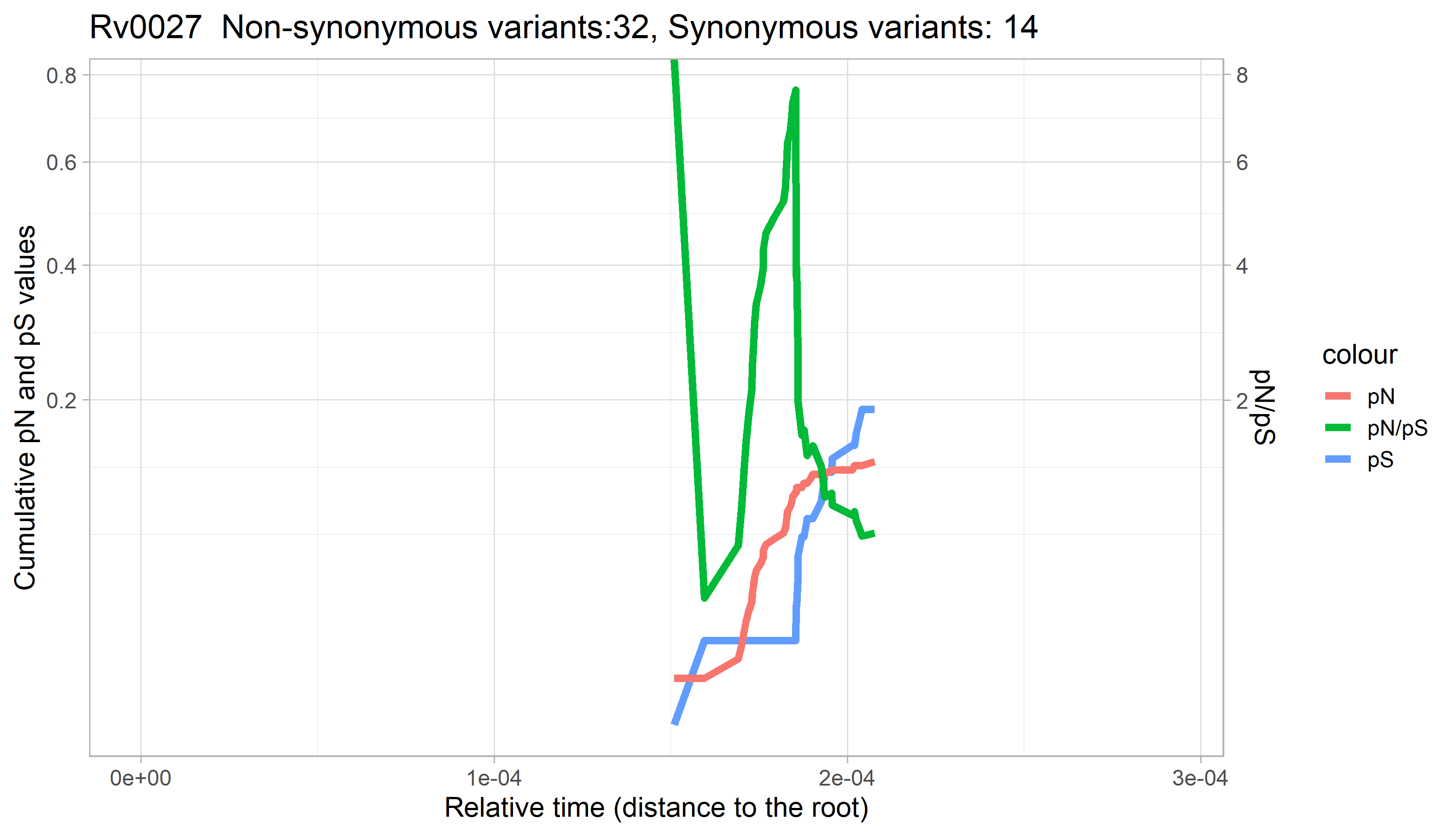

### Rv0028.png

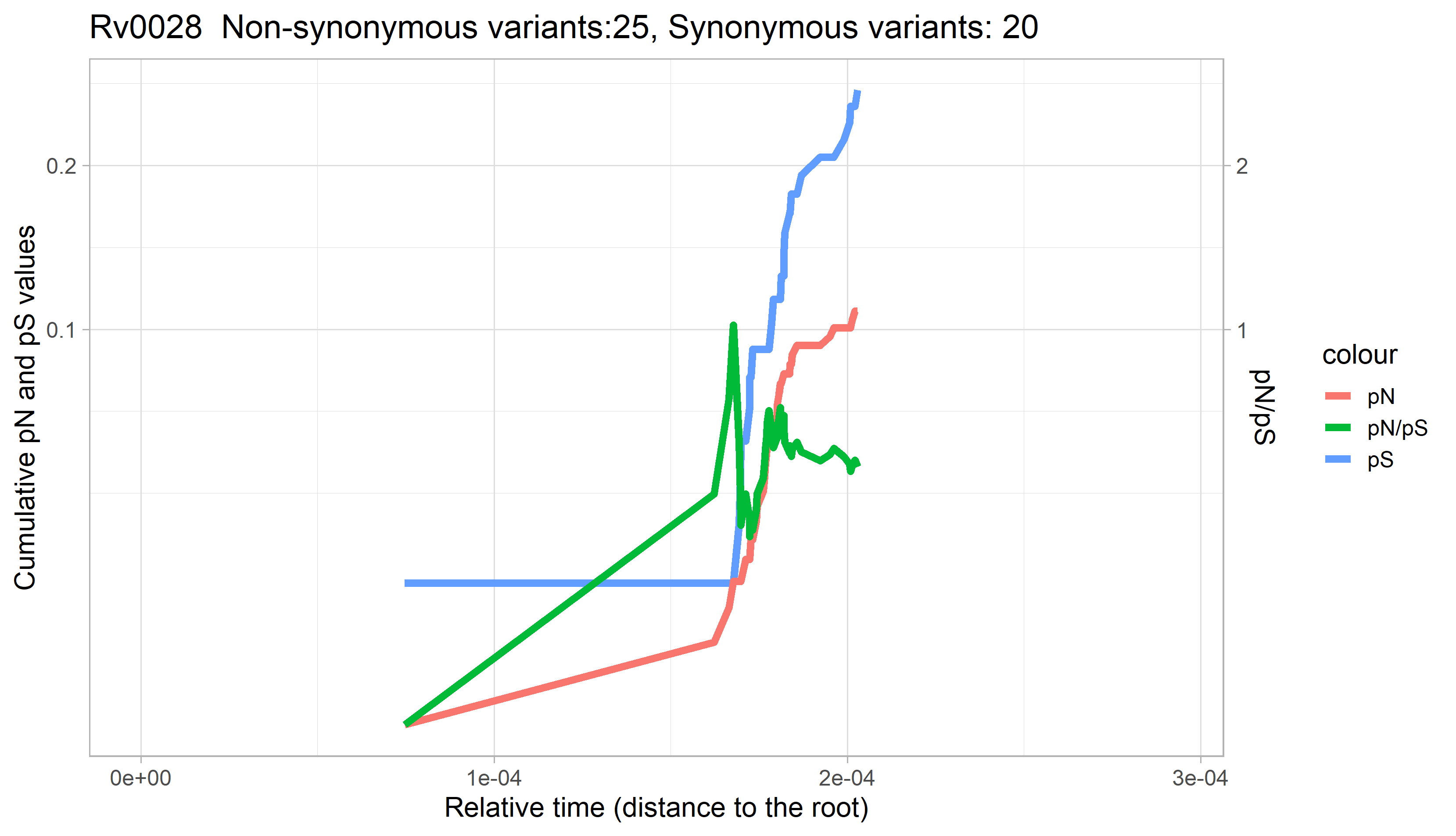

### Rv0029.png

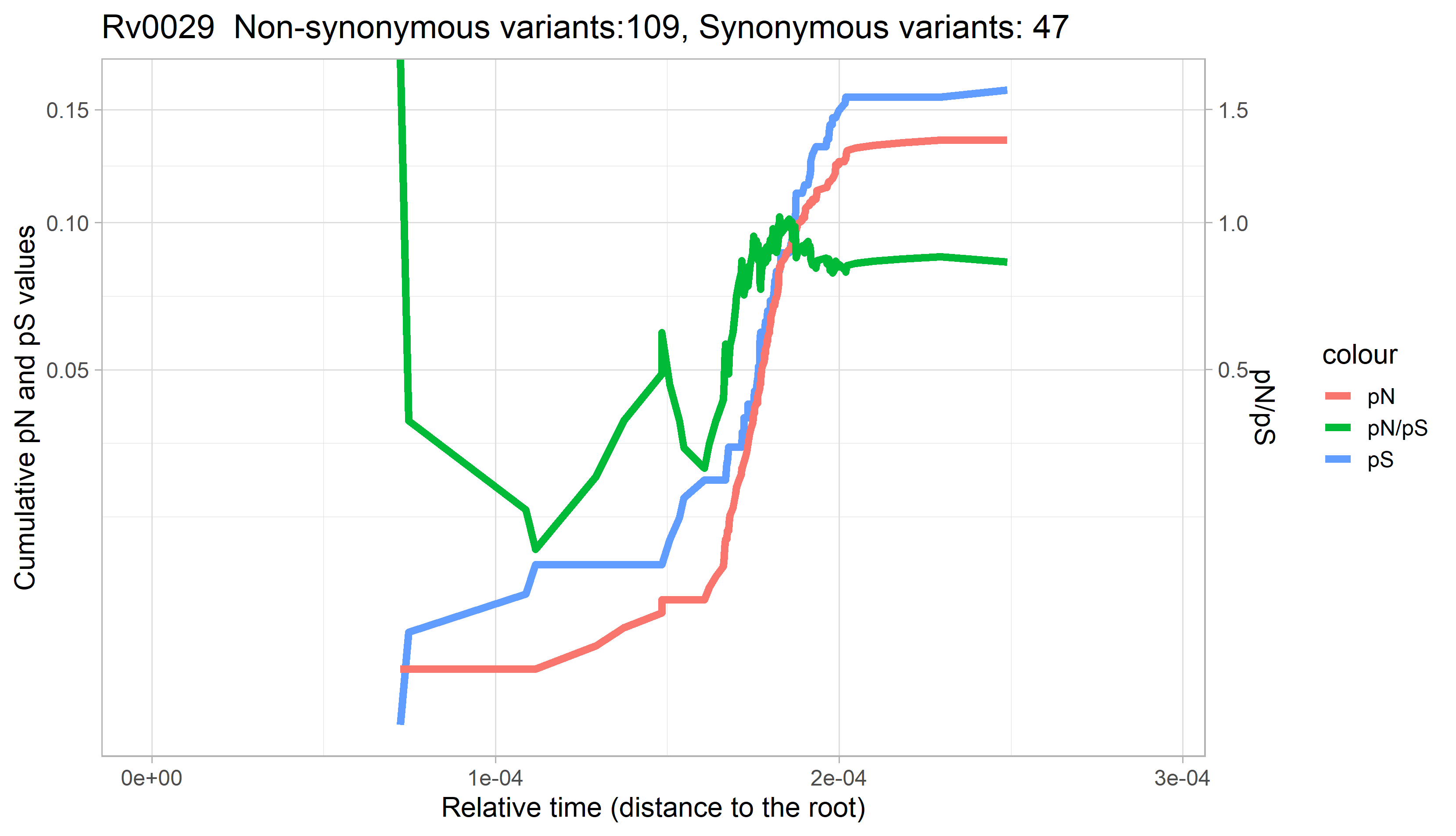

### Rv0030.png

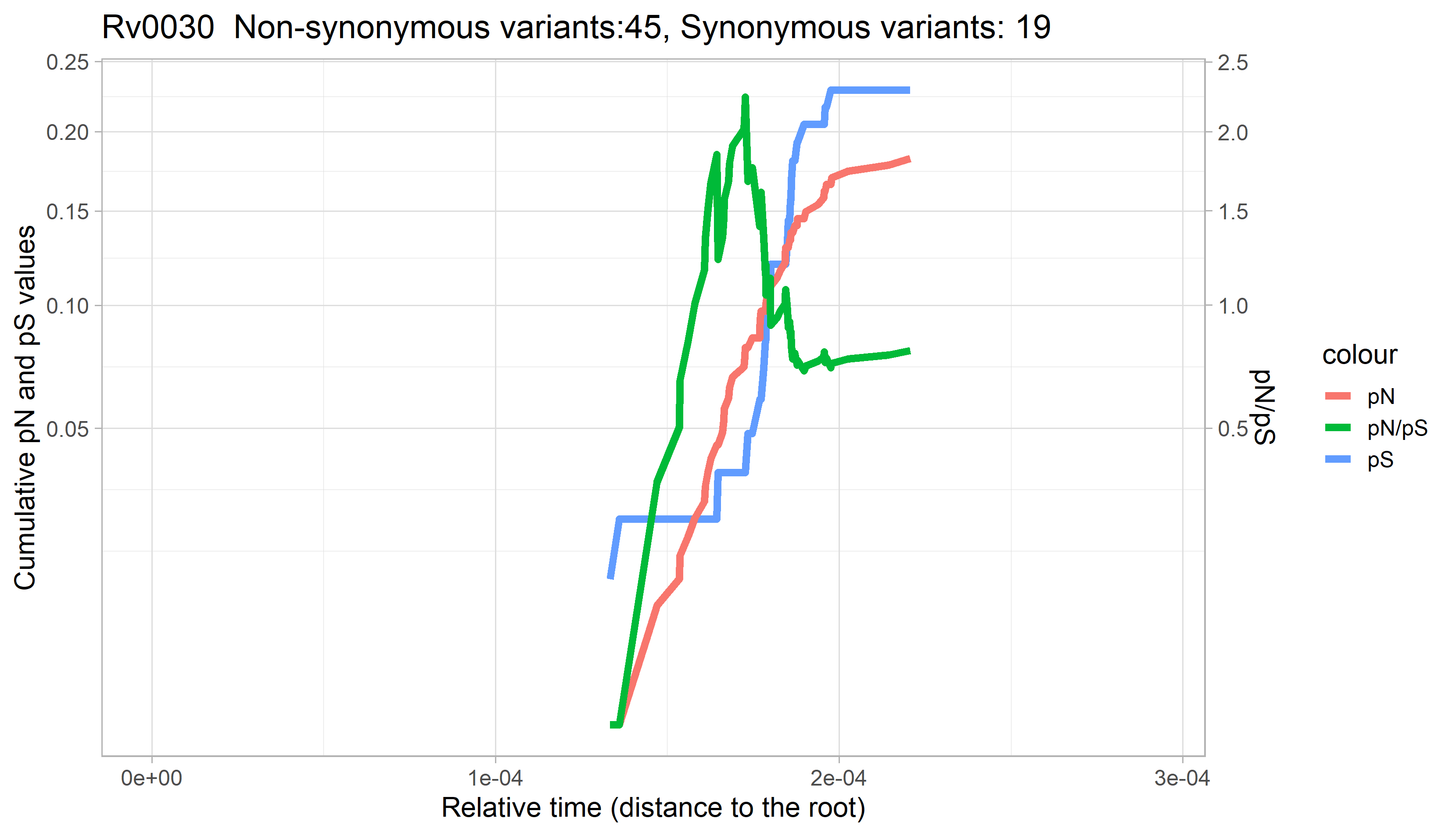

### Rv0032.png

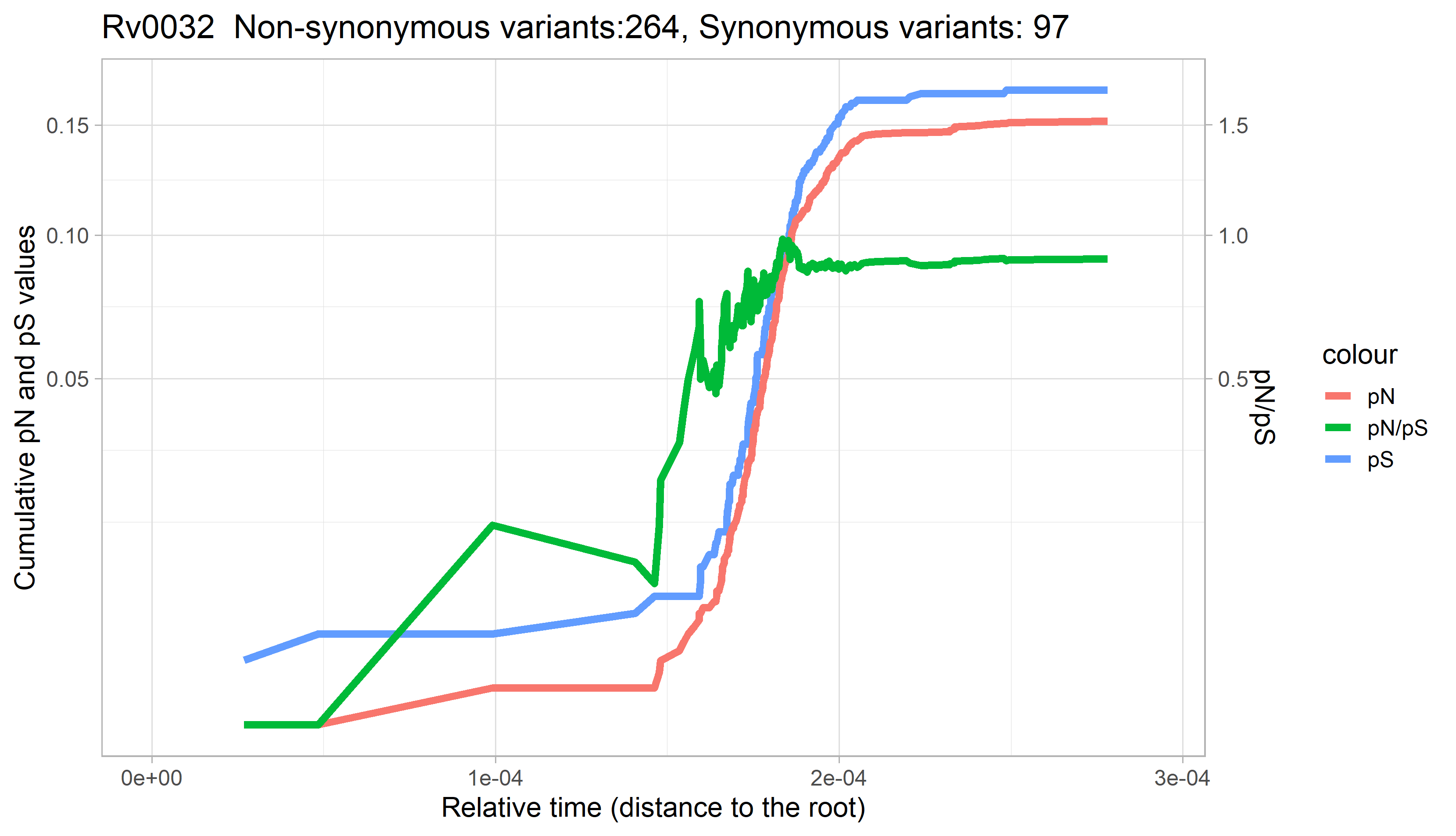

### Rv0033.png

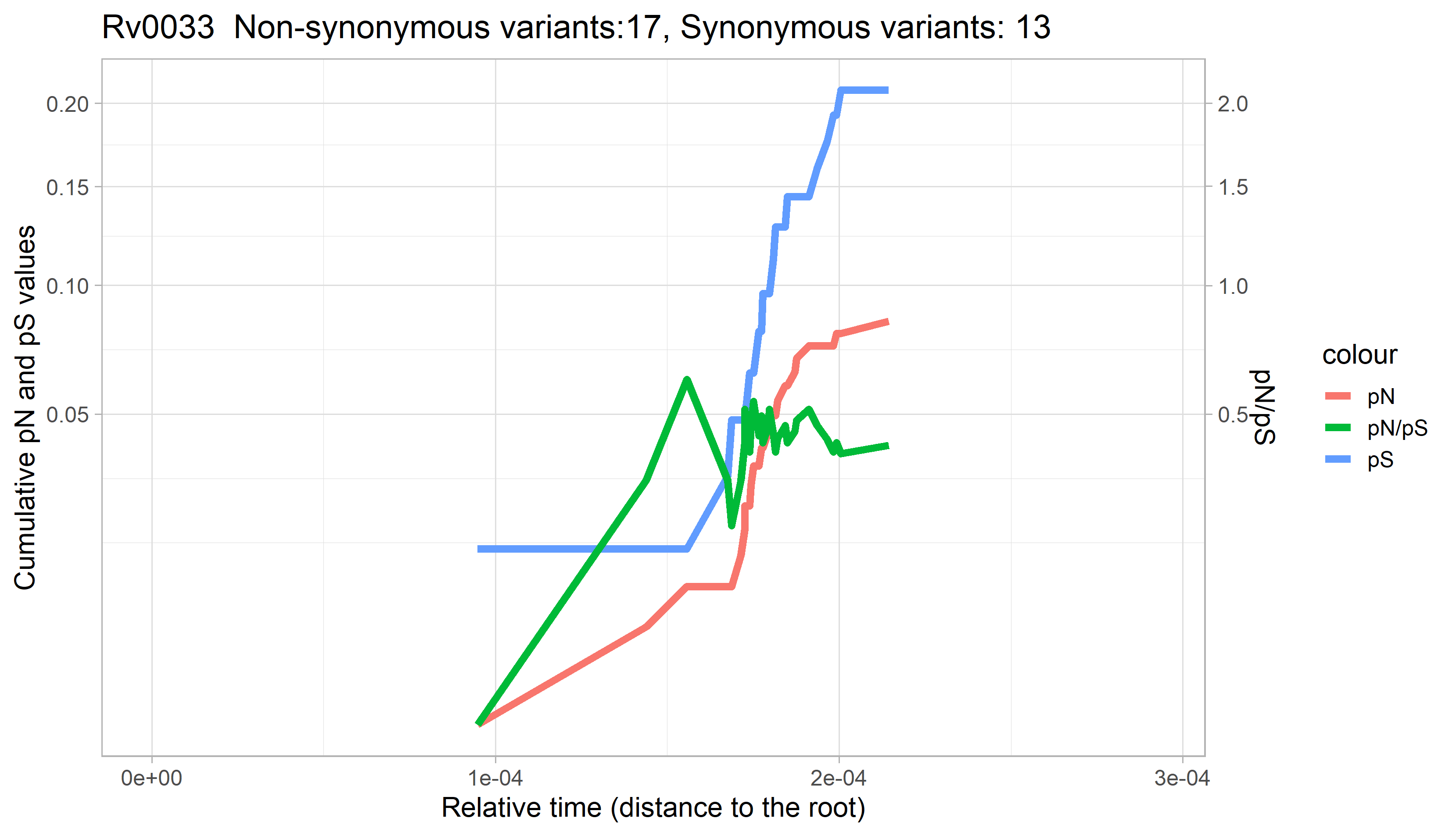

### Rv0034.png

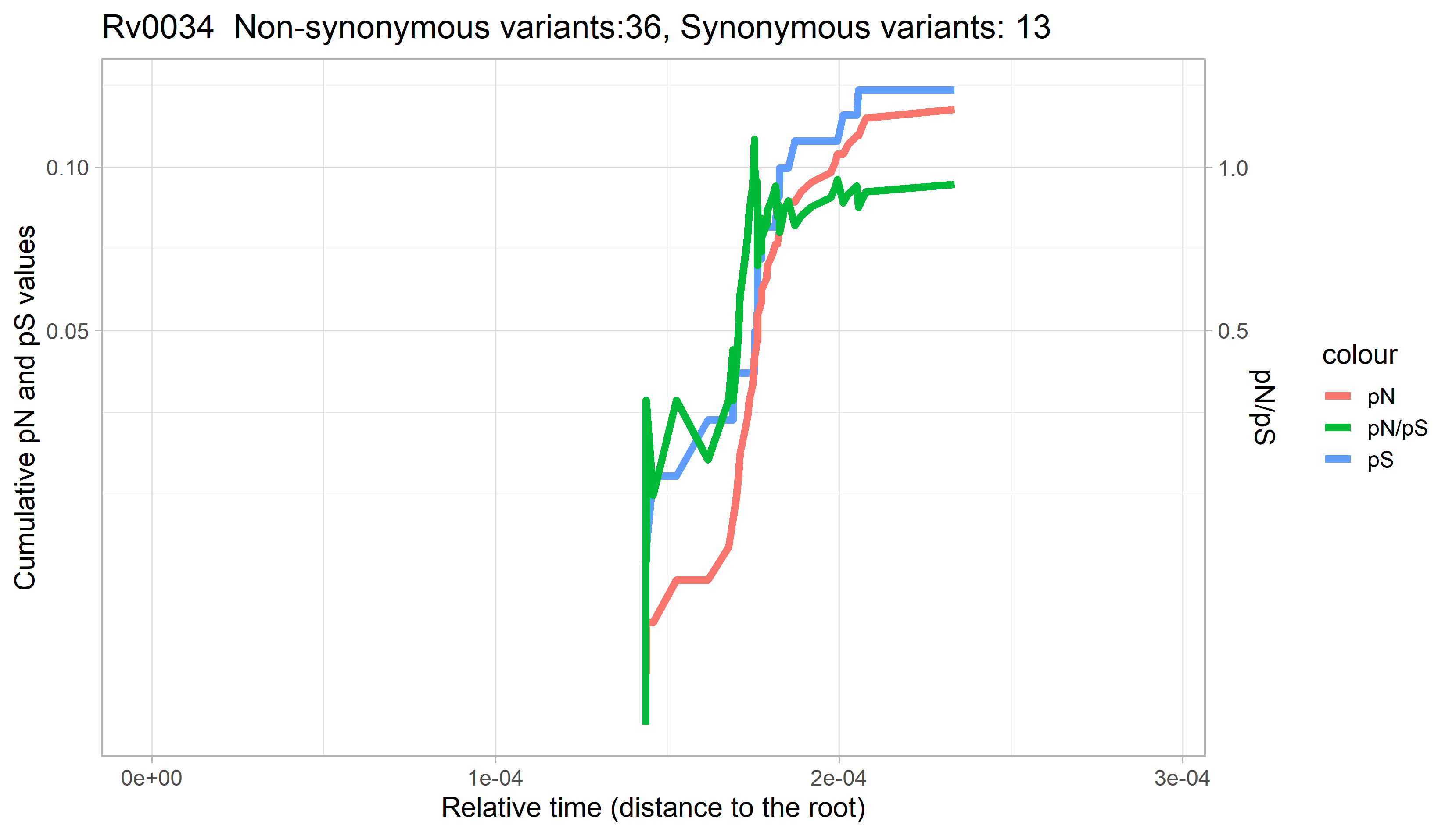

### Rv0035.png

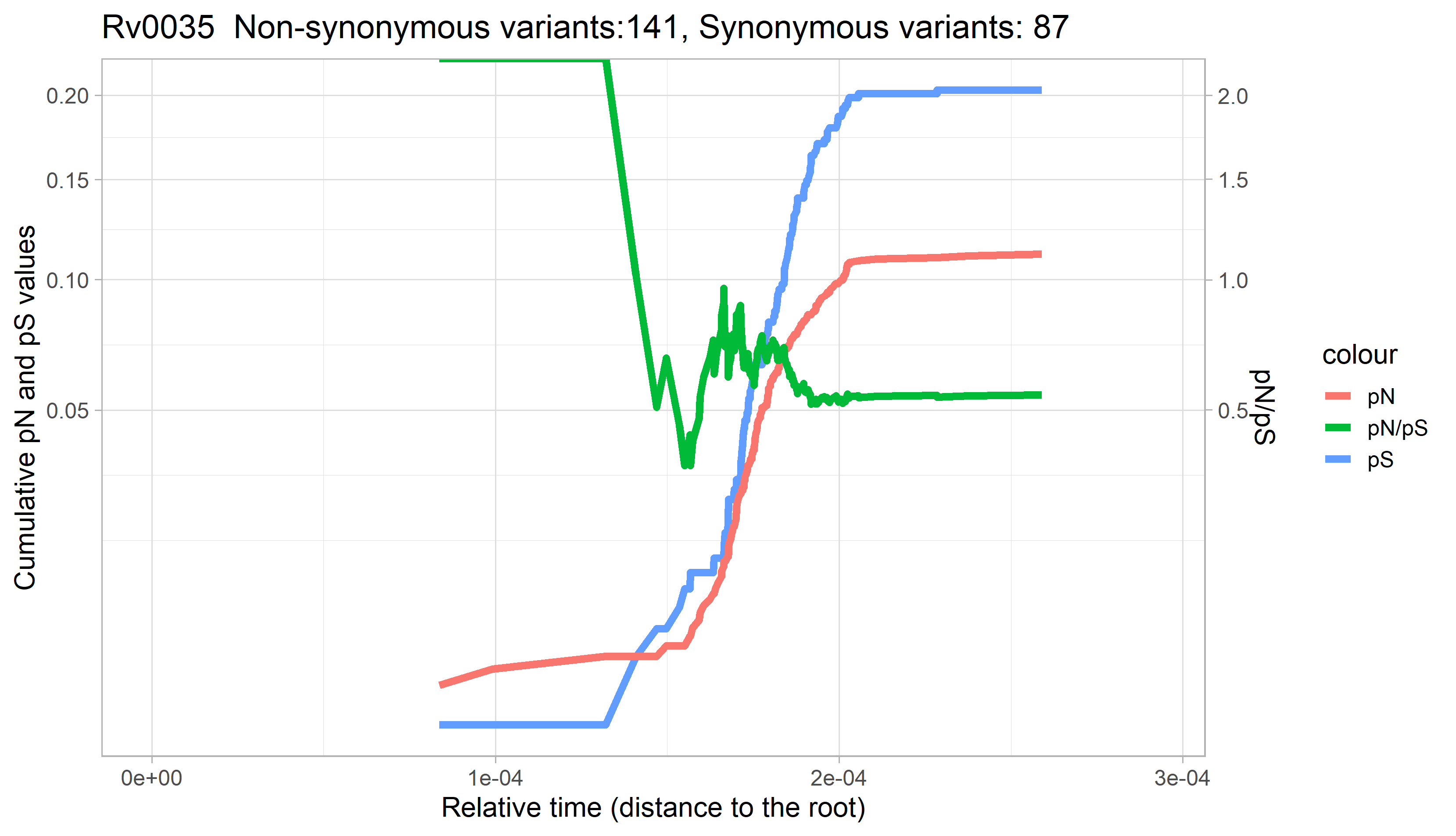

### Rv0036c.png

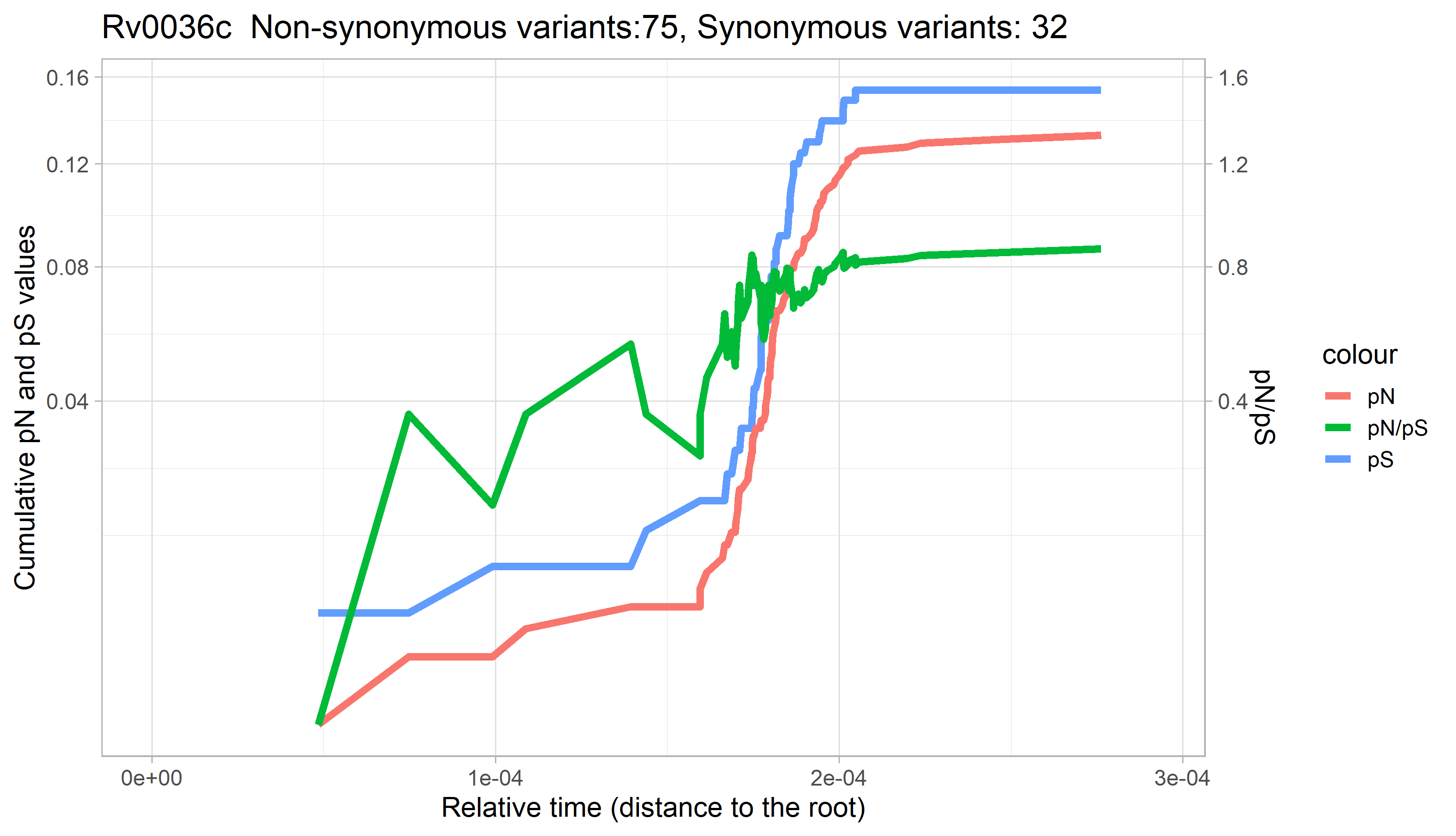

### Rv0037c.png

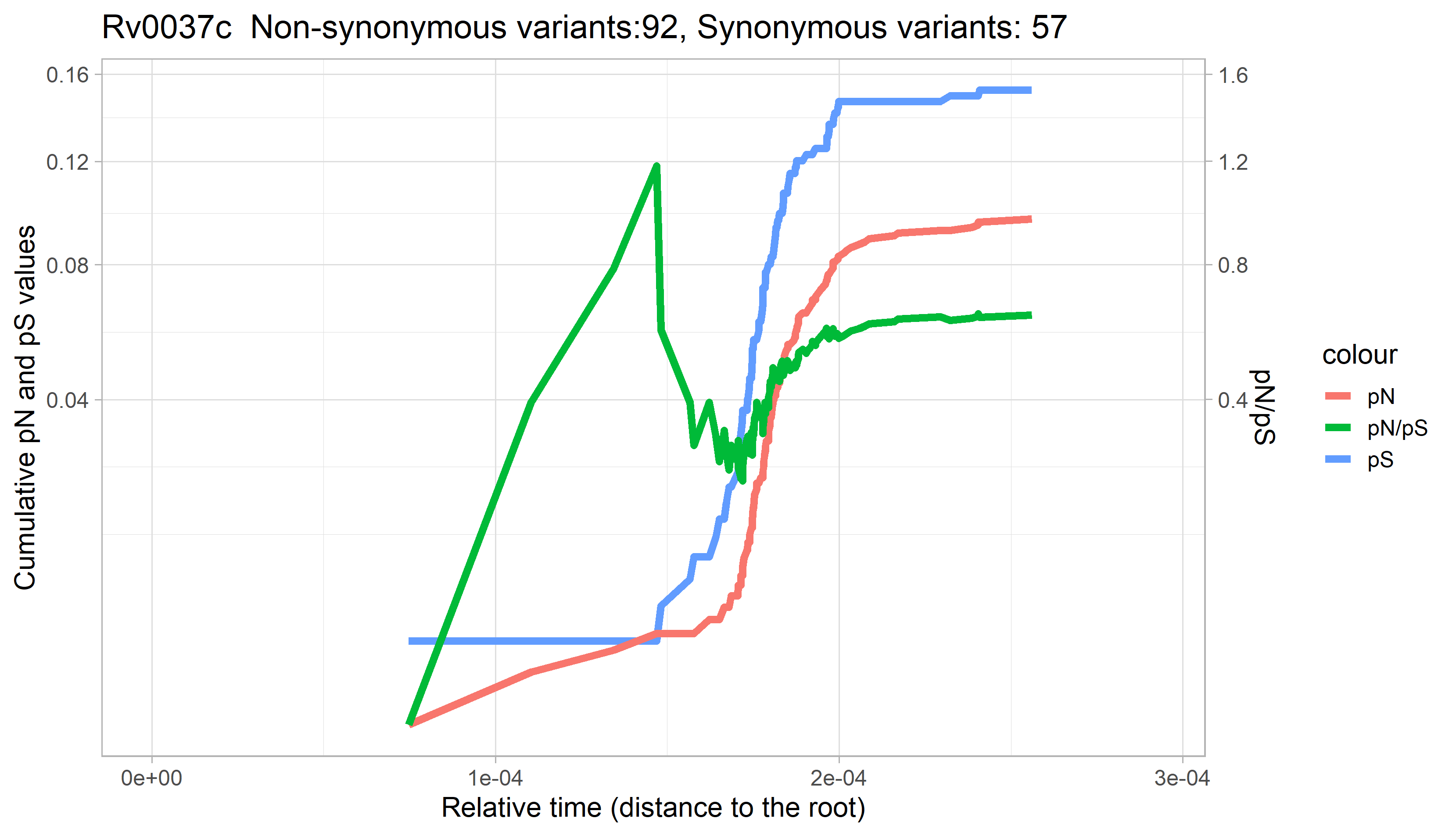

### Rv0038.png

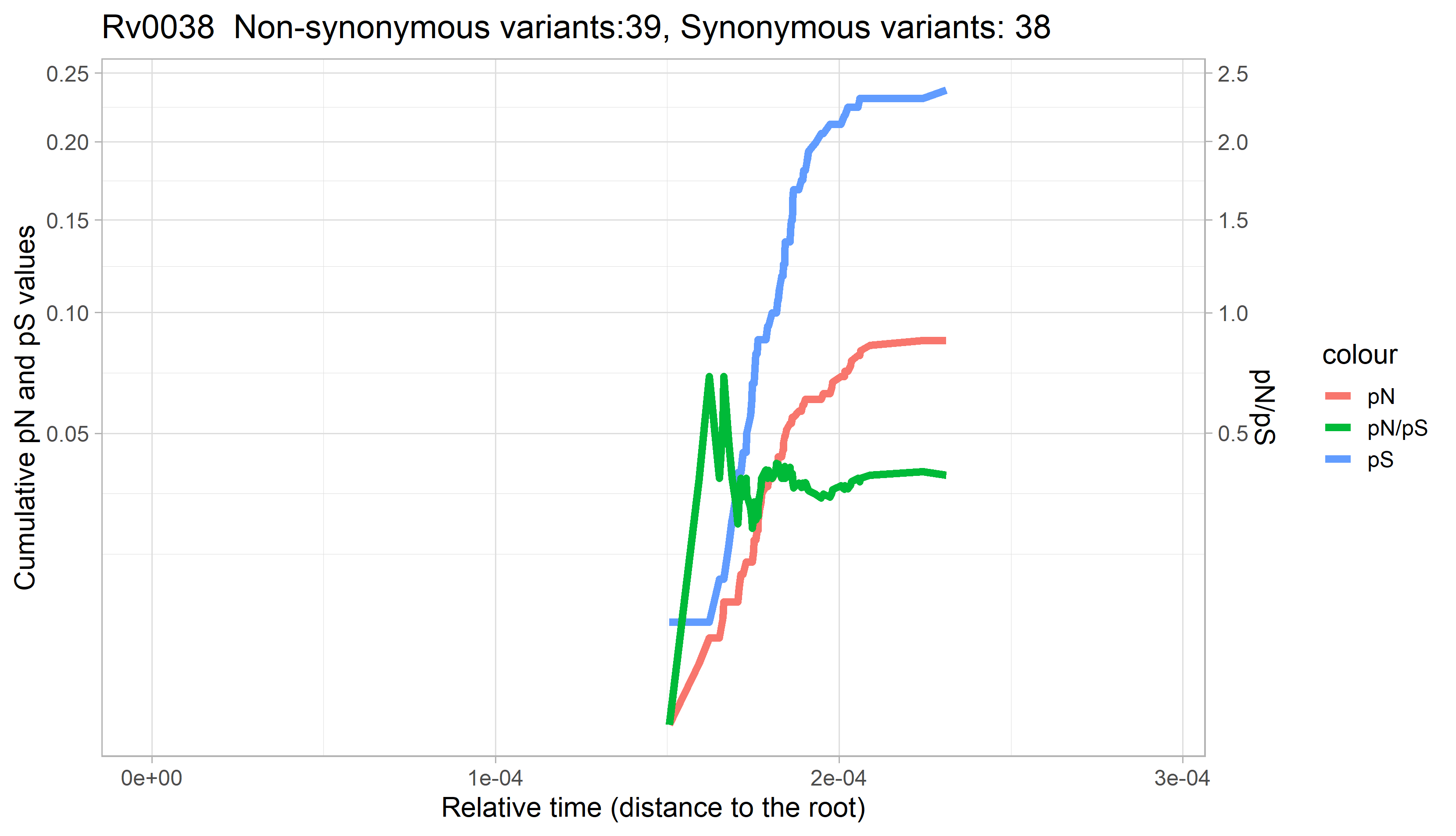

### Rv0039c.png

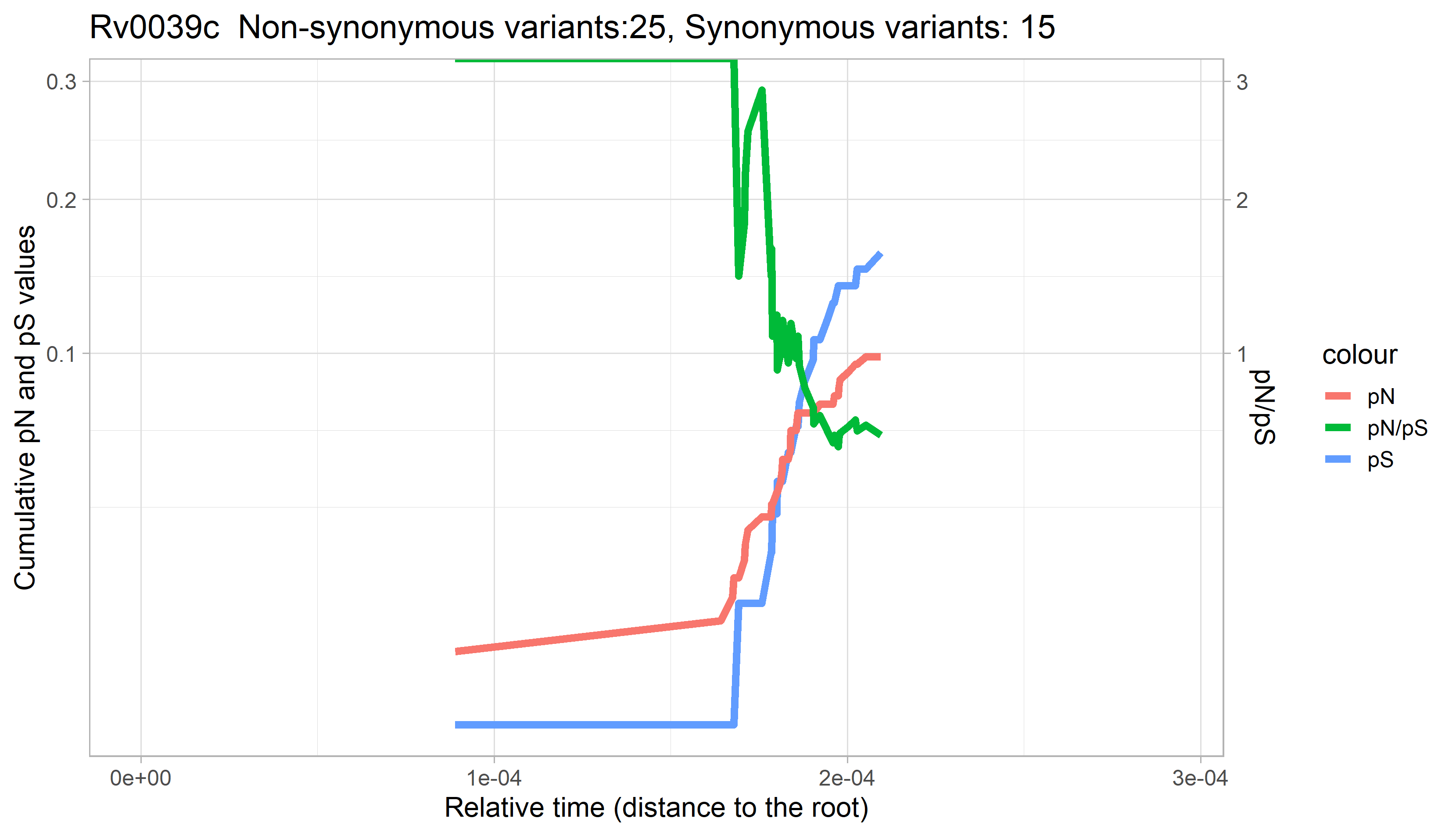

### Rv0040c.png

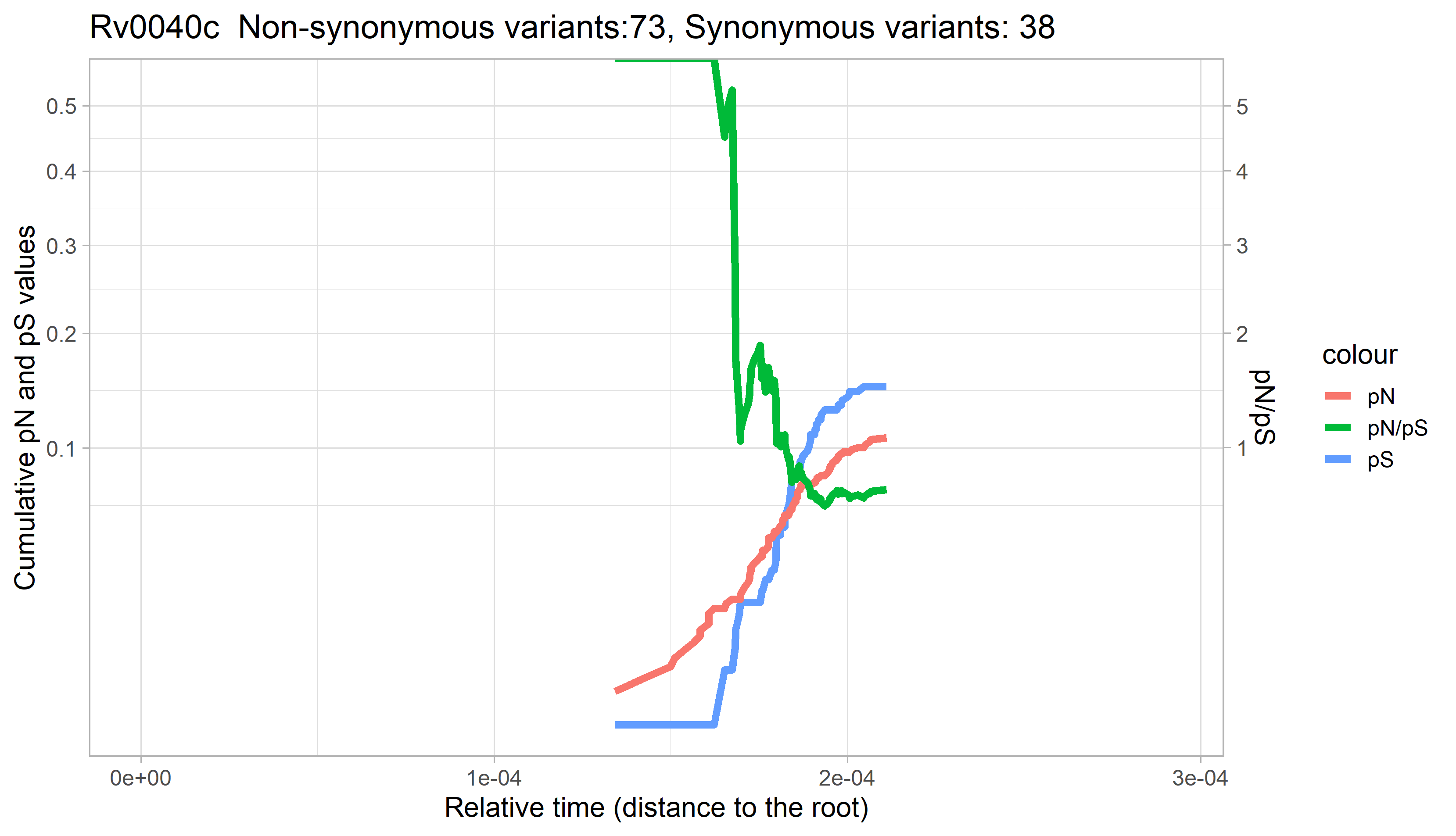

### Rv0041.png

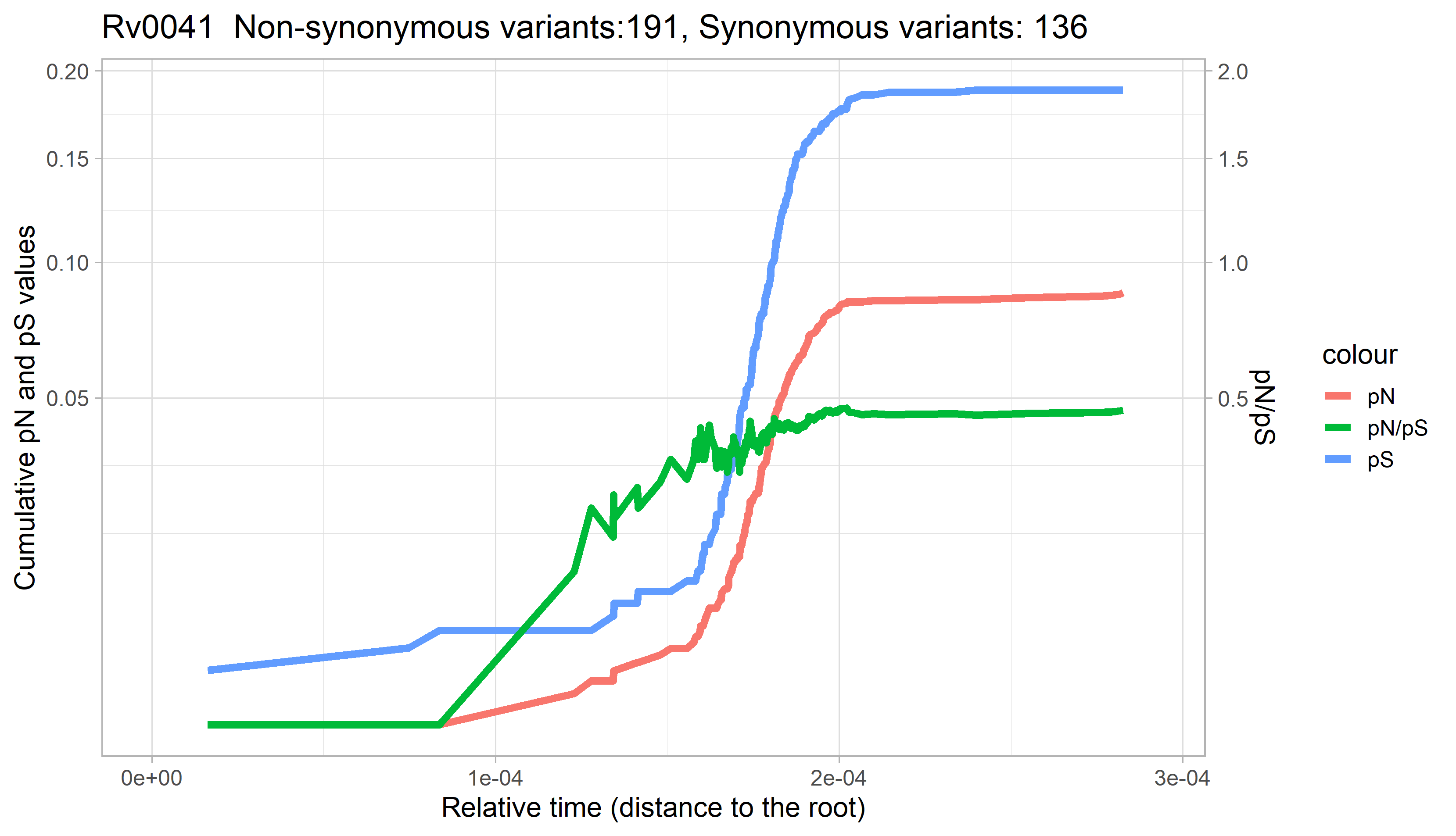

### Rv0042c.png

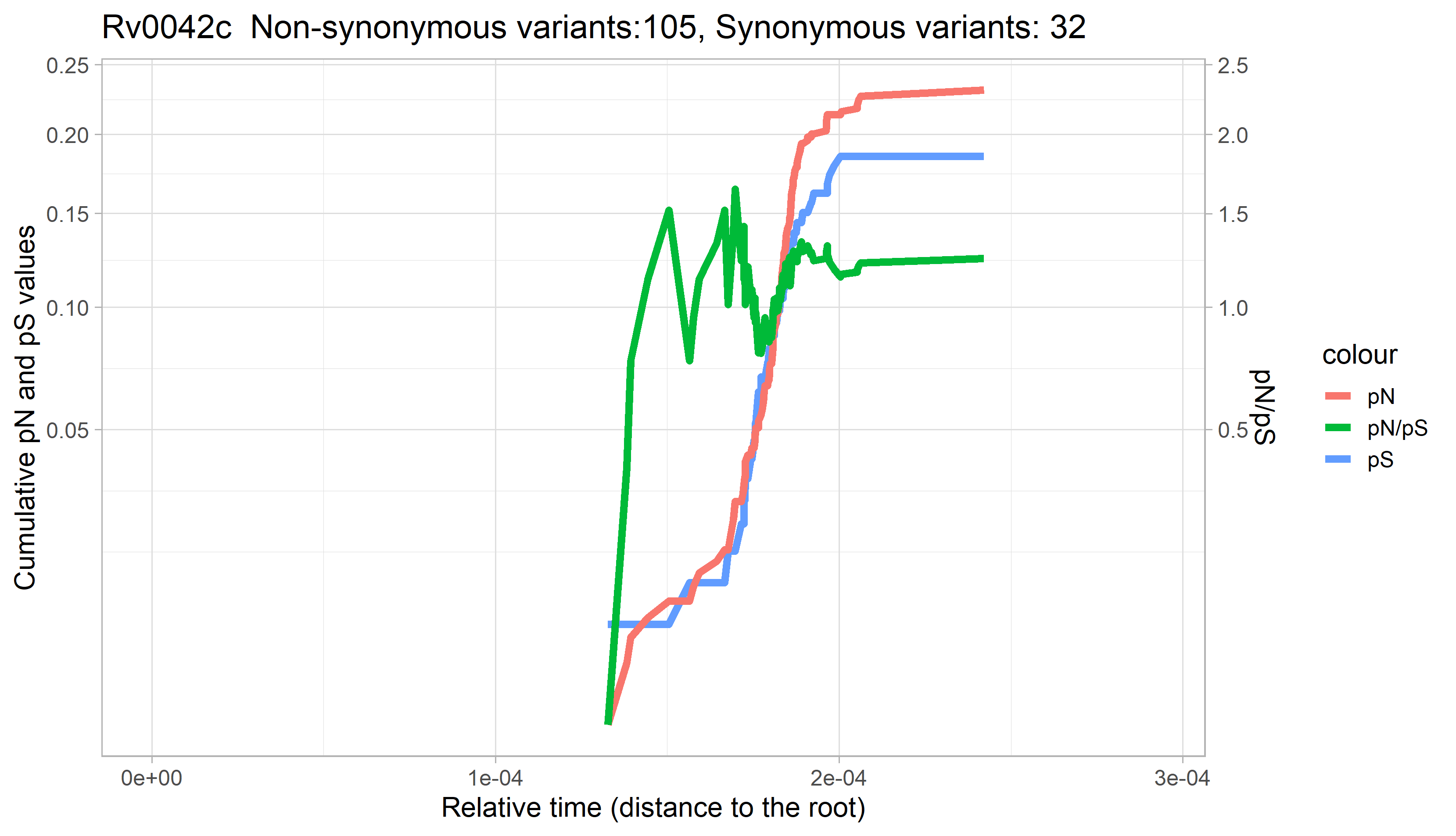

### Rv0043c.png

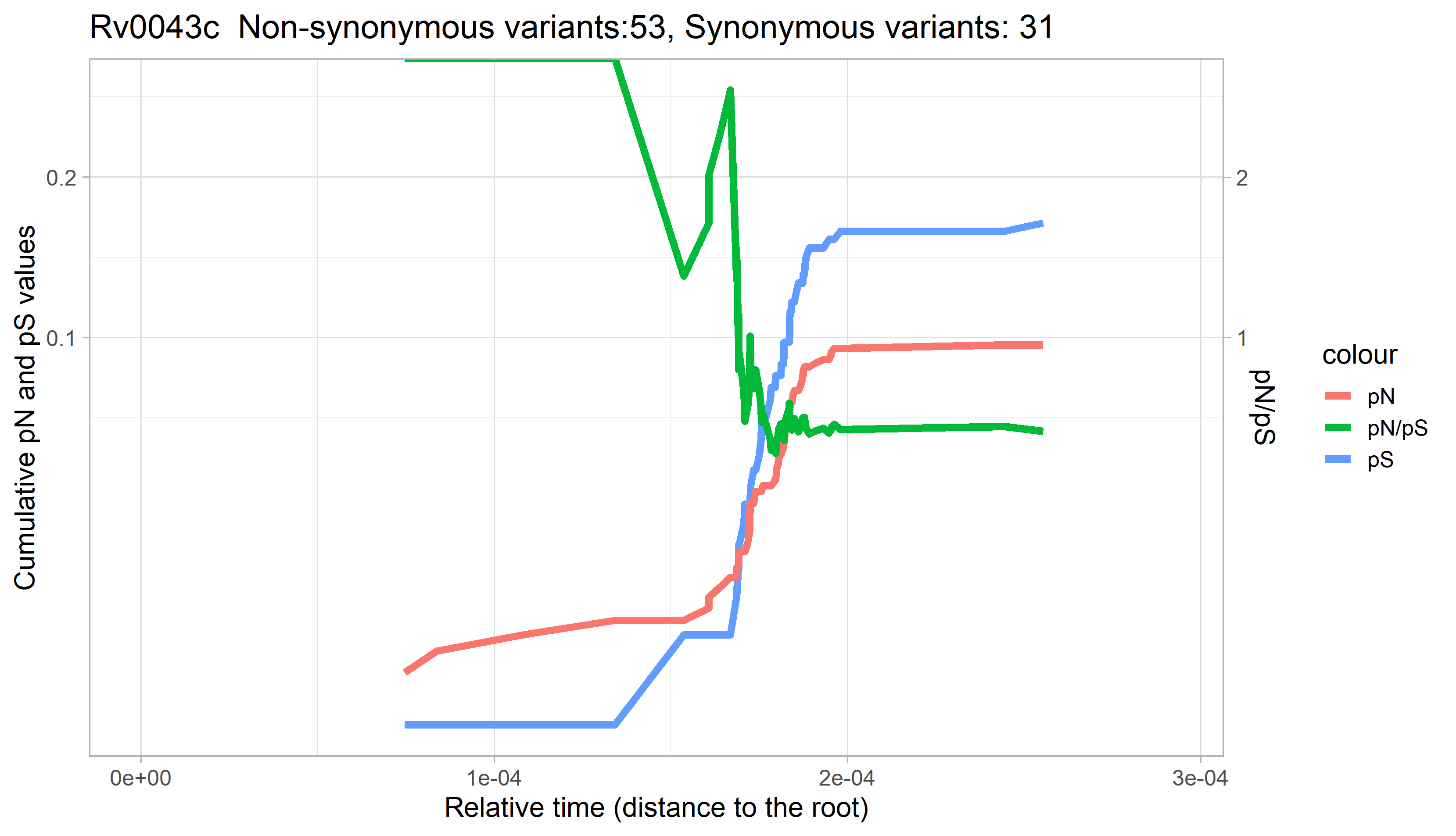

### Rv0044c.png

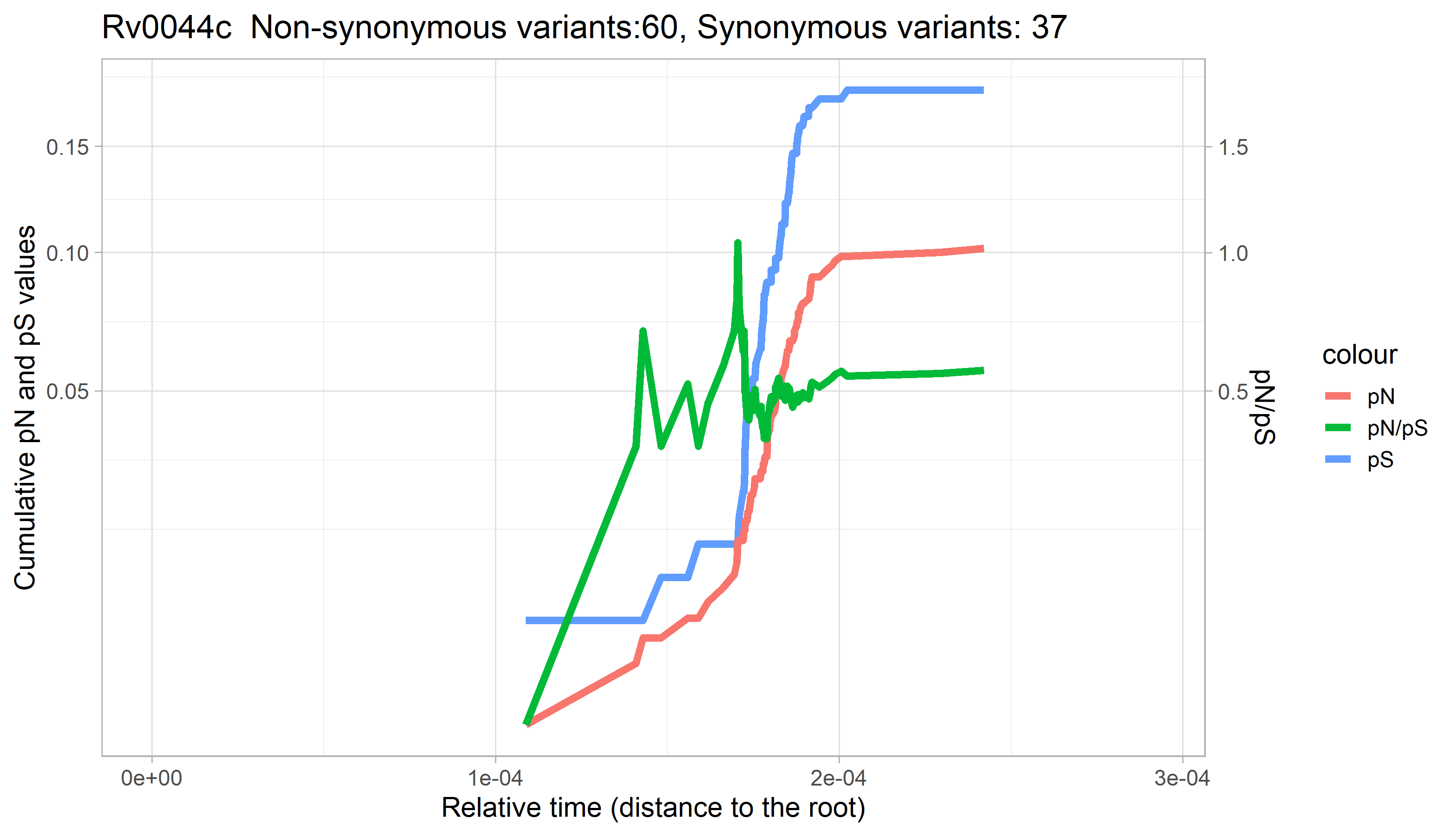

### Rv0045c.png

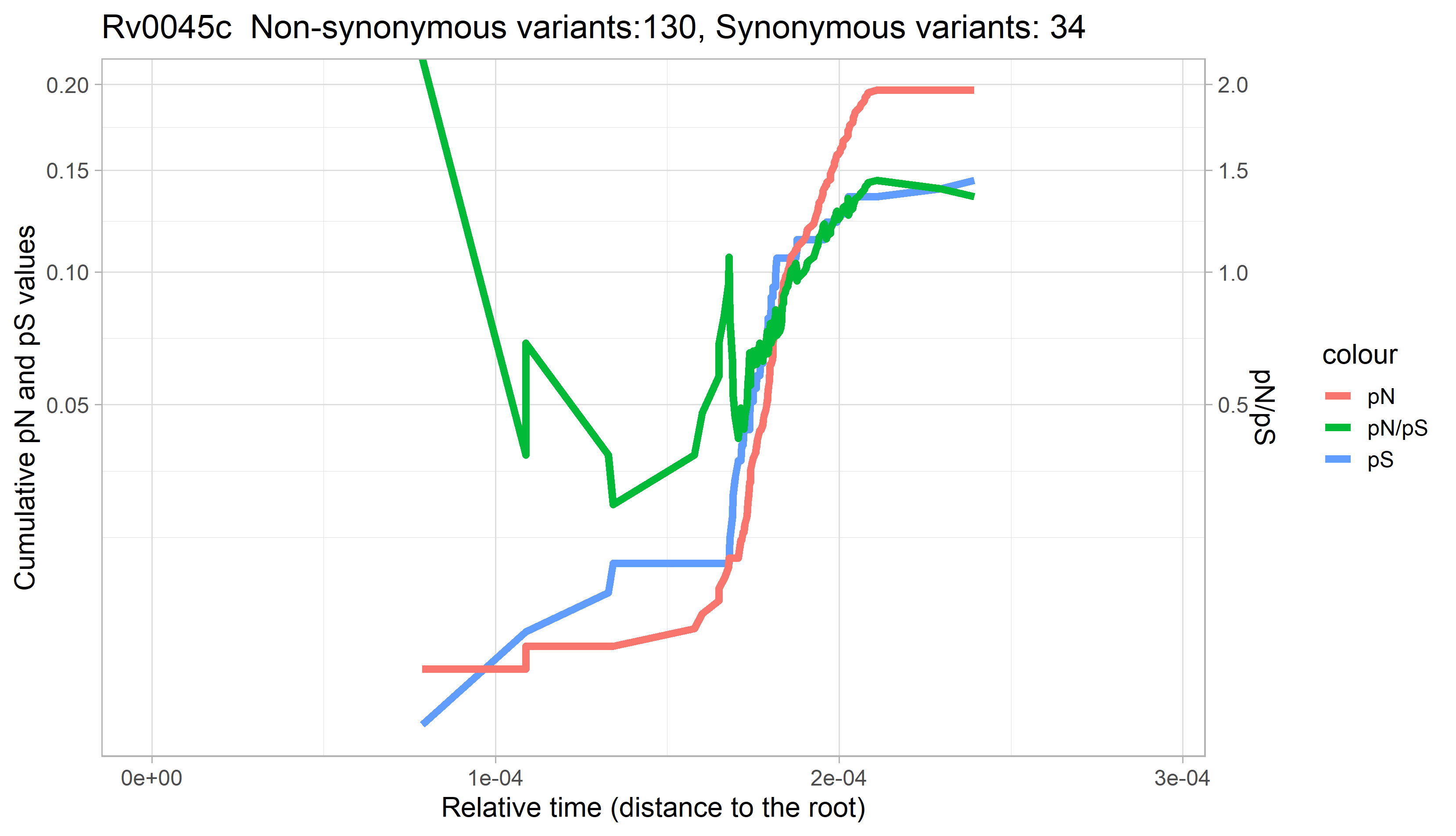

### Rv0046c.png

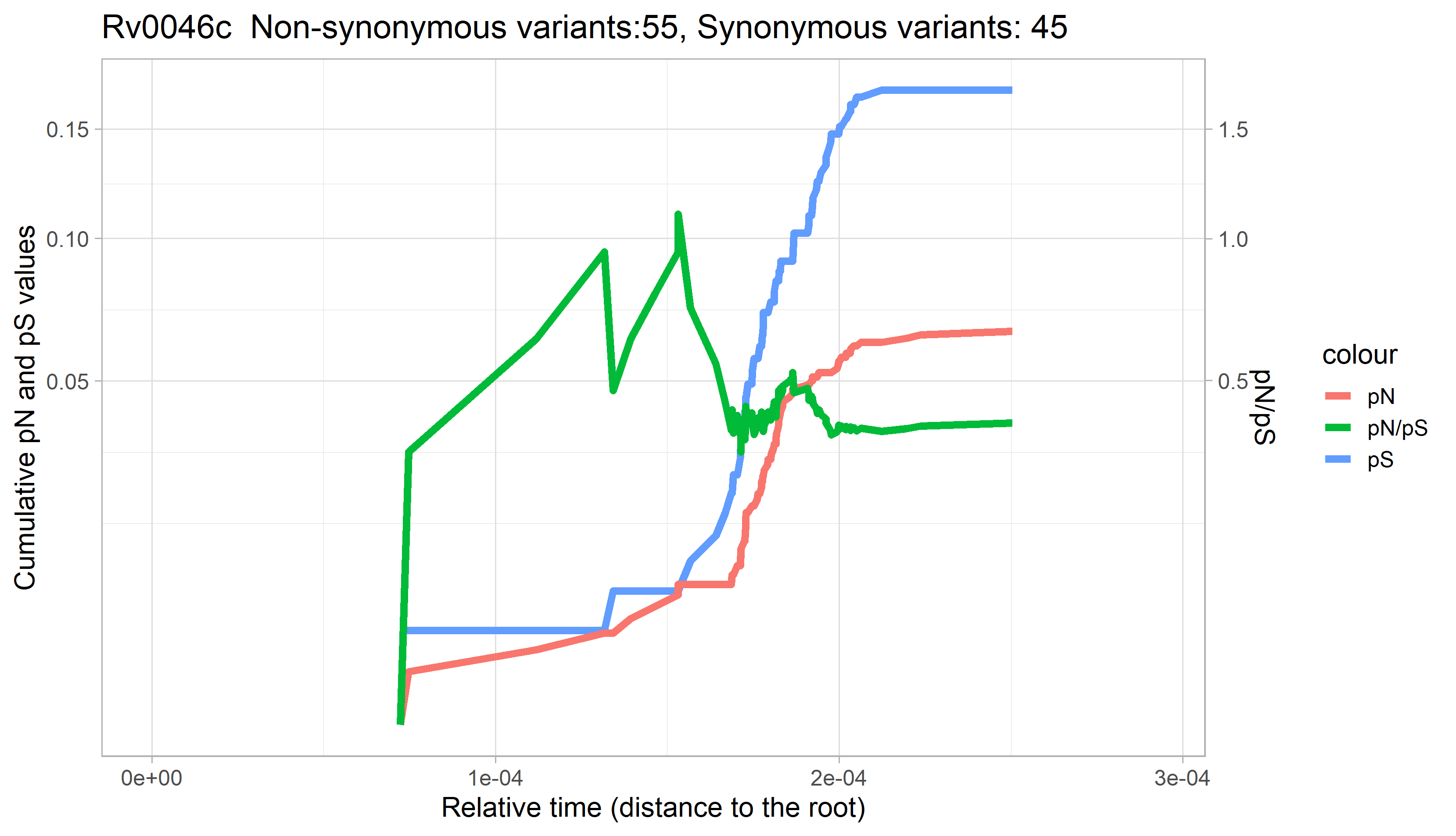

### Rv0047c.png

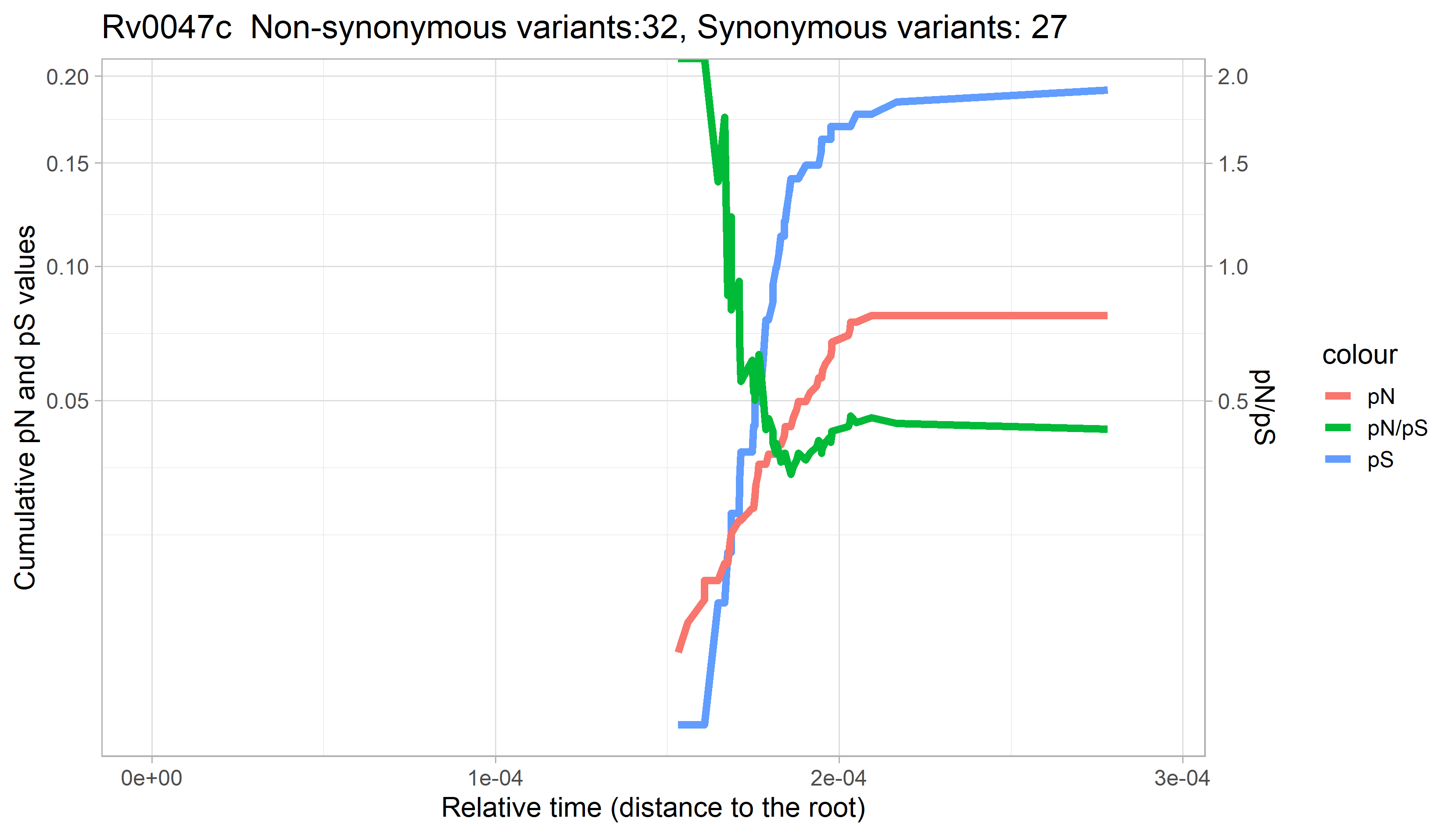

### Rv0048c.png

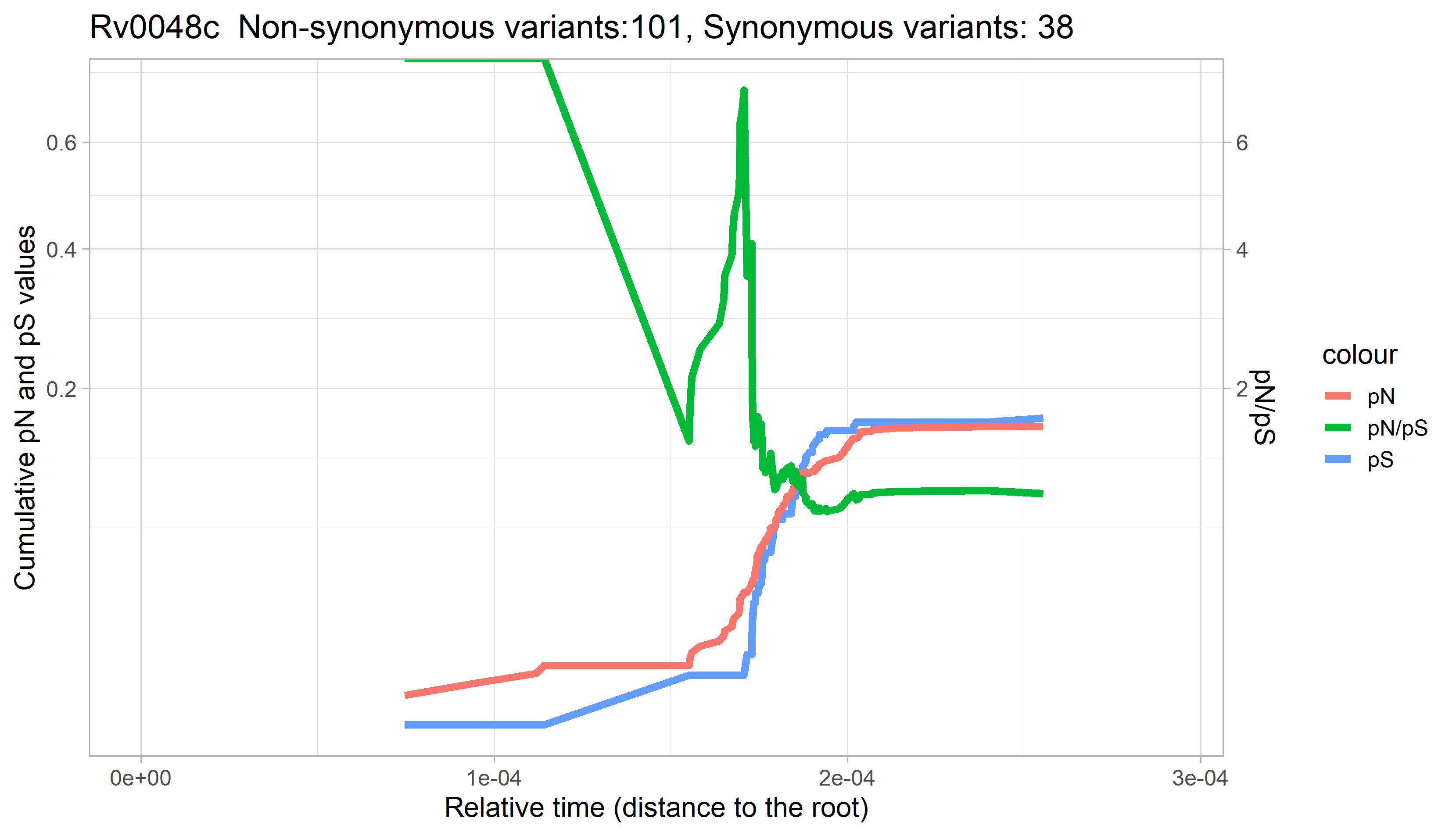

### Rv0049.png

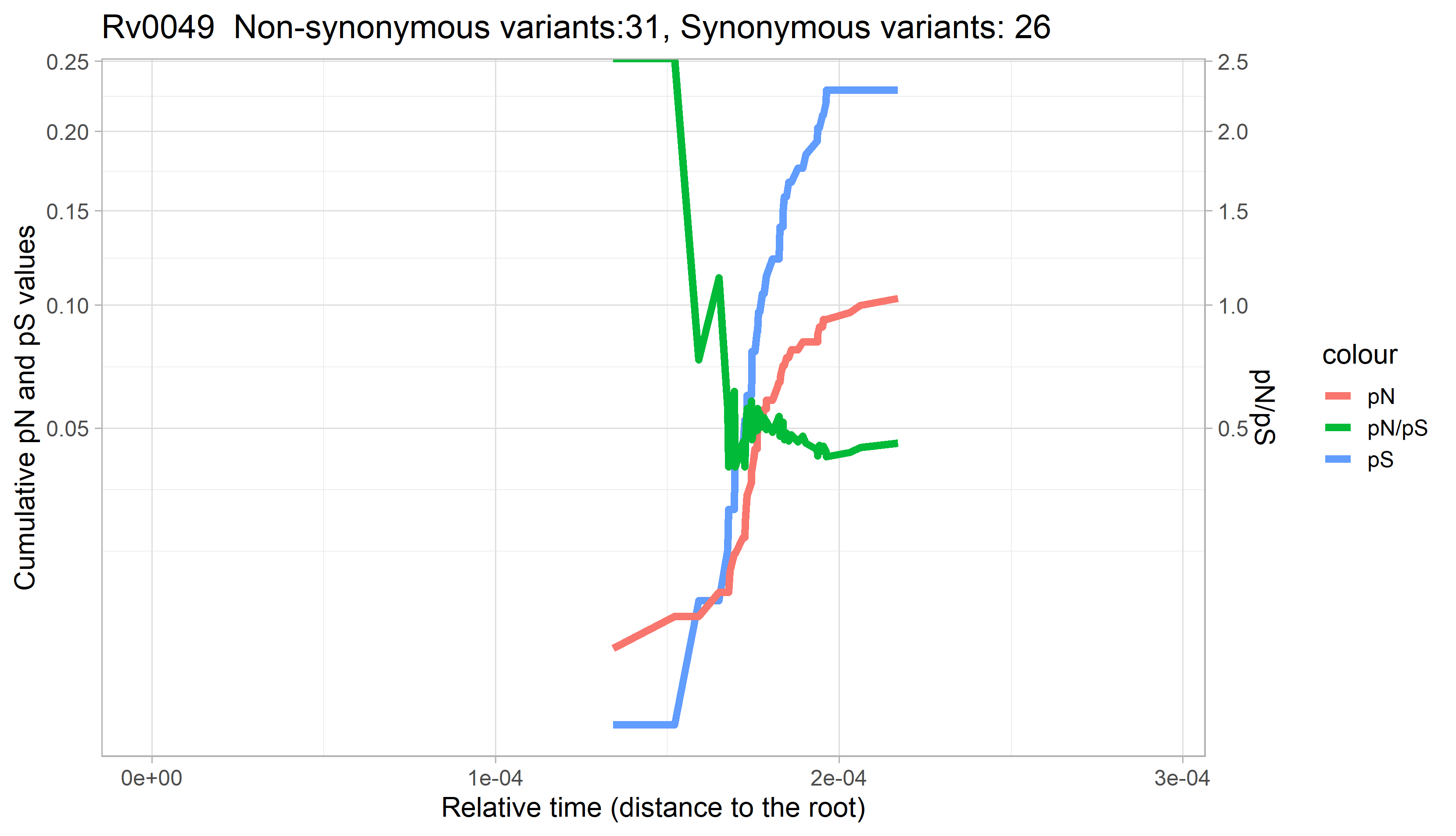

### Rv0050.png

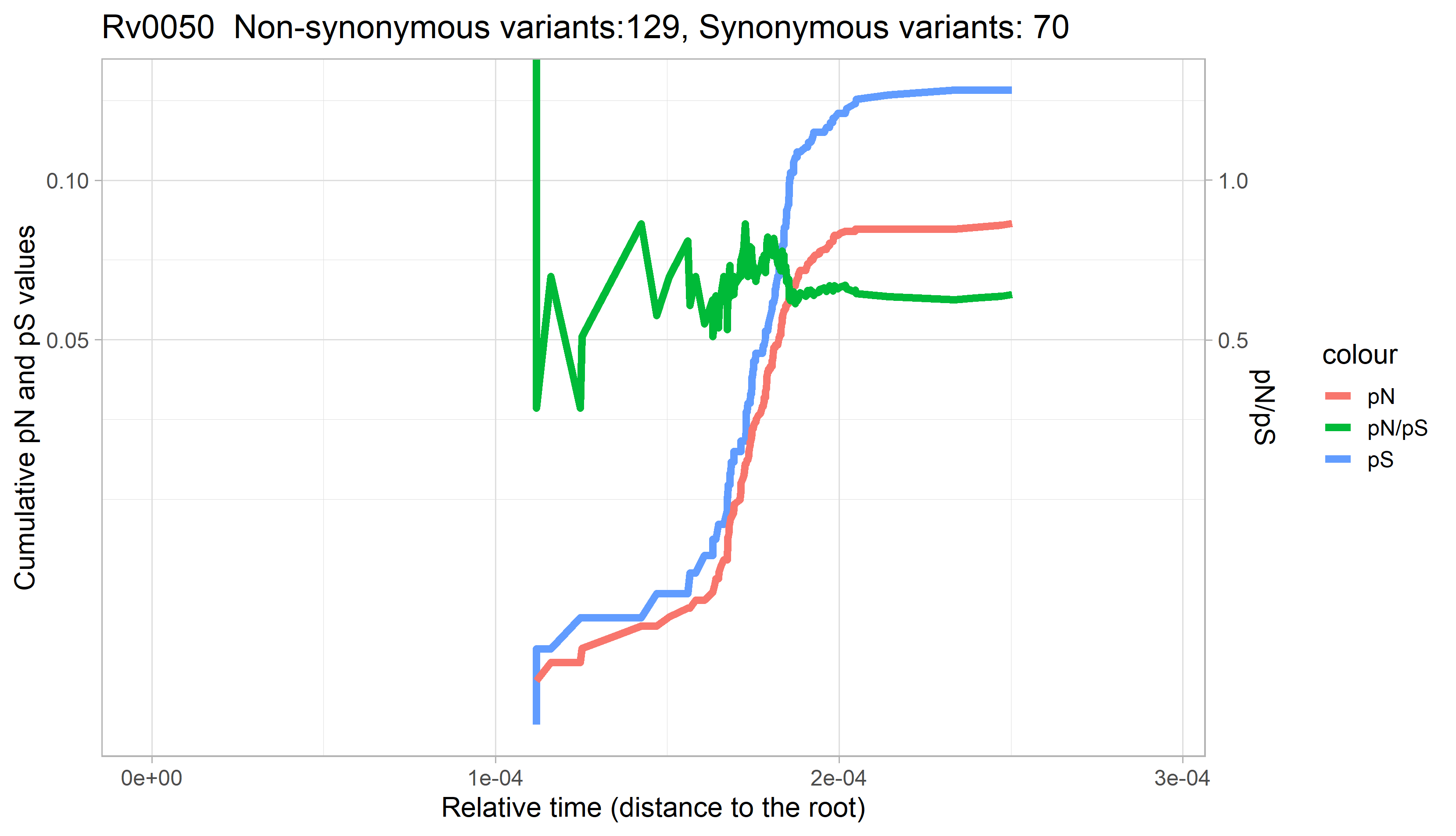

### Rv0051.png

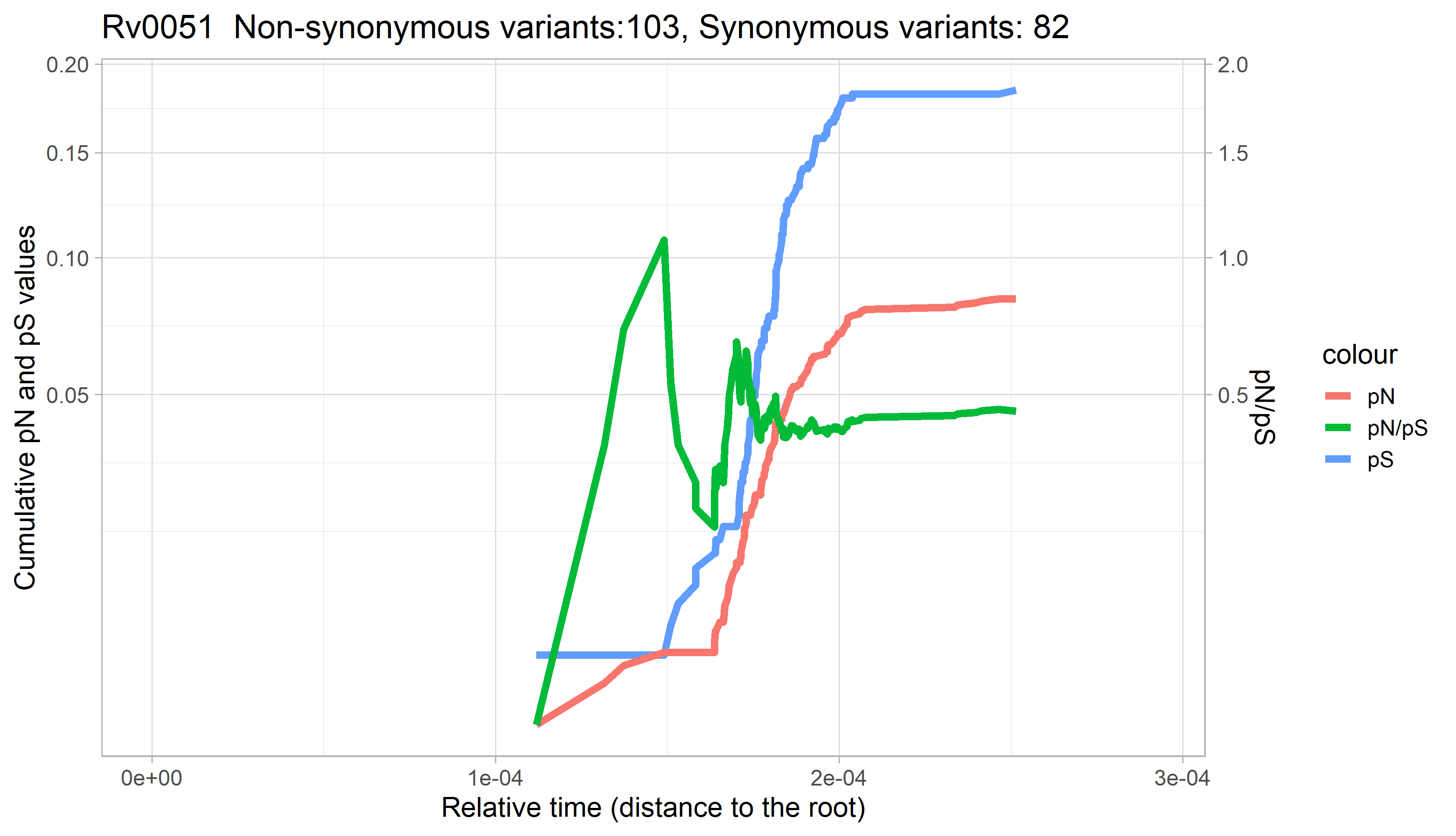

### Rv0052.png

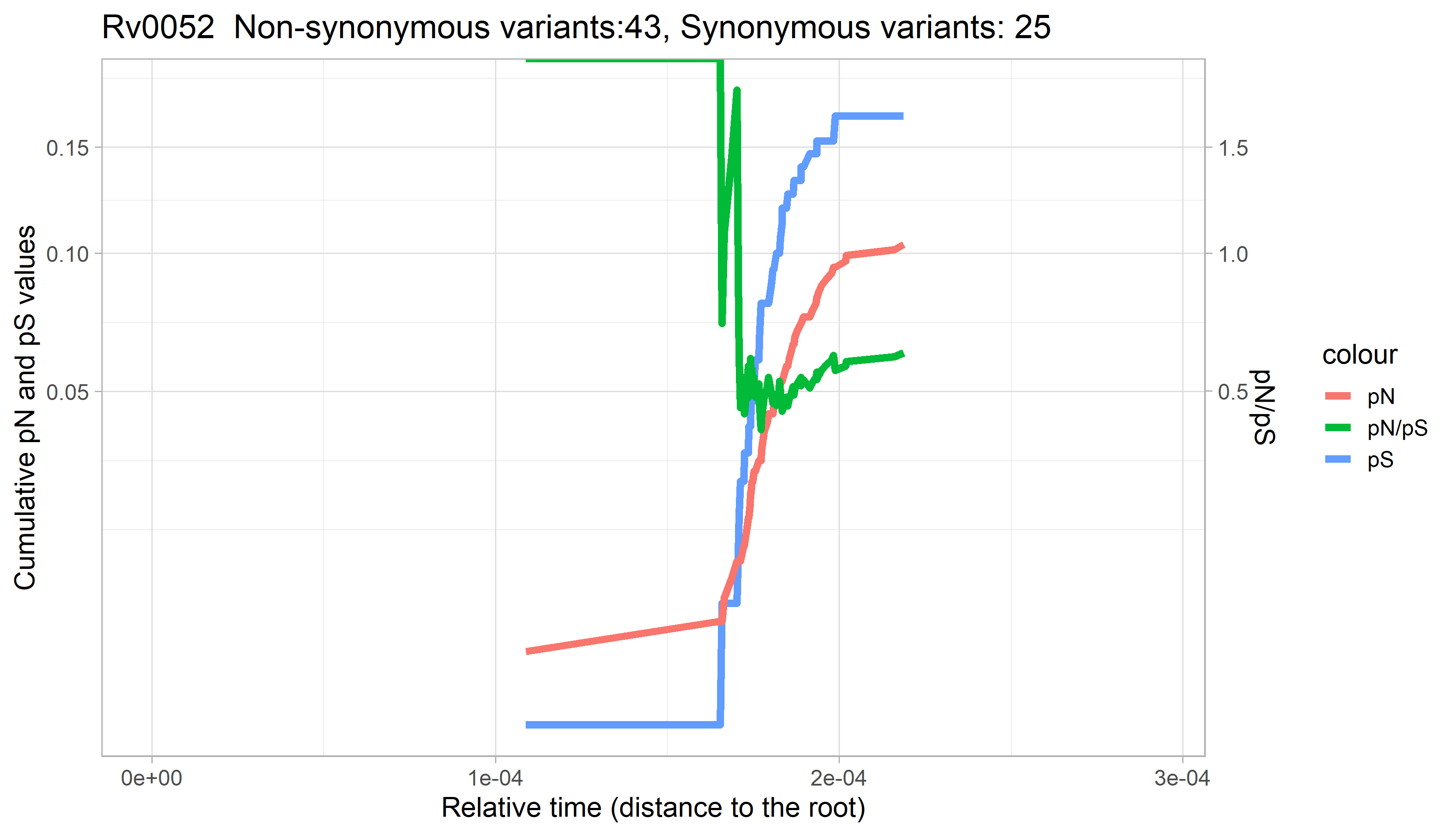

### Rv0053.png

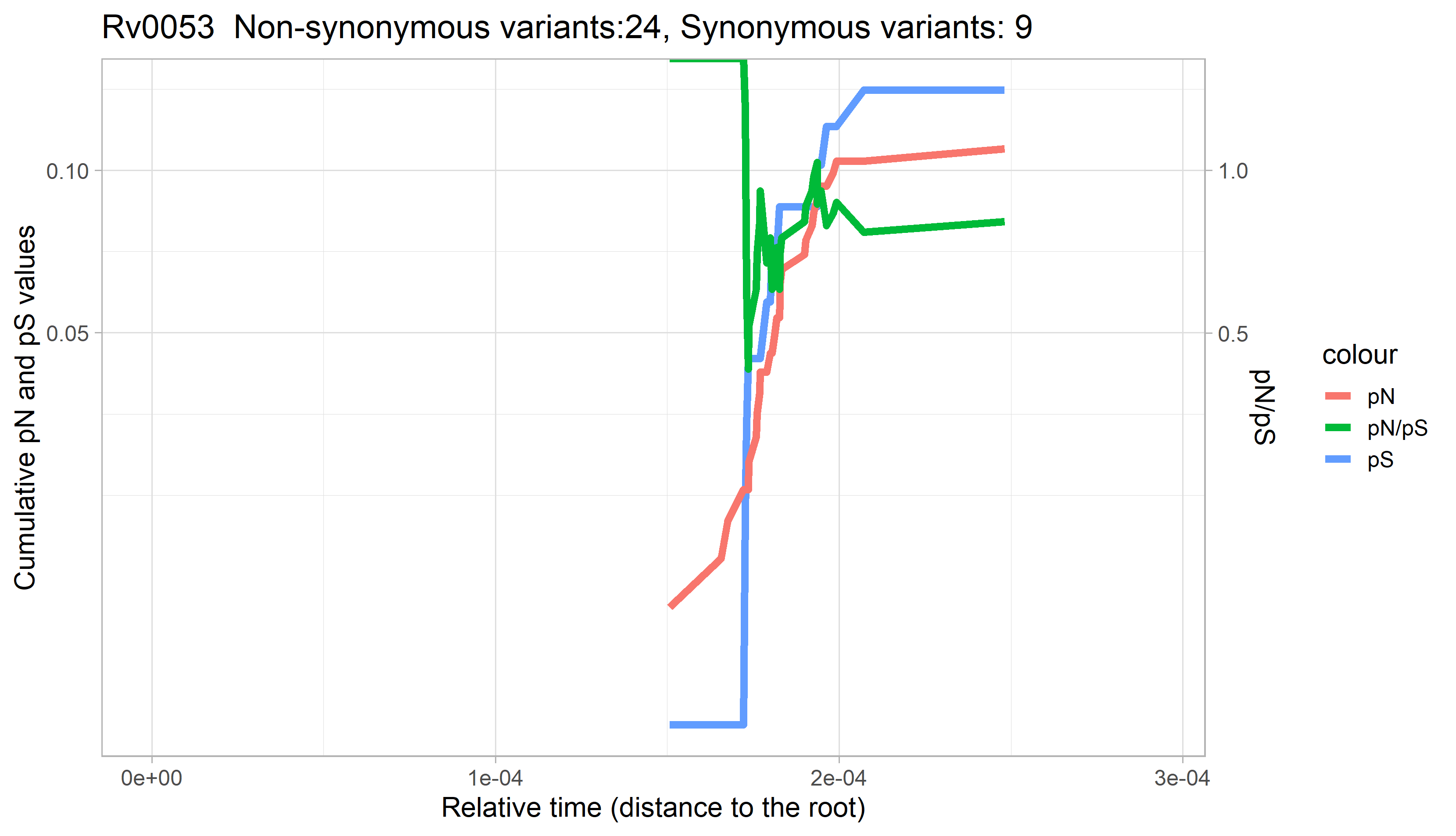

### Rv0054.png

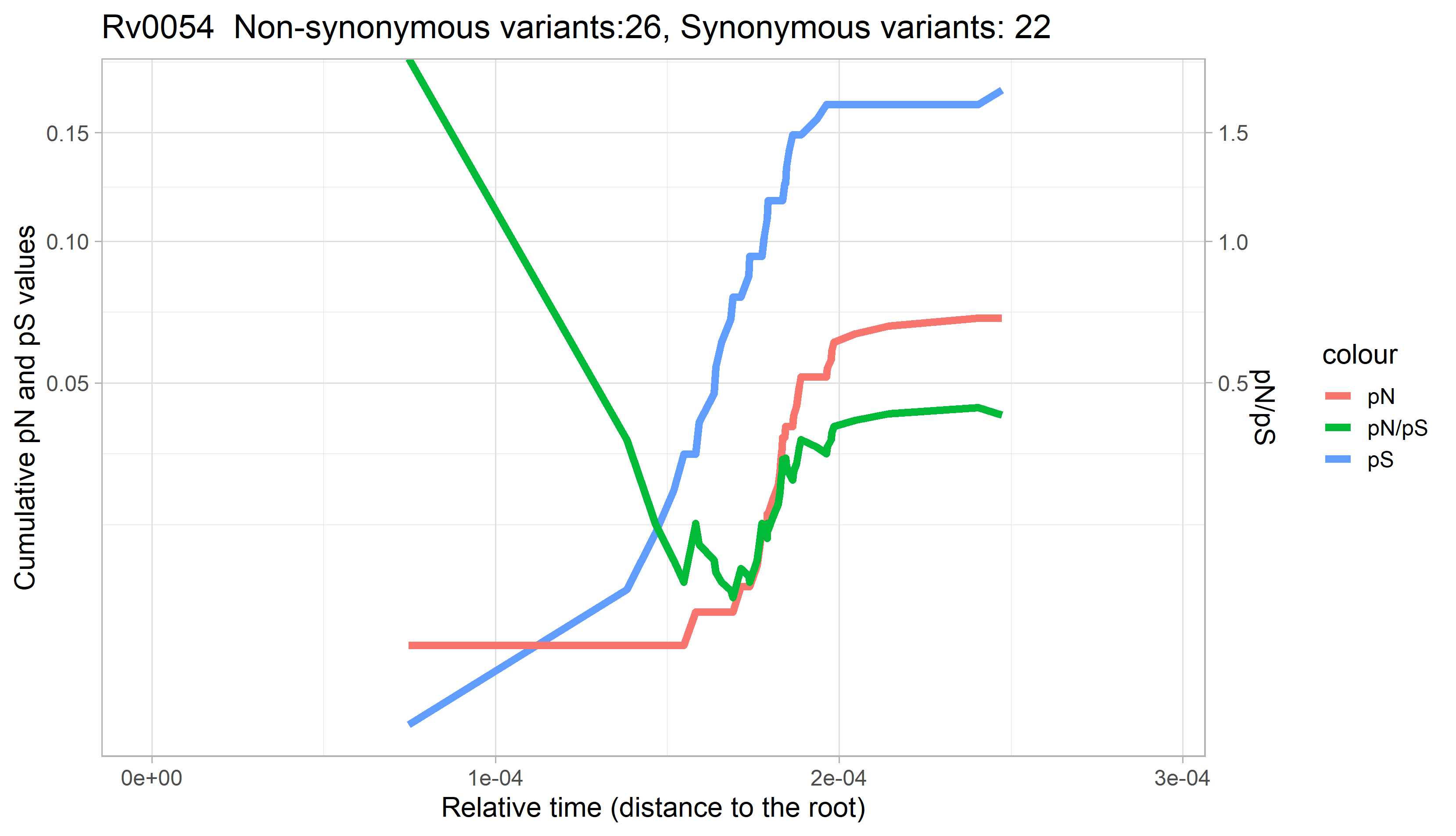

### Rv0055.png

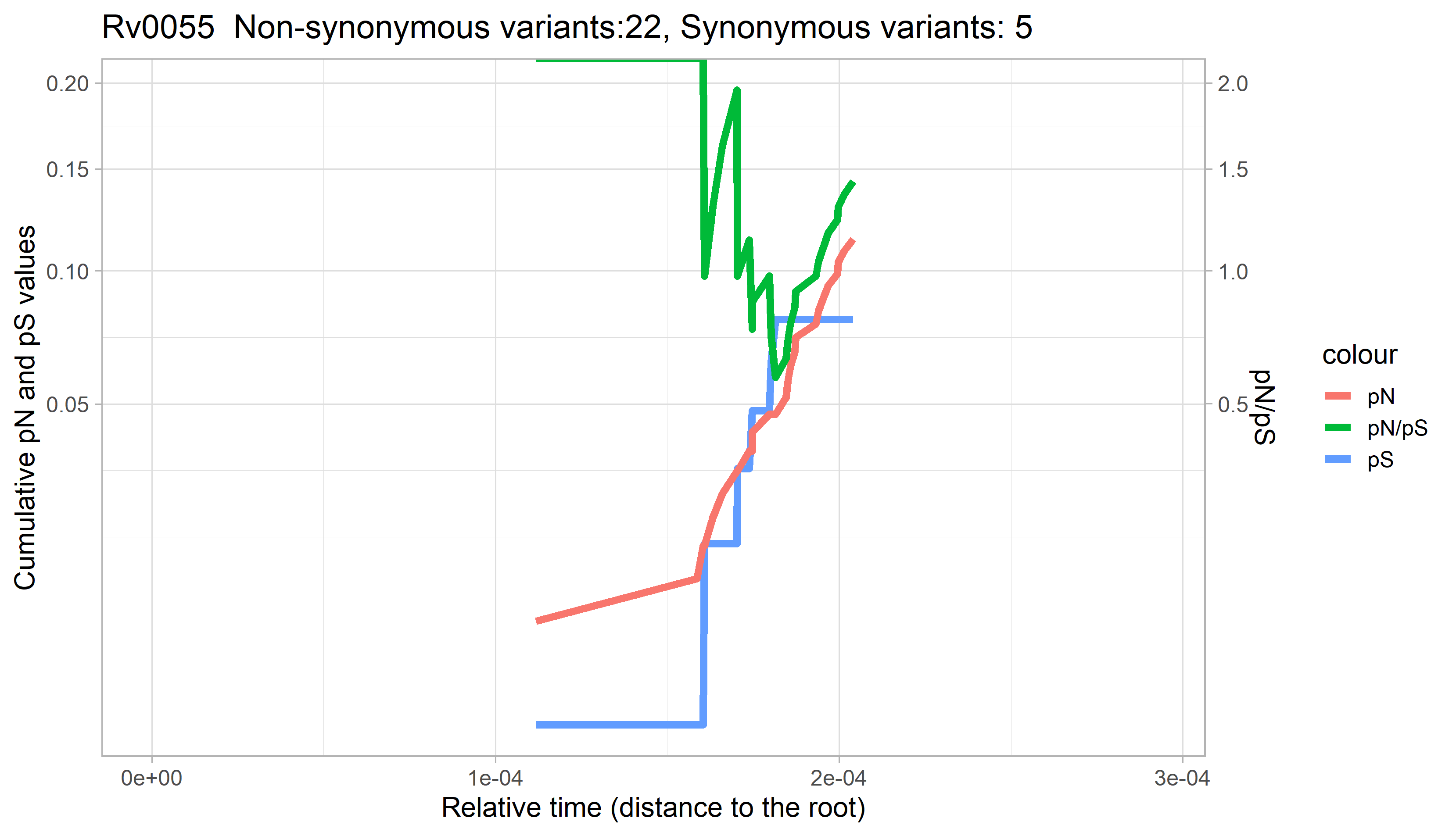

### Rv0056.png

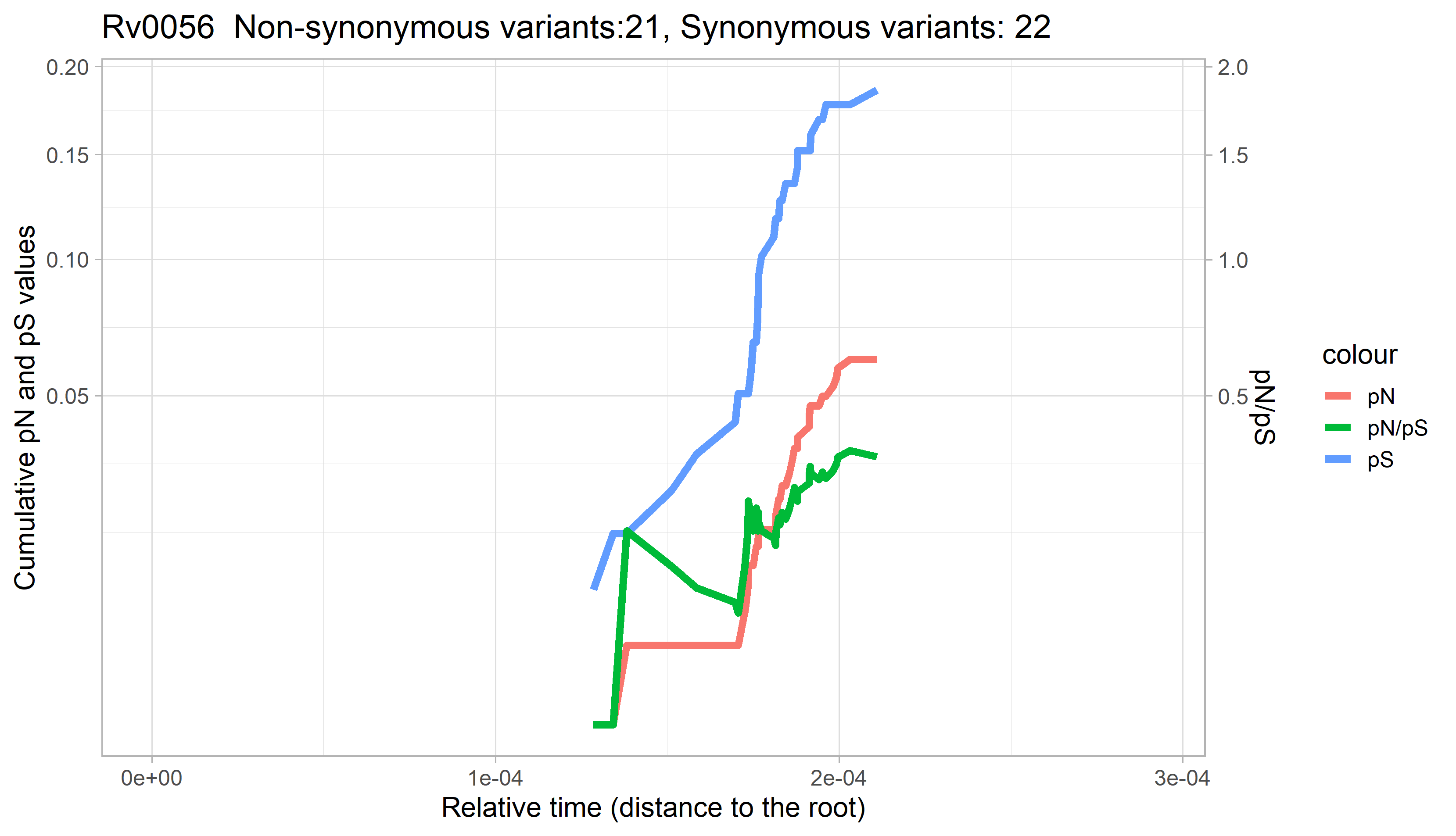

### Rv0057.png

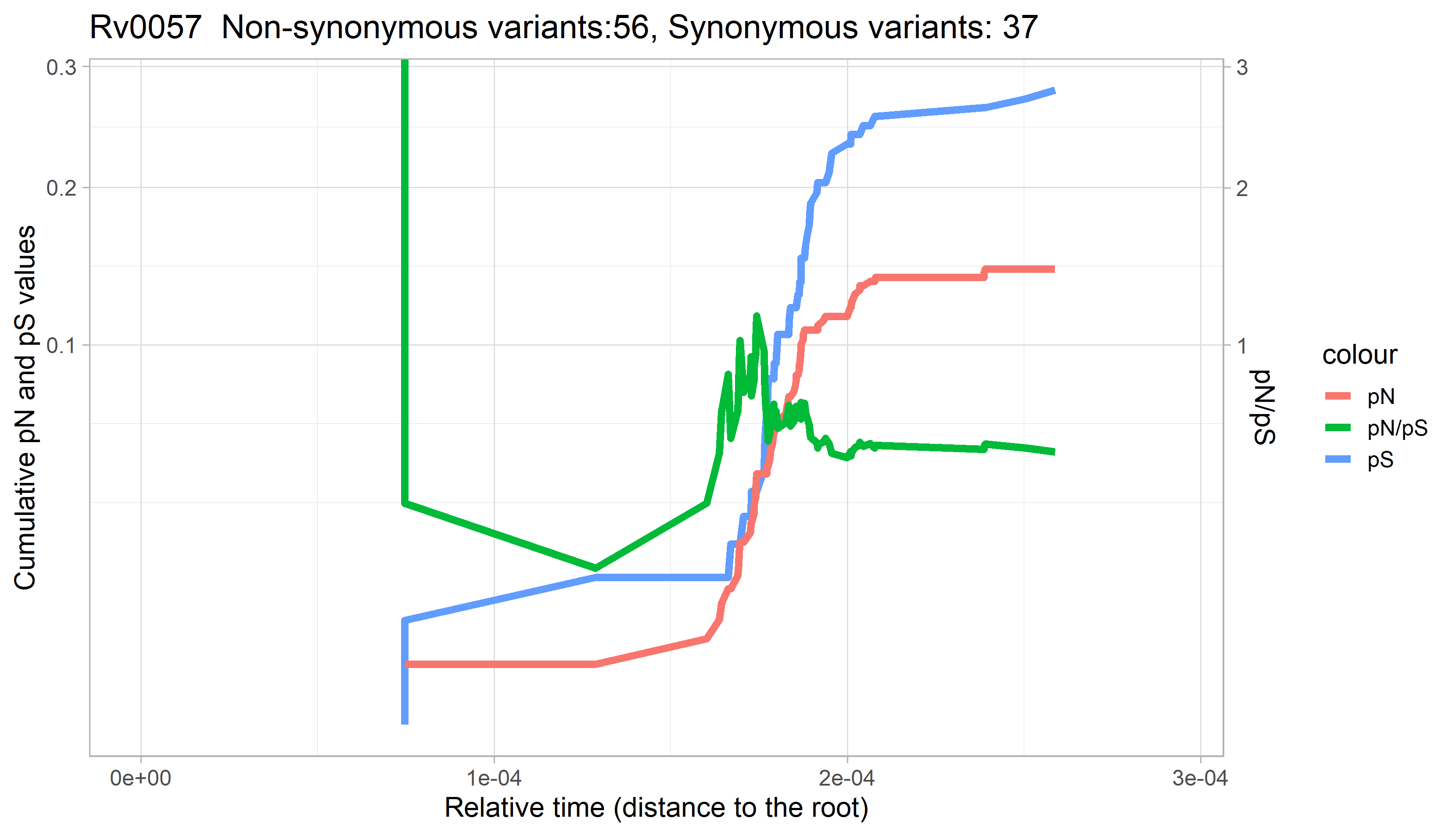

### Rv0058.png

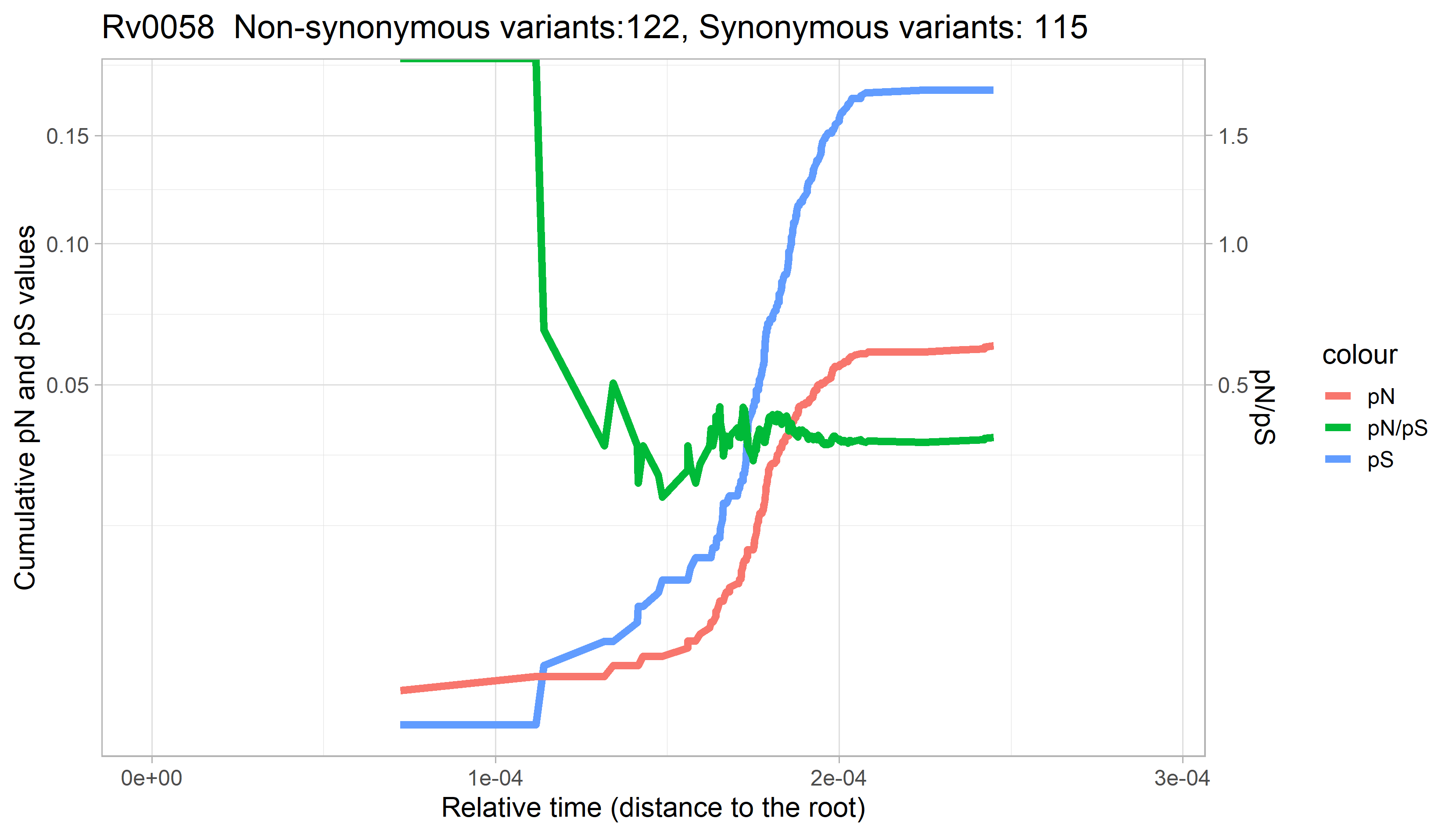

### Rv0059.png

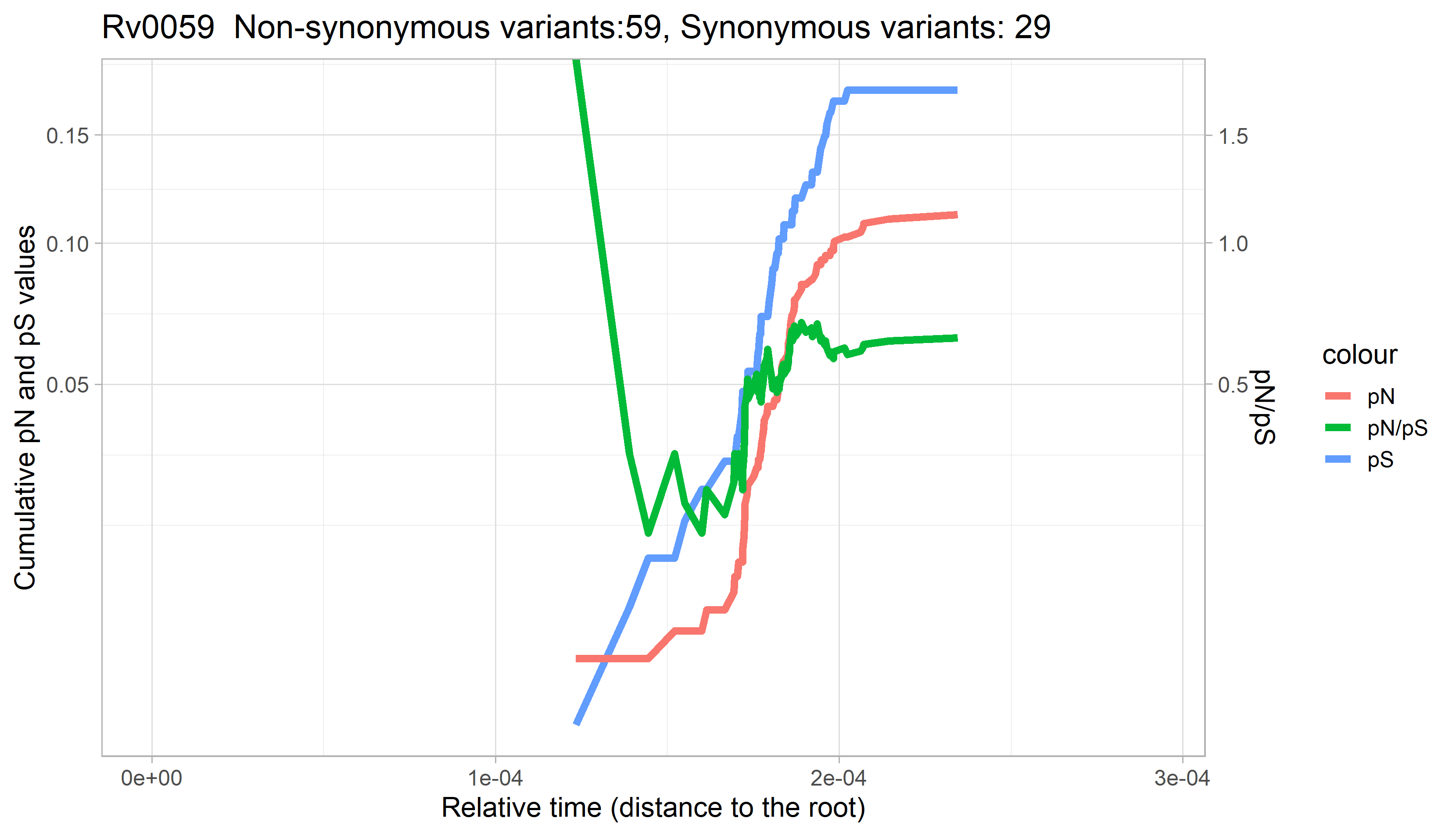

### Rv0060.png

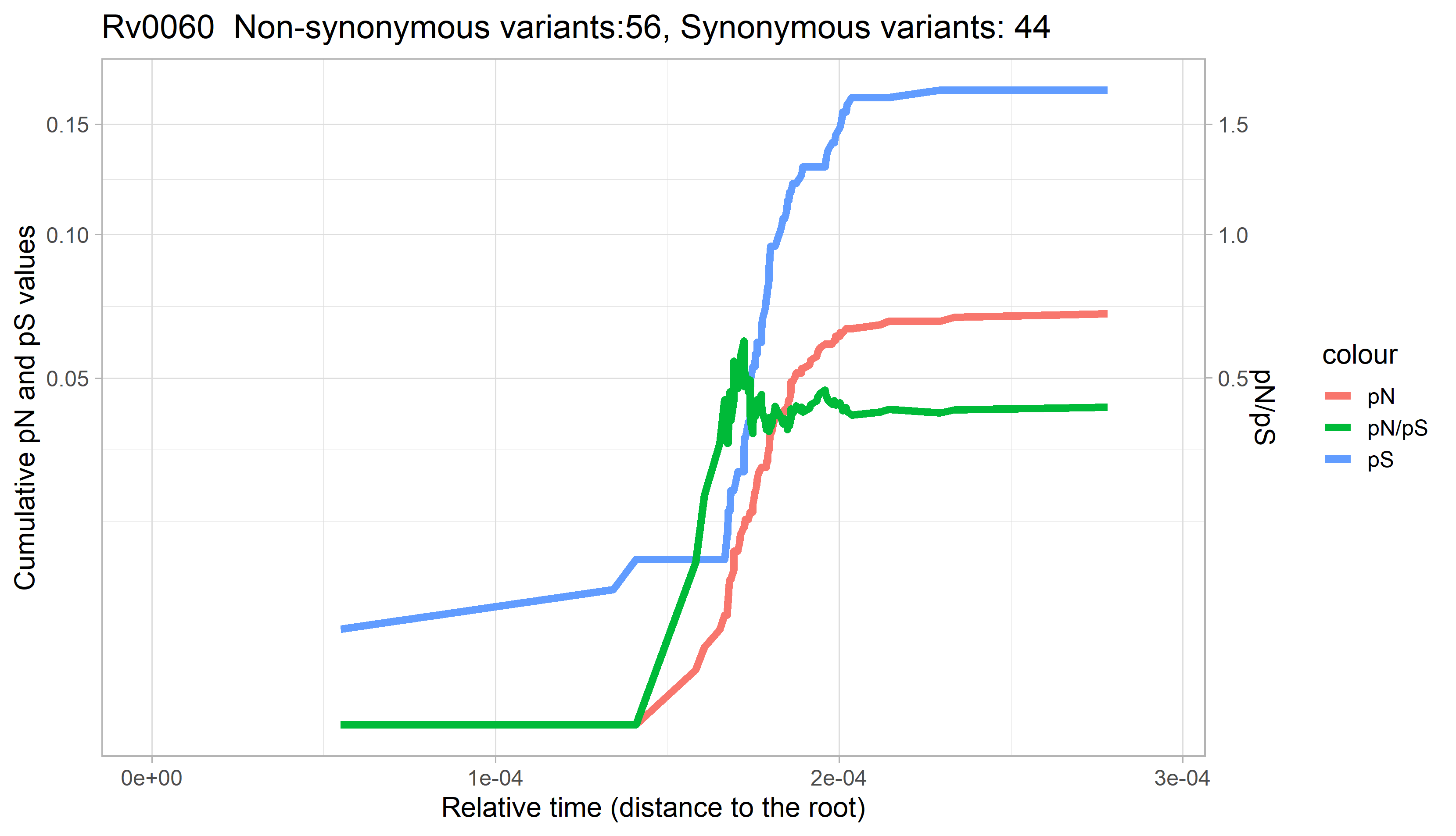

### Rv0061.png

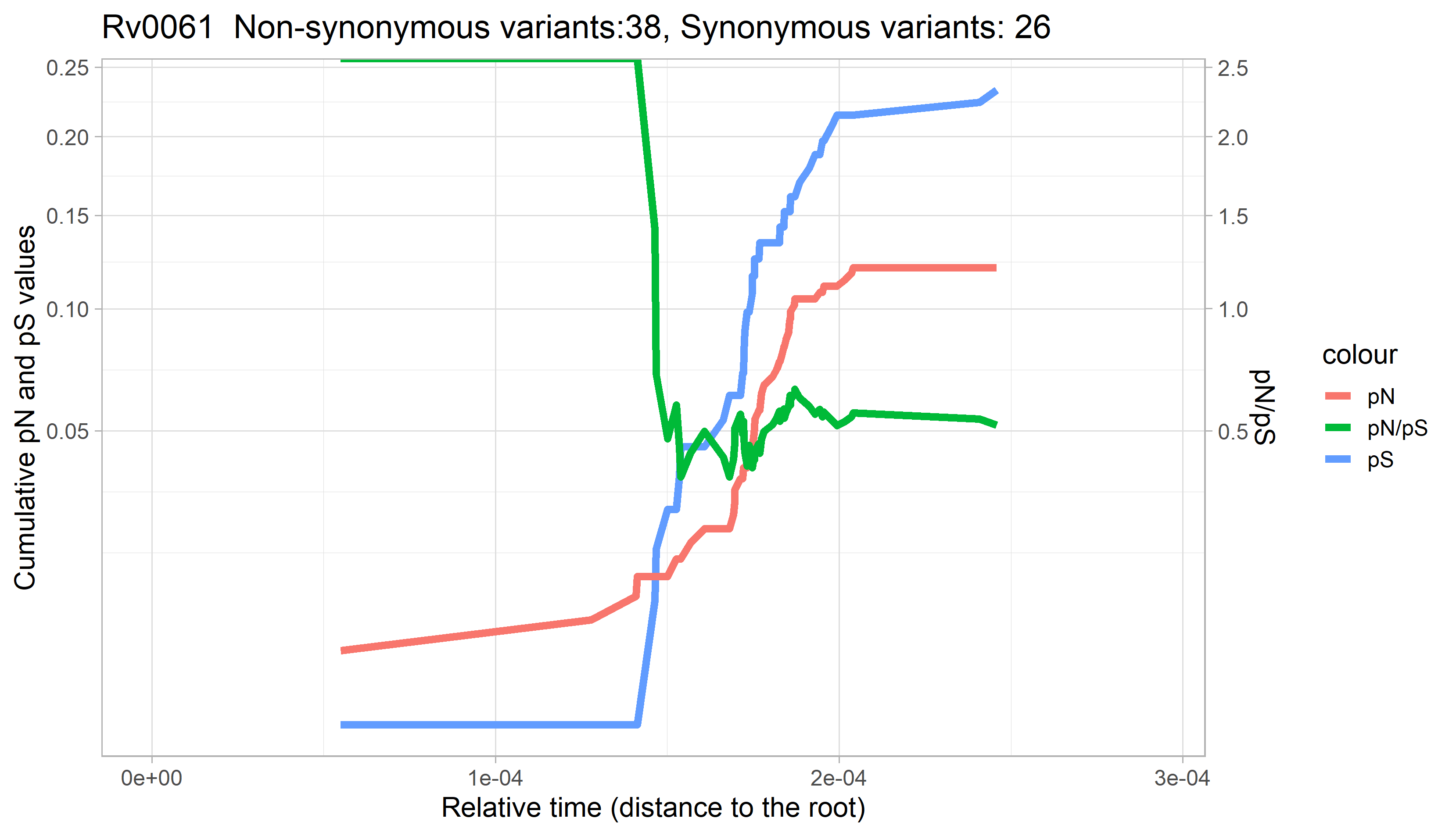

### Rv0062.png

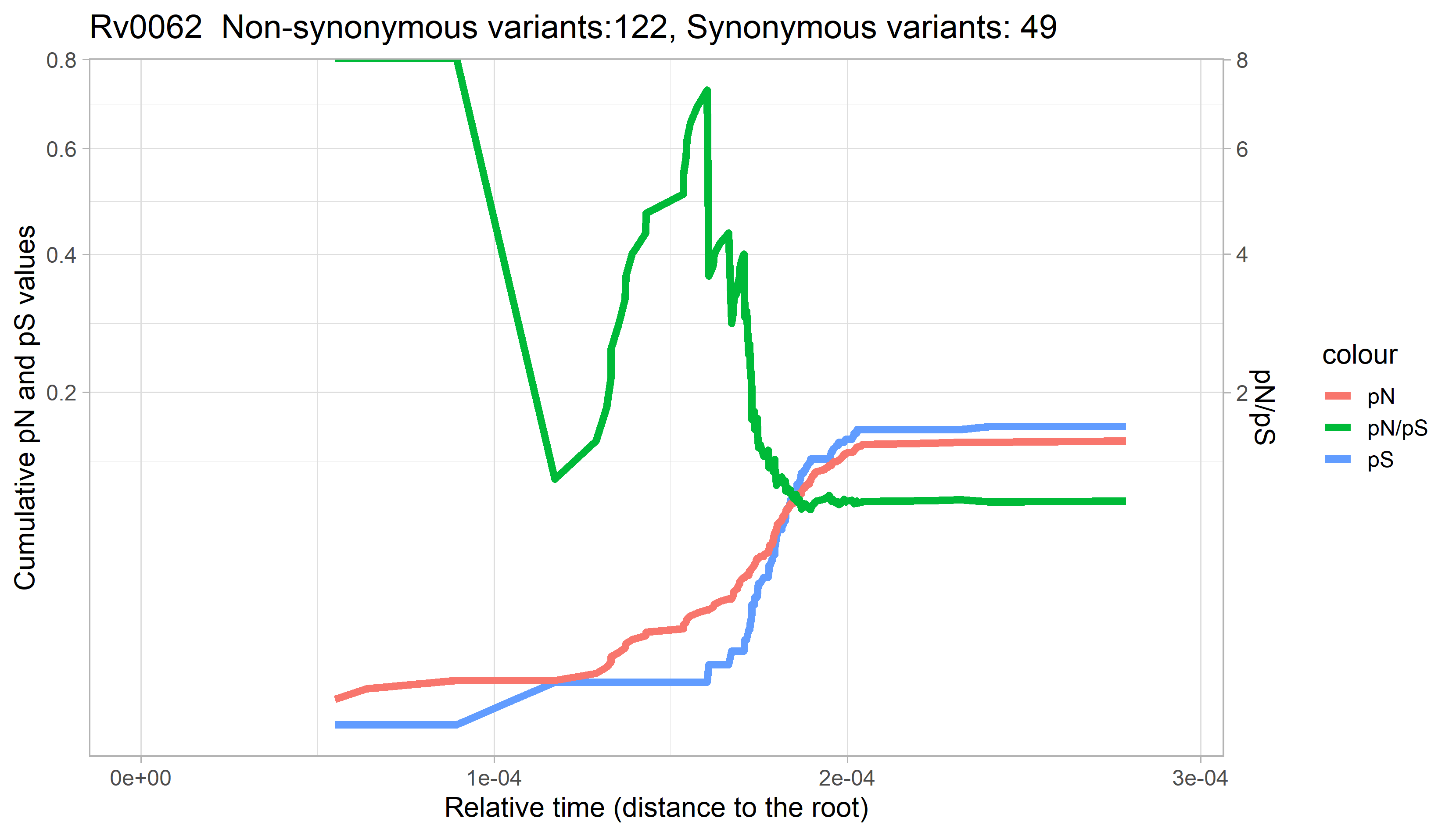

### Rv0063.png

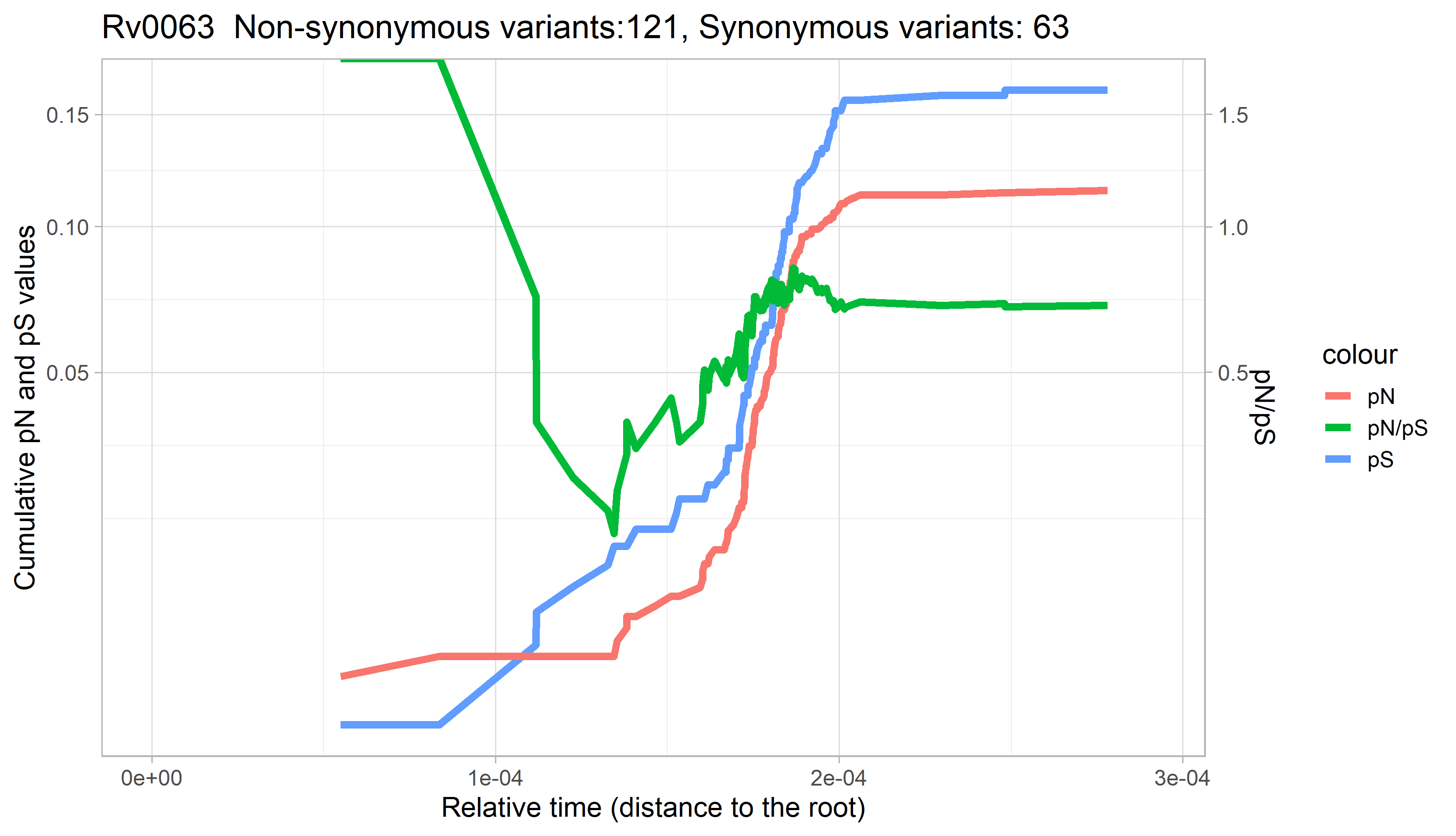

### Rv0064.png

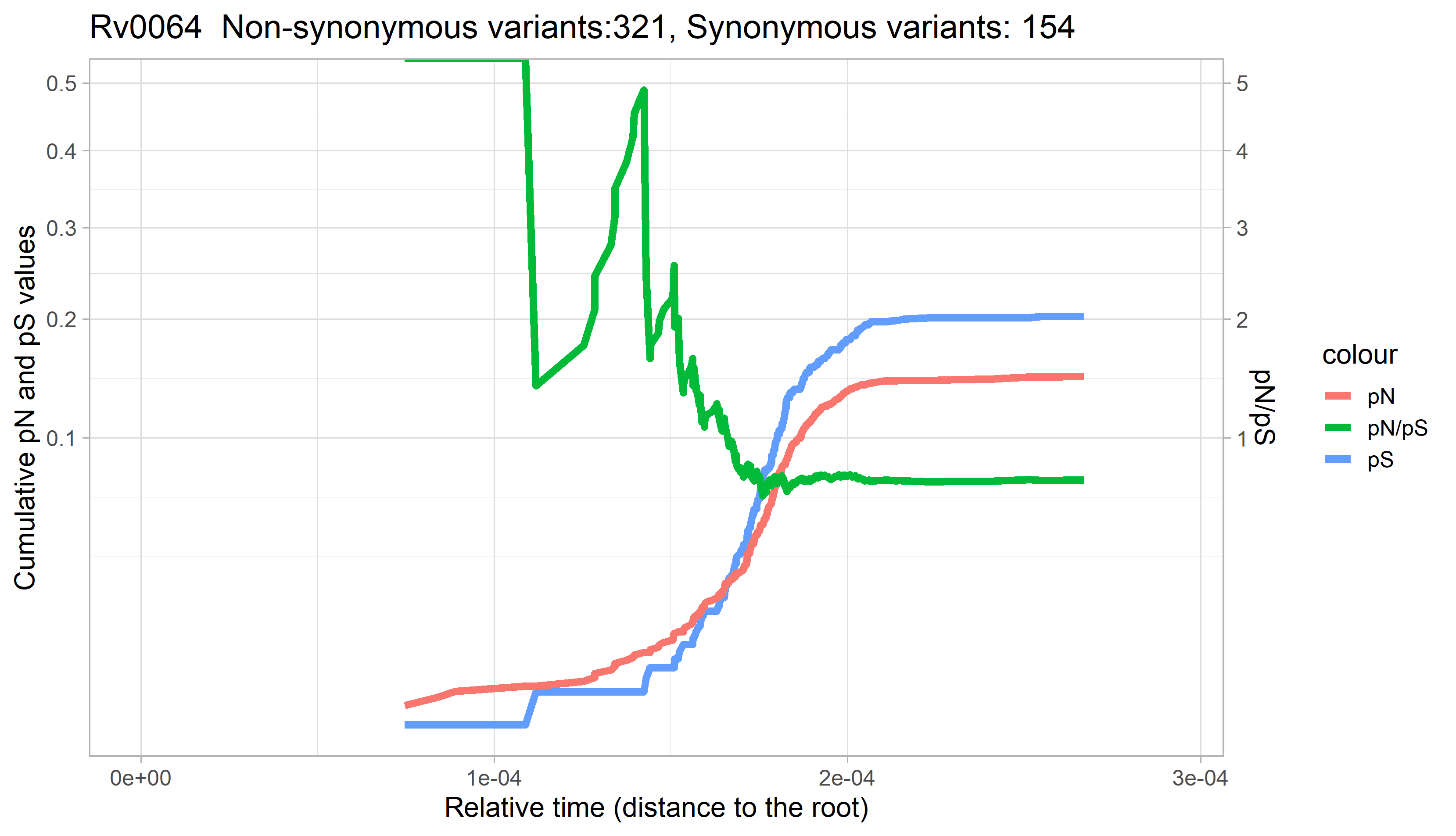

### Rv0065.png

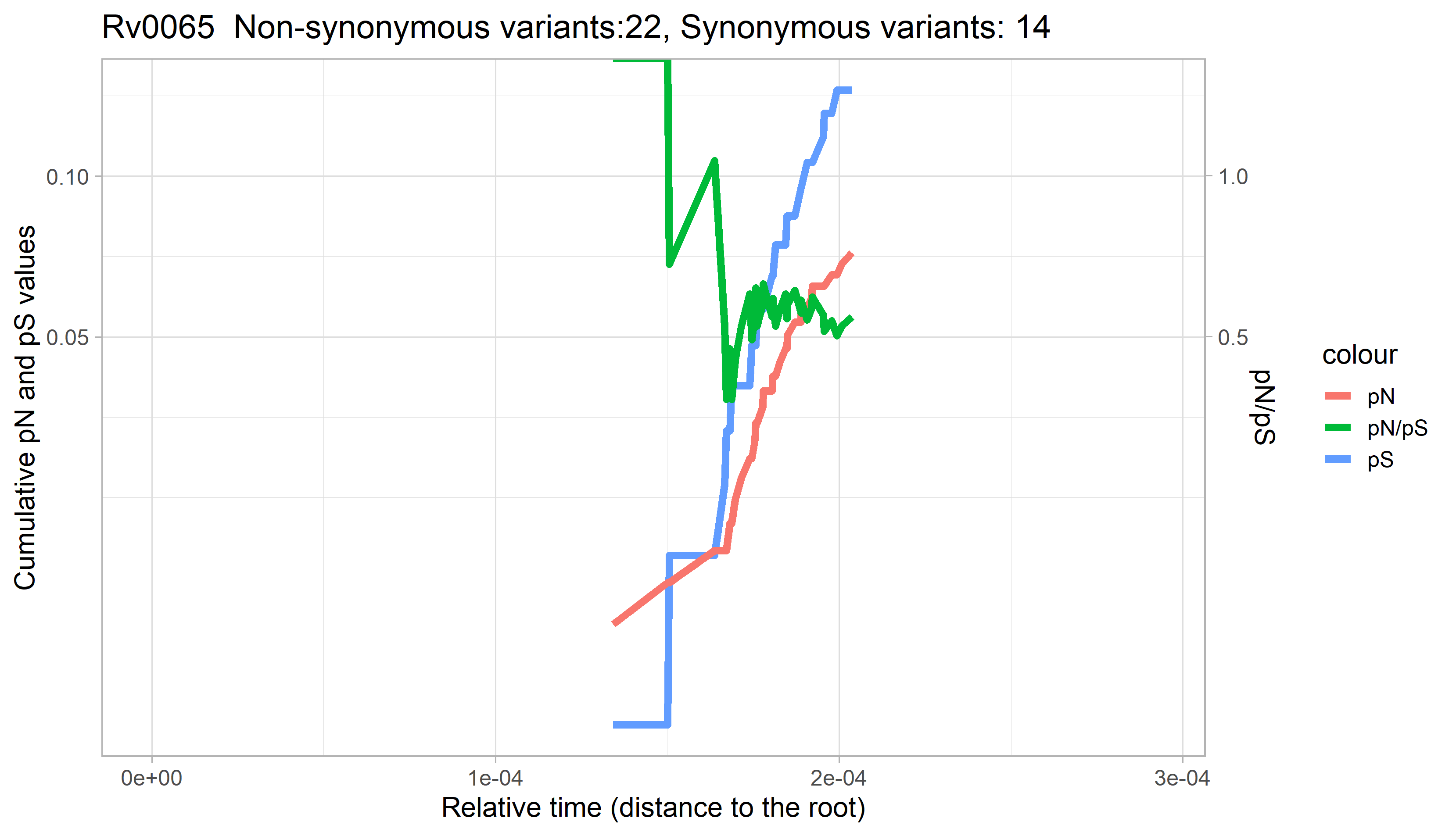

### Rv0066c.png

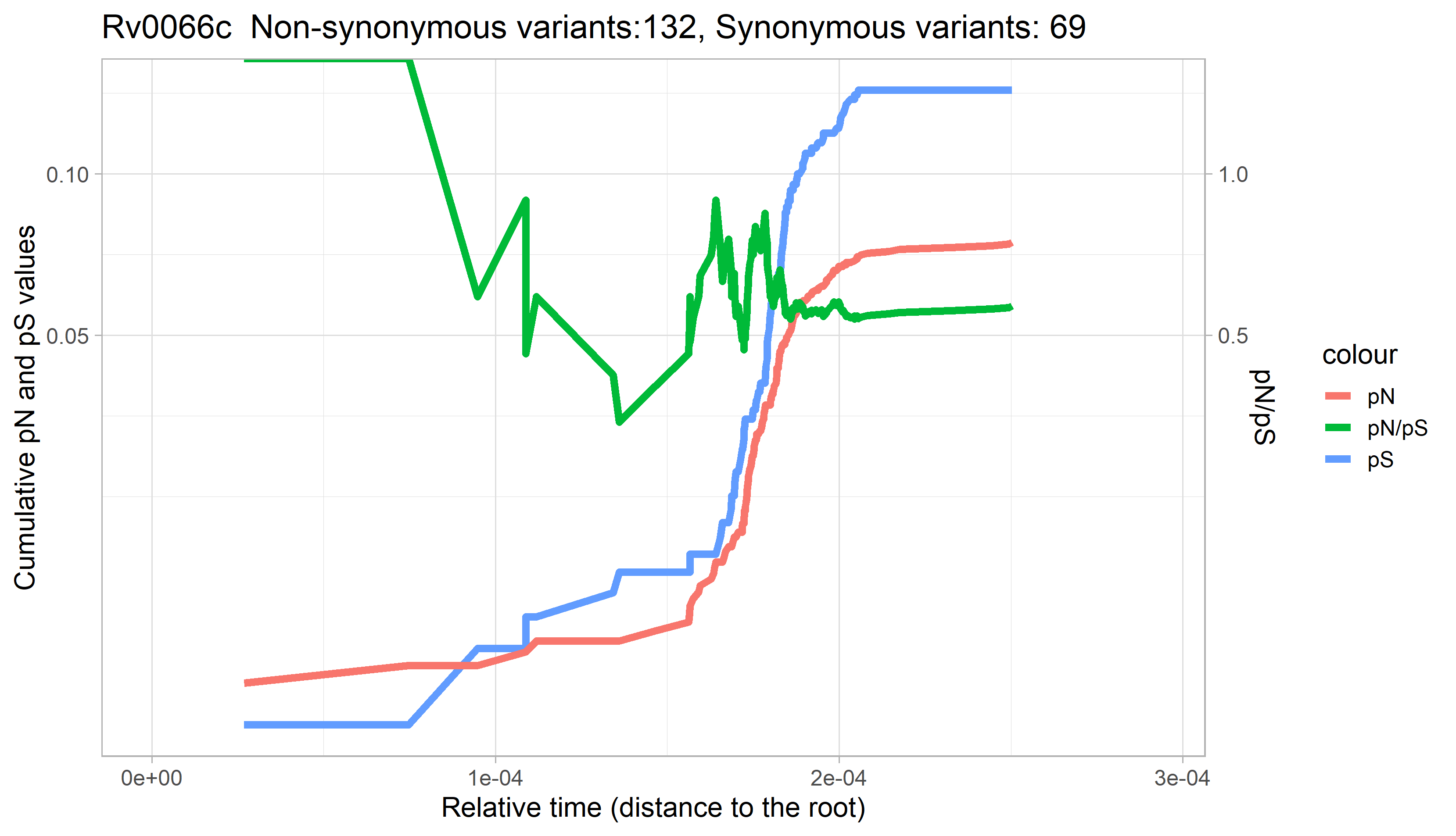

### Rv0067c.png

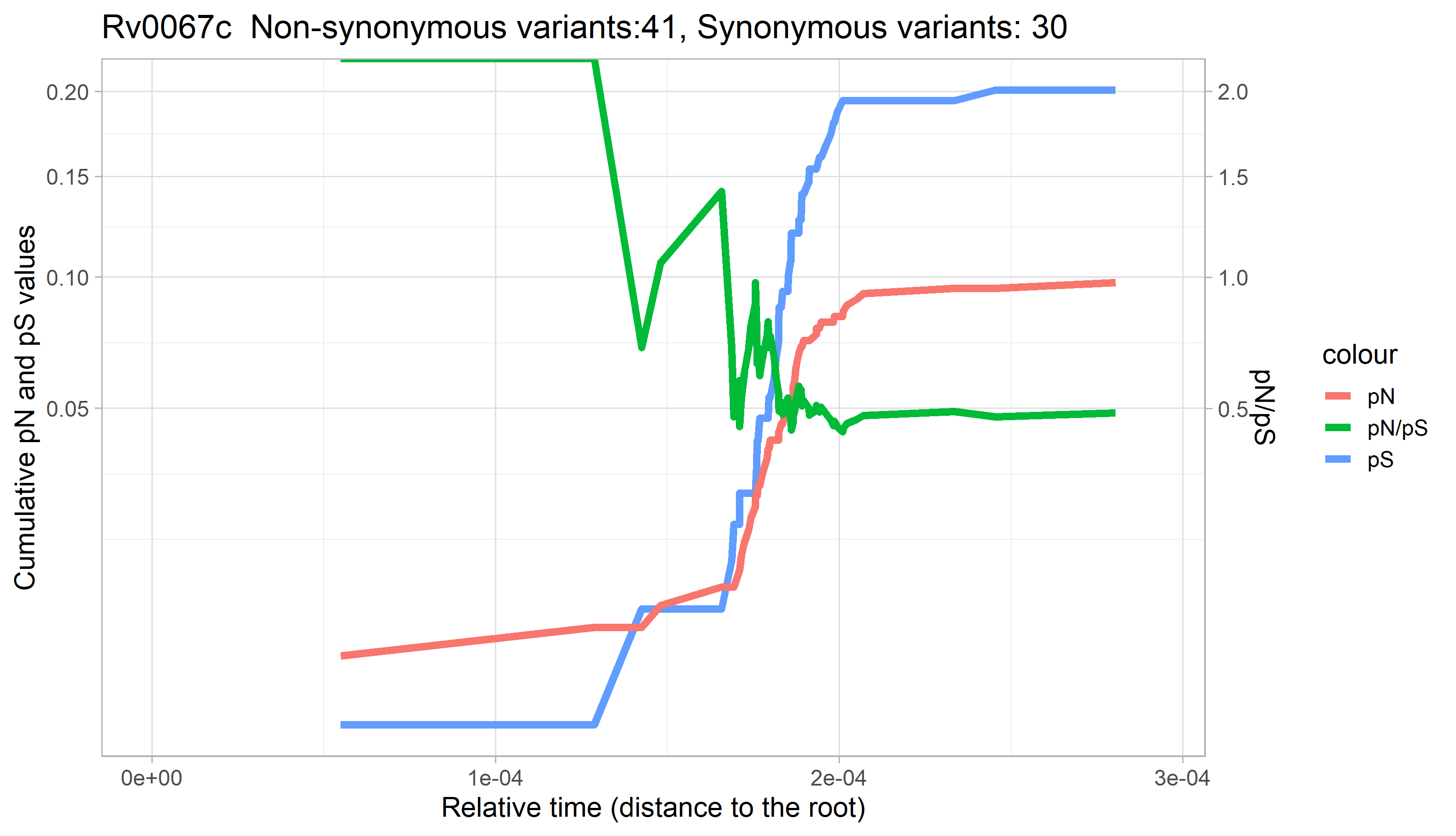

### Rv0068.png

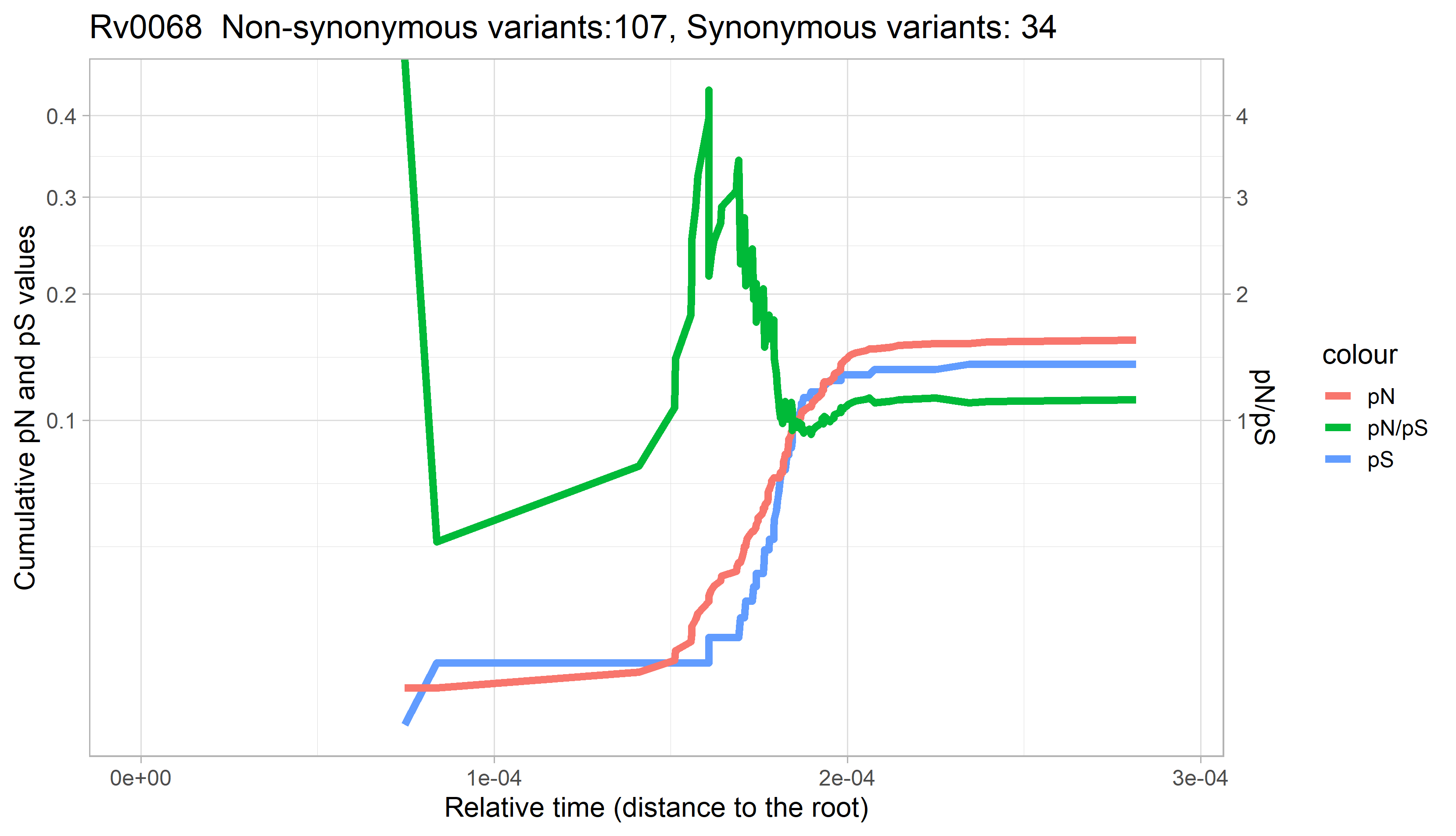

### Rv0069c.png

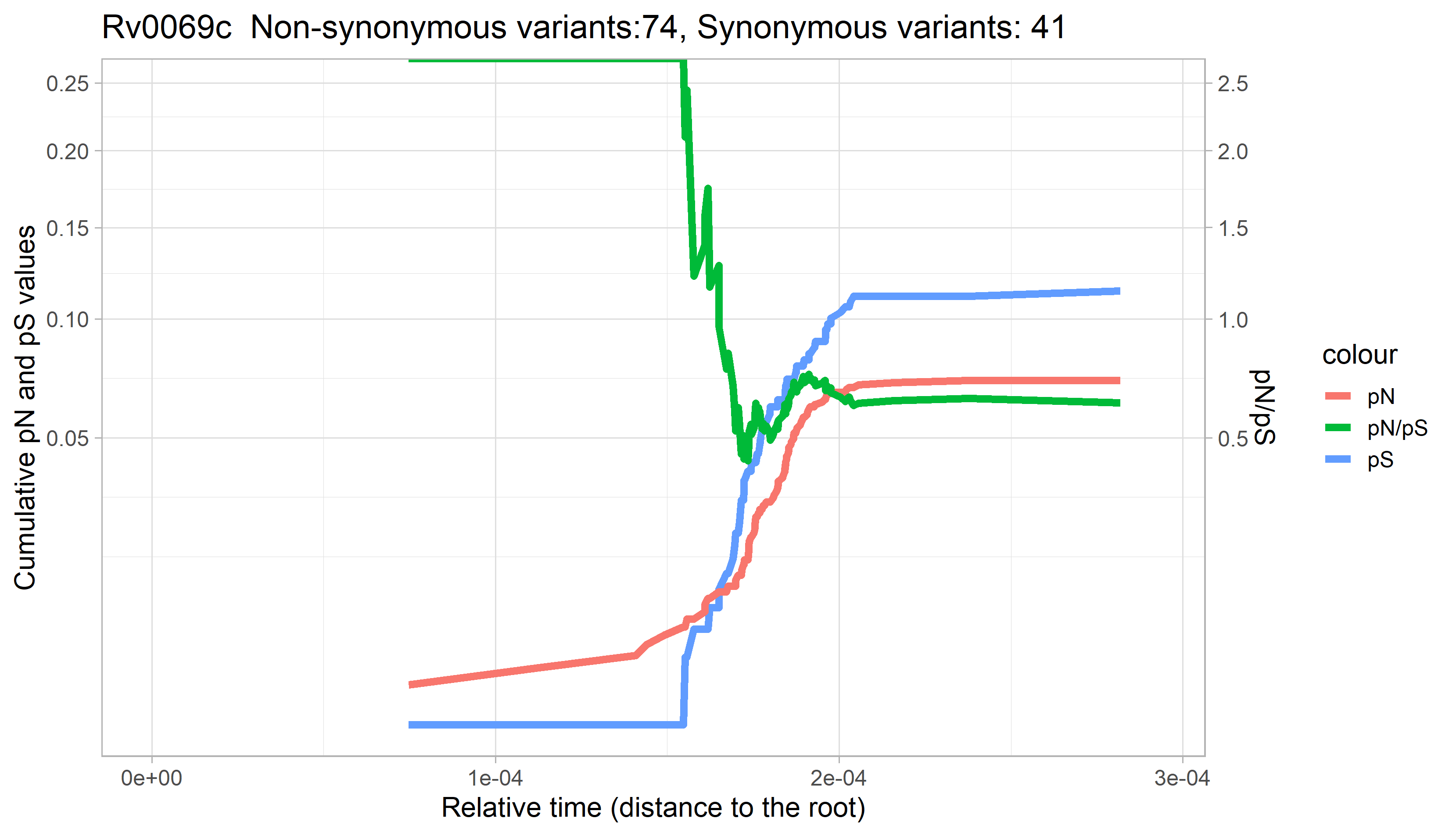

### Rv0070c.png

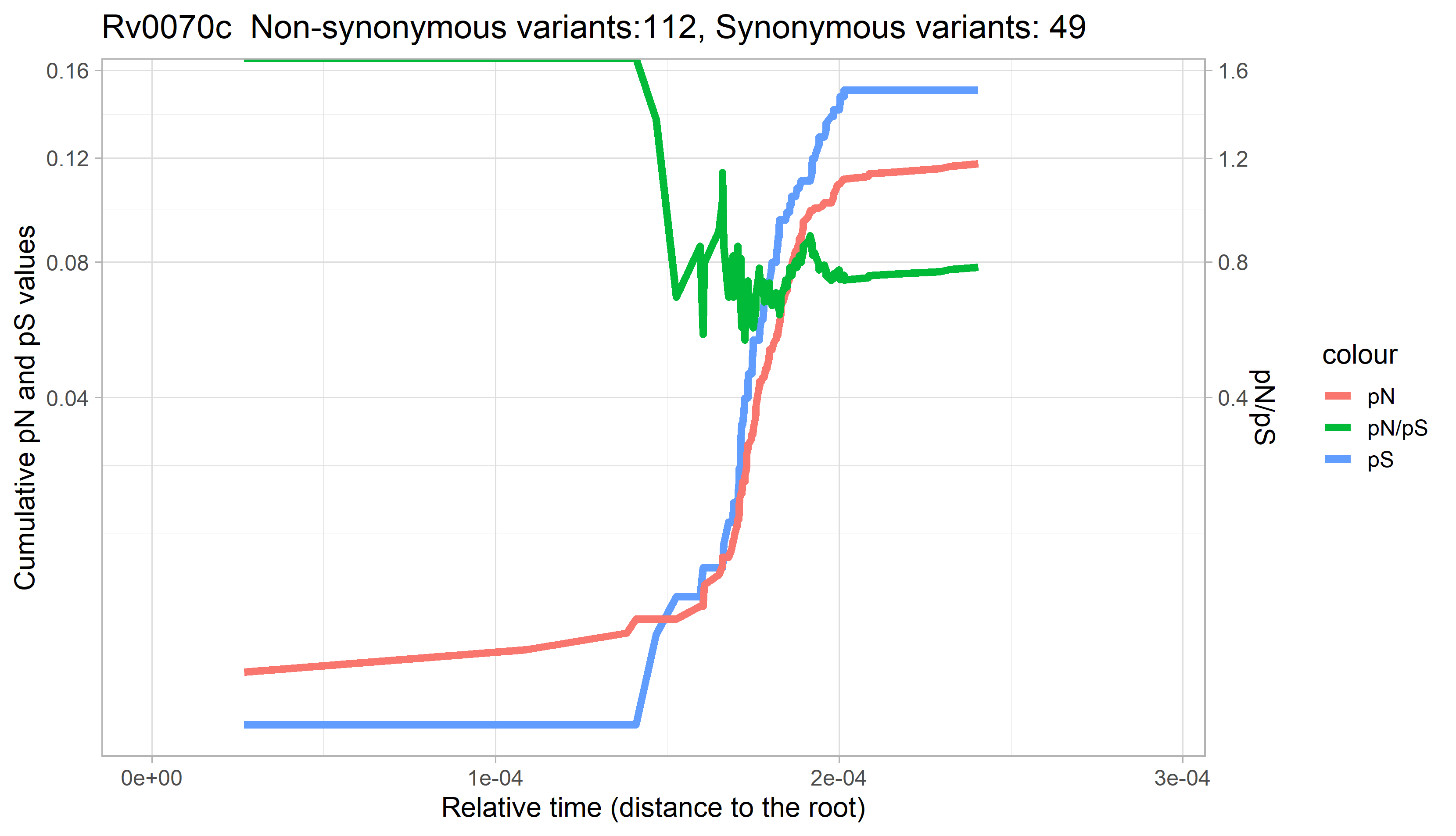

### Rv0071.png

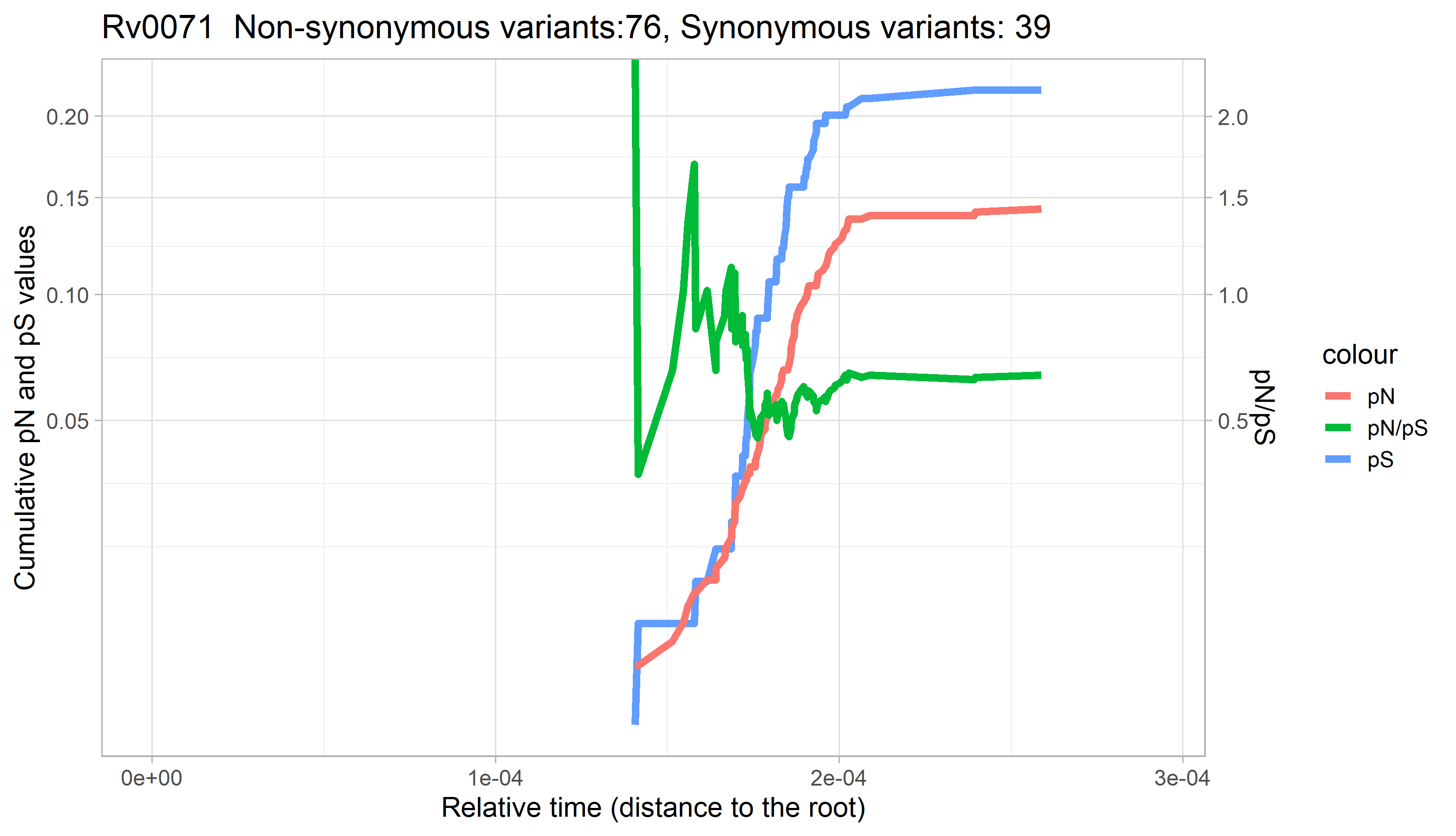

### Rv0072.png

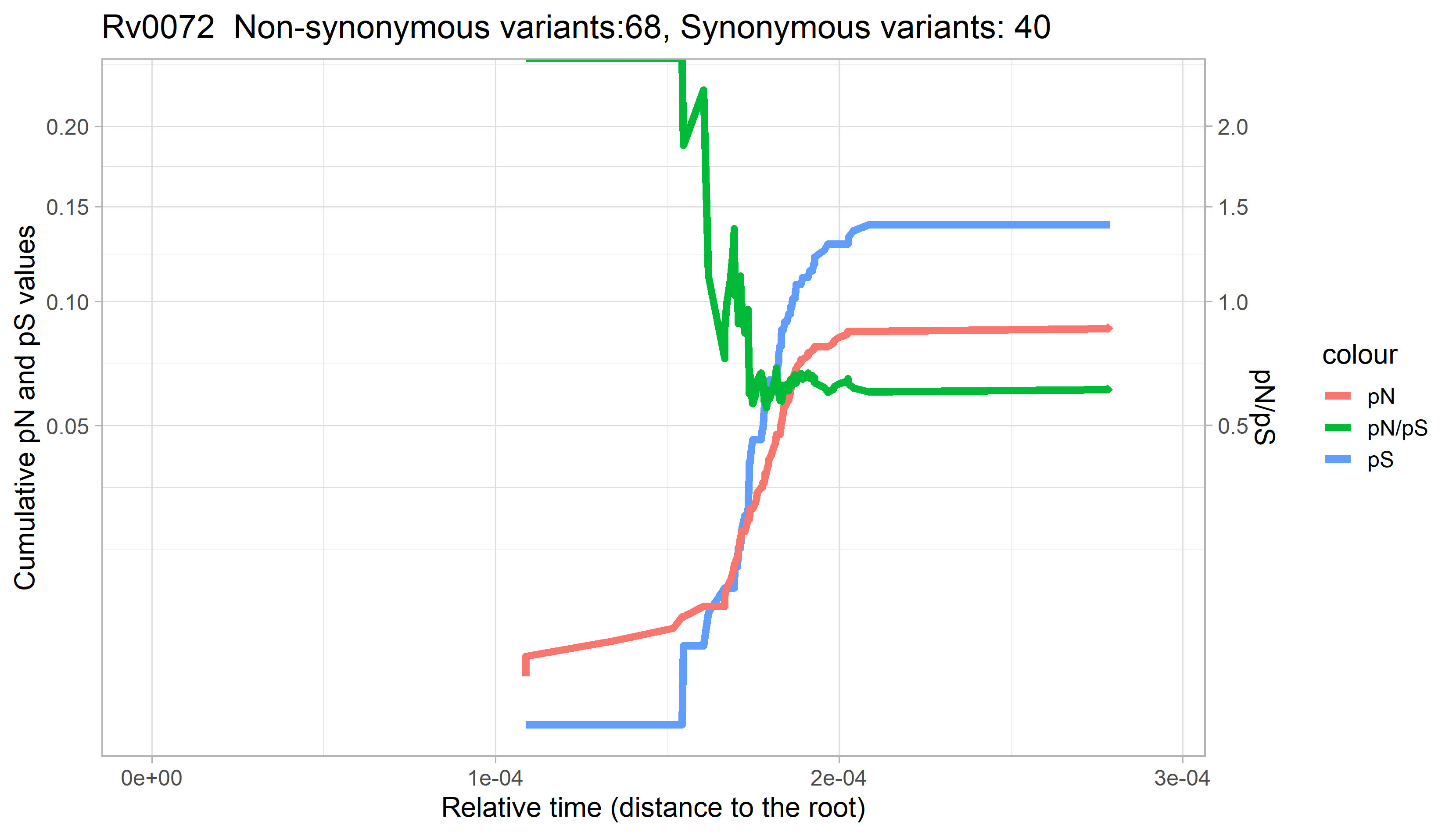

### Rv0073.png

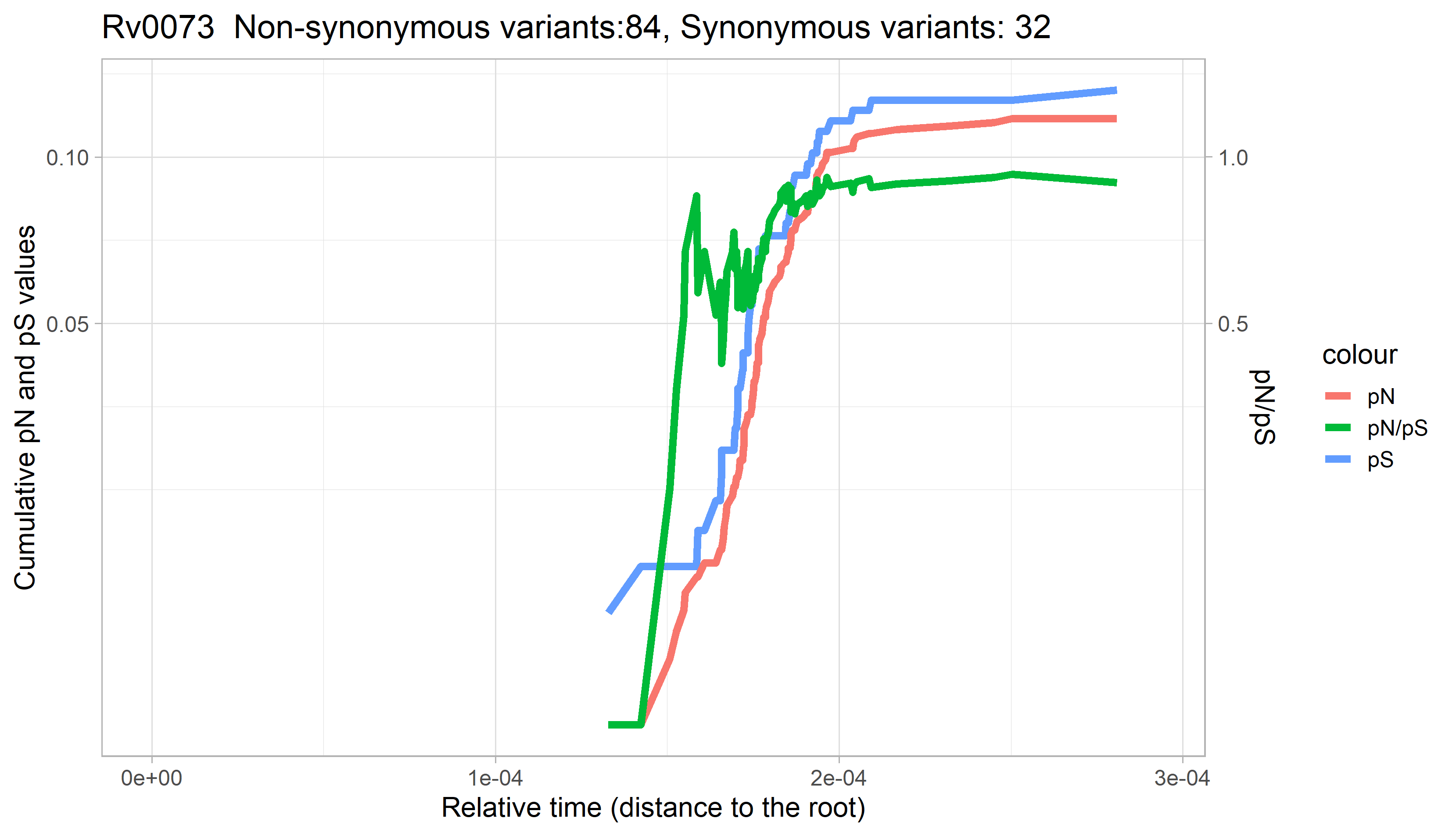

### Rv0074.png

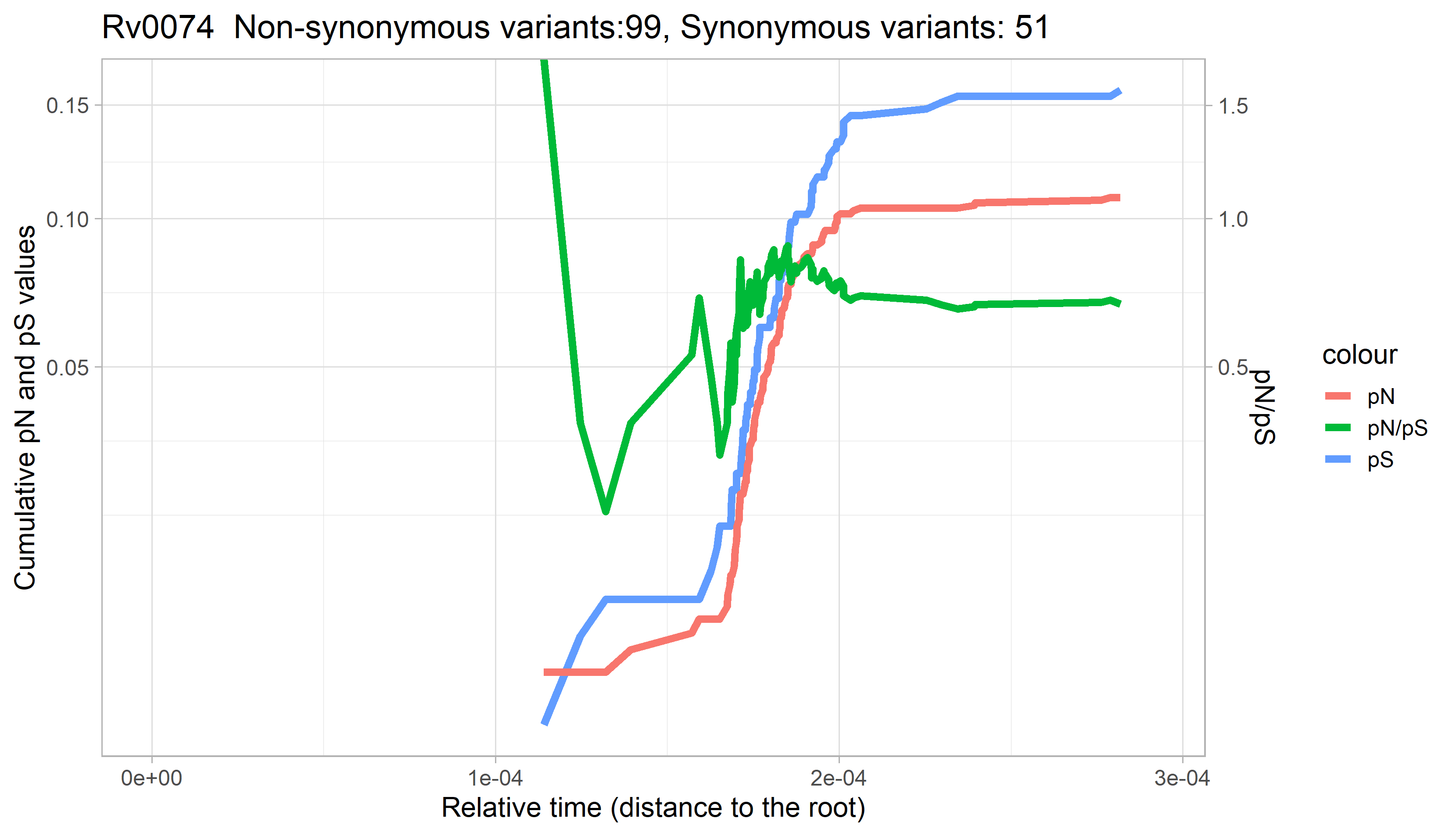

### Rv0075.png

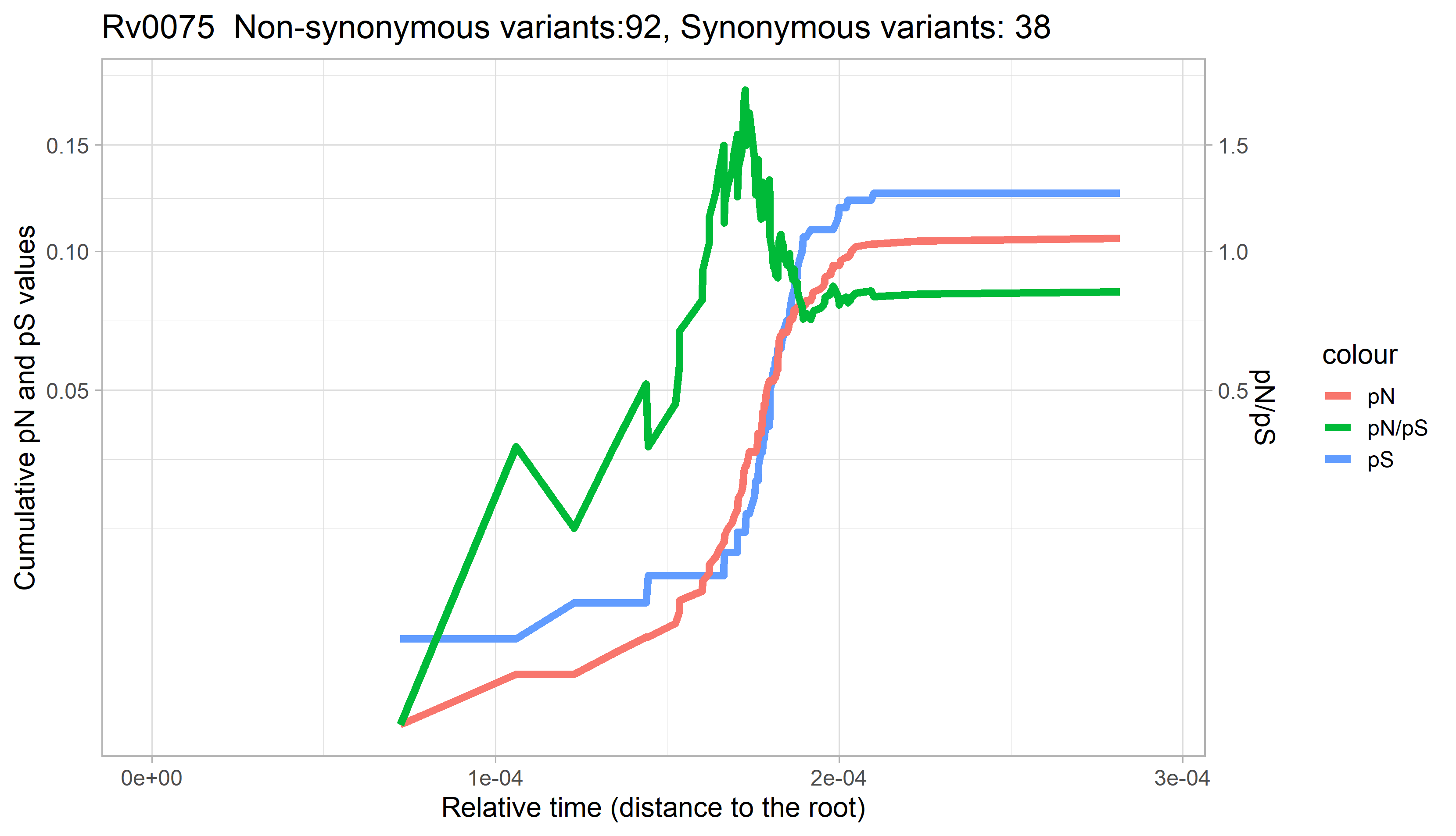

### Rv0076c.png

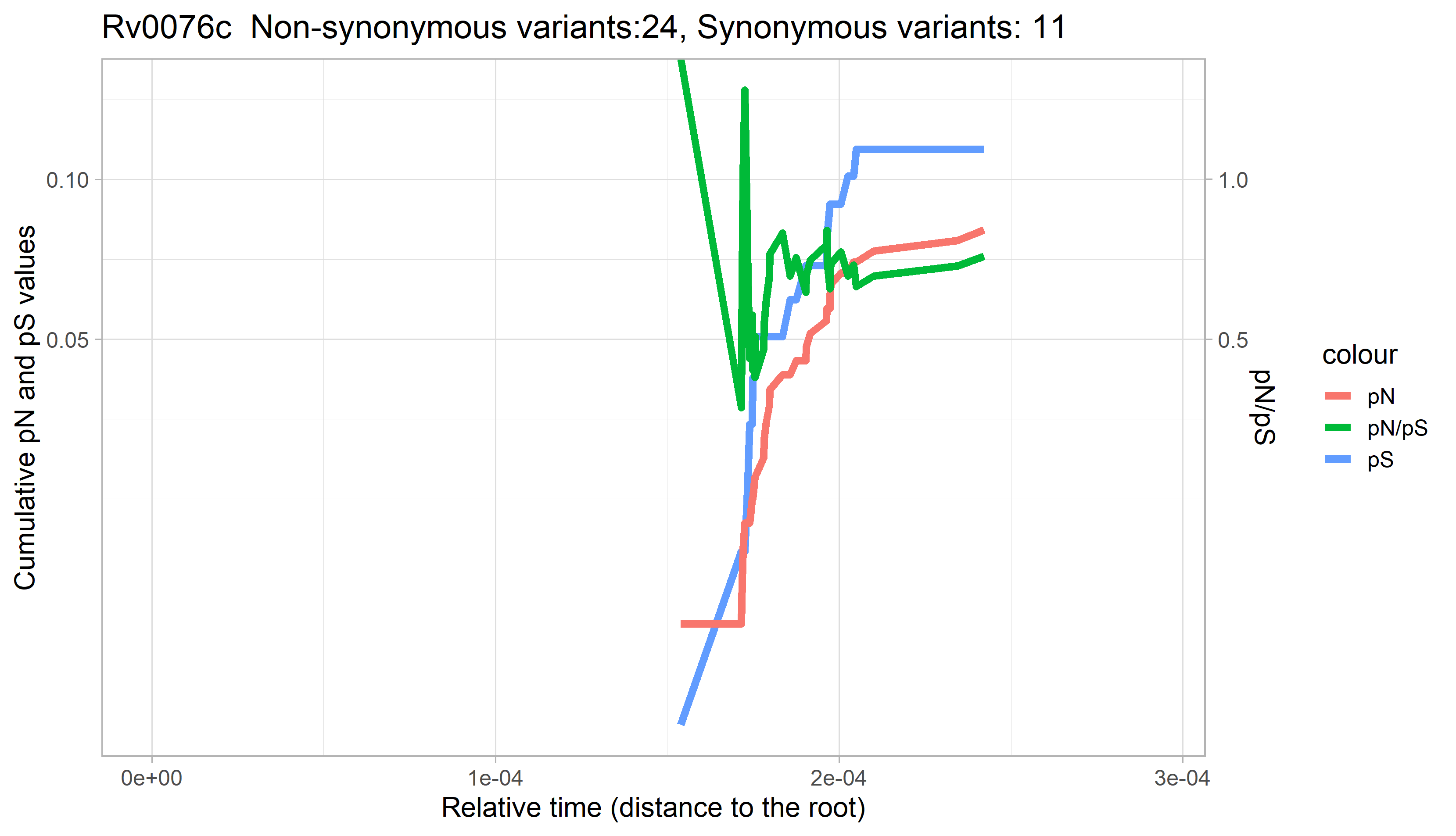

### Rv0077c.png

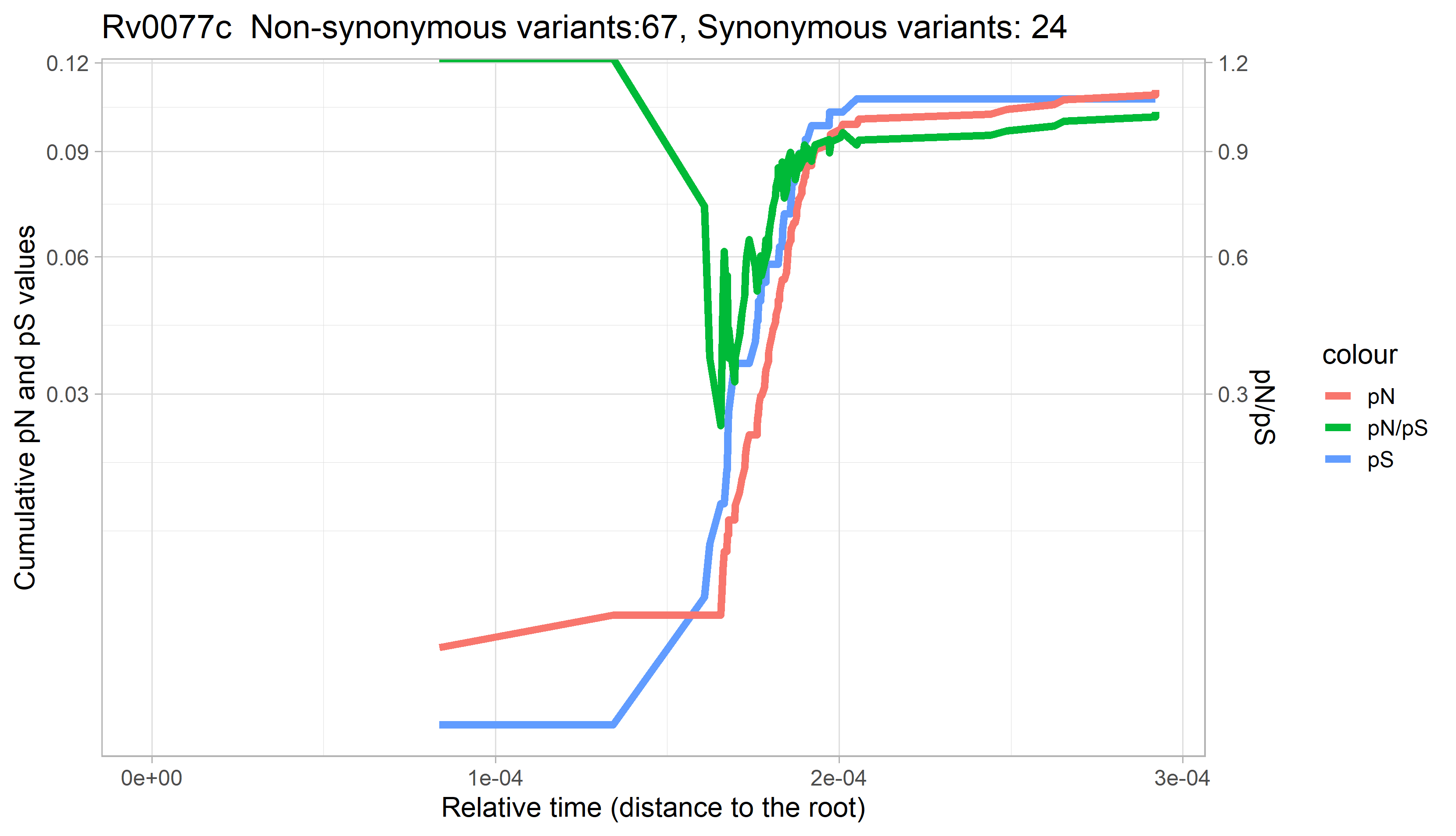

### Rv0078.png

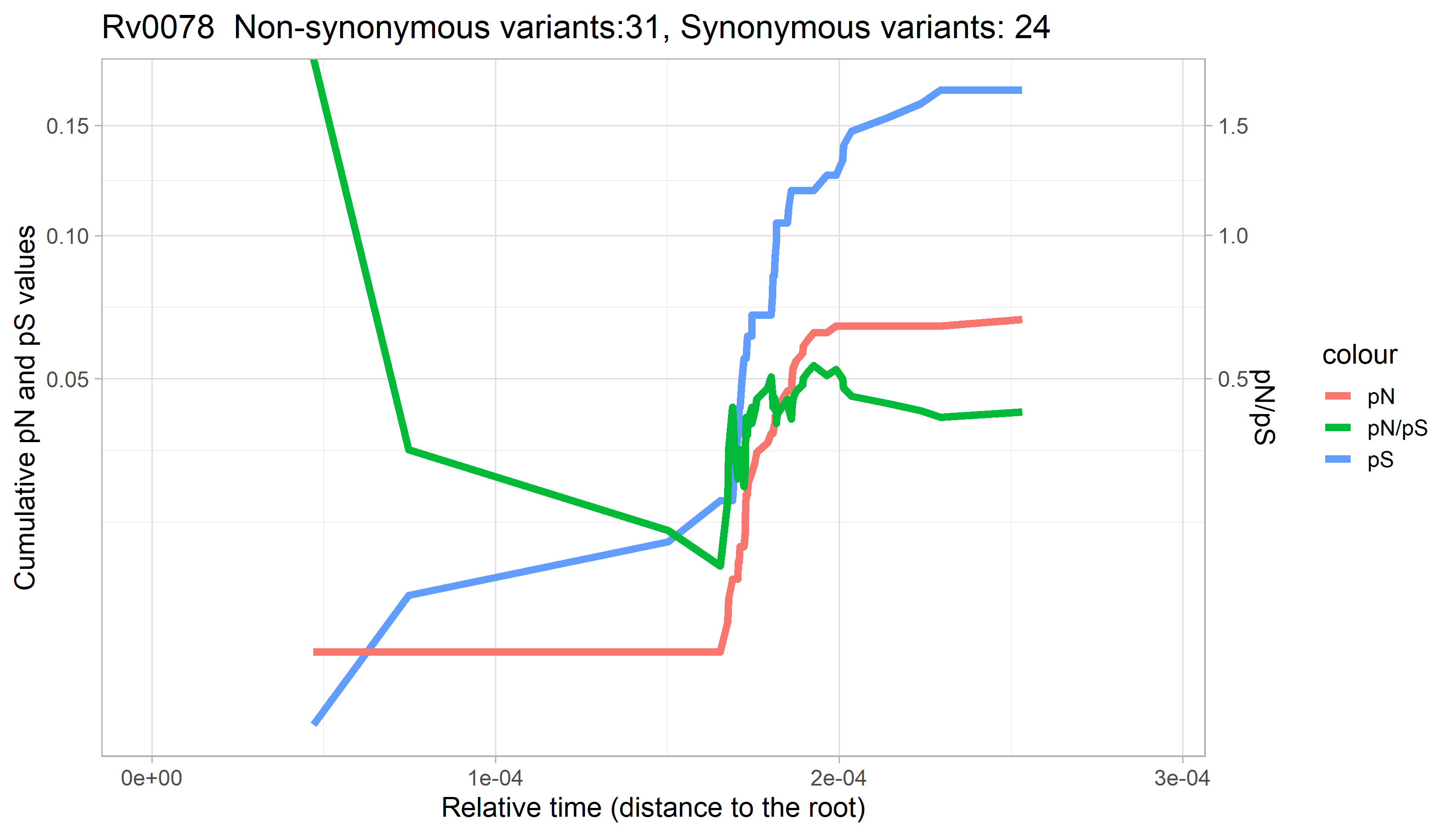

### Rv0078A.png

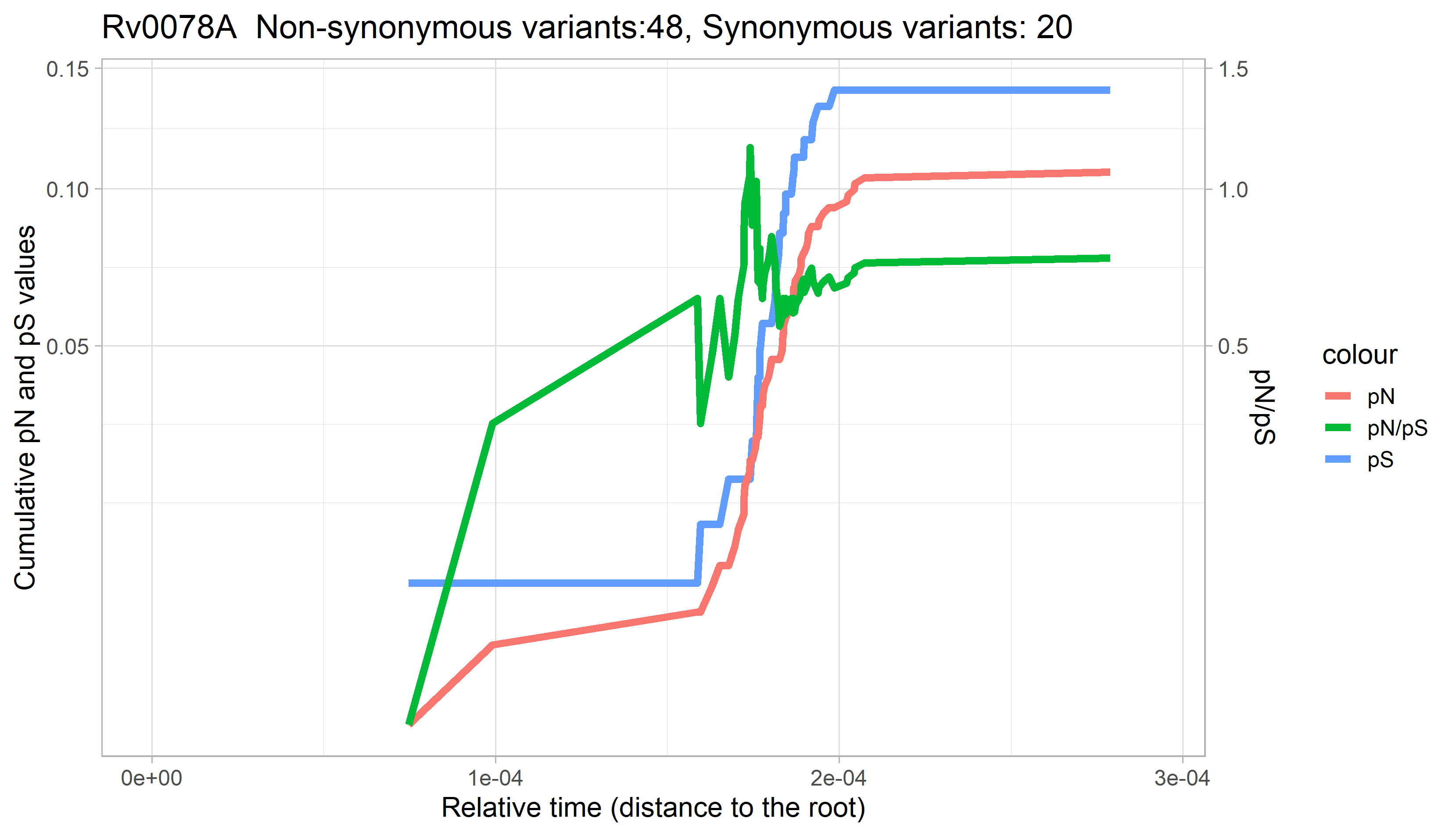

### Rv0079.png

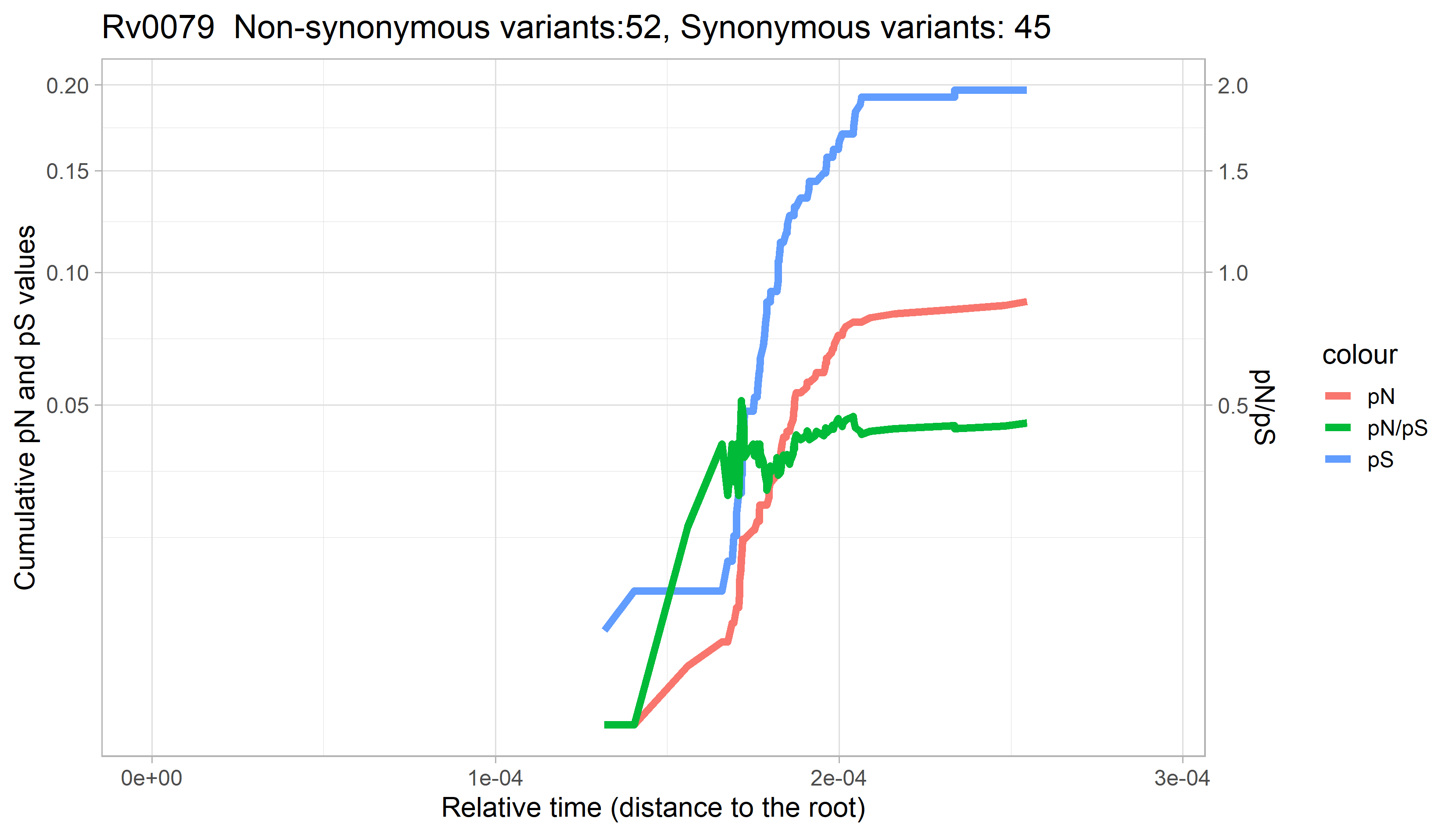

### Rv0080.png

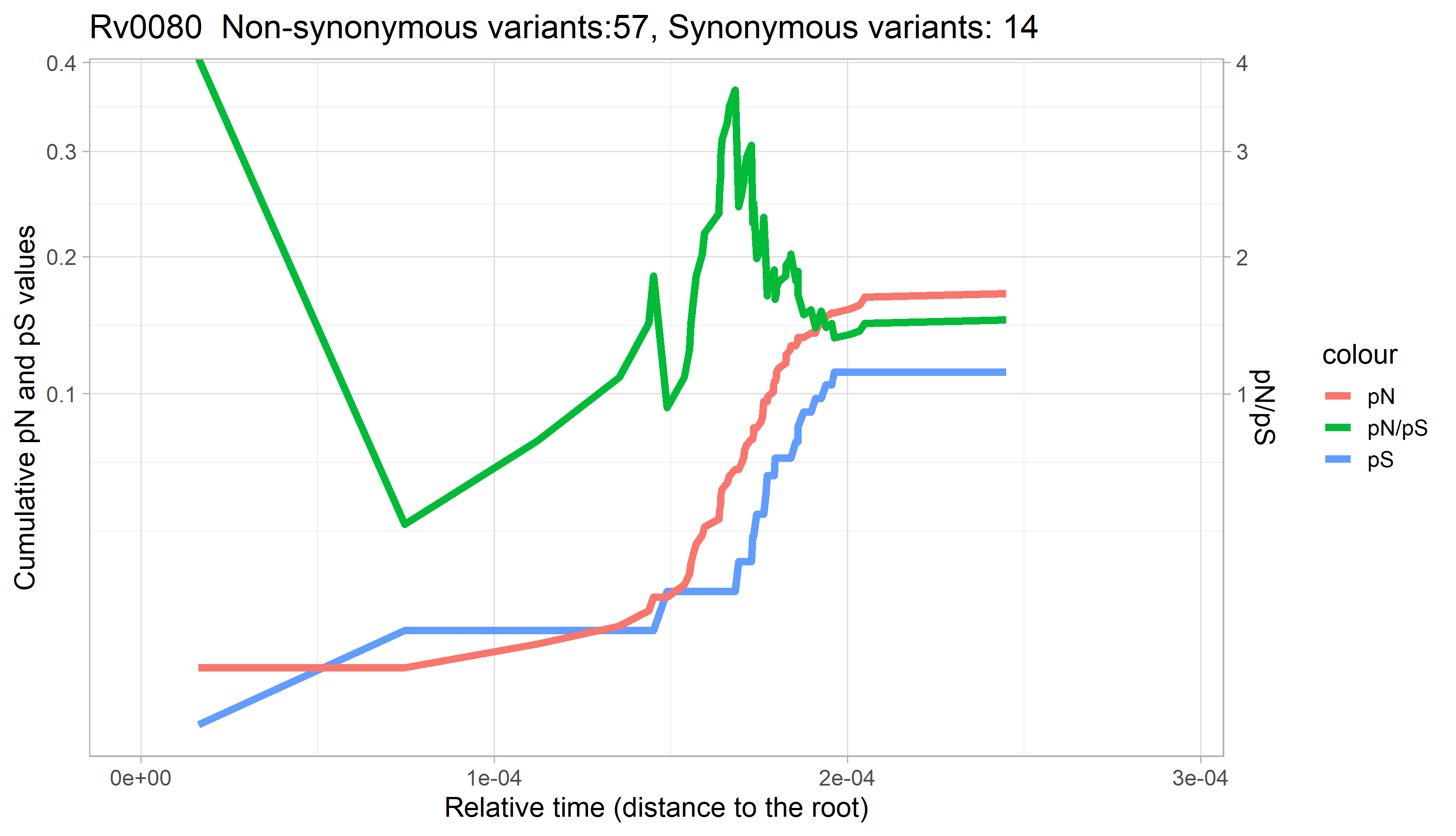

### Rv0081.png

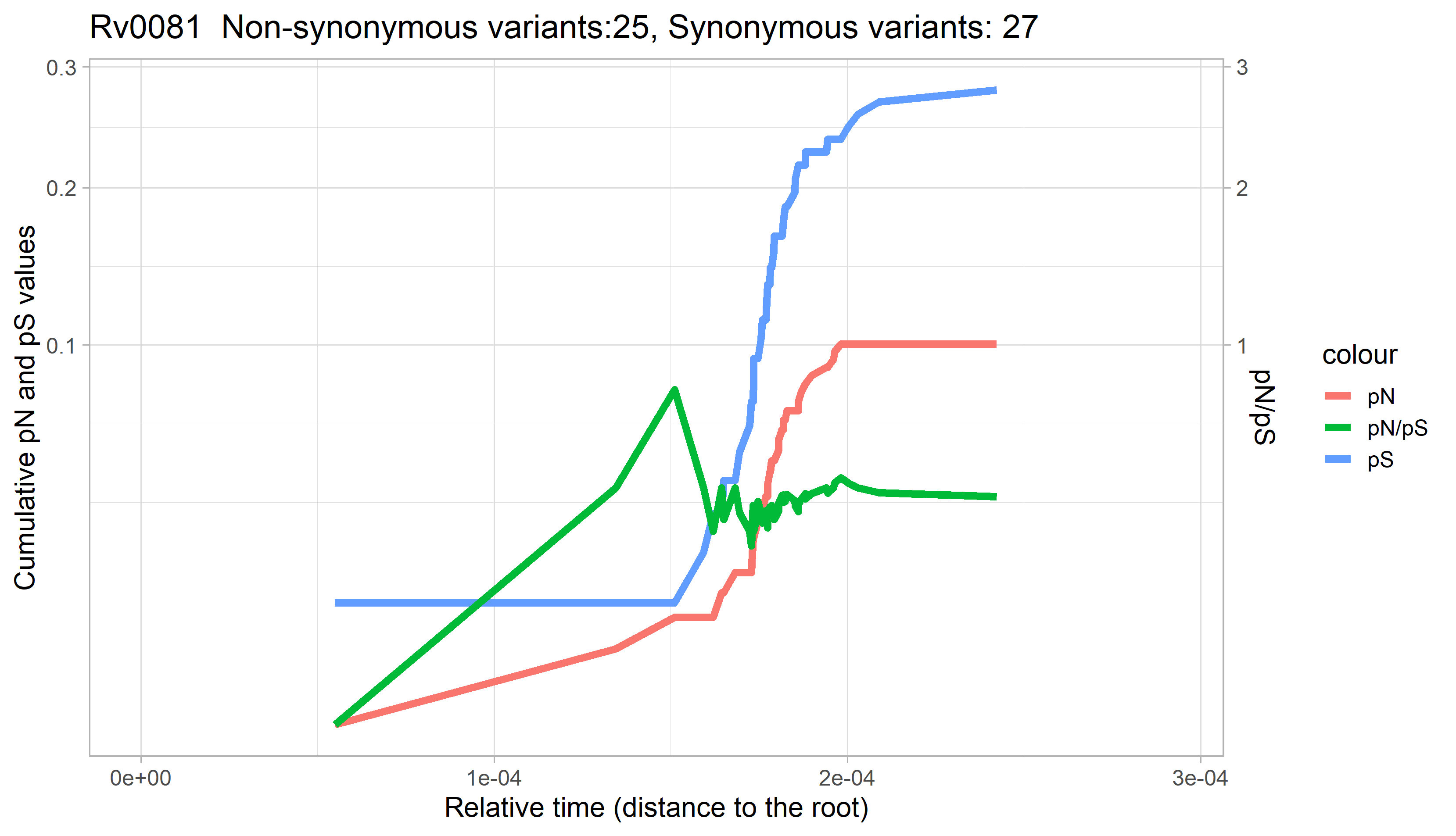

### Rv0082.png

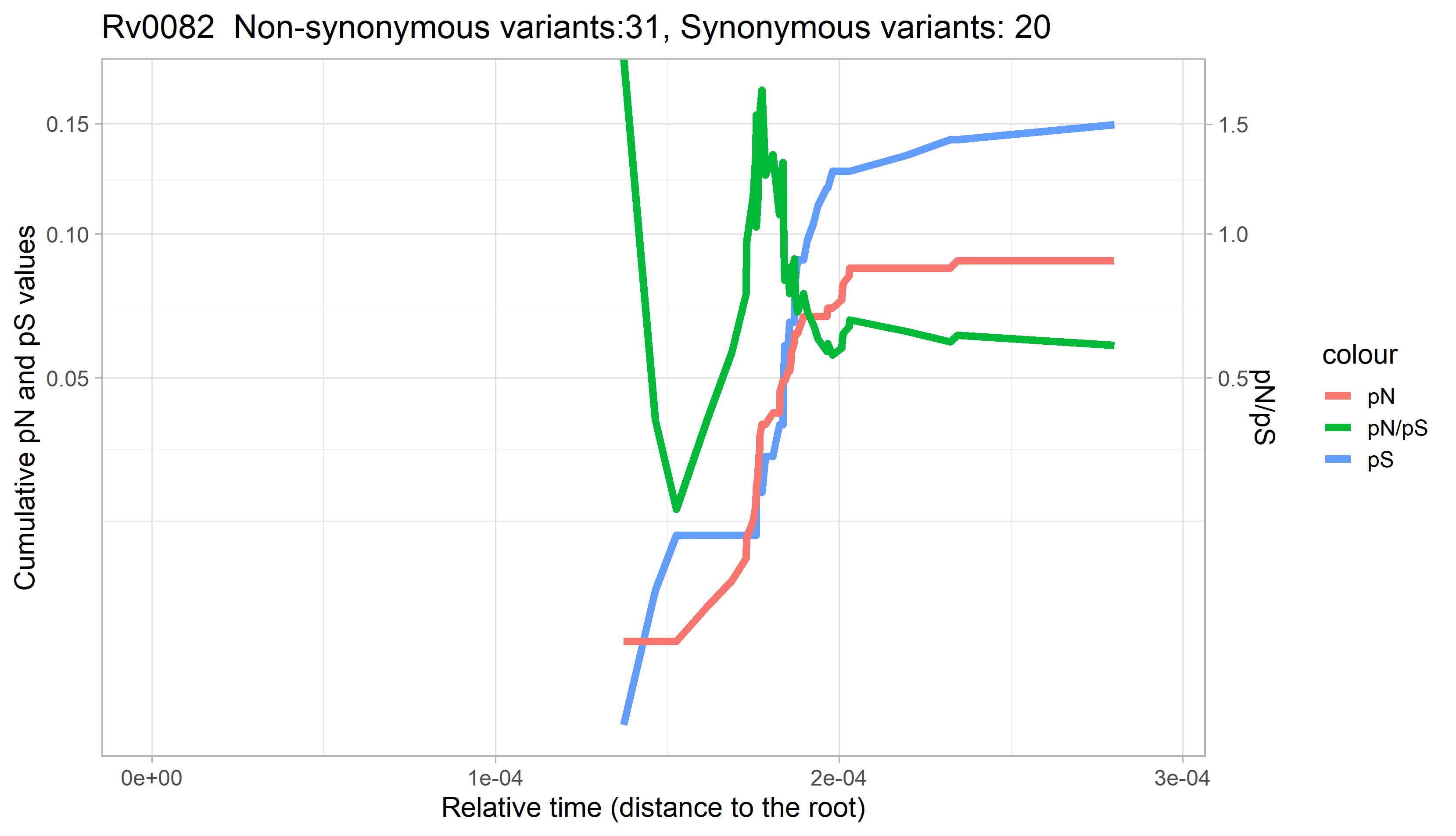

### Rv0083.png

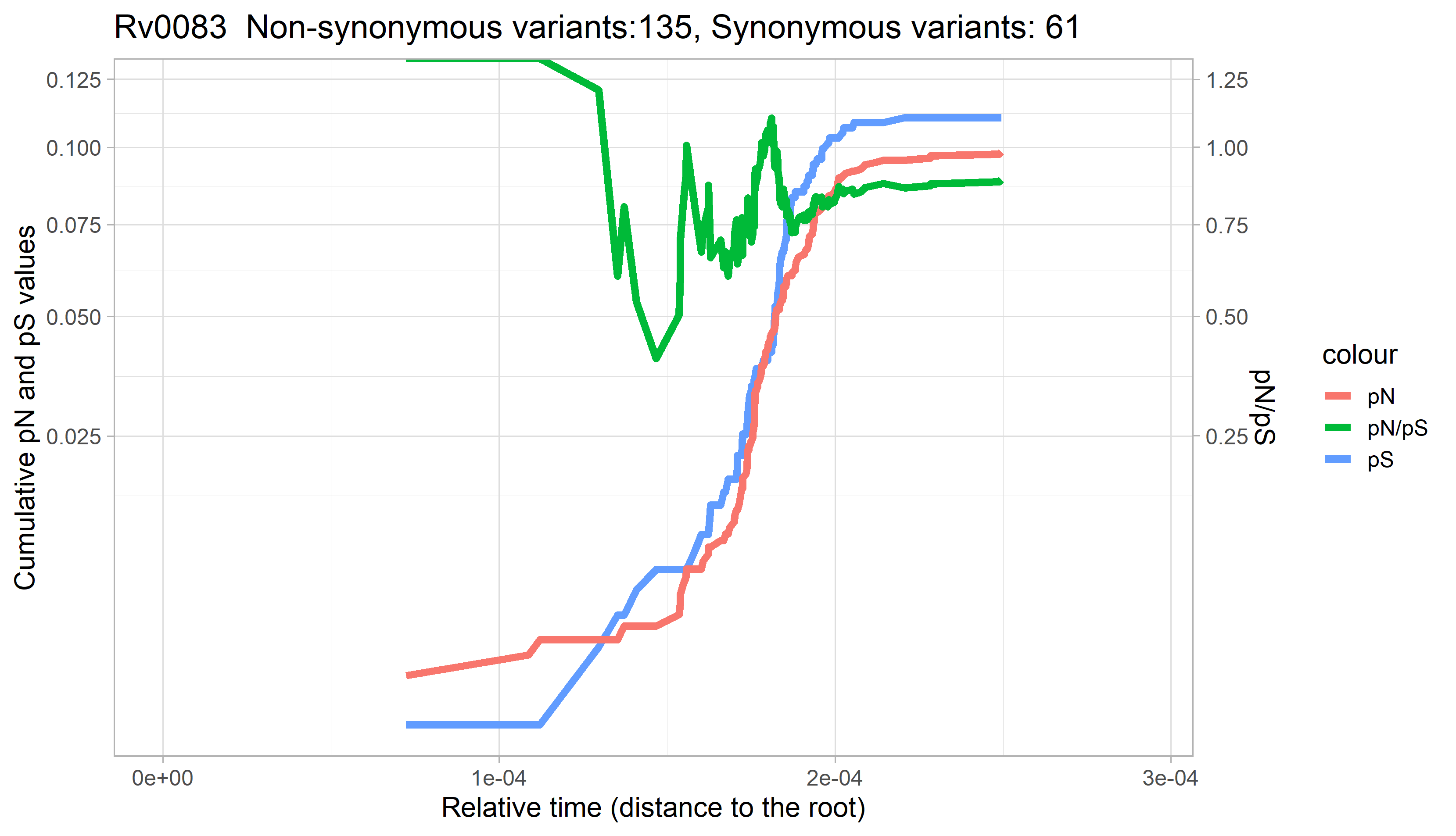

### Rv0084.png

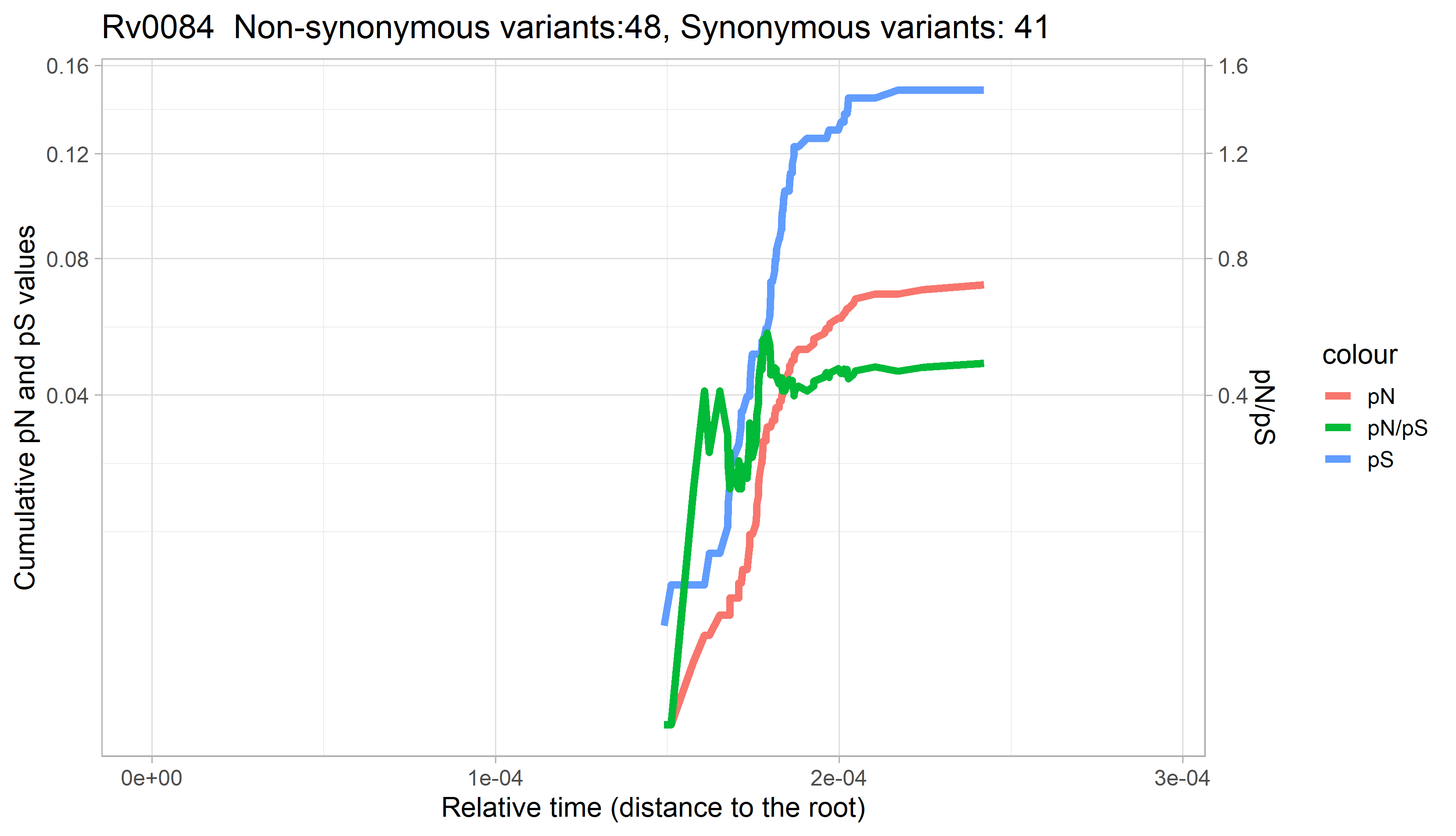

### Rv0085.png

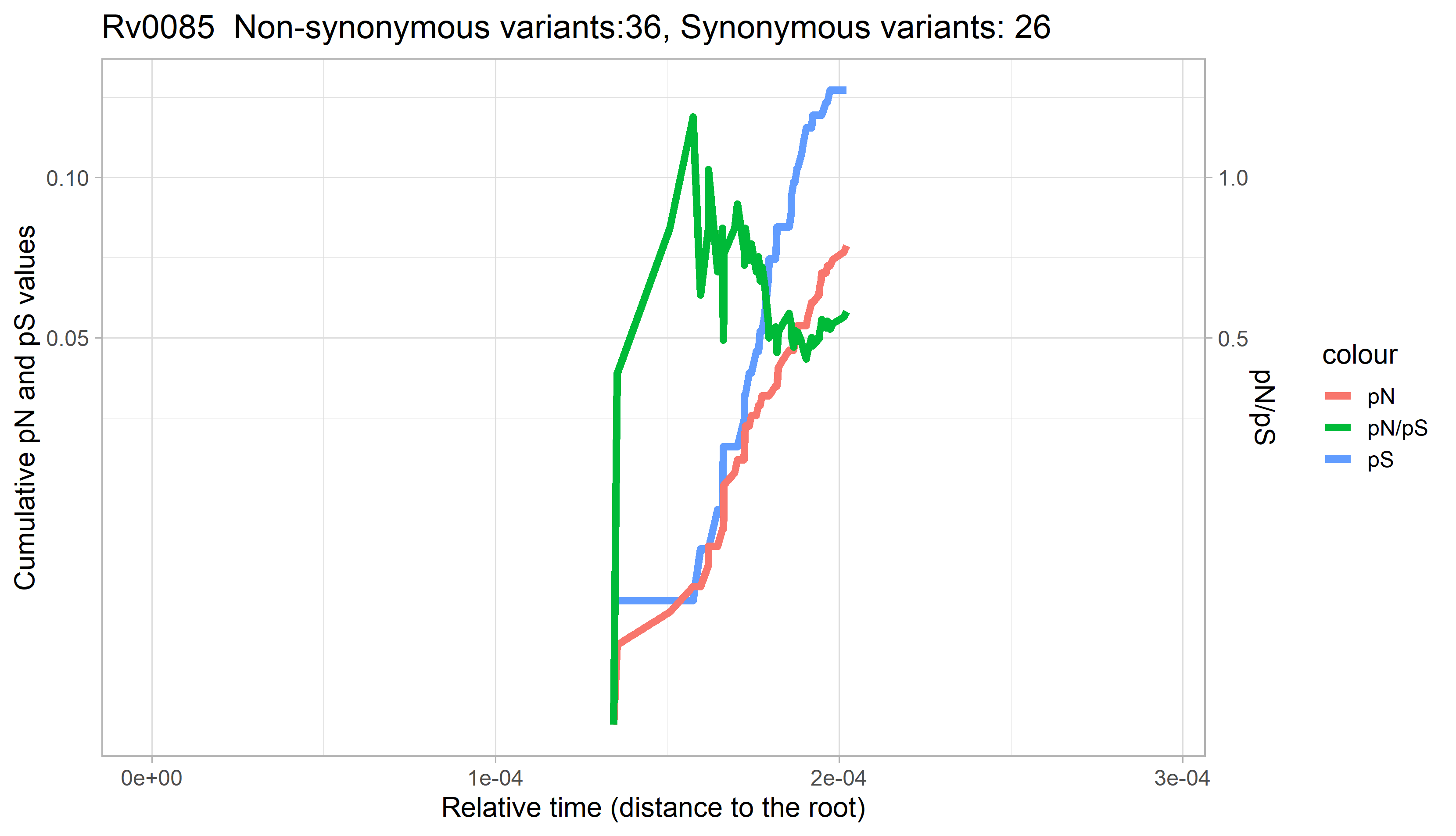

### Rv0086.png

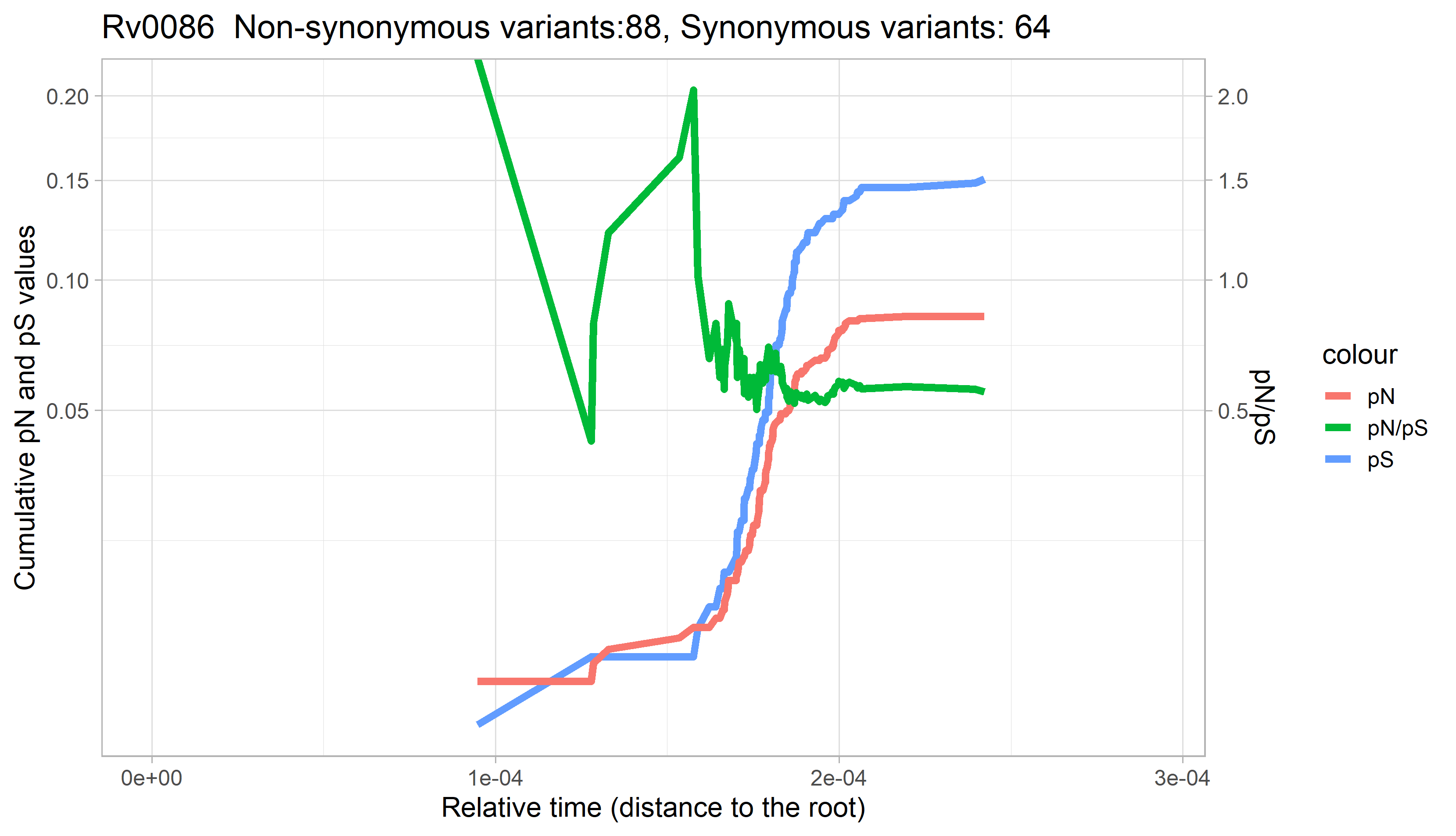

### Rv0087.png

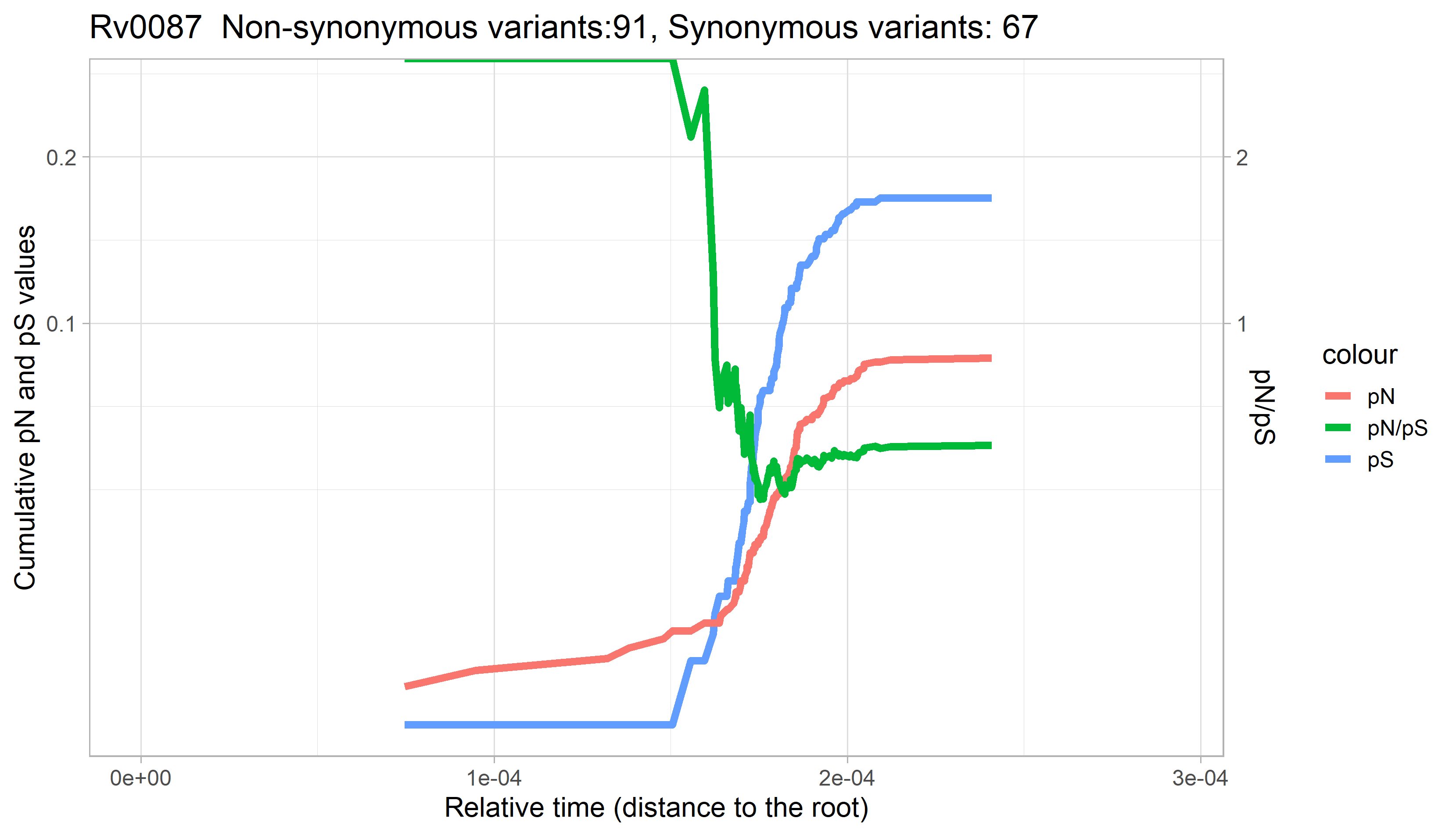

### Rv0088.png

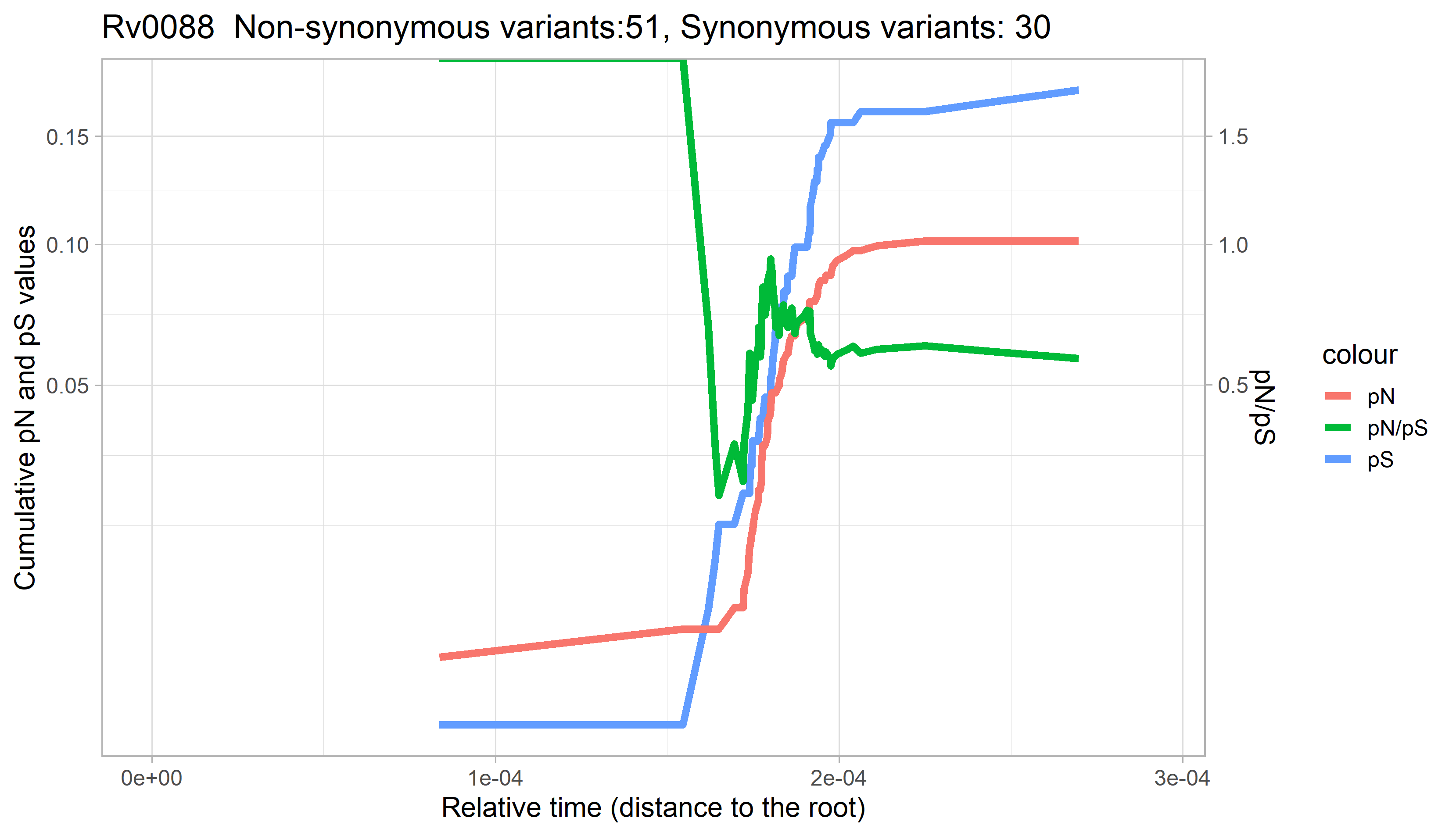

### Rv0089.png

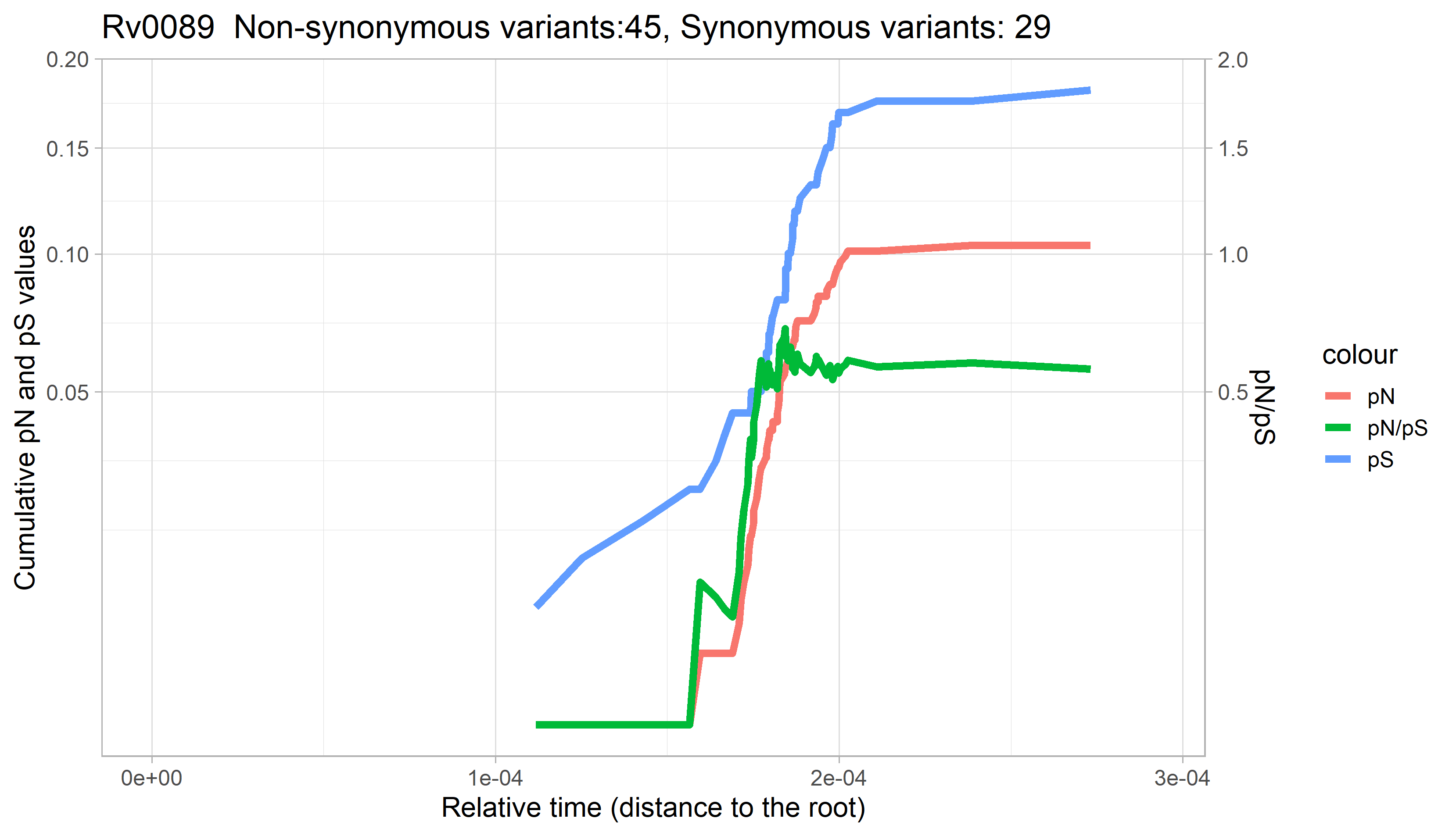

### Rv0090.png

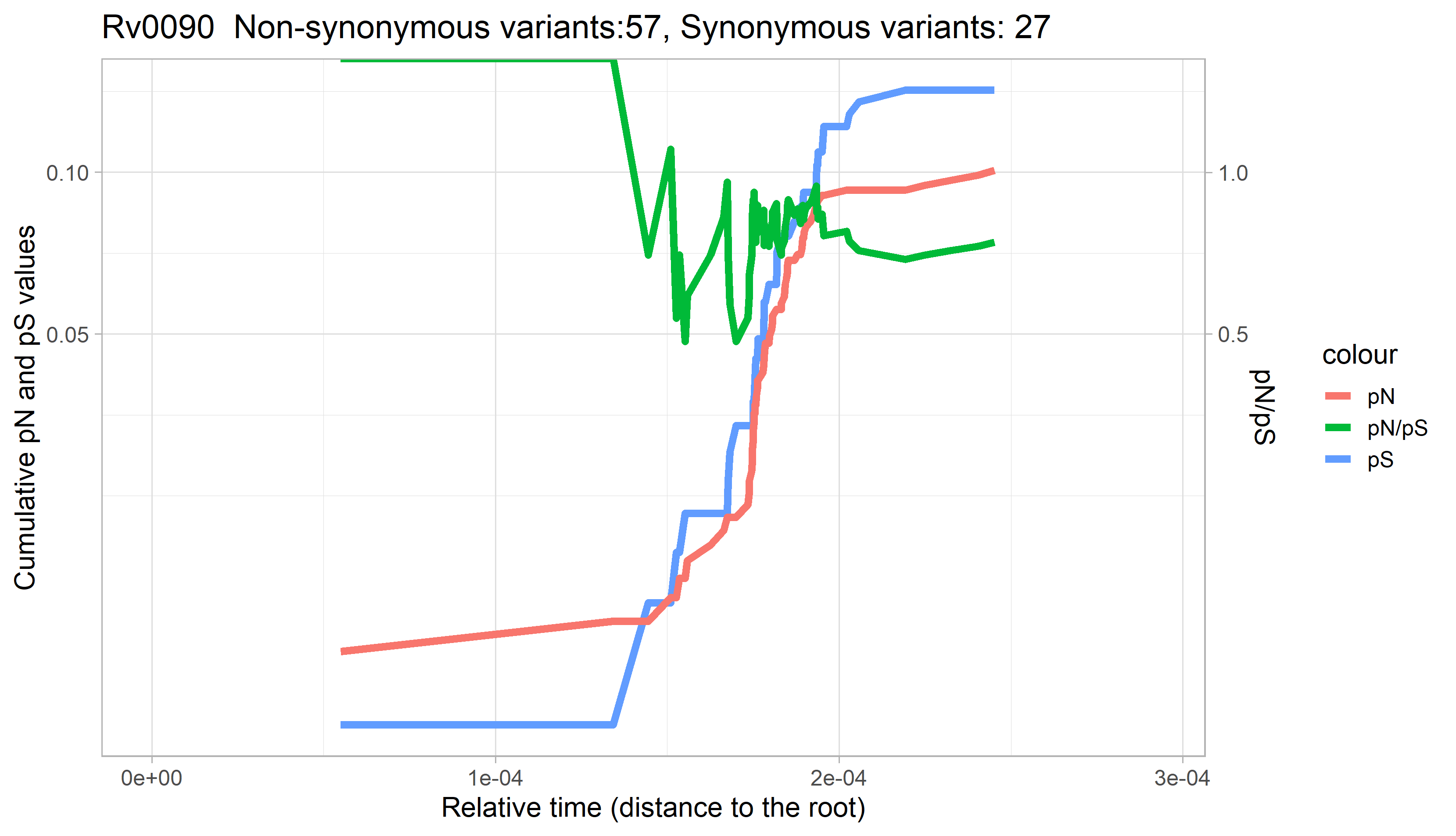

### Rv0091.png

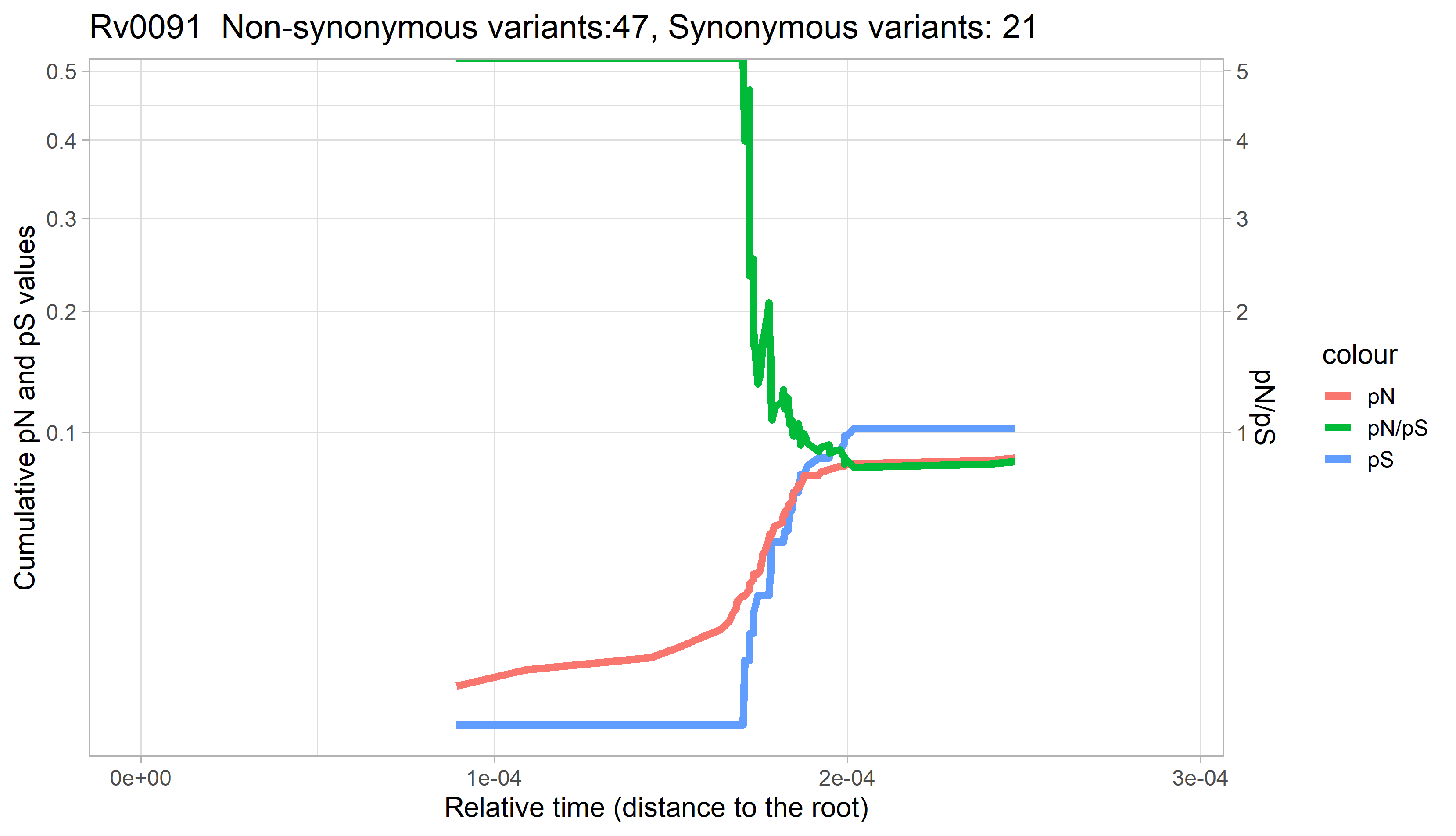

### Rv0092.png

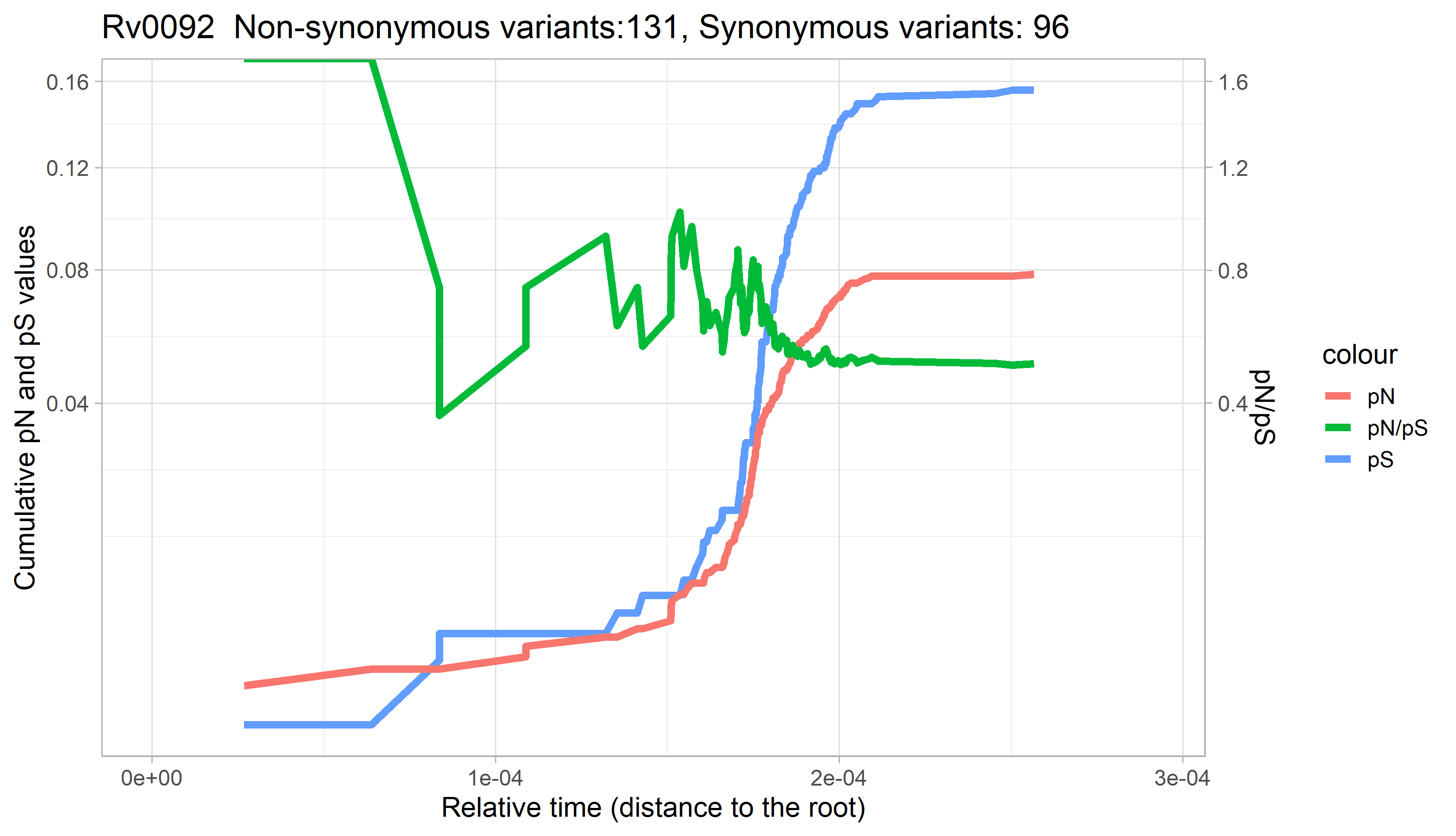

### Rv0093c.png

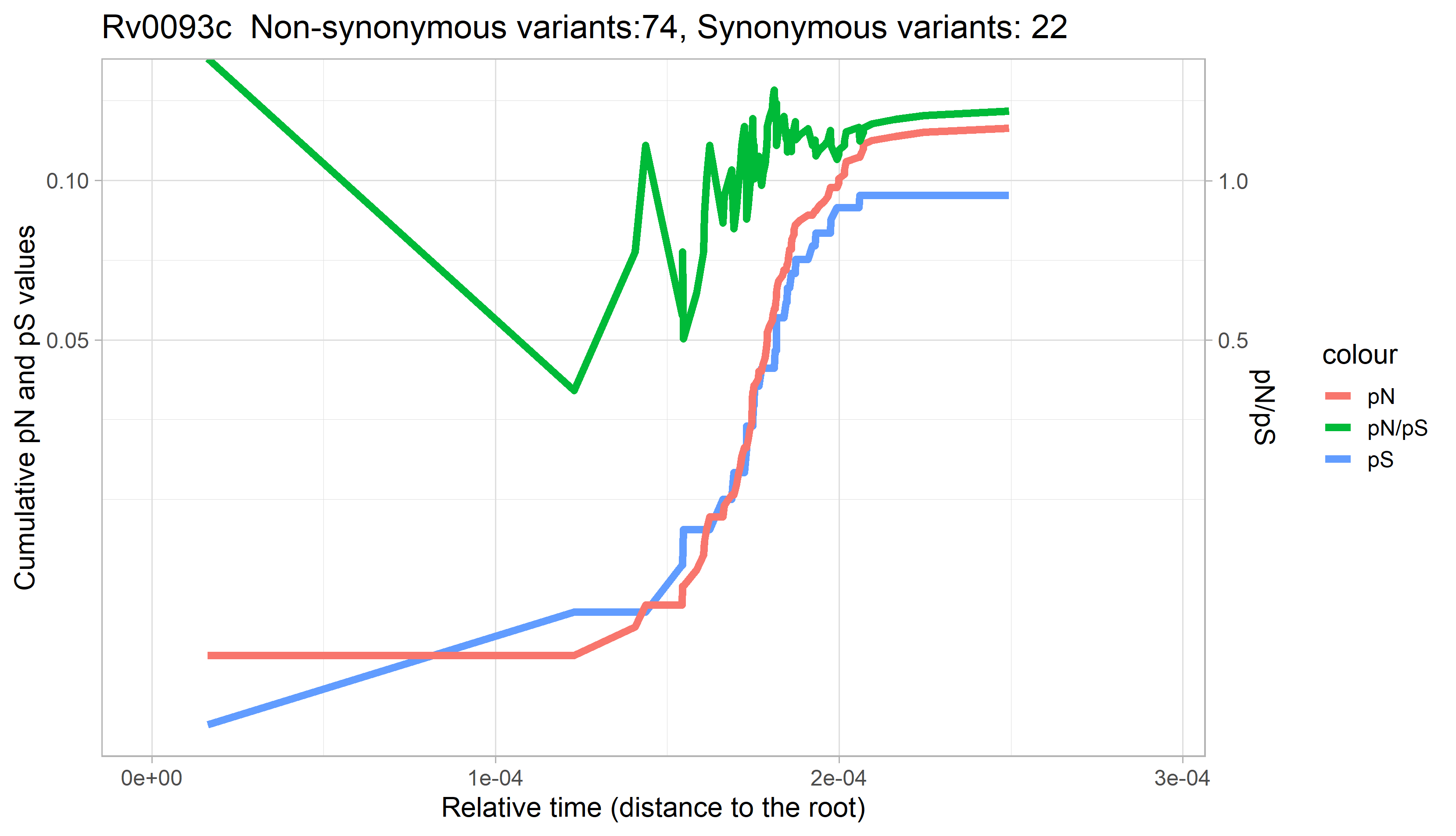

### Rv0097.png

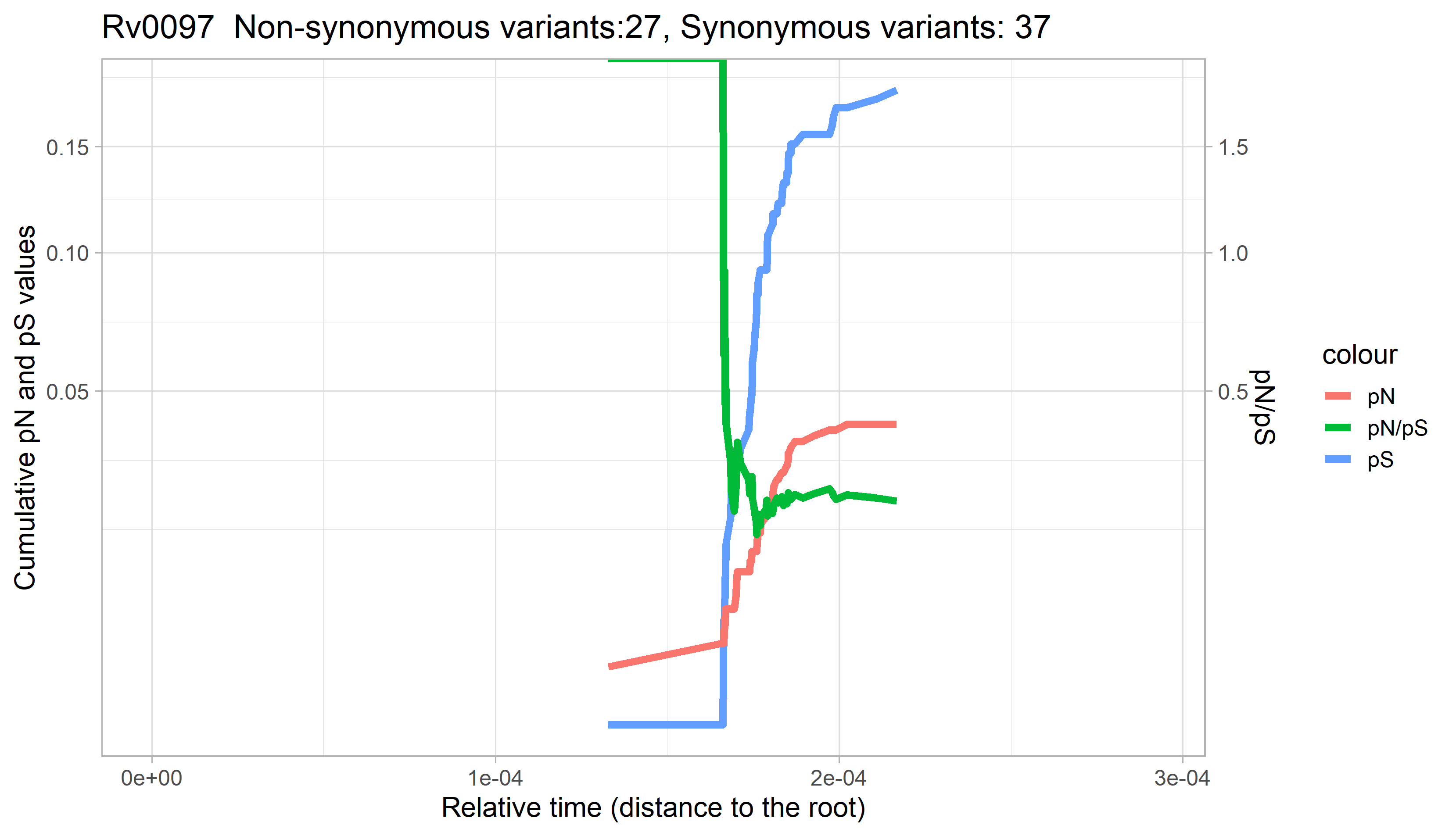

### Rv0098.png

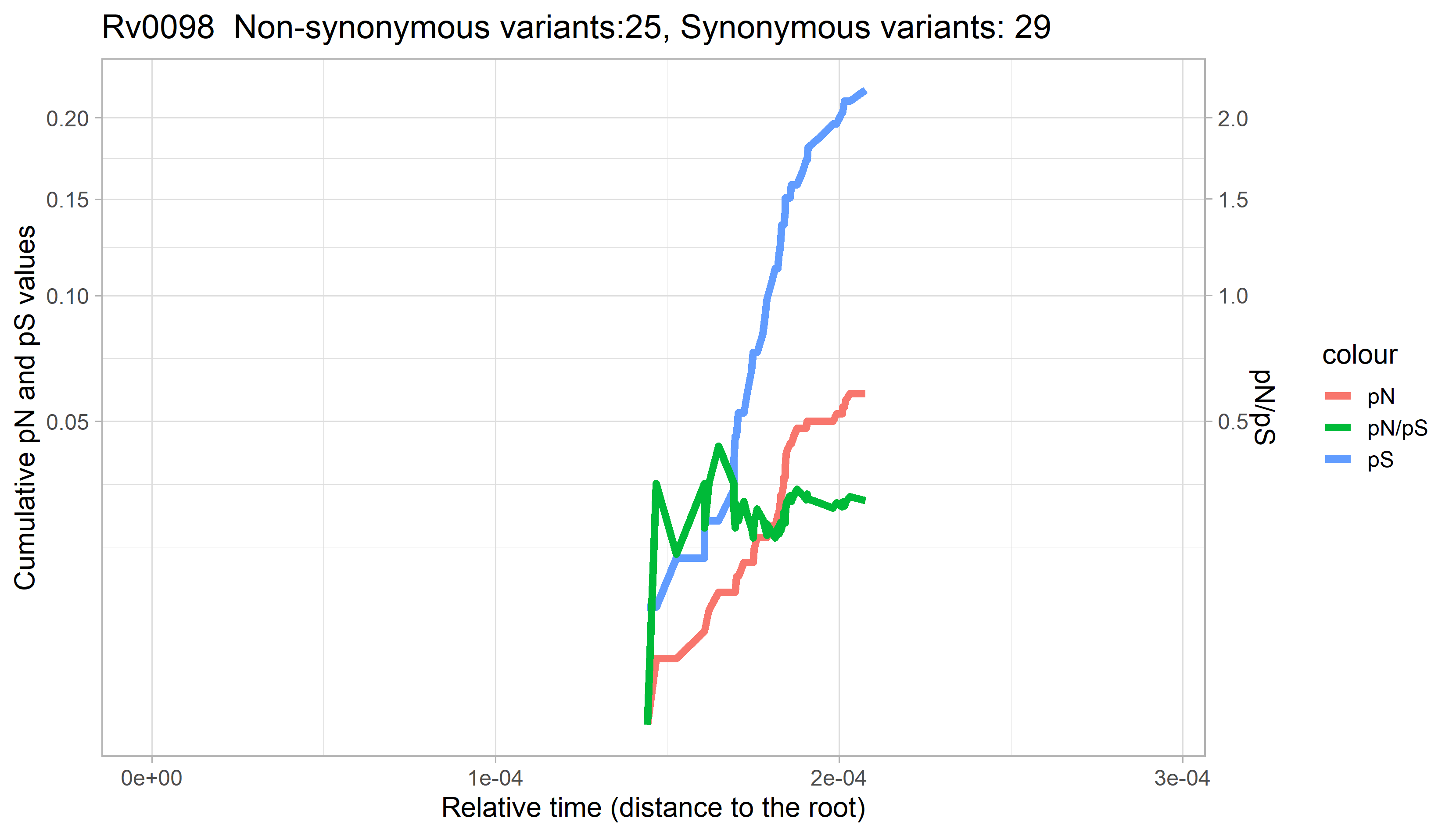

### Rv0099.png

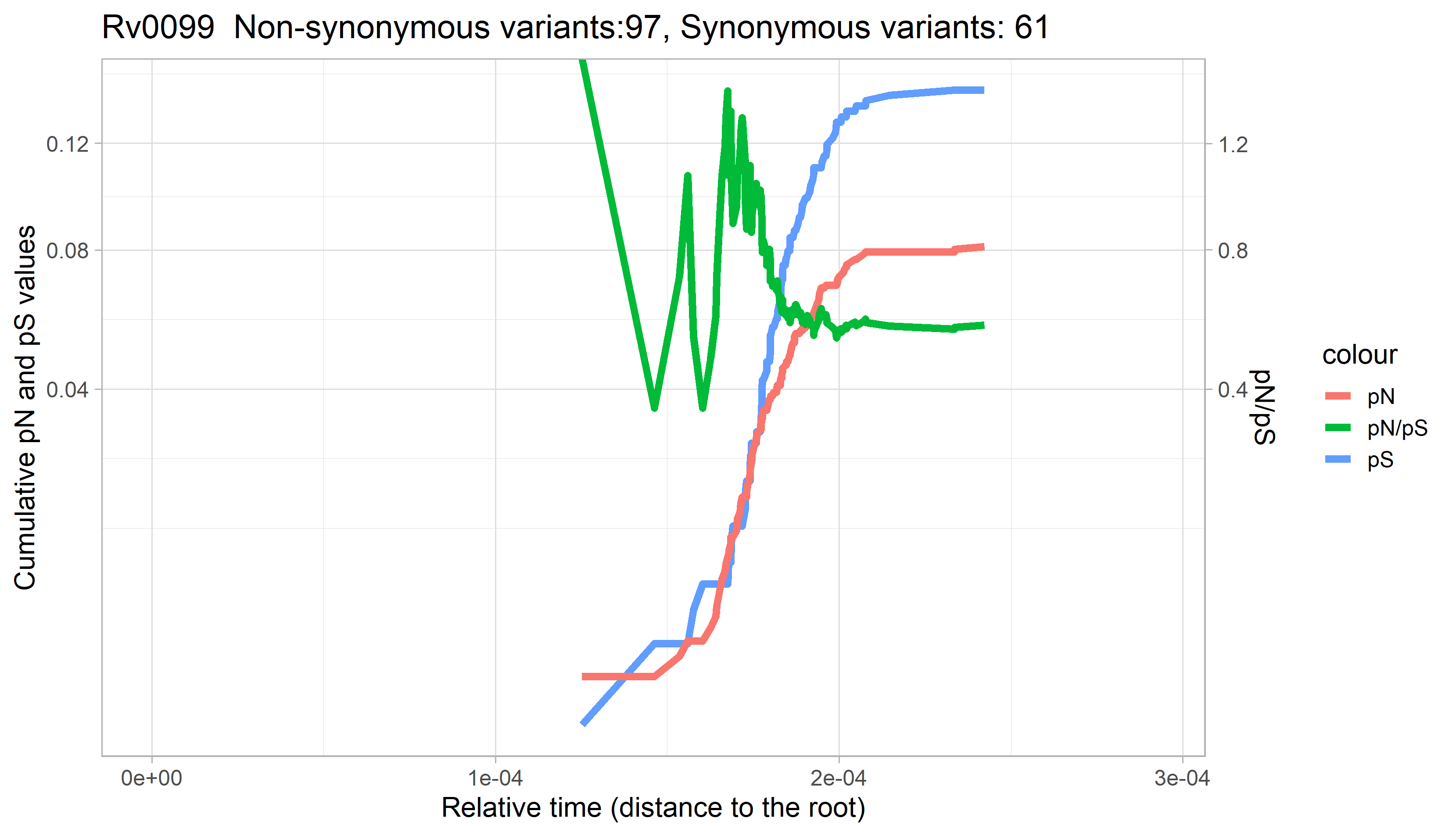

### Rv0100.png

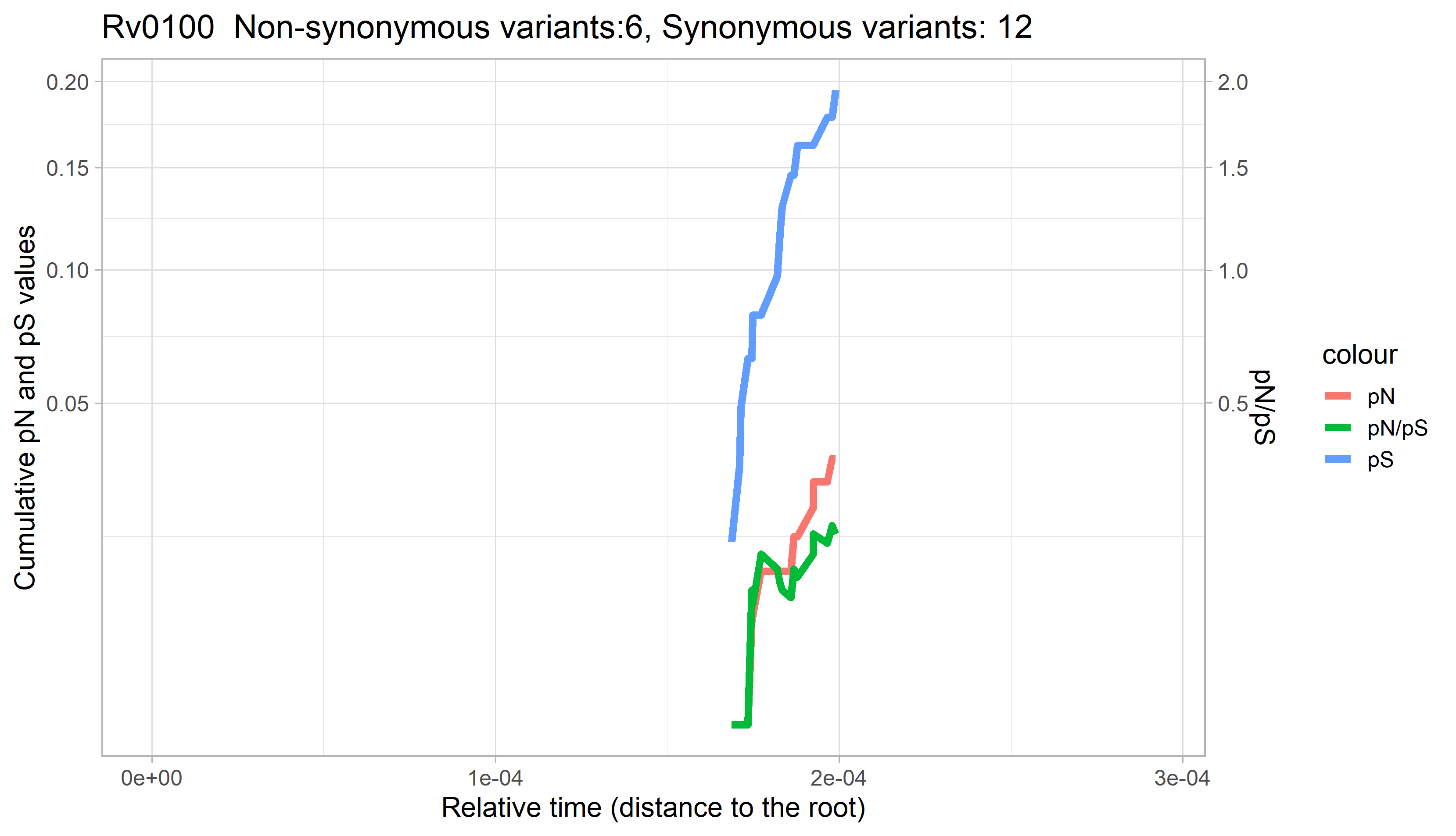

### Rv0101.png

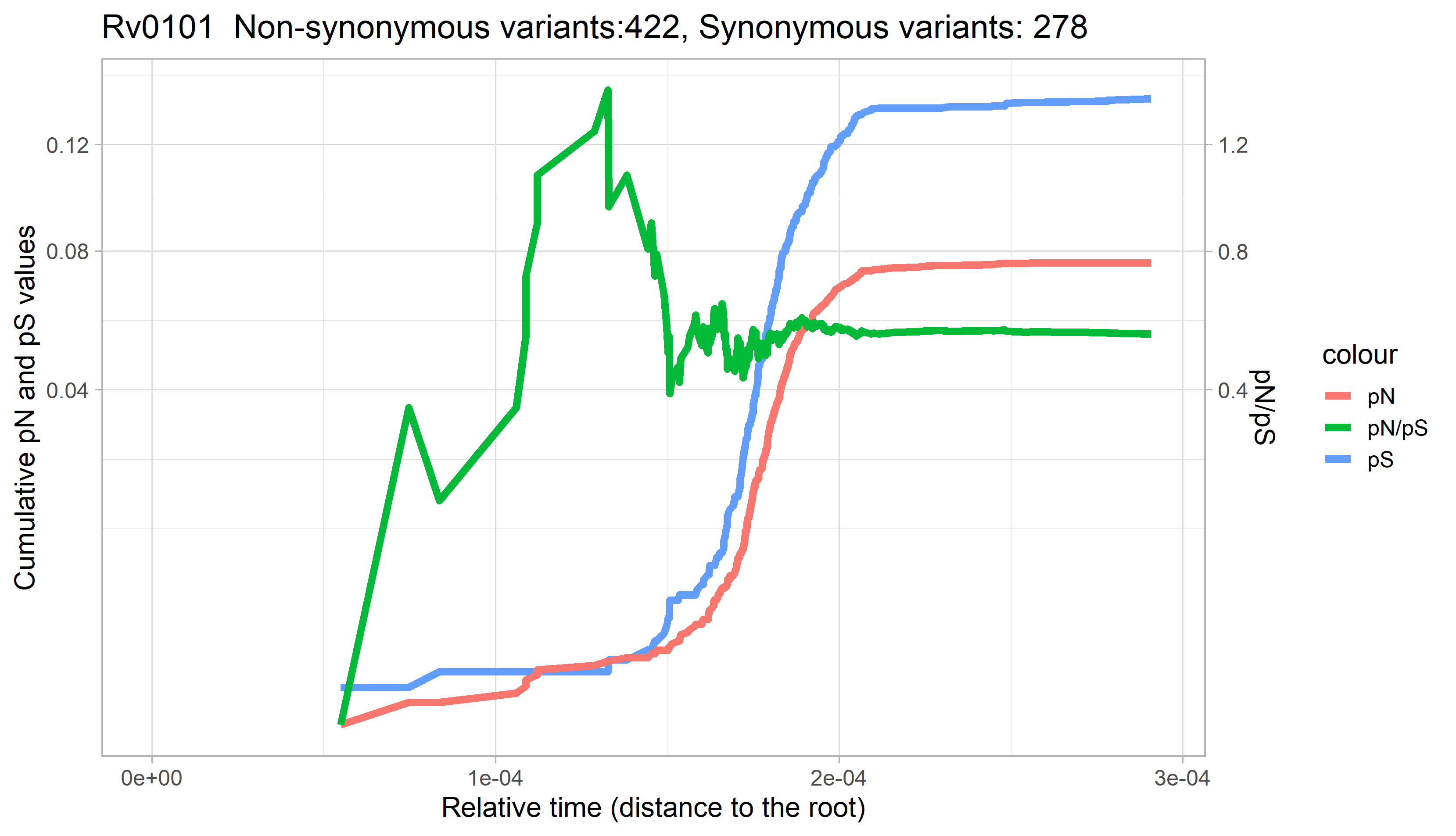

### Rv0102.png

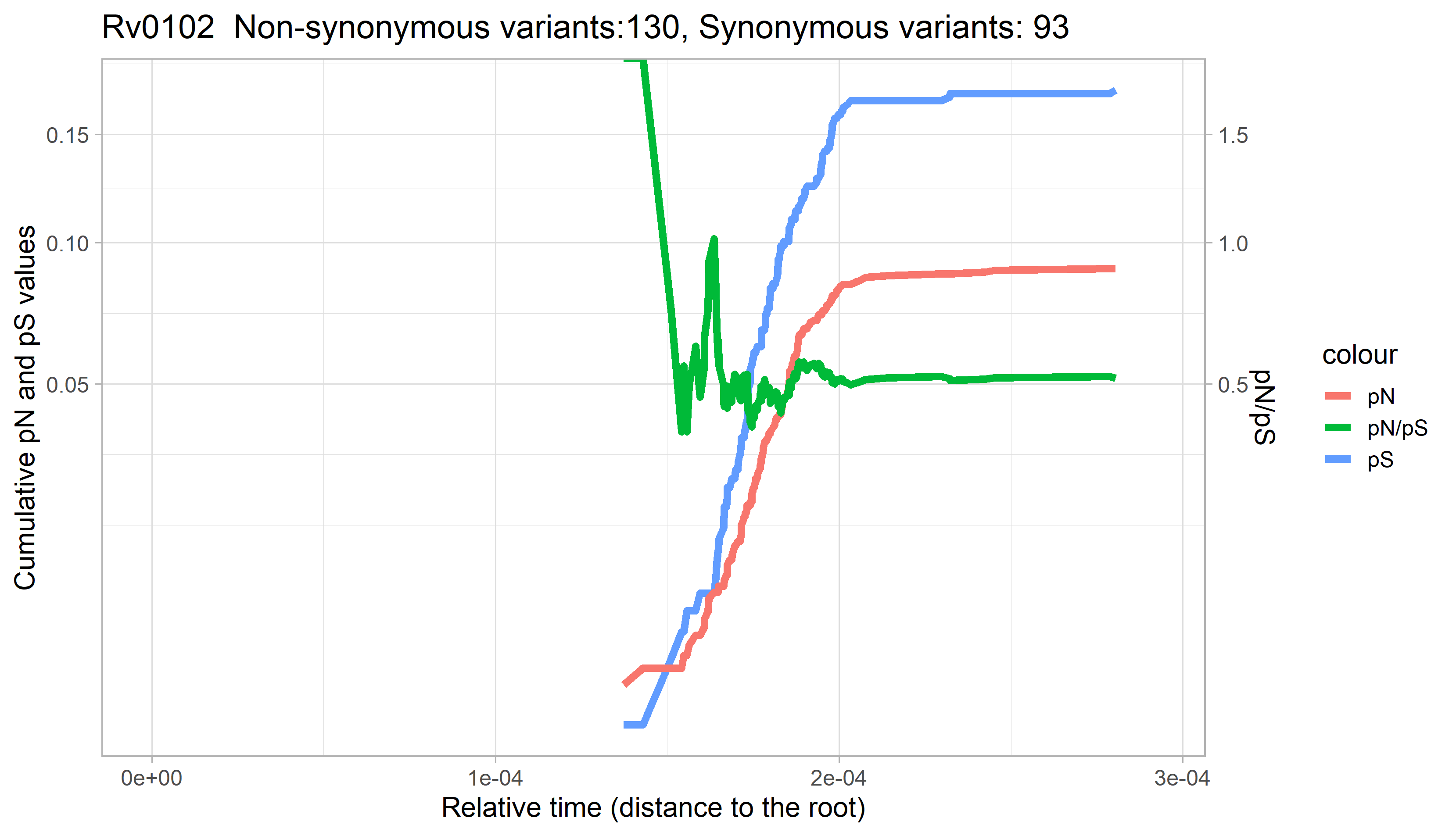

### Rv0103c.png

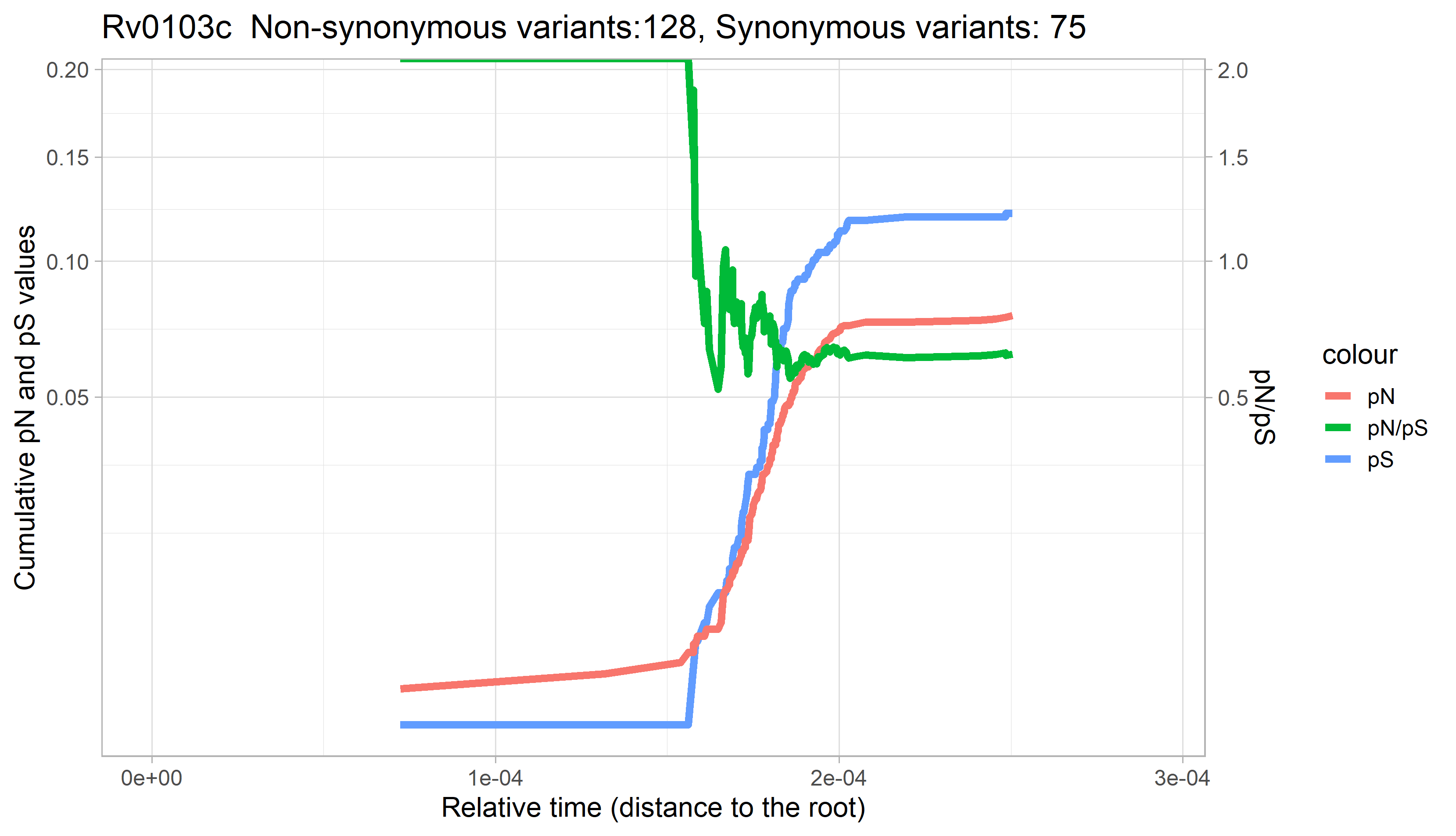
